# BRD4 Prevents R-Loop Formation and Transcription-Replication Conflicts by Ensuring Efficient Transcription Elongation

**DOI:** 10.1101/854737

**Authors:** Drake Edwards, Rohin Maganti, Jarred P. Tanksley, Jie Luo, James J.H. Park, Elena Balkanska-Sinclair, Jinjie Ling, Scott R. Floyd

**Affiliations:** Medical Scientist Training Program, Duke University School of Medicine, Durham, North Carolina 27710, USA; Department of Pharmacology and Cancer Biology, Duke University Medical Center, Durham, North Carolina 27710, USA; Department of Radiation Oncology, Duke University School of Medicine, Durham, North Carolina 27710, USA; Duke University, Durham, North Carolina 27710, USA

## Abstract

Effective spatio-temporal control of transcription and replication during S-phase is paramount to maintaining genomic integrity and cell survival. Dysregulation of these systems can lead to conflicts between the transcription and replication machinery causing DNA damage and cell death. BRD4, a BET bromodomain protein and known transcriptional regulator, interacts with P-TEFb to ensure efficient transcriptional elongation by stimulating phosphorylation of RNA Polymerase II (RNAPII). Here we report that disruption of BET bromodomain protein function causes RNAPII pausing on the chromatin and DNA damage affecting cells in S-phase. We find that this persistent, RNAPII-dependent pausing leads to accumulation of RNA:DNA hybrids (R-loops), which are known to lead to transcription-replication conflicts (TRCs), DNA damage, and cell death. Furthermore, we show that resolution of R-loops abrogates BET bromodomain inhibitor-induced DNA damage, and that BET bromodomain inhibition induces both R-loops and DNA damage at sites of BRD4 occupancy. Finally, we see that the BRD4 C-terminal domain, which interacts with P-TEFb, is required to prevent R-loop formation and DNA damage caused by BET bromodomain inhibition and that oncogenes which promote transcription and replication exacerbate BET bromodomain inhibitor-induced DNA damage. Together, these findings demonstrate that BET bromodomain inhibitors can damage DNA via induction of R-loops and TRCs in highly replicative cells.

## INTRODUCTION

Maintaining the integrity of the genome throughout the cell cycle is paramount to cell survival(Hanahan and Weinberg, 2011), and therefore complex systems have evolved to tackle various threats to the genome’s integrity(Blackford and Jackson, 2017; Cimprich and Cortez, 2008; Hamperl and Cimprich, 2016). During S-phase, areas of chromatin that are engaged with generating RNA transcripts must be coordinated with migrating replication forks. Disruption of either transcription or replication control and coordination can lead to the desynchronization of these chromatin-based activities, resulting in transcription-replication conflicts (TRCs) and subsequent replication stress, DNA damage, and cell death(Aguilera and Gómez-González, 2017; Gaillard and Aguilera, 2016; Garcia-Muse and Aguilera, 2016; Hage et al., 2010; Sollier and Cimprich, 2015). To avoid these collisions, these processes are separated in both time and space through the activity of several known chromatin-based complexes(Hamperl and Cimprich, 2016). Specifically, the processivity of both the replication machinery and the nascent RNA strand are paramount in preventing collisions between the two (Schwab et al., 2015; Zeman and Cimprich, 2014). These systems are an active area of study, especially in cancer cells, as many amplified transcription programs and more frequent replication distinguish cancer cells from normal cells(Kotsantis et al., 2016a; Stork et al., 2016). The strategies that cancer cells employ to avoid TRCs are therefore of potential therapeutic interest, as the components of these TRC avoidance mechanisms could be targeted with wide therapeutic window in variety of cancers.

One source of TRCs is the aberrant formation of RNA:DNA hybrids (R-loops), caused by nascent RNA re-annealing with its DNA template strand forming a three-stranded structure (Aguilera and Gómez-González, 2017; Costantino and Koshland, 2018; Crossley et al., 2019; Garcia-Muse and Aguilera, 2019; Hamperl et al., 2017; Hamperl and Cimprich, 2016; Richard and Manley, 2016; Santos-Pereira and Aguilera, 2015; Sollier and Cimprich, 2015). R-loops play various physiological roles, including Ig class-switching, CRISPR-Cas9 bacterial defense systems, and normal transcription regulation(Chaudhuri and Alt, 2004; Garcia-Muse and Aguilera, 2019; Shao and Zeitlinger, 2017a; Skourti-Stathaki and Proudfoot, 2014; Stuckey et al., 2015; Xiao et al., 2017). However, pathologic R-loops can also form from dysregulated transcription, and these pathologic R-loops can impede the progression of the transcription bubble(Crossley et al., 2019). In the case where RNAPII is stalled, the nascent RNA is allowed to re-anneal with its template strand and form a stable R-loop leading to tethering of RNAPII to the chromatin. During S-phase, these R-loop-tethered transcription bubbles create a roadblock for replication forks(Gan et al., 2011; Matos et al., 2019). If these roadblocks are not resolved, collisions with the replication machinery will lead to replication fork breakdown and DNA strand breaks. Important factors have been identified that prevent and resolve R-loops, including the RNAPII activator CDK9 and the RNA:DNA hybrid endonuclease RNAse H1(L. Chen et al., 2017; Grunseich et al., 2018; Matos et al., 2019; Morales et al., 2016; Nguyen et al., 2017; Parajuli et al., 2017; Shivji et al., 2018; Skourti-Stathaki et al., 2011; Wahba et al., 2011; Wessel et al., 2019a; Zatreanu et al., 2019).

BRD4, a member of the bromodomain and extra-terminal domain (BET) protein family, is a known regulator of transcription elongation. Through its C-terminal domain (CTD) it is known to activate CDK9, the RNAPII-phosphorylating component of the positive transcription elongation factor, P-TEFb(R. Chen et al., 2014; Itzen et al., 2014; Jang et al., 2005; Kanno et al., 2014; Liu et al., 2013; Patel et al., 2013; Rahman et al., 2011; Winter et al., 2017; W. Zhang et al., 2012). After RNAPII has initiated transcription and paused, at many genomic loci, BRD4 releases P-TEFb from its inhibitory complex and allows CDK9 to phosphorylate the second serine of the YSPTSPS repeat on the tail of RNAPII (RNAPIIpS2). Once this phosphorylation event occurs, RNAPII is able to enter the elongation phase of transcription. Consequently, inhibition of BRD4 function reduces transcription of many transcripts(Delmore et al., 2011; Filippakopoulos et al., 2010a; Muhar et al., 2018; Winter et al., 2017).

BET family inhibitors have shown activity in pre-clinical models of several cancers, and clinical trials have shown efficacy, yet mechanisms of action and predictive biomarkers remain elusive. Recently, members of the BET bromodomain family have been implicated in both replication stress, and R-loop biology(Bowry et al., 2018; Kim et al., 2019; Wessel et al., 2019a). In an effort to illuminate the role BRD4 plays in preventing cancer cell death, we studied how the DNA damage repair systems react to BET inhibition. We see that BET inhibitors cause double strand breaks in cells undergoing S-phase replication. Furthermore, we see that overexpression of full-length BRD4 rescues the effects of BRD4 loss, but rescue fails when BRD4 is truncated to delete the P-TEFb-interacting C-terminal domain (CTD). Finally, we see that BET inhibitors cause an RNAPII-dependent increase in the formation of R-loops, and that overexpression of RNAse H1, an endonuclease that acts on the RNA strand of R-loops, reverses BET inhibitor-induced DNA damage. These data suggest a new role for BRD4 in preventing aberrant R-loop formation and TRCs by ensuring efficient RNAPII transcription.

## RESULTS

### Inhibition or degradation of BET family proteins leads to spontaneous DNA damage in cancer cells

BRD4, through its two N-terminal bromodomains, interacts with the chromatin by binding to acetylated histones(Filippakopoulos et al., 2012). In previous work, we have described how a low abundance isoform of BRD4 (Isoform B) mediated chromatin dynamics and DNA damage signaling in the presence of radiation(Floyd et al., 2013a). However, small molecule BET bromodomain protein inhibitors are effective against cancer cells in the absence of radiation(Asangani et al., 2014; Dawson et al., 2011; Rathert et al., 2015; Zuber et al., 2011). Several groups have reported variable effects of BET bromodomain inhibitors on DNA damage signaling (Bowry et al., 2018; Floyd et al., 2013b; Kim et al., 2019; Pericole et al., 2019; Schröder et al., 2012; Sun et al., 2018; J. Zhang et al., 2018). We therefore sought to understand the DNA damage consequences of BET bromodomain inhibition. JQ1, a small molecule inhibitor of BET family proteins, binds to the bromodomains and competitively prevents BRD4 from interacting with chromatin(Filippakopoulos et al., 2010a). In order to test whether JQ1 was able to induce a DNA damage response, we treated HeLa and HCT-116 cells with (500 nM) JQ1 for 16 hours and stained for γH2AX foci, a marker of DNA damage(Rogakou et al., 1998). We observed that JQ1 was able to induce γH2AX foci formation, indicating that BET proteins can prevent spontaneous DNA damage (**Figure 1A**; **Figure 1B**; **Figure S1A; Figure S1B)**. In addition to JQ1, other small molecules that have been used in clinical trials or target a specific bromodomain also led to an increase in γH2AX signaling in HeLa cells at clinically relevant doses (Faivre et al., 2020; Odore et al., 2015; Ozer et al., 2018; Piha-Paul et al., 2019)(**Figure 1H**).

**Figure 1:**
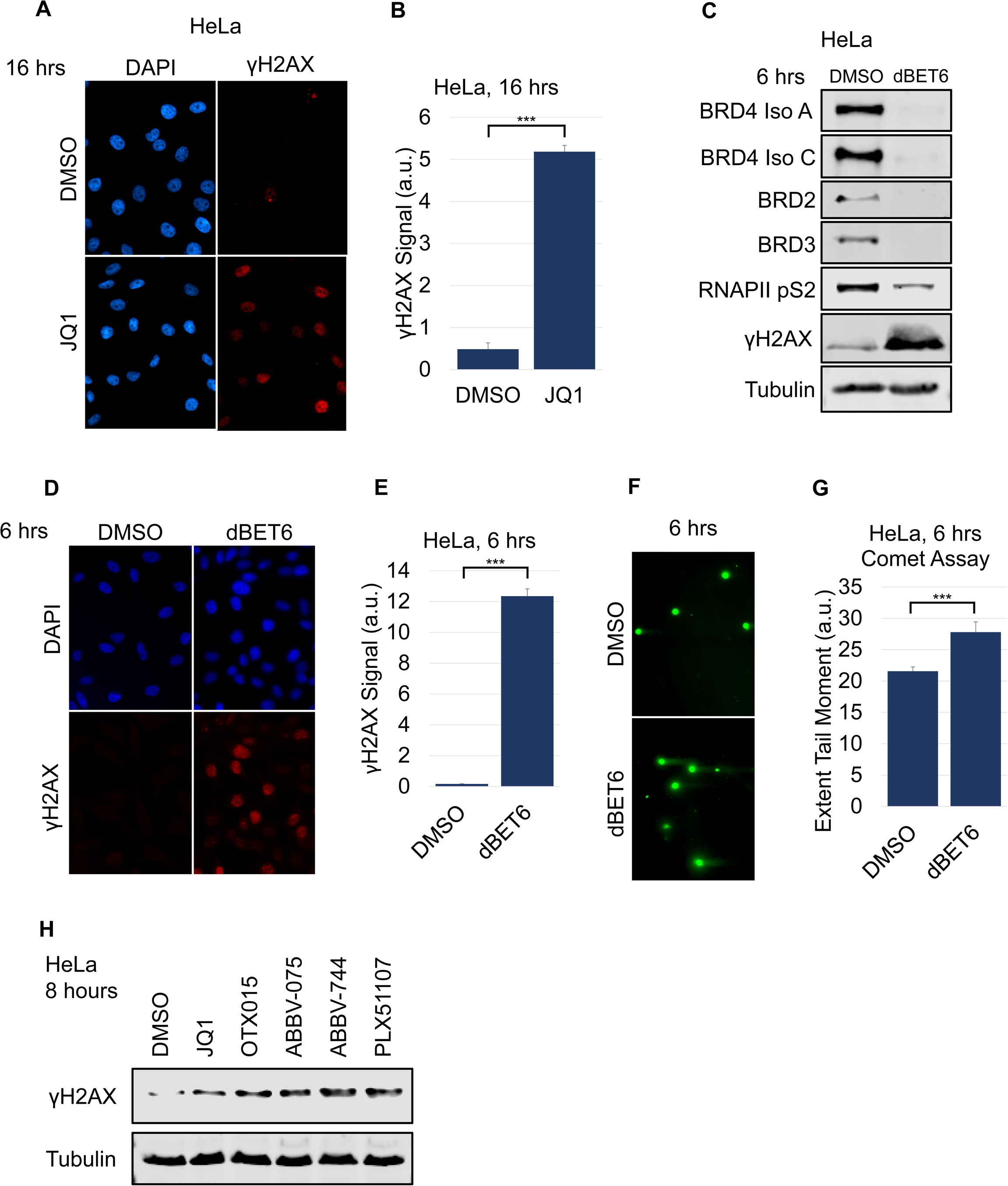
BET protein loss of function leads to spontaneous DNA damage. **A.** Representative images and **B.** quantification of γH2AX staining per nucleus in HeLa cells treated with DMSO or 500 nM JQ1 for 16 hours (>100 cells). **C.** Representative western blots from HeLa cells treated with DMSO or 100 nM dBET6 for 6 hours before harvest. **D.** Representative images and **E.** quantification of γH2AX staining per nucleus in HeLa cells treated with DMSO or 100 nM dBET6 for 6 hours. **F.** Representative images and **G.** quantification of neutral single cell electrophoresis assay of HeLa cells treated with DMSO or 100 nM dBET6 for 6 hours. **H.** Representative western blots from HeLa cells treated with various BET inhibitors for 8 hours before harvest. For western blots, lysates are probed for the epitope indicated beside each panel. Student’s t-test (two-tailed, unpaired) was performed on **B, E,** and **G**. Data represent the mean ±SEM. \**P* < 0.05; \*\**P* < 0.01; \*\*\**P* < 0.001. Source data are provided as a Source Data file.

Recently, a small molecule, dBET6, has been shown to cause rapid degradation of BET proteins(Winter et al., 2017). dBET6, as with other PROTAC molecules, links to an E3-ligase recruiter which causes ubiquitination and subsequent, rapid degradation of target proteins. Advantages of dBET6 are that it allows for the visualization of BET protein loss and acts as a more potent BET protein inhibitor with rapid kinetics. We observed that dBET6 elicited a robust DNA damage response detectable by western blot in HeLa cells at 100 nM concentration in as few as 6 hours. Concurrent with dBET6-induced loss of BET proteins, we observed a reduction in RNA Polymerase II phospho-Serine 2 and saw γH2AX signaling both by western blot and immunofluorescence (**Figure 1C**; **Figure 1D**; **Figure 1E**). These observations confirmed that loss of BET proteins can result in increased DNA damage signaling.

While γH2AX is a general marker for DNA damage signaling, we wanted to establish whether BET protein loss also leads to an increase in physical DNA damage such as double strand breaks. We therefore employed single cell electrophoresis (comet assay) to measure the amount of DNA double strand breaks after dBET6 treatment. Interestingly, we found that in addition to the DNA damage signaling increase, dBET6 increased the number of DNA double strand breaks (**Figure 1F**; **Figure 1G**). These observations indicate that loss of the BET family of proteins can cause physical DNA damage as well as a robust DNA damage response.

### BET protein loss induces DNA damage during S-phase

TRCs, by definition, occur while the cell is actively replicating its genome during S-phase. An active replication fork, when it collides with a transcription bubble in the head-on orientation, leads to fork stalling, DNA damage, and cell death(Hamperl et al., 2017). While probing for DNA damage following BET protein loss in immunofluorescence microscopy studies, we noticed heterogeneity in which cells would display γH2AX foci following dBET6 exposure. Prior work from other groups showed that BRD4 loss leads to a loss of S-phase cells(Maruyama et al., 2002a). While this has been described as a G1/S phase arrest, we decided to test whether actively replicating S-phase cells could be prone to DNA damage after BRD4 loss.

To test whether BET protein loss leads to DNA damage preferentially in actively replicating cells, we labeled HeLa cells with EdU to monitor actively replicating cells while simultaneously treating with dBET6 for two hours. Accordingly, we observed that γH2AX foci formed only in the cells that were labeled with EdU by immunofluorescence (**Figure 2A**; **Figure 2B**). We also labeled OCI-AML2 cells, another JQ1 sensitive cell line(Fiskus et al., 2014; Zhou et al., 2018), and also saw that EdU positive cells showed the most DNA damage following dBET6 treatment **(Figure S2A; Figure S2B)**. These data indicate that BET protein loss is specifically leading to DNA damage in cells that are actively replicating in S-phase.

**Figure 2:**
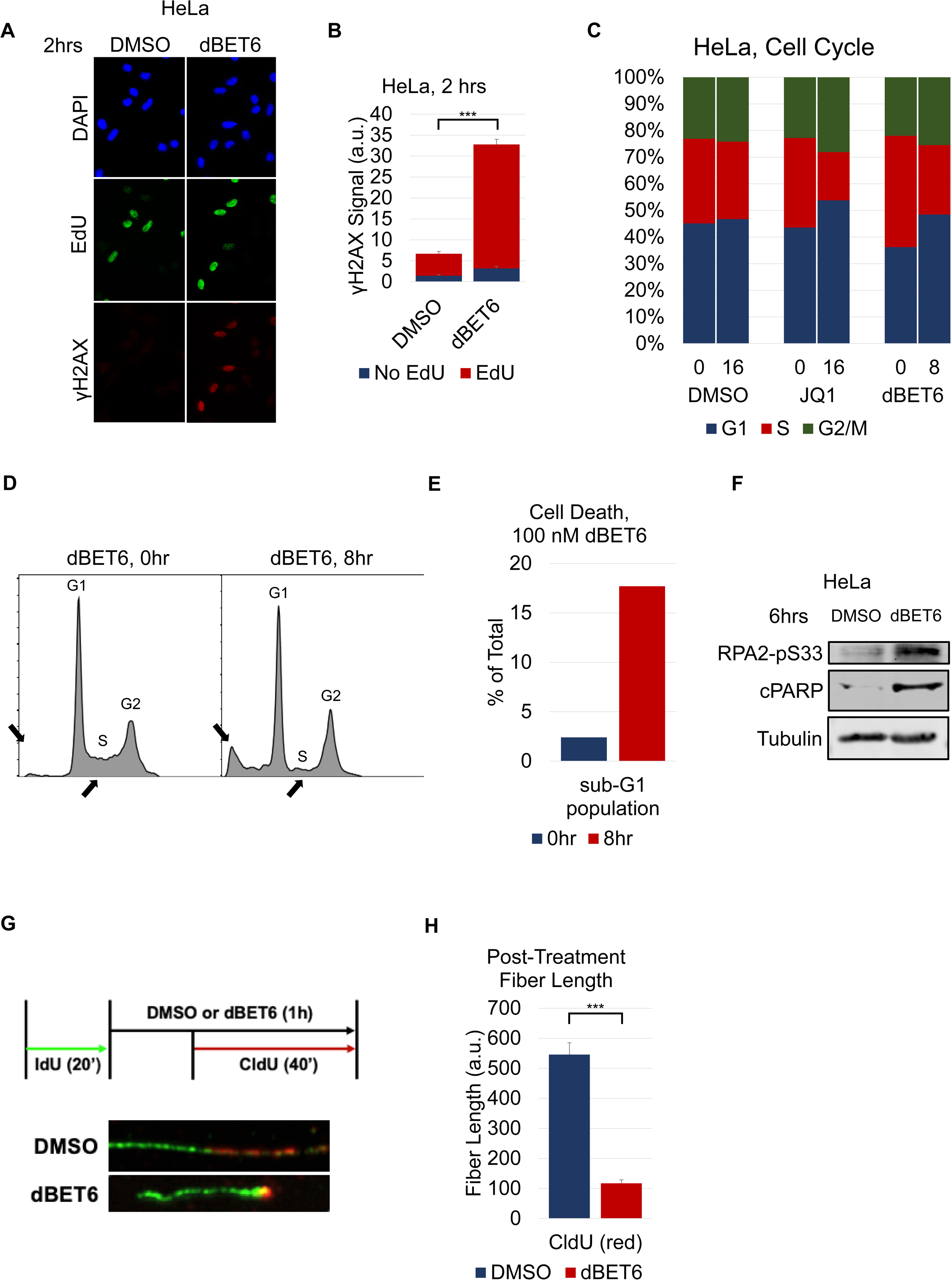
BET protein degradation leads to replication stress and S-phase-dependent DNA damage. **A.** Representative images and **B.** quantification of γH2AX staining per nucleus in HeLa cells treated simultaneously with 100 nM dBET6 and 10 µM EdU for 2 hours. **C.** Cell cycle analysis of HeLa cells treated with DMSO, 500 nM JQ1, or 100 nM dBET6 for times as shown. **D.** Histogram and **E.** quantification of sub-G1 populations of HeLa cells before and after treatment with 100 nM dBET6. **F.** Representative western blot images of lysates from HeLa cells treated with DMSO or 100 nM dBET6 for 6 hours probed for the epitope indicated beside each panel. **G.** Representative images and **H.** quantification of DNA fiber analysis of HeLa cells treated with DMSO or 100 nM dBET6. Cells in **C.**, **D.**, and **E.** were fixed after treatment, stained with PI, and quantified for DNA content using flow cytometry. Student’s t-test (two-tailed, unpaired) was performed on **B. and H.** Data represent the mean ±SEM. \**P* < 0.05; \*\**P* < 0.01; \*\*\**P* < 0.001. Source data are provided as a Source Data file.

To determine whether the S-phase-specific DNA damage following BET LOF was associated with cell death, we performed cell cycle analysis of HeLa cells treated with JQ1 or dBET6. Previous work has shown that apoptotic cells display as a broad hypodiploid (sub-G_1_) peak(Riccardi and Nicoletti, 2006). Interestingly, following BET LOF, we observed a decrease in the S-phase population of cells and a corresponding increase in the sub-G_1_ population (**Figure 2C**; **Figure 2D**; **Figure 2E**). These cell cycle changes indicate that BET LOF lead to cell death of cells in S-phase.

These observations also correlated with replication stress and apoptotic signaling. RPA2, a downstream target of the replication stress master kinase ATR, is known to be phosphorylated on Serine 33 (RPA2-pS33) by ATR in response to replication stress(Olson et al., 2006). BET inhibition with dBET6 caused a robust increase in RPA2-pS33 (**Figure 2F**), indicating that BET inhibition causes replication stress, and providing further evidence that BET protein loss leads to S-phase dependent damage. Furthermore, in dBET6-treated cells, we saw increased levels of cleaved Poly(ADP-ribose) (cPARP) indicating that this S-phase damage was not effectively repaired, and caused cell death (**Figure 2F**).

Finally, to understand whether BET LOF led to dysregulation of replication, we used DNA fiber analysis to observe the progression of replication forks following treatment of dBET6. Interestingly, we found that treatment with dBET6 led to a significant decrease in CldU incorporation, indicating that BET LOF leads to a decrease in replication fork progression (**Figure 2G**; **Figure 2H**). Taken together, these data demonstrate that BET LOF leads to an increase in replication stress and cell death in actively replicating cells.

### The C-Terminal Domain of BRD4 is necessary to prevent DNA damage caused by BET protein loss

The BET protein family consists of four members: BRD2, BRD3, BRD4, and BRDT (of note, BRDT is expressed mainly in the testes)(Pivot-Pajot et al., 2003). Inhibitors of this family of proteins, namely JQ1 and the degrader dBET6, function by binding to the bromodomains which are shared by all members. Thus, it is important to elucidate how the various BET bromodomain proteins contribute to the DNA damage seen by dBET6 treatment. To test this, we used siRNA to knock down BRD2, BRD3, and BRD4 and measured γH2AX signaling (**Figure 3A**; **Figure S3A**). After 72 hours of knock down, we observed that both BRD2 and BRD4 loss led to increased γH2AX signaling, similar to recent reports(Kim et al., 2019). Owing to a wealth of studies that established mechanisms of BRD4 in transcription regulation, and earlier work showing replication dysfunction caused by BRD4 loss(Bisgrove et al., 2007; Maruyama et al., 2002b; Wessel et al., 2019a; Winter et al., 2017), we focused on the role of BRD4 in the prevention of S-phase DNA damage.

**Figure 3:**
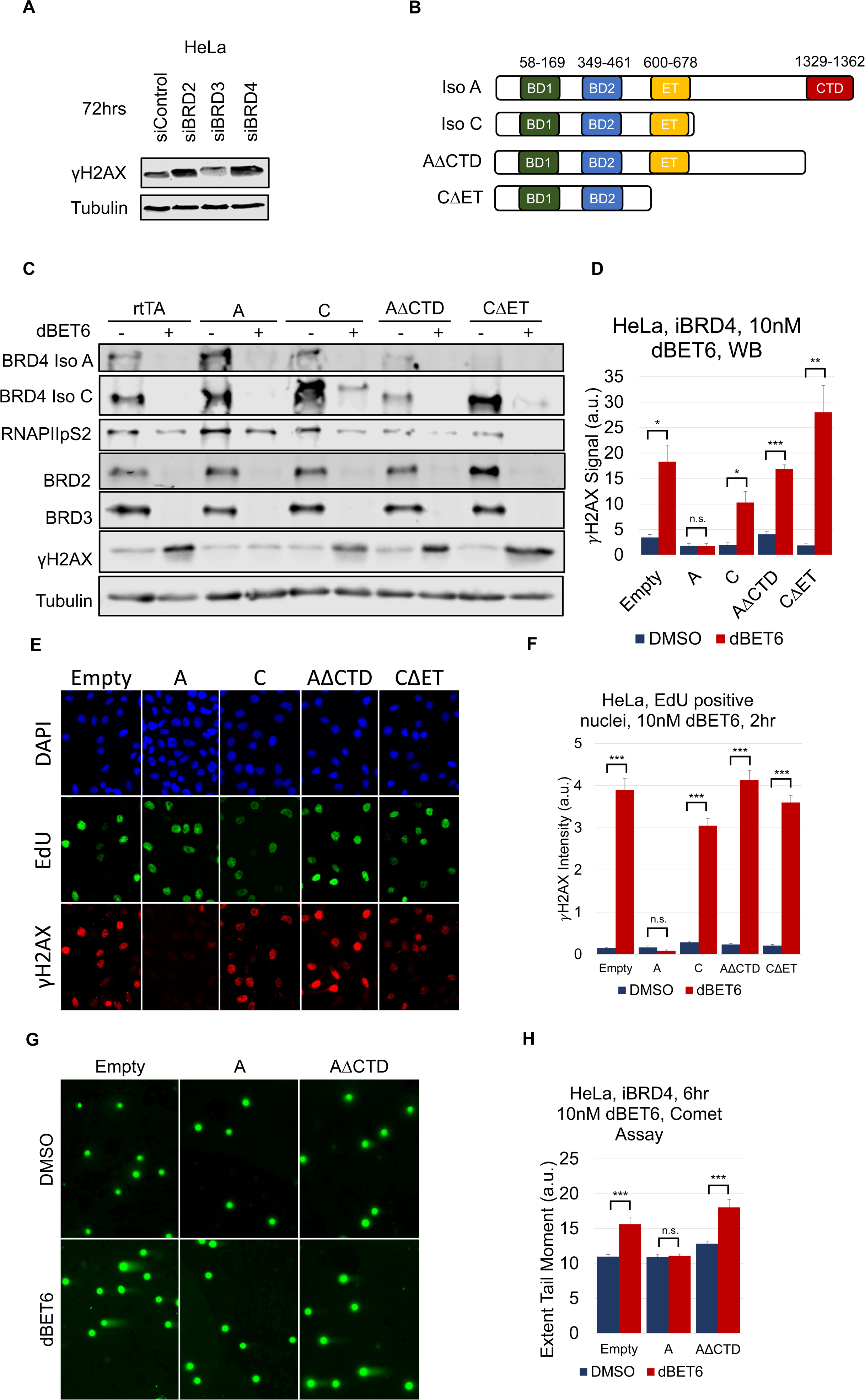
The C-terminal domain of BRD4 is required to prevent TRCs. **A.** Representative western blots of HeLa cells treated with siControl, siBRD2, siBRD3, or siBRD4 for 72 hours and probed for the epitope indicated beside each panel. **B.** Domain structure of overexpression constructs depicting the location of the bromodomains, extra-terminal domain, and C-terminal domain of BRD4. **C.** Representative images and **D.** quantification of western blots from HeLa cells stably infected with each BRD4 construct and induced with doxycycline for 24 hours before being treated with 10 nM dBET6 for 6 hours and harvested: lysates were probed for the epitope indicated beside each panel. **E.** Representative images and **F.** quantification of γH2AX staining per nucleus in EdU-positive HeLa cells induced as in **D** and then simultaneously treated with 10 nM dBET6 and 10 µM EdU for 2 hours. **G.** Representative images and **H.** quantification of neutral single cell electrophoresis assay of HeLa cells induced as in **D** followed by treatment with DMSO or 10 nM dBET6 for 6 hours. Student’s t-test (two-tailed, unpaired) was performed on **D, F,** and **H**. Data represent the mean ±SEM. \**P* < 0.05; \*\**P* < 0.01; \*\*\**P* < 0.001. Source data are provided as a Source Data file.

The full-length isoform of BRD4, isoform A, contains several known domains, including two bromodomains, an extra-terminal domain, and a C-terminal domain (**Figure 3B**). The two bromodomains, which bind to acetylated lysine on histone tails, and the extra-terminal domain are shared among all BET protein members. The C-terminal domain, however, is unique to BRD4 isoform A and interacts with the P-TEFb complex that contains CDK9, leading to Serine 2 phosphorylation of RNAPII and transcription pause-release(R. Chen et al., 2014; Itzen et al., 2014; Jang et al., 2005; Kanno et al., 2014; Liu et al., 2013; Patel et al., 2013; Rahman et al., 2011; Winter et al., 2017; W. Zhang et al., 2012). Also, previous work showed that CDK9 inhibition leads to paused RNAPII and an increase in R-loops (J.-Y. Chen et al., 2019). Thus, we hypothesized that BRD4 loss could also lead to CDK9 dysfunction, resulting in R-loops and DNA damage. Moreover, we reasoned that the P-TEFb-interacting CTD would be required to prevent TRCs and DNA damage.

To determine the mechanism behind damage caused by BET protein loss, we developed a panel of inducible BRD4 overexpression constructs in order to test their ability to rescue the effects of dBET6 (**Figure 3B**). The panel included two naturally occurring isoforms, A and C. Isoform A being the full length isoform mentioned above, and isoform C as a shorter isoform only including the two bromodomains and the extra-terminal domain(Floyd et al., 2013b) (and lacking the CTD). We also developed a truncated construct of isoform A missing only the CTD (AΔCTD) which has previously shown to interact with CDK9(Bisgrove et al., 2007). Finally, we developed a construct excluding the extra-terminal domain (CΔET). These constructs were used to develop stable cell lines under doxycycline control to overexpress the BRD4 isoforms **(Figure S3B)**.

In order to determine whether BRD4 isoform A (full length isoform) was able to rescue the DNA damage effects caused by dBET6, we induced isoform A expression with doxycycline for 24 hours before treatment with dBET6. We found that isoform A was indeed able to rescue the γH2AX signaling caused by dBET6 (**Figure 3C**; **Figure 3D**; **Figure S3C**). While we saw that isoform A was able to rescue the effects of dBET6 treatment, the protein levels of overexpression construct remaining after dBET6 treatment were difficult to detect by Western blot. To further verify rescue of TRC-induced DNA damage by BRD4 isoform A, we measured BRD4 levels by immunofluorescence staining of dBET6-treated cells that either did, or did not, contain the overexpression construct **(Figure S3D)**. As expected, isoform A was still present after dBET6 treatment only in cells expressing the induced rescue construct, confirming that the rescue of γH2AX was due to isoform A still being present. We also observed that isoform A was able to rescue the loss of RNAPIIpS2, indicating that overexpressing full-length BRD4 was able to ensure efficient transcription elongation even in the presence of dBET6. These data suggest that BRD4 is sufficient in rescuing the effects of dBET6. Next, we applied the same conditions to the entire panel of BRD4 overexpression constructs by western blot (**Figure 3C**; **Figure 3D**). Importantly, none of the other overexpression constructs was able to rescue either the γH2AX signaling or the loss of RNAPIIpS2. Furthermore, we saw that only isoform A was able to rescue the S-phase specific γH2AX foci caused by dBET6 treatment (**Figure 3E**; **Figure 3F**). These observations indicate that the C-terminal domain (CTD) is required to prevent BET inhibitor-induced loss of RNAPIIpS2 and S-phase DNA damage.

Next, we wanted to elucidate whether the CTD of BRD4 was necessary to rescue the DNA double strand breaks caused by dBET6 treatment. To test this, we used a comet assay to quantify the breaks following dBET6 treatment following overexpression of isoform A or AΔCTD. Again, we saw that isoform A, but not AΔCTD, was able to rescue the dBET6-induced DNA double strand breaks. This further indicates that the C-terminal domain of BRD4 is necessary to prevent DNA double strand breaks in S-phase, and points to a mechanism involving both transcription and replication.

### BET inhibition leads to an increase in R-loop-dependent DNA damage

R-loops have been previously shown to cause TRCs and replication stress in cancer(Aguilera and Gómez-González, 2017; Costantino and Koshland, 2018; Crossley et al., 2019; Garcia-Muse and Aguilera, 2019; Hamperl et al., 2017; Hamperl and Cimprich, 2016; Richard and Manley, 2016; Santos-Pereira and Aguilera, 2015; Sollier and Cimprich, 2015). Specifically, an R-loop is able to tether a persistently-paused RNAPII to the chromatin, creating a roadblock for the replication machinery. RNAPII, after initiation of transcription of ∼50 bp, becomes paused until a second phosphorylation event of the second serine on its tail. BRD4, through its C-terminal domain, activates CDK9 to undergo this phosphorylation event and ensure efficient transcription elongation(Bisgrove et al., 2007; R. Chen et al., 2014; Itzen et al., 2014; Jang et al., 2005; Krueger et al., 2010; Patel et al., 2013). Previous work has also shown that loss of BRD4 leads to decreased traveling ratios of RNAPII after dBET6 treatment, indicating that RNAPII is paused on the chromatin(Winter et al., 2017). Furthermore, previous studies have indicated that direct chemical inhibition of CDK9 leads to stalled RNAPII and an increase in R-loop formation(L. Chen et al., 2017; Shao and Zeitlinger, 2017a). Therefore, we hypothesized that loss of BRD4 may also lead to an increase of R-loops, and that those R-loops are responsible for the S-phase damage seen after BRD4 loss.

To determine whether BRD4 loss leads to an increase in R-loop formation, we employed the R-ChIP-seq technique which has previously been described as a way to detect R-loop formation on the chromatin(J.-Y. Chen et al., 2019). R-ChIP employs the use of a catalytically inactive form of the R-loop-specific endonuclease, RNAse H1. The mutation, D210N, allows RNase H1 to bind to, but not resolve, R-loops. The construct is tagged with a V5 peptide, which then allows it to be enriched from crosslinked cells, along with associated chromatin, for ChIP-sequencing **(Figure S4A)**. We performed R-ChIP-seq in dBET6-exposed cells and found dramatic increases in global R-loop formation (**Figure 4A**). Similarly, we saw globally increased γH2AX ChIP signal in dBET6-treated cells. Furthermore, we validated three previously described(Liu et al., 2013) BRD4 occupying loci using R-ChIP-qPCR (**Figure 4B**; **Figure 4C**). Surprisingly, while we saw most of the R-loop formation near the promoter regions, there was also increased R-loop formation throughout the length of the gene. In addition, we also saw a decrease of RNAPIIpS2 along the length of these loci as well and an increase in RNAPII travel ratio which has been reported previously (Winter et al., 2017)**(Figure S4B; Figure S4C)**. This indicates that BRD4 not only prevents pause-release of RNAPII, but also prevents the accumulation of R-loops and RNAPII stalling throughout the length of the gene.

**Figure 4:**
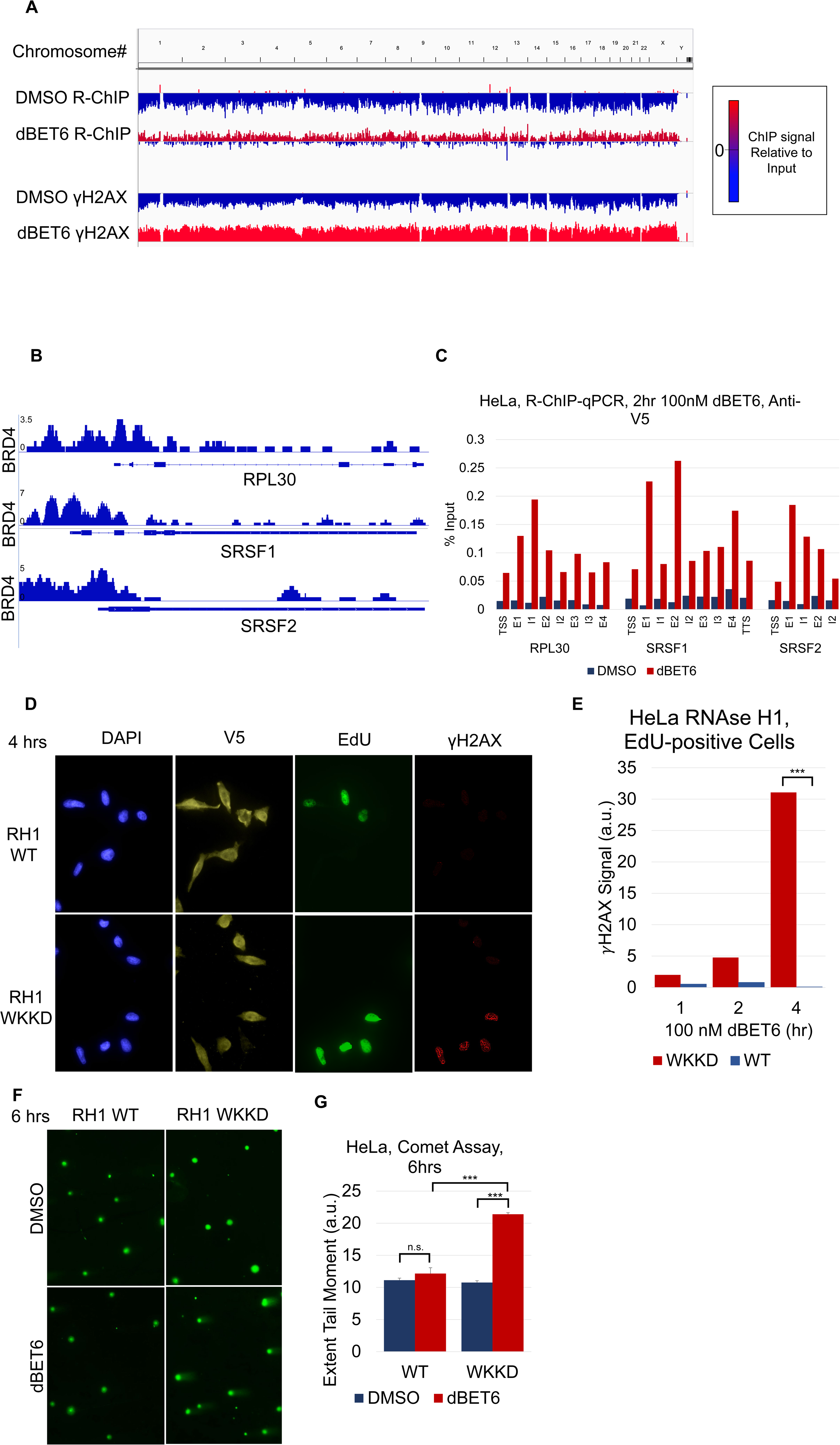
BET inhibition leads to an increase in R-loop-dependent DNA damage **A.** Global ChIP-seq and R-ChIP-seq signal relative to input for HeLa cells treated with DMSO or dBET6 as shown. The right panel depicts how different colors represent the ChIP-seq or R-ChIP-seq signal relative to input. **B.** BRD4 ChIP-seq signal of select loci from ChIP-seq data published in Liu, et al. (2013) (Liu et al., 2013) **C.** Quantification of R-ChIP-qPCR at loci shown in **B** after treatment with DMSO or 100 nM dBET6. **D.** Representative images and **E.** quantification of γH2AX staining per nucleus in HeLa cells transfected with wild-type or WKKD mutant RNAse H1 before being treated with 100 nM dBET6 or 10 µM EdU for 4 hours. **F.** Representative images and **G.** quantification of neutral single cell electrophoresis assay of HeLa cells transfected as in **E** before treatment with DMSO or 100 nM dBET6 for 6 hours. Student’s t-test (two-tailed, unpaired) was performed on **E**. ANOVA was performed on **G**. Data represent the mean ±SEM. \**P* < 0.05; \*\**P* < 0.01; \*\*\**P* < 0.001. Source data are provided as a Source Data file.

We next postulated that the R-loops formed by BRD4 loss could be the root cause of the S-phase DNA damage we observed. To elucidate this, we employed the overexpression of V5-tagged wild-type RNase H1, which is known to be able to resolve R-loops and reverse DNA damage caused by their existence(Matos et al., 2019). As a negative control, we used a V5-tagged RNase H1 mutant, containing mutations at W43A, K59A, K60A and D210N (WKKD), which has been previously described to lack both the catalytic activity as well as the DNA binding activity of RNAse H1(L. Chen et al., 2017). To test whether RNase H1 was able to rescue the S-phase DNA damage caused by BRD4 loss, we overexpressed either the WT RNase H1 or the WKKD mutant construct, treated with dBET6, and stained for V5, EdU, and γH2AX (**Figure 4D**; **Figure 4E**; **Figure S4D**). Consistent with our hypothesis that BET inhibition leads to DNA damage via increased formation of R-loops, over-expression of WT RNAseH1, but not the non-binding WKKD mutant, rescued the DNA damage induced by BRD4 loss in EdU positive cells. We then sought to test whether RNase H1 was able to rescue the DNA double strand breaks caused by dBET6 (**Figure 4F**; **Figure 4G**). Similarly, we observed that RNAse H1 was also able to rescue these DNA double strand breaks. These data indicate that following BRD4 loss, R-loops form and lead to DNA damage in S-phase, likely from TRCs.

As BRD4 plays a regulatory role in the transcription of many genes, we sought to understand whether BRD4 was playing a direct role in preventing R-loop formation, or whether it was indirectly preventing R-loop formation through the transcriptional control of other proteins implicated in R-loop processing. SETX and SRSF1 have both been previously shown to be involved with R-loop processing(Li and Manley, 2005; Sollier et al., 2014). We saw that dBET6 treatment did not impact the level of neither SETX nor SRSF1 in the timeframe when the R-loop-dependent TRCs and DNA damage occurred **(Figure S4E)**.

In order to dissect the mechanism of BET LOF-induced TRCs and DNA damage, we explored whether knock down of other proteins associated with transcription would have an effect on DNA damage caused by BET LOF. HEXIM normally holds CDK9 in an inhibitory complex until activated by BRD4(R. Chen et al., 2014; Krueger et al., 2010). In addition, the nuclear excision factors XPG and XPF have been implicated in the resolution of R-loops in transcription termination(Sollier et al., 2014). Interestingly, knock down of these proteins also did not have an effect on the DNA damage caused by dBET6 **(Figure S4F; Figure S4G)**.

Topoisomerases, which relieve torsional stress produced by the movement of both the replication fork and transcription bubble, are important in preventing replication stress caused by TRCs(Bermejo et al., 2012). Specifically, the activity of Top1 has been implicated in relieving negative supercoiling behind a transcription bubble that can lead to R-loop formation(Drolet et al., 1995; Hage et al., 2010; Massé et al., 1997). In addition, BET inhibition has been previously shown to kill cells synergistically with the topoisomerase I inhibitor, camptothecin(Baranello et al., 2016; Wessel et al., 2019b).

We therefore measured DNA damage after Top1 inhibition alone or in combination with bromodomain degradation. As expected, exposing HeLa cells to either dBET6 or camptothecin alone results in DNA damage, however, the combination showed additive effects, indicating that BRD4 may be causing DNA damage through mechanisms additional to Top1 inhibition **(Figure S4H)**. Indeed, prior work indicates that BRD4-stimulated activation of Top1 proceeds through an N-terminal kinase activity(Baranello et al., 2016). Our data indicates that this N-terminal BRD4 activity is insufficient to rescue the TRC-driven DNA damage that we observe specifically in S-phase (**Figure 3**).

Topoisomerase II has been implicated in the generation of transcription-dependent DNA double strand breaks (Canela et al., 2019; Kim et al., 2019). Therefore, we measured the effect of topoisomerase II inhibition and knock down on BETi-induced DNA damage. In contrast to recently reported findings involving BRD2, we saw that Top2 inhibition with dexrazoxane acted synergistically with dBET6 in HeLa cells to increase DNA damage signaling **(Figure S4H)**. Similarly, we observed that siRNA knock down of Top2α or Top2β in HeLa cells had a small but not significant effect on the DNA damage caused by dBET6 **(Figure S4F; Figure S4G)**. Lastly, generation of transcription-dependent DNA double strand breaks by Top2 has been linked to proteasomal degradation of Top2 (Canela et al., 2019; Kim et al., 2019). However, as shown in **Figure 6A**, co-treatment of OCI-AML2 cells with dBET6 and the proteasome inhibitor MG-132 lead to greatly enhanced DNA damage. Taken together, these findings point to a mechanism of BET loss-induced DNA damage that is distinct from BRD2 effects and involves BRD4 protecting against S-phase dependent damage and TRCs through suppression of R-loop formation.

**Figure 5:**
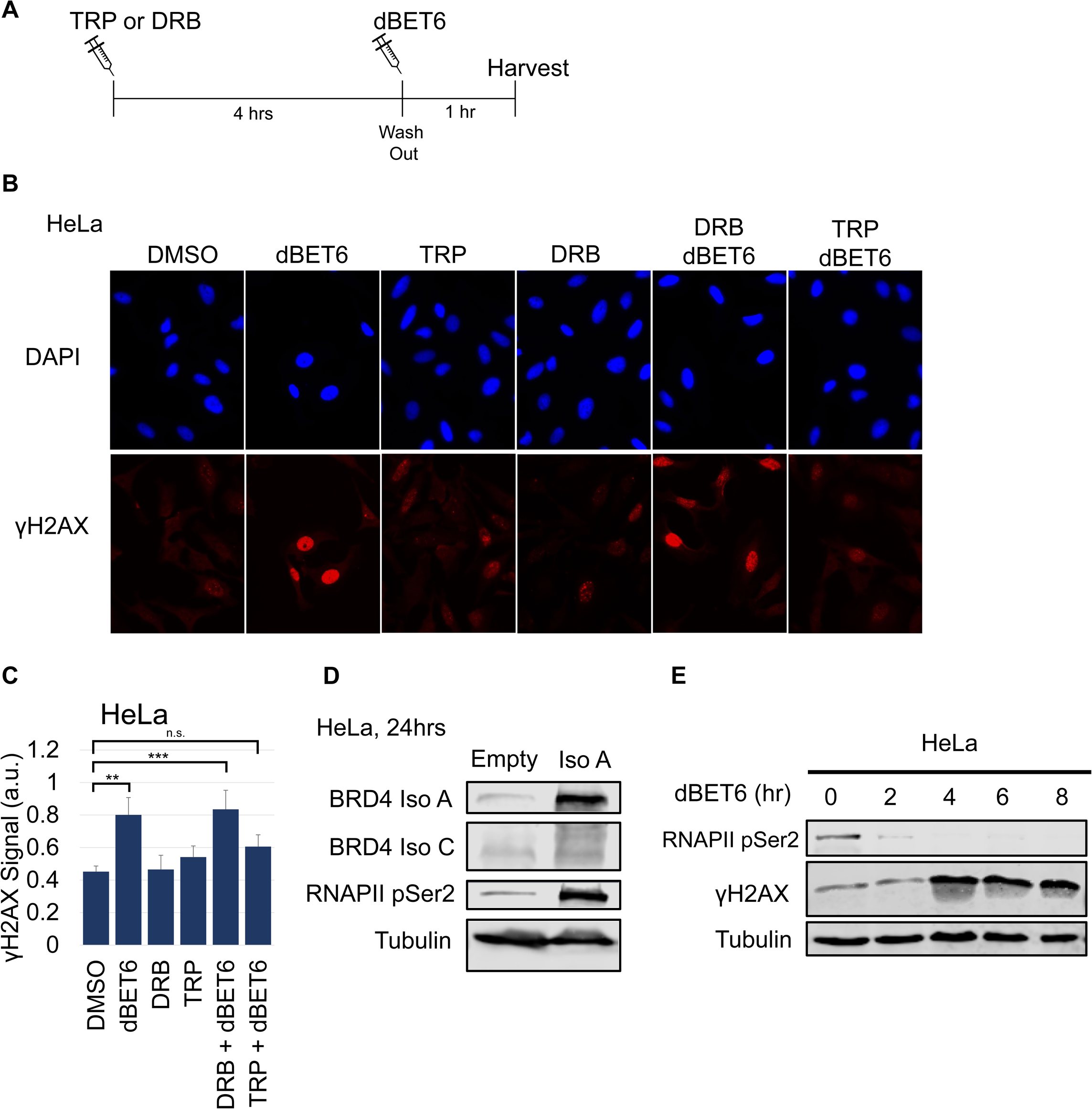
RNAPII loss rescues TRCs caused by BET inhibition **A.** Depiction of experimental design. HeLa cells were treated with 250 nM Triptolide or 100 µM DRB for four hours before being washed out. Subsequently, cells were treated with 100 nM dBET6 for one hour before fixation. **B.** Representative images and **C.** quantification of γH2AX staining per nucleus from HeLa cells treated as described in **A. D.** Representative images of western blots from HeLa cells stably induced with the expression construct shown above each column for 24 hours. **E.** Representative images of western blots from HeLa cells treated with 100 nM dBET6 for indicated times. ANOVA was performed on **c**. Data represent the mean ±SEM. \**P* < 0.05; \*\**P* < 0.01; \*\*\**P* < 0.001. Source data are provided as a Source Data file.

**Figure 6:**
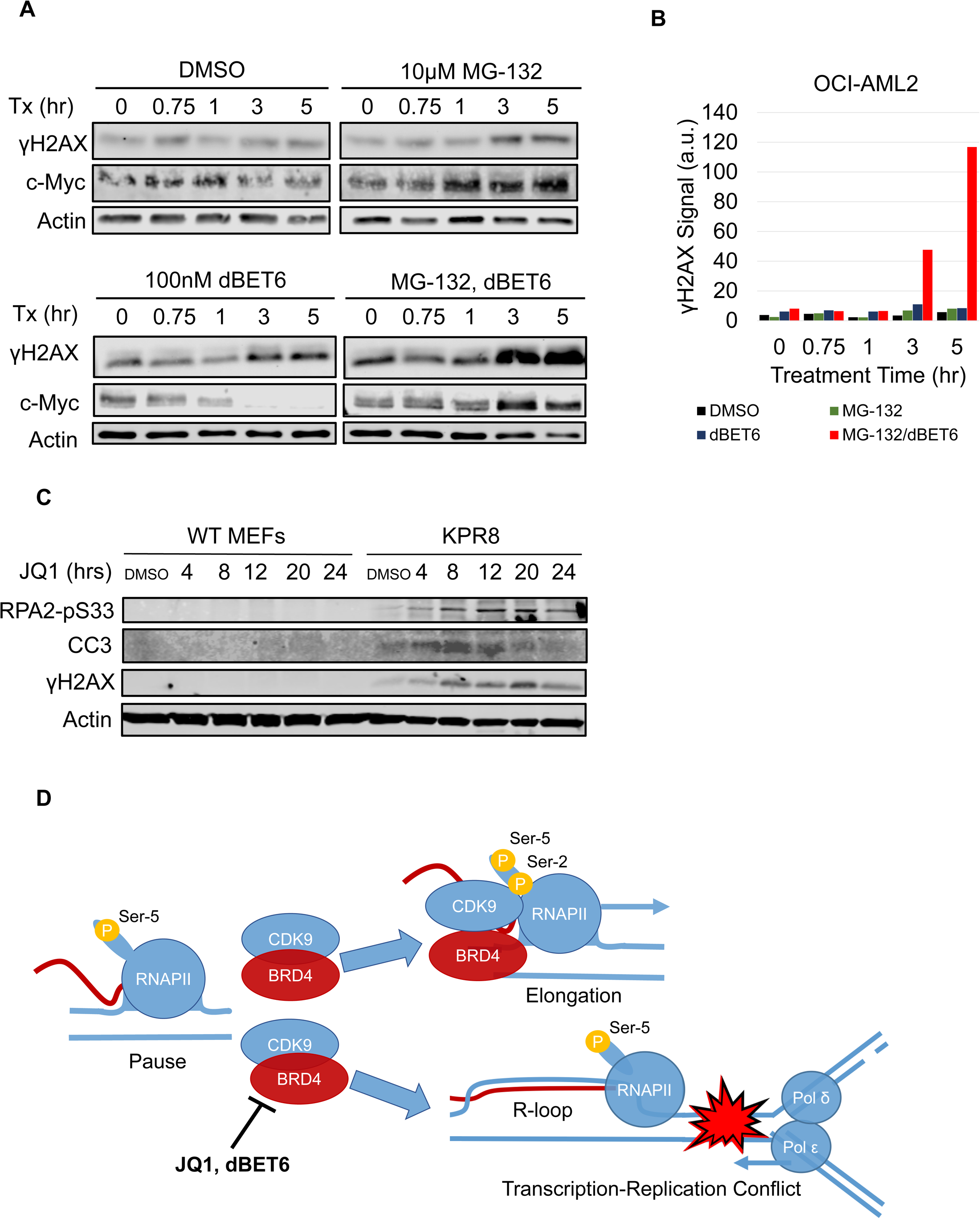
Model depicting the role of BRD4 in the prevention of R-loop-dependent TRCs **A.** Representative images and **B** quantification of western blots from OCI-AML2 cells harvested at various time points following the labeled treatment. **C.** Representative images of western blots from wild-type MEF or KPR8 cells treated with JQ1. **D.** Depiction of proposed model. In normal conditions, BRD4 interacts with CDK9 to ensure the efficient phosphorylation of Serine-2 on the tail of RNAPII to release from transcriptional pause and allow transcription elongation. When BRD4 is inhibited or degraded by JQ1 or dBET6 respectively, RNAPII is unable to release from transcriptional pause or undergo elongation. This results in the build-up of R-loops which lead to TRCs and subsequent DNA damage. For western blots, lysates were probed for the epitope indicated beside each panel. Source data are provided as a Source Data file.

### Active transcription and RNAPII occupancy are required for BET protein-loss induced damage

There are five stages of transcription: RNAPII recruitment, initiation, pause/release, elongation, and termination(Haberle and Stark, 2018; Porrua and Libri, 2015). Transcription initiation is denoted by a phosphorylation event in which CDK7, a subunit of TFIIH, phosphorylates Serine-5 on the tail of RNAPII(Komarnitsky et al., 2000). After ∼50bp of nascent transcription, RNAPII undergoes a pausing event until CDK9, a subunit of P-TEFb, phosphorylates Serine-2 on the tail of RNAPII(Baumli et al., 2012). Inhibitors of these two kinases exist and have been shown to have different effects on RNAPII occupation of chromatin(Shao and Zeitlinger, 2017a). Triptolide (TRP) inhibits TFIIH and results in the blocking of transcription initiation and the degradation of RNAPII **(Figure S5A)**. DRB inhibits CDK9 and leads loss of RNAPIIpS2 and stalling of RNAPII on the chromatin, resulting in R-loops and TRCs(L. Chen et al., 2017; Shao and Zeitlinger, 2017b) **(Figure S5B)**. With this understanding, we hypothesized that these two molecules would have differing effects on the DNA damage caused by BRD4 loss.

To test whether degradation of RNAPII with TRP would be able to rescue the DNA damage effects of dBET6 treatment, we designed an experiment to pre-treat and manipulate RNAPII prior to dBET6 exposure, as described in **Figure 5A**. After pre-treating with either TRP or DRB, we washed out the drugs and treated with dBET6 for one hour. Following the dBET6 treatment, cells were fixed and stained for γH2AX (**Figure 5B**; **Figure 5C**). Remarkably, we saw that TRP was able to rescue the DNA damage effects of dBET6, while DRB was not. We then co-treated HCT-116 cells with TRP and dBET6 in and saw that TRP was also able to rescue the DNA damage effects caused by dBET6 in this cell line **(Figure S5C; Figure S5D)**. These data indicate that RNAPII occupation on the chromatin is necessary for DNA damage caused by BRD4 loss.

We also wanted to explore the relationship between RNAPIIpS2 and DNA damage caused by dBET6 treatment. We observed that when BRD4 isoform A is overexpressed, there is an increase in RNAPIIpS2 (**Figure 5D**). In addition, we see that RNAPIIpS2 negatively correlates with γH2AX following dBET6 treatment both in HeLa cells and HEK-293T cells (**Figure 5E**; **Figure S5E; Figure S5F**). These data again suggest that the loss of BRD4 leads to loss of transcription and pausing of RNAPII on the chromatin causing TRCs and subsequent DNA damage.

### Oncogene-induced enhanced transcription exacerbates BET loss-induced DNA damage

BET proteins have been previously linked to regulation of the oncoprotein, c-Myc (Delmore et al., 2011; Fowler et al., 2014; Mertz et al., 2011; Muhar et al., 2018; Winter et al., 2017). In addition, studies have shown that increased levels of transcription due to increased expression of MYC leads to increased replication stress(Kotsantis et al., 2016b; Lin et al., 2012; Puccetti et al., 2019). We hypothesized that stabilizing c-Myc would sensitize cancer cells to the effects of BET LOF.

To test this, we co-treated OCI-AML2 cells with dBET6 and MG-132, a small molecule which targets the proteasome. When cells were treated with dBET6, we observed the expected rapid loss of c-Myc (**Figure 6A**). However, co-treating cells with dBET6 and MG-132 stabilized c-Myc. Surprisingly, this also led to a synergistic increase in γH2AX signal by western blot (**Figure 6A**; **Figure 6B**). This indicated to us that increased transcriptional signaling from c-Myc primed cells for even more replication stress following BET LOF.

We hypothesized that additional oncogene signaling that enhances transcription and is known to enhance R-loops (Kotsantis et al., 2016b) would also enhance DNA damage and other effects from BET LOF. Therefore, we measured the DNA damage response in normal mouse embryonic fibroblasts exposed to JQ1 and compared this to responses in murine cells expressing mutant KRAS (**Figure 6C**). As expected, the mutant KRAS-expressing cells showed greatly enhanced γH2AX, RPApS33, and cleaved caspase 3 indicating that oncogene expression sensitizes cells BET bromodomain inhibition-dependent DNA damage and cell death.

## DISCUSSION

Inhibitors of BRD4 have been shown to be effective treatments for several cancers, yet the mechanism of action remains unclear(Asangani et al., 2014; Dawson et al., 2011; Rathert et al., 2015; Zuber et al., 2011). Specifically, questions remain as to the mechanism by which inhibition of BRD4, which controls global transcription(Winter et al., 2017), may preferentially impact cancer cells more than normal cells – a feature that is required of all effective chemotherapies. Here, we propose a novel role for BRD4 in the prevention of R-loops, TRCs, S phase-dependent DNA damage, and cell death in highly transcription and replication-driven cells (**Figure 6D**).

Our data show that inhibition or degradation of BET proteins, with JQ1 or dBET6 respectively, leads to an accumulation of DNA damage signaling and DNA double strand breaks. When we characterized the nature of the DNA damage, we found that, in the several cell types we investigated, the cell cycle state dictated whether or not a cell accumulated this damage. Specifically, we saw that cells actively undergoing replication in S-phase preferentially exhibited DNA damage and cell death following BET protein LOF. Historically, BET proteins have been shown to play a major role in transcription regulation, thus we postulated that the S-phase dependent DNA damage caused by BET protein loss could be working through a mechanism of increased TRCs.

Due to the fact that BET protein inhibitors such as JQ1 and degraders such as dBET6 target the bromodomains of BRD2, BRD3, and BRD4, it was previously unclear if one member of the family is responsible for the DNA damage caused by BET protein loss. Several works have shown that the different BET proteins have both unique and shared roles in the cell(Cheung et al., 2017; Hsu et al., 2017; LeRoy et al., 2008),. Our data show that while both BRD2 and BRD4 show increased γH2AX signaling after 72 hours, overexpression of the full-length isoform (isoform A) was sufficient to effectively rescue the DNA damage effects of BET protein degradation by treatment with dBET6. Specifically, we observed that the C-terminal domain of BRD4 was necessary to rescue this effect. Our data and the literature show that the C-terminal domain plays a critical role in the activation of RNAPII to ensure efficient elongation(R. Chen et al., 2014; Itzen et al., 2014; Jang et al., 2005; Kanno et al., 2014; Liu et al., 2013; Patel et al., 2013; Rahman et al., 2011; Winter et al., 2017; W. Zhang et al., 2012). BRD4, through its C-terminal domain, interacts with CDK9 to phosphorylate Serine-2 on the heptapeptide repeat on the tail of RNAPII. This phosphorylation event allows RNAPII to proceed with transcription elongation on schedule. Previous studies have identified DRB, a small molecule inhibitor of CDK9, as a factor that increases R-loop formation (L. Chen et al., 2017). Our work adds to this finding by identifying BRD4, a physiological activator or CDK9, as an important R-loop regulator. Our findings show that BRD4 loss of function causes S-phase-dependent DNA damage through a novel TRC mechanism, specifically in highly transcription-replication driven cells. This novel mechanism is distinct from that proposed for BET bromodomain proteins in other recent work (Bowry et al., 2018; Kim et al., 2019) and impacts the use of BET bromodomain inhibitors, which are in clinical trials for a number of diseases..

In recent years, the importance of R-loops has become more apparent. While they play critical roles normal physiological activity(Chaudhuri and Alt, 2004; Garcia-Muse and Aguilera, 2019; Shao and Zeitlinger, 2017a; Skourti-Stathaki and Proudfoot, 2014; Stuckey et al., 2015; Xiao et al., 2017), it has also come to light that aberrant R-loops can lead to TRCs, DNA damage, and cell death(Aguilera and Gómez-González, 2017; Costantino and Koshland, 2018; Crossley et al., 2019; Garcia-Muse and Aguilera, 2019; Hamperl et al., 2017; Hamperl and Cimprich, 2016; Richard and Manley, 2016; Santos-Pereira and Aguilera, 2015; Sollier and Cimprich, 2015). Our data show that DNA damage caused by BRD4 loss is correlated with an increase in R-loop formation on the chromatin. Furthermore, this damage can be rescued by overexpressing RNAse H1, an endonuclease that resolves R-loops. These observations indicate that some cells may depend on BRD4 to ensure that efficient transcription during S-phase prevents R-loop dependent conflicts between transcription and replication. We believe this is an important observation, especially for cells with elevated replicative and transcriptional drive. Thus, cancer cells and other highly-driven cells may be more dependent on BRD4 to prevent the transcription and replication machinery from colliding. This finding may shed light on additional prior studies. Early work on BRD4 knockout mice showed both embryonic lethality and replication deficits (Houzelstein et al., 2002; Maruyama et al., 2002b). Additionally, studies of the normal tissue toxicities of whole-animal knockout of BRD4 could indicate vulnerability in rapidly replicating normal tissues(Bolden et al., 2014). Finally, it informs BET inhibition synergy with ATR inhibitors resulting in increased γH2AX signaling and cell death(Pericole et al., 2019). Our proposed mechanism would predict that this synergism exists by increasing the number of TRCs while simultaneously inhibiting a cell’s ability to handle replication stress.

One outstanding question that remains to be completely resolved is what makes a cancer cell more or less sensitive to BRD4 loss. It has been shown that certain cancer cell lines are more sensitive to BET protein inhibition(Rathert et al., 2015), yet it is unclear as to why this is the case. For example, our group and others have shown that BRD4 loss in certain cell lines do not result in an increase in DNA damage signaling, although recent reports have disputed this (Bowry et al., 2018; Floyd et al., 2013b; Kim et al., 2019). Notably, it is reported that that certain cell lines do not exhibit a decrease in RNAPIIpS2 following BRD4 loss, nor do R-loops formed by BRD4 loss exhibit replication stress (Bowry et al., 2018; Kim et al., 2019). As is well known, different cells operate under different transcriptional programming. We hypothesize that certain cancer cell lines may be more globally dependent on BRD4-mediated transcriptional activation, leading to R-loops, TRCs, DNA damage, and cell death upon BET inhibition. We believe that RNAPIIpS2 loss after BRD4 degradation could be predictive of whether a cancer cell line exhibits DNA damage following treatment. Through further study of both BRD4 and the role of R-loops in cancer, we hope that we can identify new chemotherapeutic targets and broaden the effectiveness of BET inhibitors in cancer therapies.

## METHODS

### Cell Culture

HeLa (ATCC), HEK-293T (ATCC), mouse embryonic fibroblast (MEF), and K-rasV12D-p53 deleted (KPR8) cells were cultured in Dulbecco’s modified Eagle’s medium (DMEM) (Genesee Scientific) supplemented with 10% fetal bovine serum (FBS) (Summerlin Scientific Products) and 1% penicillin/streptomycin (P/S) (Thermo Fisher Scientific). HCT-116 (Duke Cell Culture Facility-verified) cells were cultured in McCoy’s 5A medium (Thermo Fisher Scientific) supplemented with 10% FBS and 1% P/S. OCI-AML2 cells were cultured in Roswell Park Memorial Institute 1640 medium (RPMI) (Thermo Fisher Scientific) supplemented with 10% FBS and 1% P/S. MEF and KPR8 cells were kind gift from Tyler Jacks.

### Antibodies and stains

The following antibodies were used for western blot (WB), immunofluorescence (IF), or ChIP experiments: BRD4 N-terminus (1:1000WB, 1:1000IF, ab128874, Abcam); BRD2 (1:500WB, 5848S, Cell Signaling Technology); BRD3 (1:100WB, ab50818, Abcam); RNAPIIpS2 (1:1000WB, 1:50ChIP, 04-1571, EMD Millipore); γH2AX (1:1000WB, 1:1000IF, 1:50ChIP, 9718S, Cell Signaling Technology); ⍺-Tubulin (1:1000WB, 2144S, Cell Signaling Technology); RPA2pS33 (1:500WB, ab211877, Abcam); cleaved PARP (1:1000WB, 5625, Cell Signaling Technology); BrdU (1:20IF, 347580, BD Biosciences); BrdU (1:80IF, ab6326, Abcam); V5 (1:1000IF, 1:50ChIP, ab9116, Abcam); Total RNAPII (1:000WB, 1:50ChIP, ab817, Abcam); c-Myc (1:1000WB, 5605S, Cell Signaling Technology); Cleaved Caspase 3 (1:500WB, 9664S, Cell Signaling Technology); SETX (1:500WB, ab220827, Abcam); SRSF1 (1:500WB, 324600, Thermo Fisher Scientific); DHX9 (1:1000WB, PA519542, Thermo Fisher Scientific); HEXIM (1:1000WB, ab25388, Abcam); Top2a (1:1000WB, ab52934, Abcam); Top 2b (1:1000, ab72334, Abcam); XPF (1:1000WB, ab76948, Abcam); XPG (1:1000WB, ab224815, Abcam); Goat Anti-Rabbit IgG 800CW (1:6000WB, 926-32211, LI-COR Biosciences); Goat Anti-Mouse IgG 680RD (1:6000WB, 926-68070, LI-COR Biosciences); Goat Anti-Rat IgG 680LT (1:6000, 926-68029, LI-COR Biosciences); Goat Anti-Rabbit IgG Alexa Fluor 647nm (1:500IF, A211245, Life Technologies); Goat Anti-Rabbit IgG Alexa Fluor 555nm (1:500IF, A21428, Invitrogen); Goat Anti-Rabbit IgG Alexa Fluor 488nm (1:500IF, A11008, Life Technologies); Goat Anti-Mouse IgG Alexa Fluor 488 (1:500IF, A11001, Invitrogen); Goat Anti-Rat IgG Alexa Fluor 647 (1:500IF, A21247, Invitrogen). DAPI (1:2000IF, Thermo Fisher Scientific) was used to stain nuclei. SYBR Gold (1X, Thermo Fisher Scientific) was used to stain single cell electrophoresis (comet) assay. Propidium Iodide (50 µg/mL, VWR) was used to stain nuclei for cell cycle analysis.

### Immunofluorescence

Cells were grown on coverslips or in micro-chamber wells (Ibidi) overnight before induction or treatment. When the experiment was completed, cells were washed with ice cold PBS and fixed with 4% paraformaldehyde for 20 minutes at room temperature (RT). After fixation, cells were washed with PBS and then blocked in 5% goat serum and .25% Triton-X for 1 hour at RT, rocking. Following blocking, primary antibodies were diluted in the same blocking buffer and incubated at 4°C overnight, rocking. Following incubation with primary antibody, cells were washed three times with PBS and stained with the appropriate secondary antibody diluted and DAPI in blocking buffer at RT for 1 hour, rocking. After incubation with secondary antibody, cells were washed three times with PBS. In the case of cells grown on coverslips, cells were mounted on slides using Prolong Gold (Thermo Fisher Scientific) before imaging. Cells grown in micro-chamber wells were left in PBS before immediate imaging. Immunofluorescence images were taken either on a Zeiss Axio Observer or EVOS microscope using a 40X objective. Quantification of γH2AX foci was done using the speckle counting pipeline in CellProfiler. All images within a single experiment were fed into the same pipeline and speckles (foci) were counted in an unbiased fashion using the automated program. γH2AX signal is defined as the multiplication of foci count of a nucleus with the mean integrated intensity of the foci within that nucleus.

### Western Blotting

Whole cell lysates were prepared with a whole cell lysis buffer (50mM Tris-HCl pH 8.0, 10mM EDTA, 1% SDS) with protease and phosphatase inhibitors (Thermo, 78440) added fresh. Lysates were then sonicated using a QSonica Q700 sonicator for two minutes with an amplitude of 35. After sonication, protein concentrations were determined using BCA reagents (Pierce), compared to protein assay standards (Thermo Fischer Scientific), and scanned using a Spectramax i3x. Equivalent amounts of protein were resolved by SDS-PAGE gels and transferred to nitrocellulose membranes. Membranes were then blocked with a 1:1 solution of PBS and Odyssey Blocking Buffer (LI-COR Biosciences) at RT for one hour, rocking. Primary antibodies were then diluted in the blocking buffer as described above and incubated with the membranes at 4°C overnight. Membranes were then washed three times with 0.2% Tween-20 in PBS (PBS-T). The appropriate secondary antibodies were also diluted in the blocking buffer and incubated with the membranes at RT for one hour. Membranes were then washed with PBS-T three times and scanned using a LI-COR Odyssey scanner. Quantification and normalization of western blot signal was done using the LI-COR software, Image Studio.

### Single Cell Electrophoresis (Comet) Assay

Neutral comet assays were performed using the CometAssay Reagent Kit (Trevigen) according to the manufacturer’s protocol. Briefly, cells were washed in ice cold PBS, scraped from the plate, mixed with low melt agarose and spread onto supplied microscope slides in the dark. The agarose was gelled at 4°C for 30 minutes before being submerged in the supplied lysis buffer 4°C overnight in the dark. Slides were then incubated with chilled neutral electrophoresis buffer at 4°C for 30 minutes before being subjected to 21V for 45 minutes. Slides were submerged with DNA precipitation at RT for 30 minutes and then 70% ethanol at RT for 30 minutes. Slides were then dried and stained with 1X SYBR gold as described above. Comets were imaged on a Zeiss Axio Observer using a 10X objective. Comets were quantified using the comet pipeline from CellProfiler. All images within a single experiment were fed into the same pipeline and comets were quantified in an unbiased fashion using the automated program. Extent Tail moment is defined as Tail DNA % multiplied with the length of the comet tail.

### Transfections

For RNA interference, cells were incubated with Thermo Fisher *Silencer*® Select Pre-designed or Dharmacon ON-TARGETplus siRNAs for BRD2 (Thermo, s12071), BRD3 (Thermo, s15545), BRD4 (Thermo, 23902), HEXIM (Thermo, s20843), Top2a (Dharmacon, L-004239-00-0005), Top2b (Dharmacon, L-004240-00-0005), XPF (Dharmacon, L-019946-00-0005), XPG (Dharmacon, L-006626-00-0005), or negative control (Thermo, 4390846). Transfections were done with Lipofectamine RNAiMAX transfection reagent (Invitrogen) according to the manufacturer’s protocol.

For transfection of RNAse H1 constructs, cells were transfected with WT RNase H1 (Addgene, 111906), the D210N mutant (Addgene, 111904), or WKKD mutant (Addgene, 111904) which were a gift from Xiang-Dong Fu and previously described(L. Chen et al., 2017). 750 fmol of plasmid was incubated with a 6:1 ratio of Xtremegene HP transfection reagent at RT for 20 minutes in 1 mL of Opti-mem media. The transfection mixture was then added dropwise to a 10cm dish containing Cells at 70% confluence for 24 hours. Cells were then selected with 100 µg/mL hygromycin for 24 hours before fixing (for immunofluorescence experiments) or immediately fixed (for ChIP experiments).

### DNA Fiber Analysis

DNA fiber analysis was performed as previously described(Quinet et al., 2017). Briefly, cells were plated at 1×10^5^ cells per well in a 6-well plate and incubated overnight. Cells were then pulsed with the appropriate thymidine analog and treated as shown in **Figure 2G**. Cells were then washed with PBS and placed on Superfrost Plus Microscope slides and lysed. Following lysis, slides were tilted by raising the edge of the slide 2.2 cm to allow DNA fibers to stretch along the slide and left to dry. DNA was then fixed in 3:1 methanol:acetic acid, dried, and washed in PBS before HCl denaturation of the DNA. The slides were then blocked in 5% BSA before being stained with primary antibodies to detect IdU (mouse anti-BrdU) or CldU (rat anti BrdU). Slides were then stained with the appropriate secondary antibodies before imaging. Images were taken on a Leica SP5 microscope using a 100X objective. ImageJ was used to measure lengths of fibers.

### Plasmid Construction

The iBRD4 plasmids were constructed using the pCW57-GFP-2A-MCS backbone (Addgene, 71783), which was a gift from Adam Karpf and previously described(Barger et al., 2019). Gibson assembly was used to insert either mCherry-2A-Flag-BRD4 isoform A or isoform C into the backbone in place of the TurboGFP-P2A-hPGK promoter-PuroR-T2A-rTetR region. The C-terminal domain was deleted from isoform A using PCR (AΔCTD). The extra-terminal domain was deleted from isoform C using PCR (CΔET). Sanger sequencing was performed to verify the cloning products.

### Small Molecule Inhibitors

The BET protein degrader dBET6 was a gift from Nathanael Gray and previously described(Winter et al., 2017). dBET6 was used at a concentration of 100 nM in all experiments except those involving the iBRD4 system, in which it was used at 10 nM. The BET bromodomain inhibitor JQ1 was a gift from James Bradner and previously described(Filippakopoulos et al., 2010b). JQ1 was used at a concentration of 500 nM for all experiments. The CDK9 inhibitor DRB (Cayman Chemical Company, 10010302) was used at a concentration of 100 µM for all experiments. The TFIIH inhibitor triptolide (EMD Millipore, 645900) was used at a concentration of 250 nM (HeLa) or 1 µM (HCT-116). The proteasome inhibitor, MG-132 (Selleck Chemicals) was used at a concentration of 10 µM. The topoisomerase I inhibitor, camptothecin (Selleck Chemicals), was used at a concentration of 10 µM. The topoisomerase II inhibitor, dexrazoxane (Selleck Chemicals), was used at a concentration of 50 µM. The BET bromodomain inhibitors OTX015 (Selleck Chemicals), ABBV-075 (Selleck Chemicals), ABBV-744 (Selleck Chemicals), and PLX51107 (Selleck Chemicals) where used at 2 µM, 20 nM, 50 nM, and 2 µM, respectively.

### EdU Detection

EdU detection was done according using the EdU-Click Chemistry 488 kit (Sigma-Aldrich, BCK-EDU488) according to manufacturer’s instructions. In brief, cells were pulsed with 10 µM EdU alongside simultaneous treatment with DMSO or dBET6. Cells were then fixed, washed with PBS, and blocked as described above. Cells were then incubated at RT for 30 minutes in the click chemistry cocktail. Following incubation, cells were washed three times with PBS. After the click chemistry was completed, cells were further process according to the immunofluorescence methods described above.

### Flow Cytometry and Cell Cycle Analysis

For cell cycle analysis, cells were trypsinized and washed with ice cold PBS. Cells were then fixed with 70% ethanol at 4°C for 30 minutes. Cells were then washed with PBS twice before being incubated with 100 µg/mL RNAse A and 50 µg/mL propidium iodide overnight at 4°C. Cells were then quantified by flow cytometry for DNA content on a BD FACSCanto II machine. For analysis, flow results were entered into the univariate cell cycle modeling in FlowJo for the distribution of cell cycle. Analysis of sub-G_1_ populations were done as previously described(Riccardi and Nicoletti, 2006).

For EdU and γH2AX flow experiments, cells were fixed and stained according to the EdU click chemistry and immunofluorescence methods described above. Cells were then quantified for EdU and γH2AX signal on a BD FACSCanto II machine. FlowJo was then used to generate the figures. Cells that were not pulsed with EdU were used as a negative control for EdU click chemistry. Cells not stained with γH2AX primary antibody were used as a negative control for γH2AX staining.

### Chromatin Immunoprecipitation Followed by Next Generation Sequencing (ChIP-seq)

Wild-type HeLa cells (ChIP) or cells transfected with RNAse H1 D210N (R-ChIP) were both prepared for qPCR or sequencing using the SimpleChIP® Plus Sonication Chromatin IP Kit according to the manufacturer’s instructions. In brief, cells were washed with ice cold PBS and then fixed with 1% formaldehyde in PBS at RT for 13 minutes. The fixation reaction was then halted using a 1X Glycine solution. Cells were then scraped from the plates and pelleted. Cells were then incubated with 1X ChIP sonication cell lysis buffer plus protease inhibitors (PIC) on ice for 10 minutes. Cells were then pelleted and the previous step was repeated. Nuclei were then pelleted and resuspended in ice cold ChIP Sonication Nuclear Lysis buffer with PIC and incubated on ice for 10 minutes. Lysates were then fragmented by sonication with a QSonica Q700 at 4°C for 15 minutes ON-time with a 15s on, 45s off program. After sonication, a sample for 2% input was removed. 10 µg of lysates were then incubated with a ChIP grade antibody at 4°C overnight. 30 uL of magnetic beads were then added to the mixture and incubated at 4°C for two hours before going through a series of salt washes. Chromatin was then eluted from the magnetic beads in the elution buffer at 65°C for 30 minutes while vortexing. The supernatant was removed and treated with RNAse A followed by Proteinase K. ChIP DNA was then purified using the supplied columns. Library preparation, Next Generation Sequencing, and analysis was performed by GeneWiz to determine the level of ChIP-seq or R-ChIP-seq signal following DMSO or dBET6 treatment for two hours. Log2 ratio normalization to input was done using the bamCompare function of deepTools with default inputs.

### Chromatin Immunoprecipitation Followed by qPCR (ChIP-qPCR)

DNA for ChIP-qPCR and R-ChIP-qPCR was prepared the same was as described for ChIP-seq experiments. Equal volumes of DNA template were subjected to qPCR with qPCR primers designed against the transcription start sites, exons, introns, and transcription termination sites of candidate genes using iTaq Universal SYBR Green Supermix. Samples were normalized to input to determine the relative amounts of ChIP and R-ChIP signal after DMSO or dBET6 treatment for two hours. Primer sequences can be found in the Source File. RNAPII travel ratios were calculated as previously described(Winter et al., 2017).

## ACKNOWLEDGEMENTS

We thank Xiang-Dong Fu for providing the RNAse H1 constructs, Adam Karpf for providing the pCW57 construct, Nathanael Gray for providing dBET6, James Bradner for providing JQ1, and Tyler Jacks for providing MEF and KPR8 cells. We also thank Duke MSTP for providing funding for D.E. to conduct this work. The work was funded by Burroughs Wellcome Career Award for Medical Scientists and American Cancer Society Research Scholar Grant 133394-RSG-19-030-01-DMC to S.R.F.

## AUTHOR CONTRIBUTIONS

D.E and S.R.F. designed the project. D.E., R.M., J.P.T., J-H.P., E.B-S., Jie L., and Jin. L. conducted the experiments. D.E., R.M., Jie L., and S.R.F. analyzed the data. D.E. and S.R.F. wrote the manuscript.

---

**Figure S1:**
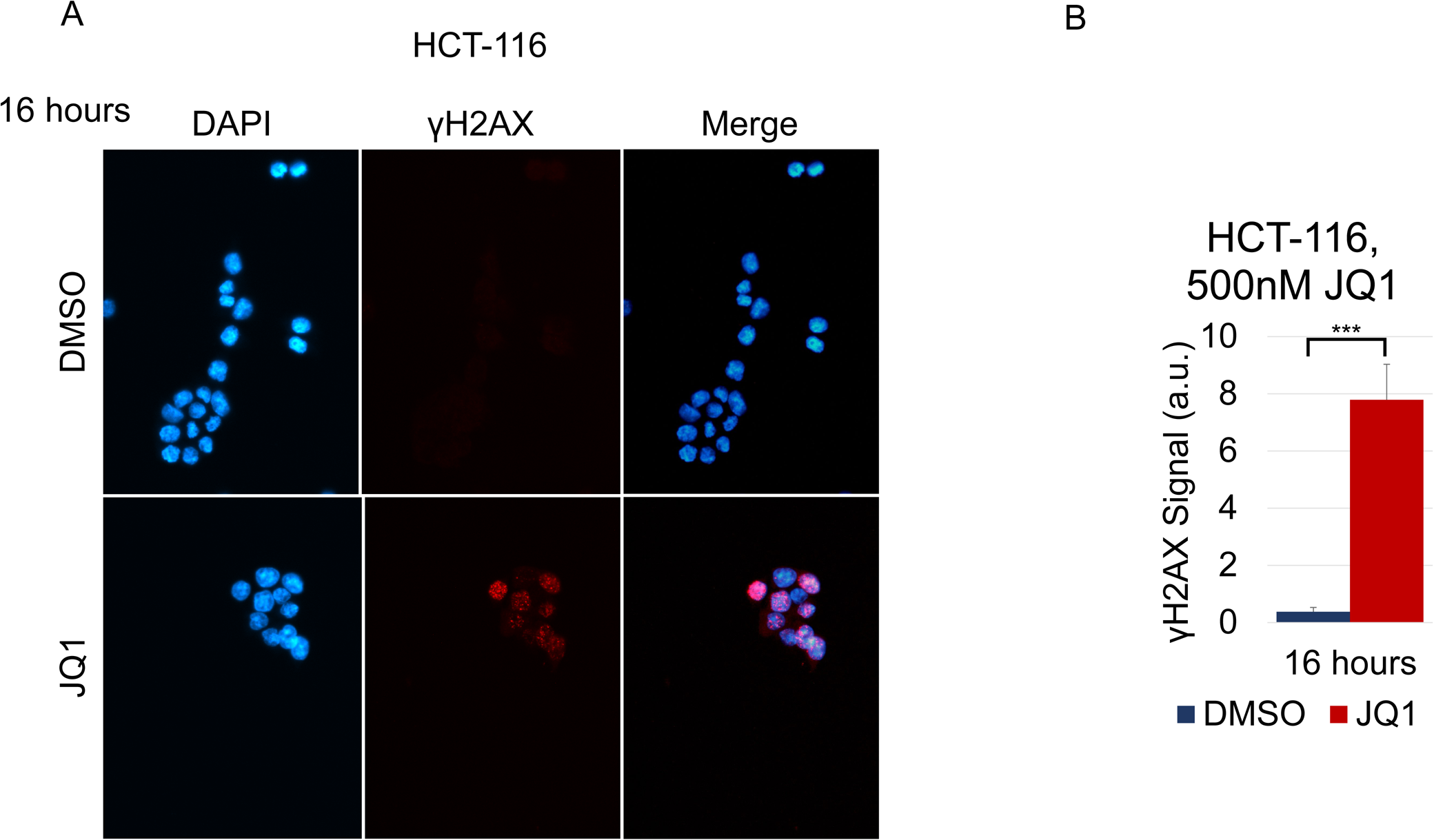
BET protein loss of function leads to spontaneous DNA damage. **A.** Representative images and **B.** quantification of γH2AX staining per nucleus in HCT-116 cells treated with DMSO or 500 nM JQ1 for 16 hours. Student’s t-test (two-tailed, unpaired) was performed on **B**. Data represent the mean ±SEM. \**P* < 0.05; \*\**P* < 0.01; \*\*\**P* < 0.001. Source data are provided as a Source Data file.

**Figure S2:**
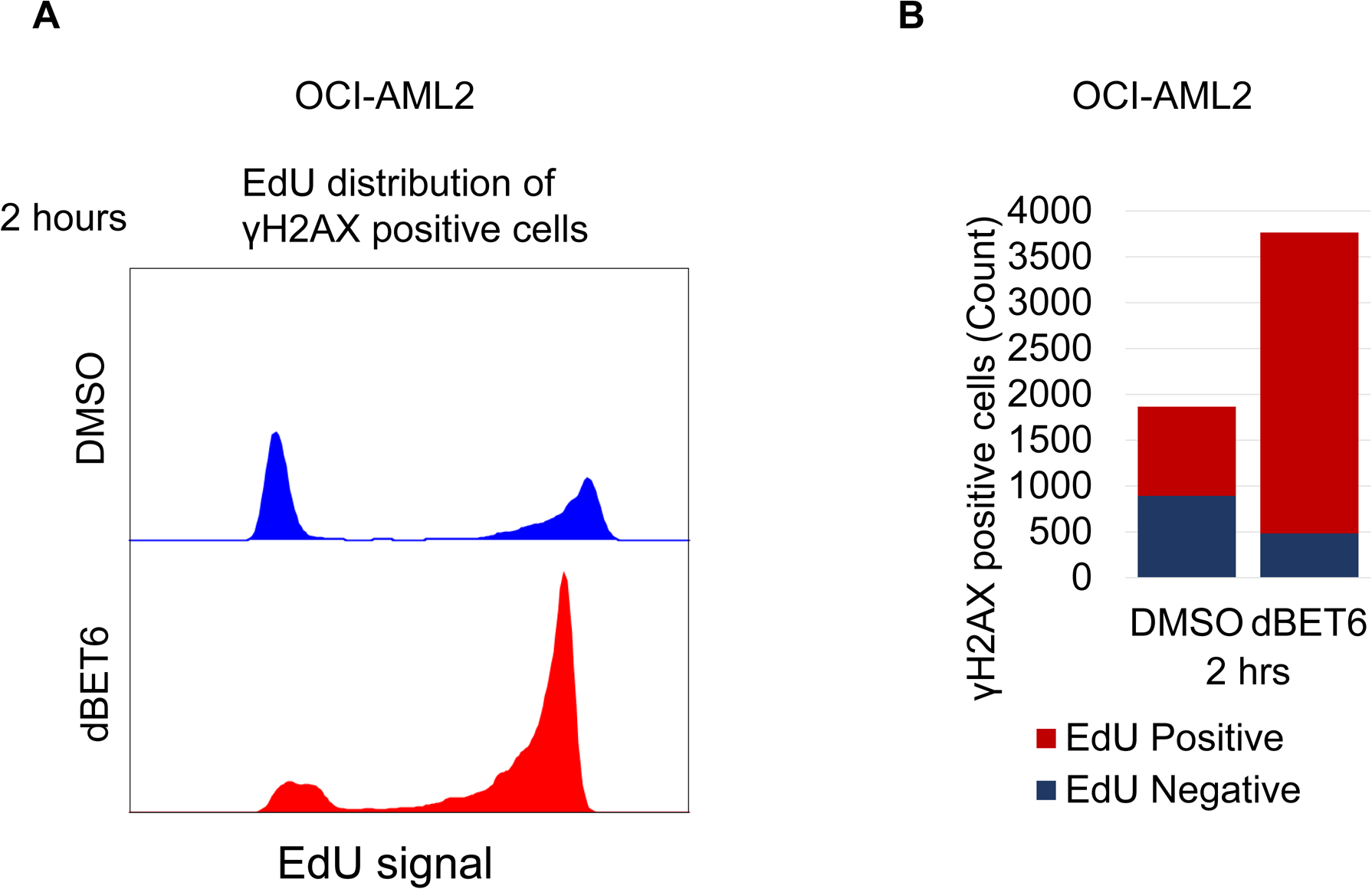
BET protein degradation leads to replication stress and S-phase-dependent DNA damage. **A.** Flow cytometry distribution of EdU in cells that positively stained for γH2AX signaling. OCI-AML2 cells were treated with DMSO or dBET6 for 2 hours before fixation. **B.** Quantification of γH2AX staining per nucleus in OCI-AML2 cells treated simultaneously with 100 nM dBET6 and 10 µM EdU for 2 hours. Source data are provided as a Source Data file.

**Figure S3:**
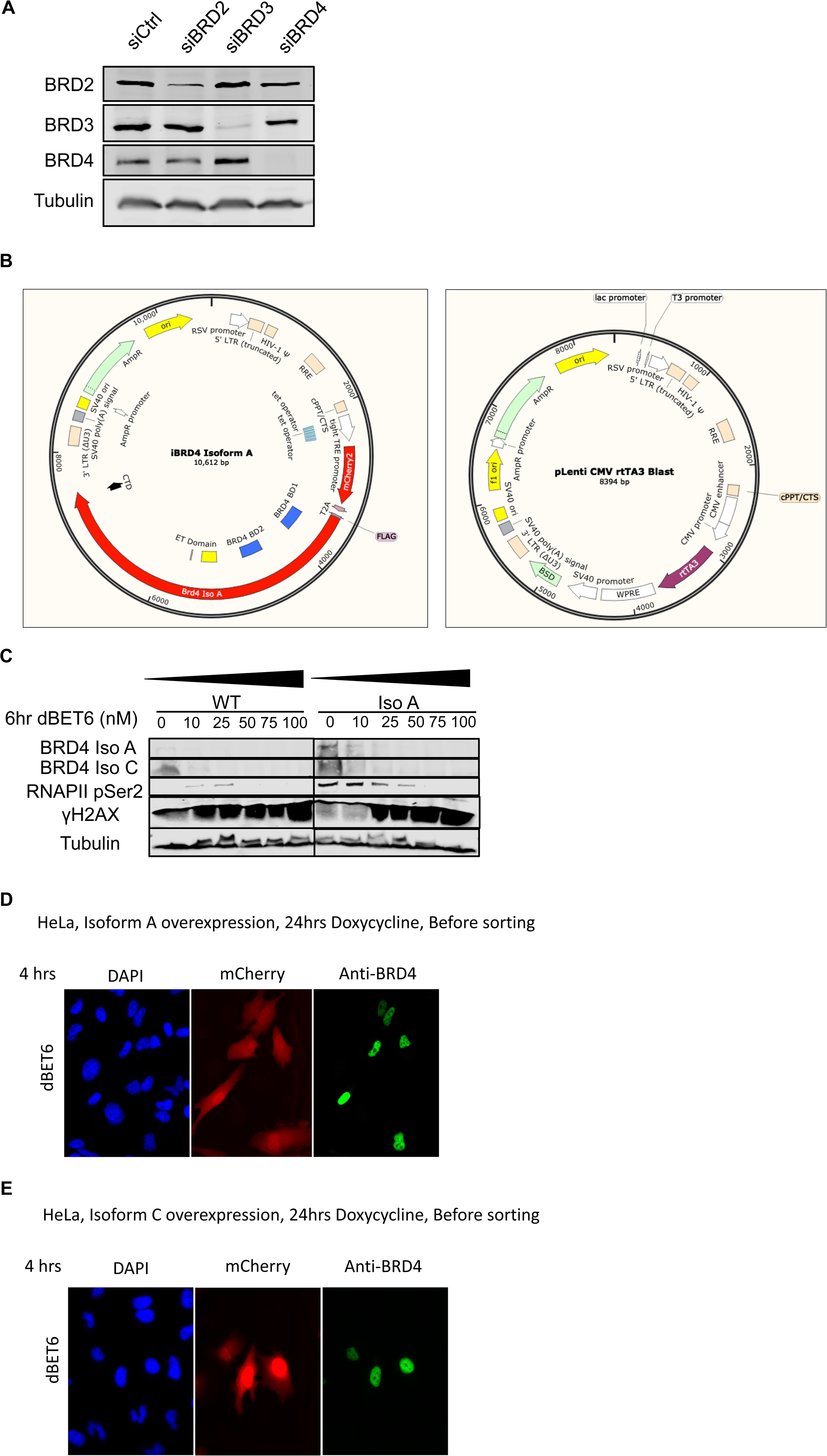
The C-terminal domain of BRD4 is required to prevent transcription-replication conflicts. **A.** Representative western blot images of HeLa cells treated with the described siRNA for 72 hours. **B.** Snapgene files depicting the 2-vector iBRD4 system. Lentiviral, doxycycline-inducible BRD4 isoform A construct (left panel) and rtTA3 (right panel) were co-infected and selected by blasticidin and mCherry flow sorting to obtain a pure population. **C.** Representative western blot images of HeLa cells induced with doxycycline for 24 hours and then treated with increasing levels of dBET6 for 6 hours. **D.** Representative images of HeLa cells harboring BRD4 Isoform A construct induced with doxycycline for 24 hours before being treated with 100 nM dBET6 for 4 hours. **E.** Representative images of HeLa cells harboring BRD4 Isoform C construct induced with doxycycline for 24 hours before being treated with 100 nM dBET6 for 4 hours. For western blots, lysates are probed for the epitope as described beside each panel. Source data are provided as a Source Data file.

**Figure S4:**
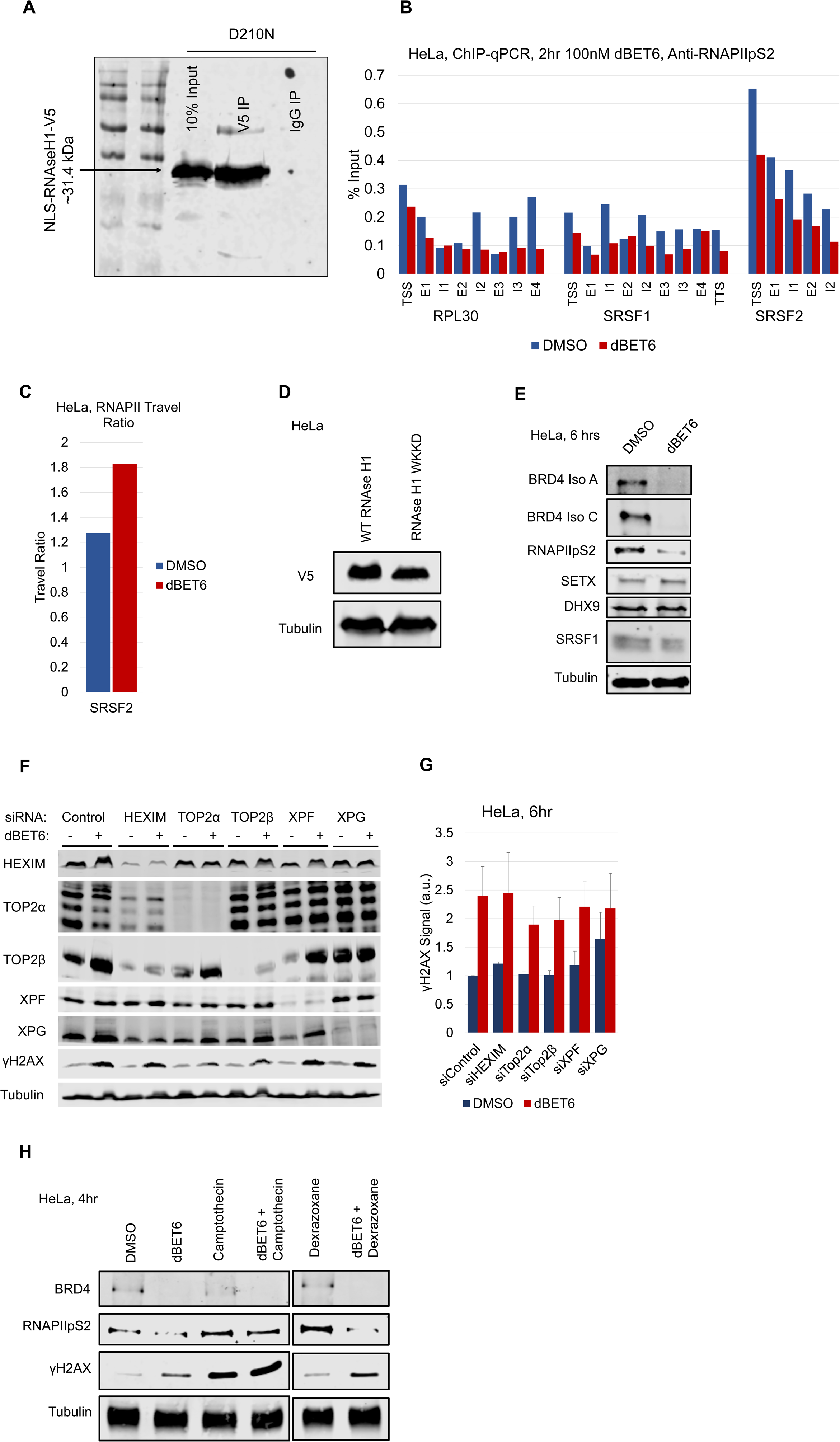
BET inhibition leads to an increase in R-loop-dependent DNA damage **A.** Western blot image depicting immunoprecipitation of RNAse H1 D210N to validate V5 specificity. HEK-293T cells were induced with RNAseH1-D210N-V5 before harvest and immunoprecipitated with an anti-V5 or anti-IgG antibody and compared to input. **B.** ChIP-qPCR signal for RNAPIIpS2 following treatment of HeLa cells with DMSO or dBET6 for 2 hours at loci described in Figure 4B. **C.** Comparison of RNA Pol II traveling ratios between DMSO and dBET6 treatment for genes at the SRSF2 locus based on RNAPII ChIP-qPCR. **D.** Western blot image confirming validation of wild-type or WKKD mutant RNAseH1-V5 constructs. **E.** Representative western blot images from HeLa cells treated with DMSO or dBET6 for 6 hours. **F.** Representative western blot images and **G.** quantification of HeLa cells with various siRNA knock downs and treated with dBET6. **H.** Representative western blot images of HeLa cells treated with dBET6 in combination with camptothecin or dexrazoxane. For western blots, lysates are probed for the epitope as described beside each panel. Source data are provided as a Source Data file.

**Figure S5:**
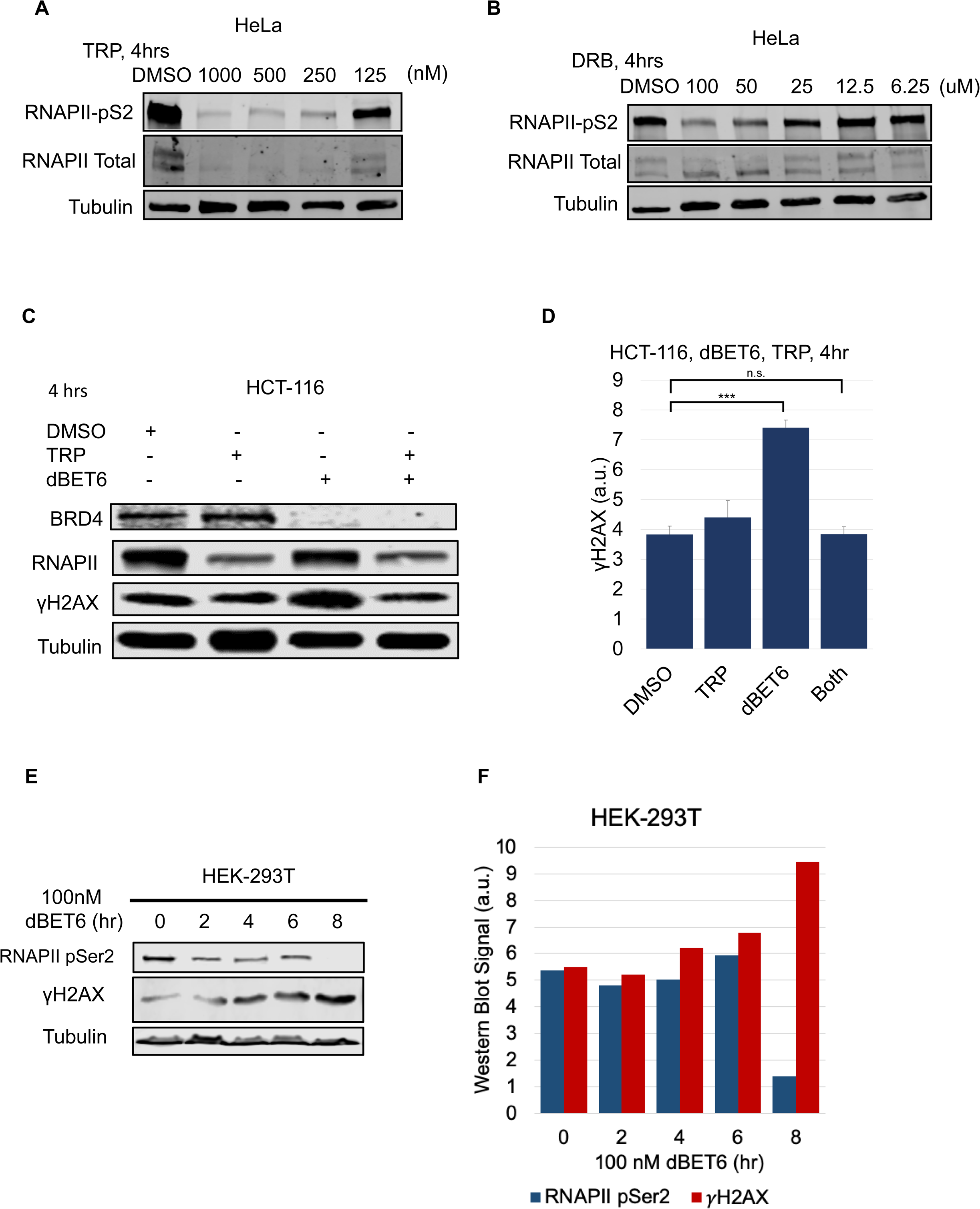
RNAPII loss rescues TRCs caused by BET inhibition **A.** Western blot images of HeLa cells treated with DMSO or decreasing levels of triptolide for four hours: lysates probed for the epitope as described beside each panel. **B.** Western blot images of HeLa cells treated with DMSO or decreasing levels of DRB for four hours: lysates probed for the epitope as described beside each panel. **C.** Representative images and **D.** quantification of western blot images of HCT-116 cells treated with DMSO, 1 µM triptolide, and/or 100 nM dBET6 as described for four hours before harvest: lysates probed for the epitope described beside each panel. **E.** Representative images and **F.** quantification of western blots of HeLa cells treated with 100 nM dBET6 for indicated times: lysates probed for the epitope described next to each panel. ANOVA was performed on **D**. Data represent the mean ±SEM. \**P* < 0.05; \*\**P* < 0.01; \*\*\**P* < 0.001. Source data are provided as a Source Data file.

## REFERENCES

1. Aguilera, A., Gómez-González, B., 2017. DNA–RNA hybrids: the risks of DNA breakage during transcription. Nat Struct Mol Biol 24, 439–443. doi:10.1038/nsmb.3395

2. Asangani, I.A., Dommeti, V.L., Wang, X., Malik, R., Cieslik, M., Yang, R., Escara-Wilke, J., Wilder-Romans, K., Dhanireddy, S., Engelke, C., Iyer, M.K., Jing, X., Wu, Y.-M., Cao, X., Qin, Z.S., Wang, S., Feng, F.Y., Chinnaiyan, A.M., 2014. Therapeutic targeting of BET bromodomain proteins in castration-resistant prostate cancer. Nature 510, 278–282. doi:10.1038/nature13229

3. Baranello, L., Wojtowicz, D., Cui, K., Devaiah, B.N., Chung, H.-J., Chan-Salis, K.Y., Guha, R., Wilson, K., Zhang, X., Zhang, H., Piotrowski, J., Thomas, C.J., Singer, D.S., Pugh, B.F., Pommier, Y., Przytycka, T.M., Kouzine, F., Lewis, B.A., Zhao, K., Levens, D., 2016. RNA Polymerase II Regulates Topoisomerase 1 Activity to Favor Efficient Transcription. Cell 165, 357–371. doi:10.1016/j.cell.2016.02.036

4. Barger, C.J., Branick, C., Chee, L., Karpf, A.R., 2019. Pan-Cancer Analyses Reveal Genomic Features of FOXM1 Overexpression in Cancer. Cancers (Basel) 11. doi:10.3390/cancers11020251

5. Baumli, S., Hole, A.J., Wang, L.-Z., Noble, M.E.M., Endicott, J.A., 2012. The CDK9 Tail Determines the Reaction Pathway of Positive Transcription Elongation Factor b. Structure/Folding and Design 20, 1788–1795. doi:10.1016/j.str.2012.08.011

6. Bermejo, R., Lai, M.S., Foiani, M., 2012. Preventing replication stress to maintain genome stability: resolving conflicts between replication and transcription. Mol Cell 45, 710–718. doi:10.1016/j.molcel.2012.03.001

7. Bisgrove, D.A., Mahmoudi, T., Henklein, P., Verdin, E., 2007. Conserved P-TEFb-interacting domain of BRD4 inhibits HIV transcription. Proc. Natl. Acad. Sci. U.S.A. 104, 13690–13695. doi:10.1073/pnas.0705053104

8. Blackford, A.N., Jackson, S.P., 2017. ATM, ATR, and DNA-PK: The Trinity at the Heart of the DNA Damage Response. Mol Cell 66, 801–817. doi:10.1016/j.molcel.2017.05.015

9. Bolden, J.E., Tasdemir, N., Dow, L.E., van Es, J.H., Wilkinson, J.E., Zhao, Z., Clevers, H., Lowe, S.W., 2014. Inducible in vivo silencing of Brd4 identifies potential toxicities of sustained BET protein inhibition. Cell Reports 8, 1919–1929. doi:10.1016/j.celrep.2014.08.025

10. Bowry, A., Piberger, A.L., Rojas, P., Saponaro, M., Petermann, E., 2018. BET Inhibition Induces HEXIM1- and RAD51-Dependent Conflicts between Transcription and Replication. Cell Reports 25, 2061–2069.e4. doi:10.1016/j.celrep.2018.10.079

11. Canela, A., Maman, Y., Huang, S.-Y.N., Wutz, G., Tang, W., Zagnoli-Vieira, G., Callen, E., Wong, N., Day, A., Peters, J.-M., Caldecott, K.W., Pommier, Y., Nussenzweig, A., 2019. Topoisomerase II-Induced Chromosome Breakage and Translocation Is Determined by Chromosome Architecture and Transcriptional Activity. Mol Cell 75, 252–266.e8. doi:10.1016/j.molcel.2019.04.030

12. Chaudhuri, J., Alt, F.W., 2004. Class-switch recombination: interplay of transcription, DNA deamination and DNA repair. Nat. Rev. Immunol. 4, 541–552. doi:10.1038/nri1395

13. Chen, J.-Y., Zhang, X., Fu, X.-D., Chen, L., 2019. R-ChIP for genome-wide mapping of R-loops by using catalytically inactive RNASEH1. Nat Protoc 46, 115. doi:10.1038/s41596-019-0154-6

14. Chen, L., Chen, J.-Y., Zhang, X., Gu, Y., Xiao, R., Shao, C., Tang, P., Qian, H., Luo, D., Li, H., Zhou, Y., Zhang, D.-E., Fu, X.-D., 2017. R-ChIP Using Inactive RNase H Reveals Dynamic Coupling of R-loops with Transcriptional Pausing at Gene Promoters. Mol Cell. doi:10.1016/j.molcel.2017.10.008

15. Chen, R., Yik, J.H.N., Lew, Q.J., Chao, S.-H., 2014. Brd4 and HEXIM1: multiple roles in P-TEFb regulation and cancer. Biomed Res Int 2014, 232870–11. doi:10.1155/2014/232870

16. Cheung, K.L., Zhang, F., Jaganathan, A., Sharma, R., Zhang, Q., Konuma, T., Shen, T., Lee, J.-Y., Ren, C., Chen, C.-H., Lu, G., Olson, M.R., Zhang, W., Kaplan, M.H., Littman, D.R., Walsh, M.J., Xiong, H., Zeng, L., Zhou, M.-M., 2017. Distinct Roles of Brd2 and Brd4 in Potentiating the Transcriptional Program for Th17 Cell Differentiation. Mol Cell 65, 1068–1080.e5. doi:10.1016/j.molcel.2016.12.022

17. Cimprich, K.A., Cortez, D., 2008. ATR: an essential regulator of genome integrity. Nat. Rev. Mol. Cell Biol. 9, 616–627. doi:10.1038/nrm2450

18. Costantino, L., Koshland, D., 2018. Genome-wide Map of R-Loop-Induced Damage Reveals How a Subset of R-Loops Contributes to Genomic Instability. Mol Cell 71, 487–497.e3. doi:10.1016/j.molcel.2018.06.037

19. Crossley, M.P., Bocek, M., Cimprich, K.A., 2019. R-Loops as Cellular Regulators and Genomic Threats. Mol Cell 73, 398–411. doi:10.1016/j.molcel.2019.01.024

20. Dawson, M.A., Prinjha, R.K., Dittmann, A., Giotopoulos, G., Bantscheff, M., Chan, W.-I., Robson, S.C., Chung, C.-W., Hopf, C., Savitski, M.M., Huthmacher, C., Gudgin, E., Lugo, D., Beinke, S., Chapman, T.D., Roberts, E.J., Soden, P.E., Auger, K.R., Mirguet, O., Doehner, K., Delwel, R., Burnett, A.K., Jeffrey, P., Drewes, G., Lee, K., Huntly, B.J.P., Kouzarides, T., 2011. Inhibition of BET recruitment to chromatin as an effective treatment for MLL-fusion leukaemia. Nature 478, 529–533. doi:10.1038/nature10509

21. Delmore, J.E., Issa, G.C., Lemieux, M.E., Rahl, P.B., Shi, J., Jacobs, H.M., Kastritis, E., Gilpatrick, T., Paranal, R.M., Qi, J., Chesi, M., Schinzel, A.C., McKeown, M.R., Heffernan, T.P., Vakoc, C.R., Bergsagel, P.L., Ghobrial, I.M., Richardson, P.G., Young, R.A., Hahn, W.C., Anderson, K.C., Kung, A.L., Bradner, J.E., Mitsiades, C.S., 2011. BET bromodomain inhibition as a therapeutic strategy to target c-Myc. Cell 146, 904–917. doi:10.1016/j.cell.2011.08.017

22. Drolet, M., Phoenix, P., Menzel, R., Massé, E., Liu, L.F., Crouch, R.J., 1995. Overexpression of RNase H partially complements the growth defect of an Escherichia coli delta topA mutant: R-loop formation is a major problem in the absence of DNA topoisomerase I. Proc. Natl. Acad. Sci. U.S.A. 92, 3526–3530. doi:10.1073/pnas.92.8.3526

23. Faivre, E.J., McDaniel, K.F., Albert, D.H., Mantena, S.R., Plotnik, J.P., Wilcox, D., Zhang, L., Bui, M.H., Sheppard, G.S., Wang, L., Sehgal, V., Lin, X., Huang, X., Lu, X., Uziel, T., Hessler, P., Lam, L.T., Bellin, R.J., Mehta, G., Fidanze, S., Pratt, J.K., Liu, D., Hasvold, L.A., Sun, C., Panchal, S.C., Nicolette, J.J., Fossey, S.L., Park, C.H., Longenecker, K., Bigelow, L., Torrent, M., Rosenberg, S.H., Kati, W.M., Shen, Y., 2020. Selective inhibition of the BD2 bromodomain of BET proteins in prostate cancer. Nature 578, 306–310. doi:10.1038/s41586-020-1930-8

24. Filippakopoulos, P., Picaud, S., Mangos, M., Keates, T., Lambert, J.-P., Barsyte-Lovejoy, D., Felletar, I., Volkmer, R., Müller, S., Pawson, T., Gingras, A.-C., Arrowsmith, C.H., Knapp, S., 2012. Histone recognition and large-scale structural analysis of the human bromodomain family. Cell 149, 214–231. doi:10.1016/j.cell.2012.02.013

25. Filippakopoulos, P., Qi, J., Picaud, S., Shen, Y., Smith, W.B., Fedorov, O., Morse, E.M., Keates, T., Hickman, T.T., Felletar, I., Philpott, M., Munro, S., McKeown, M.R., Wang, Y., Christie, A.L., West, N., Cameron, M.J., Schwartz, B., Heightman, T.D., La Thangue, N., French, C.A., Wiest, O., Kung, A.L., Knapp, S., Bradner, J.E., 2010a. Selective inhibition of BET bromodomains. Nature 468, 1067–1073. doi:10.1038/nature09504

26. Filippakopoulos, P., Qi, J., Picaud, S., Shen, Y., Smith, W.B., Fedorov, O., Morse, E.M., Keates, T., Hickman, T.T., Felletar, I., Philpott, M., Munro, S., McKeown, M.R., Wang, Y., Christie, A.L., West, N., Cameron, M.J., Schwartz, B., Heightman, T.D., La Thangue, N., French, C.A., Wiest, O., Kung, A.L., Knapp, S., Bradner, J.E., 2010b. Selective inhibition of BET bromodomains. Nature 468, 1067–1073. doi:10.1038/nature09504

27. Fiskus, W., Sharma, S., Qi, J., Valenta, J.A., Schaub, L.J., Shah, B., Peth, K., Portier, B.P., Rodriguez, M., Devaraj, S.G.T., Zhan, M., Sheng, J., Iyer, S.P., Bradner, J.E., Bhalla, K.N., 2014. Highly active combination of BRD4 antagonist and histone deacetylase inhibitor against human acute myelogenous leukemia cells. Mol. Cancer Ther. 13, 1142–1154. doi:10.1158/1535-7163.MCT-13-0770

28. Floyd, S.R., Pacold, M.E., Huang, Q., Clarke, S.M., Lam, F.C., Cannell, I.G., Bryson, B.D., Rameseder, J., Lee, M.J., Blake, E.J., Fydrych, A., Ho, R., Greenberger, B.A., Chen, G.C., Maffa, A., Del Rosario, A.M., Root, D.E., Carpenter, A.E., Hahn, W.C., Sabatini, D.M., Chen, C.C., White, F.M., Bradner, J.E., Yaffe, M.B., 2013a. The bromodomain protein Brd4 insulates chromatin from DNA damage signalling. Nature 498, 246–250. doi:10.1038/nature12147

29. Floyd, S.R., Pacold, M.E., Huang, Q., Clarke, S.M., Lam, F.C., Cannell, I.G., Bryson, B.D., Rameseder, J., Lee, M.J., Blake, E.J., Fydrych, A., Ho, R., Greenberger, B.A., Chen, G.C., Maffa, A., Del Rosario, A.M., Root, D.E., Carpenter, A.E., Hahn, W.C., Sabatini, D.M., Chen, C.C., White, F.M., Bradner, J.E., Yaffe, M.B., 2013b. The bromodomain protein Brd4 insulates chromatin from DNA damage signalling. Nature 498, 246–250. doi:10.1038/nature12147

30. Fowler, T., Ghatak, P., Price, D.H., Conaway, R., Conaway, J., Chiang, C.-M., Bradner, J.E., Shilatifard, A., Roy, A.L., 2014. Regulation of MYC expression and differential JQ1 sensitivity in cancer cells. PLoS ONE 9, e87003. doi:10.1371/journal.pone.0087003

31. Gaillard, H., Aguilera, A., 2016. Transcription as a Threat to Genome Integrity. Annu. Rev. Biochem. 85, 291–317. doi:10.1146/annurev-biochem-060815-014908

32. Gan, W., Guan, Z., Liu, J., Gui, T., Shen, K., Manley, J.L., Li, X., 2011. R-loop-mediated genomic instability is caused by impairment of replication fork progression. Genes Dev. 25, 2041–2056. doi:10.1101/gad.17010011

33. Garcia-Muse, T., Aguilera, A., 2019. R Loops: From Physiological to Pathological Roles. Cell 179, 604–618. doi:10.1016/j.cell.2019.08.055

34. Garcia-Muse, T., Aguilera, A., 2016. Transcription-replication conflicts: how they occur and how they are resolved. Nat. Rev. Mol. Cell Biol. 17, 553–563. doi:10.1038/nrm.2016.88

35. Grunseich, C., Wang, I.X., Watts, J.A., Burdick, J.T., Guber, R.D., Zhu, Z., Bruzel, A., Lanman, T., Chen, K., Schindler, A.B., Edwards, N., Ray-Chaudhury, A., Yao, J., Lehky, T., Piszczek, G., Crain, B., Fischbeck, K.H., Cheung, V.G., 2018. Senataxin Mutation Reveals How R-Loops Promote Transcription by Blocking DNA Methylation at Gene Promoters. Mol Cell 69, 426–437.e7. doi:10.1016/j.molcel.2017.12.030

36. Haberle, V., Stark, A., 2018. Eukaryotic core promoters and the functional basis of transcription initiation. Nature Publishing Group 19, 621–637. doi:10.1038/s41580- 018-0028-8

37. Hage, El, A., French, S.L., Beyer, A.L., Tollervey, D., 2010. Loss of Topoisomerase I leads to R-loop-mediated transcriptional blocks during ribosomal RNA synthesis. Genes Dev. 24, 1546–1558. doi:10.1101/gad.573310

38. Hamperl, S., Bocek, M.J., Saldivar, J.C., Swigut, T., Cimprich, K.A., 2017. Transcription-Replication Conflict Orientation Modulates R-Loop Levels and Activates Distinct DNA Damage Responses. Cell 170, 774–786.e19. doi:10.1016/j.cell.2017.07.043

39. Hamperl, S., Cimprich, K.A., 2016. Conflict Resolution in the Genome: How Transcription and Replication Make It Work 1–13. doi:10.1016/j.cell.2016.09.053

40. Hanahan, D., Weinberg, R.A., 2011. Hallmarks of cancer: the next generation. Cell 144, 646–674. doi:10.1016/j.cell.2011.02.013

41. Houzelstein, D., Bullock, S.L., Lynch, D.E., Grigorieva, E.F., Wilson, V.A., Beddington, R.S.P., 2002. Growth and early postimplantation defects in mice deficient for the bromodomain-containing protein Brd4. Mol. Cell. Biol. 22, 3794–3802. doi:10.1128/MCB.22.11.3794-3802.2002

42. Hsu, S.C., Gilgenast, T.G., Bartman, C.R., Edwards, C.R., Stonestrom, A.J., Huang, P., Emerson, D.J., Evans, P., Werner, M.T., Keller, C.A., Giardine, B., Hardison, R.C., Raj, A., Phillips-Cremins, J.E., Blobel, G.A., 2017. The BET Protein BRD2 Cooperates with CTCF to Enforce Transcriptional and Architectural Boundaries. Mol Cell 66, 102–116.e7. doi:10.1016/j.molcel.2017.02.027

43. Itzen, F., Greifenberg, A.K., Bösken, C.A., Geyer, M., 2014. Brd4 activates P-TEFb for RNA polymerase II CTD phosphorylation. Nucleic Acids Res 42, 7577–7590. doi:10.1093/nar/gku449

44. Jang, M.K., Mochizuki, K., Zhou, M., Jeong, H.-S., Brady, J.N., Ozato, K., 2005. The Bromodomain Protein Brd4 Is a Positive Regulatory Component of P-TEFb and Stimulates RNA Polymerase II-Dependent Transcription. Mol Cell 19, 523–534. doi:10.1016/j.molcel.2005.06.027

45. Kanno, T., Kanno, Y., LeRoy, G., Campos, E., Sun, H.-W., Brooks, S.R., Vahedi, G., Heightman, T.D., Garcia, B.A., Reinberg, D., Siebenlist, U., O’Shea, J.J., Ozato, K., 2014. BRD4 assists elongation of both coding and enhancer RNAs by interacting with acetylated histones. Nat Struct Mol Biol 21, 1047–1057. doi:10.1038/nsmb.2912

46. Kim, J.J., Lee, S.Y., Gong, F., Battenhouse, A.M., Boutz, D.R., Bashyal, A., Refvik, S.T., Chiang, C.-M., Xhemalce, B., Paull, T.T., Brodbelt, J.S., Marcotte, E.M., Miller, K.M., 2019. Systematic bromodomain protein screens identify homologous recombination and R-loop suppression pathways involved in genome integrity. Genes Dev. doi:10.1101/gad.331231.119

47. Komarnitsky, P., Cho, E.J., Buratowski, S., 2000. Different phosphorylated forms of RNA polymerase II and associated mRNA processing factors during transcription. Genes Dev. 14, 2452–2460. doi:10.1101/gad.824700

48. Kotsantis, P., Silva, L.M., Irmscher, S., Jones, R.M., Folkes, L., Gromak, N., Petermann, E., 2016a. Increased global transcription activity as a mechanism of replication stress in cancer. Nat Commun 7, 13087. doi:10.1038/ncomms13087

49. Kotsantis, P., Silva, L.M., Irmscher, S., Jones, R.M., Folkes, L., Gromak, N., Petermann, E., 2016b. Increased global transcription activity as a mechanism of replication stress in cancer. Nat Commun 7, 13087. doi:10.1038/ncomms13087

50. Krueger, B.J., Varzavand, K., Cooper, J.J., Price, D.H., 2010. The mechanism of release of P-TEFb and HEXIM1 from the 7SK snRNP by viral and cellular activators includes a conformational change in 7SK. PLoS ONE 5, e12335. doi:10.1371/journal.pone.0012335

51. LeRoy, G., Rickards, B., Flint, S.J., 2008. The double bromodomain proteins Brd2 and Brd3 couple histone acetylation to transcription. Mol Cell 30, 51–60. doi:10.1016/j.molcel.2008.01.018

52. Li, X., Manley, J.L., 2005. Inactivation of the SR protein splicing factor ASF/SF2 results in genomic instability. Cell 122, 365–378. doi:10.1016/j.cell.2005.06.008

53. Lin, C.Y., Lovén, J., Rahl, P.B., Paranal, R.M., Burge, C.B., Bradner, J.E., Lee, T.I., Young, R.A., 2012. Transcriptional amplification in tumor cells with elevated c-Myc. Cell 151, 56–67. doi:10.1016/j.cell.2012.08.026

54. Liu, W., Ma, Q., Wong, K., Li, W., Ohgi, K., Zhang, J., Aggarwal, A., Rosenfeld, M.G., 2013. Brd4 and JMJD6-associated anti-pause enhancers in regulation of transcriptional pause release. Cell 155, 1581–1595. doi:10.1016/j.cell.2013.10.056

55. Maruyama, T., Farina, A., Dey, A., Cheong, J., Bermudez, V.P., Tamura, T., Sciortino, S., Shuman, J., Hurwitz, J., Ozato, K., 2002a. A Mammalian bromodomain protein, brd4, interacts with replication factor C and inhibits progression to S phase. Mol. Cell. Biol. 22, 6509–6520. doi:10.1128/MCB.22.18.6509-6520.2002

56. Maruyama, T., Farina, A., Dey, A., Cheong, J., Bermudez, V.P., Tamura, T., Sciortino, S., Shuman, J., Hurwitz, J., Ozato, K., 2002b. A Mammalian bromodomain protein, brd4, interacts with replication factor C and inhibits progression to S phase. Mol. Cell. Biol. 22, 6509–6520. doi:10.1128/mcb.22.18.6509-6520.2002

57. Massé, E., Phoenix, P., Drolet, M., 1997. DNA topoisomerases regulate R-loop formation during transcription of the rrnB operon in Escherichia coli. J Biol Chem 272, 12816–12823. doi:10.1074/jbc.272.19.12816

58. Matos, D.A., Zhang, J.-M., Ouyang, J., Nguyen, H.D., Genois, M.-M., Zou, L., 2019. ATR Protects the Genome against R Loops through a MUS81-Triggered Feedback Loop. Mol Cell. doi:10.1016/j.molcel.2019.10.010

59. Mertz, J.A., Conery, A.R., Bryant, B.M., Sandy, P., Balasubramanian, S., Mele, D.A., Bergeron, L., Sims, R.J., 2011. Targeting MYC dependence in cancer by inhibiting BET bromodomains. Proc. Natl. Acad. Sci. U.S.A. 108, 16669–16674. doi:10.1073/pnas.1108190108

60. Morales, J.C., Richard, P., Patidar, P.L., Motea, E.A., Dang, T.T., Manley, J.L., Boothman, D.A., 2016. XRN2 Links Transcription Termination to DNA Damage and Replication Stress. PLoS Genet 12, e1006107.

61. Muhar, M., Ebert, A., Neumann, T., Umkehrer, C., Jude, J., Wieshofer, C., Rescheneder, P., Lipp, J.J., Herzog, V.A., Reichholf, B., Cisneros, D.A., Hoffmann, T., Schlapansky, M.F., Bhat, P., Haeseler, von, A., Köcher, T., Obenauf, A.C., Popow, J., Ameres, S.L., Zuber, J., 2018. SLAM-seq defines direct gene-regulatory functions of the BRD4-MYC axis. Science 360, 800–805. doi:10.1126/science.aao2793

62. Nguyen, H.D., Yadav, T., Giri, S., Saez, B., Graubert, T.A., Zou, L., 2017. Functions of Replication Protein A as a Sensor of R Loops and a Regulator of RNaseH1. Mol Cell 65, 832–847.e4. doi:10.1016/j.molcel.2017.01.029

63. Odore, E., Lokiec, F., Cvitkovic, E., Bekradda, M., Herait, P., Bourdel, F., Kahatt, C., Raffoux, E., Stathis, A., Thieblemont, C., Quesnel, B., Cunningham, D., Riveiro, M.E., Rezaï, K., 2015. Phase I Population Pharmacokinetic Assessment of the Oral Bromodomain Inhibitor OTX015 in Patients with Haematologic Malignancies. Clinical Pharmacokinetics 55, 397–405. doi:10.1007/s40262-015-0327-6

64. Olson, E., Nievera, C.J., Klimovich, V., Fanning, E., Wu, X., 2006. RPA2 is a direct downstream target for ATR to regulate the S-phase checkpoint. J Biol Chem 281, 39517–39533. doi:10.1074/jbc.M605121200

65. Ozer, H.G., El-Gamal, D., Powell, B., Hing, Z.A., Blachly, J.S., Harrington, B., Mitchell, S., Grieselhuber, N.R., Williams, K., Lai, T.-H., Alinari, L., Baiocchi, R.A., Brinton, L., Baskin, E., Cannon, M., Beaver, L., Goettl, V.M., Lucas, D.M., Woyach, J.A., Sampath, D., Lehman, A.M., Yu, L., Zhang, J., Ma, Y., Zhang, Y., Spevak, W., Shi, S., Severson, P., Shellooe, R., Carias, H., Tsang, G., Dong, K., Ewing, T., Marimuthu, A., Tantoy, C., Walters, J., Sanftner, L., Rezaei, H., Nespi, M., Matusow, B., Habets, G., Ibrahim, P., Zhang, C., Mathé, E.A., Bollag, G., Byrd, J.C., Lapalombella, R., 2018. BRD4 Profiling Identifies Critical Chronic Lymphocytic Leukemia Oncogenic Circuits and Reveals Sensitivity to PLX51107, a Novel Structurally Distinct BET Inhibitor. Cancer Discov 8, 458–477. doi:10.1158/2159-8290.CD-17-0902

66. Parajuli, S., Tealsey, D.C., Murali, B., Jackson, J., Vindigni, A., Stewart, S.A., 2017. Human Ribonuclease H1 resolves R loops and thereby enables progression of the DNA replication fork. J Biol Chem jbc.M117.787473–19. doi:10.1074/jbc.M117.787473

67. Patel, M.C., Debrosse, M., Smith, M., Dey, A., Huynh, W., Sarai, N., Heightman, T.D., Tamura, T., Ozato, K., 2013. BRD4 coordinates recruitment of pause release factor P-TEFb and the pausing complex NELF/DSIF to regulate transcription elongation of interferon-stimulated genes. Mol. Cell. Biol. 33, 2497–2507. doi:10.1128/MCB.01180-12

68. Pericole, F.V., Lazarini, M., de Paiva, L.B., Duarte, A.D.S.S., Vieira Ferro, K.P., Niemann, F.S., Roversi, F.M., Olalla Saad, S.T., 2019. BRD4 Inhibition Enhances Azacitidine Efficacy in Acute Myeloid Leukemia and Myelodysplastic Syndromes. Front Oncol 9, 16. doi:10.3389/fonc.2019.00016

69. Piha-Paul, S.A., Sachdev, J.C., Barve, M., LoRusso, P., Szmulewitz, R., Patel, S.P., Lara, P.N., Chen, X., Hu, B., Freise, K.J., Modi, D., Sood, A., Hutti, J.E., Wolff, J., O’Neil, B.H., 2019. First-in-Human Study of Mivebresib (ABBV-075), an Oral Pan-Inhibitor of Bromodomain and Extra Terminal Proteins, in Patients with Relapsed/Refractory Solid Tumors. Clin. Cancer Res. 25, 6309–6319. doi:10.1158/1078-0432.CCR-19-0578

70. Pivot-Pajot, C., Caron, C., Govin, J., Vion, A., Rousseaux, S., Khochbin, S., 2003. Acetylation-dependent chromatin reorganization by BRDT, a testis-specific bromodomain-containing protein. Mol. Cell. Biol. 23, 5354–5365. doi:10.1128/mcb.23.15.5354-5365.2003

71. Porrua, O., Libri, D., 2015. Transcription termination and the control of the transcriptome: why, where and how to stop. Nature Publishing Group 16, 190–202. doi:10.1038/nrm3943

72. Puccetti, M.V., Adams, C.M., Kushinsky, S., Eischen, C.M., 2019. Smarcal1 and Zranb3 Protect Replication Forks from Myc-Induced DNA Replication Stress. Cancer Res 79, 1612–1623. doi:10.1158/0008-5472.CAN-18-2705

73. Quinet, A., Carvajal-Maldonado, D., Lemacon, D., Vindigni, A., 2017. DNA Fiber Analysis: Mind the Gap! Meth. Enzymol. 591, 55–82. doi:10.1016/bs.mie.2017.03.019

74. Rahman, S., Sowa, M.E., Ottinger, M., Smith, J.A., Shi, Y., Harper, J.W., Howley, P.M., 2011. The Brd4 extraterminal domain confers transcription activation independent of pTEFb by recruiting multiple proteins, including NSD3. Mol. Cell. Biol. 31, 2641– 2652. doi:10.1128/MCB.01341-10

75. Rathert, P., Roth, M., Neumann, T., Muerdter, F., Roe, J.-S., Muhar, M., Deswal, S., Cerny-Reiterer, S., Peter, B., Jude, J., Hoffmann, T., Boryń, Ł.M., Axelsson, E., Schweifer, N., Tontsch-Grunt, U., Dow, L.E., Gianni, D., Pearson, M., Valent, P., Stark, A., Kraut, N., Vakoc, C.R., Zuber, J., 2015. Transcriptional plasticity promotes primary and acquired resistance to BET inhibition. Nature 525, 543–547. doi:10.1038/nature14898

76. Riccardi, C., Nicoletti, I., 2006. Analysis of apoptosis by propidium iodide staining and flow cytometry. Nat. Protocols 1, 1458–1461. doi:10.1038/nprot.2006.238

77. Richard, P., Manley, J.L., 2016. R Loops and Links to Human Disease. J. Mol. Biol. doi:10.1016/j.jmb.2016.08.031

78. Rogakou, E.P., Pilch, D.R., Orr, A.H., Ivanova, V.S., Bonner, W.M., 1998. DNA double-stranded breaks induce histone H2AX phosphorylation on serine 139. J Biol Chem 273, 5858–5868. doi:10.1074/jbc.273.10.5858

79. Santos-Pereira, J.M., Aguilera, A., 2015. R loops: new modulators of genome dynamics and function. Nature Reviews Genetics 16, 583–597. doi:10.1038/nrg3961

80. Schröder, S., Cho, S., Zeng, L., Zhang, Q., Kaehlcke, K., Mak, L., Lau, J., Bisgrove, D., Schnölzer, M., Verdin, E., Zhou, M.-M., Ott, M., 2012. Two-pronged binding with bromodomain-containing protein 4 liberates positive transcription elongation factor b from inactive ribonucleoprotein complexes. J Biol Chem 287, 1090–1099. doi:10.1074/jbc.M111.282855

81. Schwab, R.A., Nieminuszczy, J., Shah, F., Langton, J., Lopez Martinez, D., Liang, C.-C., Cohn, M.A., Gibbons, R.J., Deans, A.J., Niedzwiedz, W., 2015. The Fanconi Anemia Pathway Maintains Genome Stability by Coordinating Replication and Transcription. Mol Cell 60, 351–361. doi:10.1016/j.molcel.2015.09.012

82. Shao, W., Zeitlinger, J., 2017a. Paused RNA polymerase II inhibits new transcriptional initiation. Nature Publishing Group 49, 1045–1051. doi:10.1038/ng.3867

83. Shao, W., Zeitlinger, J., 2017b. Paused RNA polymerase II inhibits new transcriptional initiation. Nature Publishing Group 49, 1045–1051. doi:10.1038/ng.3867

84. Shivji, M.K.K., Renaudin, X., Williams, Ç.H., Venkitaraman, A.R., 2018. BRCA2 Regulates Transcription Elongation by RNA Polymerase II to Prevent R-Loop Accumulation. Cell Reports 22, 1031–1039. doi:10.1016/j.celrep.2017.12.086

85. Skourti-Stathaki, K., Proudfoot, N.J., 2014. A double-edged sword: R loops as threats to genome integrity and powerful regulators of gene expression. Genes Dev. 28, 1384–1396. doi:10.1101/gad.242990.114

86. Skourti-Stathaki, K., Proudfoot, N.J., Gromak, N., 2011. Human senataxin resolves RNA/DNA hybrids formed at transcriptional pause sites to promote Xrn2-dependent termination. Mol Cell 42, 794–805. doi:10.1016/j.molcel.2011.04.026

87. Sollier, J., Cimprich, K.A., 2015. Breaking bad: R-loops and genome integrity. Trends in Cell Biology 25, 514–522. doi:10.1016/j.tcb.2015.05.003

88. Sollier, J., Stork, C.T., García-Rubio, M.L., Paulsen, R.D., Aguilera, A., Cimprich, K.A., 2014. Transcription-Coupled Nucleotide Excision Repair Factors Promote R-Loop-Induced Genome Instability. Mol Cell 56, 777–785. doi:10.1016/j.molcel.2014.10.020

89. Stork, C.T., Bocek, M., Crossley, M.P., Sollier, J., Sanz, L.A., Chédin, F., Swigut, T., Cimprich, K.A., 2016. Co-transcriptional R-loops are the main cause of estrogen-induced DNA damage. eLife 5, e17548. doi:10.7554/eLife.17548

90. Stuckey, R., García-Rodríguez, N., Aguilera, A., Wellinger, R.E., 2015. Role for RNA:DNA hybrids in origin-independent replication priming in a eukaryotic system. Proc. Natl. Acad. Sci. U.S.A. 112, 5779–5784. doi:10.1073/pnas.1501769112

91. Sun, C., Yin, J., Fang, Y., Chen, J., Jeong, K.J., Chen, X., Vellano, C.P., Ju, Z., Zhao, W., Zhang, D., Lu, Y., Meric-Bernstam, F., Yap, T.A., Hattersley, M., O’Connor, M.J., Chen, H., Fawell, S., Lin, S.-Y., Peng, G., Mills, G.B., 2018. BRD4 Inhibition Is Synthetic Lethal with PARP Inhibitors through the Induction of Homologous Recombination Deficiency. Cancer Cell 33, 401–416.e8. doi:10.1016/j.ccell.2018.01.019

92. Wahba, L., Amon, J.D., Koshland, D., Vuica-Ross, M., 2011. RNase H and Multiple RNA Biogenesis Factors Cooperate to Prevent RNA:DNA Hybrids from Generating Genome Instability. MOLCEL 44, 978–988. doi:10.1016/j.molcel.2011.10.017

93. Wessel, S.R., Mohni, K.N., Luzwick, J.W., Dungrawala, H., Cortez, D., 2019a. Functional Analysis of the Replication Fork Proteome Identifies BET Proteins as PCNA Regulators. Cell Reports 28, 3497–3509.e4. doi:10.1016/j.celrep.2019.08.051

94. Wessel, S.R., Mohni, K.N., Luzwick, J.W., Dungrawala, H., Cortez, D., 2019b. Functional Analysis of the Replication Fork Proteome Identifies BET Proteins as PCNA Regulators. Cell Reports 28, 3497–3509.e4. doi:10.1016/j.celrep.2019.08.051

95. Winter, G.E., Mayer, A., Buckley, D.L., Erb, M.A., Roderick, J.E., Vittori, S., Reyes, J.M., di Iulio, J., Souza, A., Ott, C.J., Roberts, J.M., Zeid, R., Scott, T.G., Paulk, J., Lachance, K., Olson, C.M., Dastjerdi, S., Bauer, S., Lin, C.Y., Gray, N.S., Kelliher, M.A., Churchman, L.S., Bradner, J.E., 2017. BET Bromodomain Proteins Function as Master Transcription Elongation Factors Independent of CDK9 Recruitment. Mol Cell. doi:10.1016/j.molcel.2017.06.004

96. Xiao, Y., Luo, M., Hayes, R.P., Kim, J., Ng, S., Ding, F., Liao, M., Ke, A., 2017. Structure Basis for Directional R-loop Formation and Substrate Handover Mechanisms in Type I CRISPR-Cas System. Cell 170, 48–60.e11. doi:10.1016/j.cell.2017.06.012

97. Zatreanu, D., Han, Z., Mitter, R., Tumini, E., Williams, H., Gregersen, L., Dirac-Svejstrup, A.B., Roma, S., Stewart, A., Aguilera, A., Svejstrup, J.Q., 2019. Elongation Factor TFIIS Prevents Transcription Stress and R-Loop Accumulation to Maintain Genome Stability. MOLCEL 76, 57–69.e9. doi:10.1016/j.molcel.2019.07.037

98. Zeman, M.K., Cimprich, K.A., 2014. Causes and consequences of replication stress. Nat. Cell Biol. 16, 2–9. doi:10.1038/ncb2897

99. Zhang, J., Dulak, A.M., Hattersley, M.M., Willis, B.S., Nikkilä, J., Wang, A., Lau, A., Reimer, C., Zinda, M., Fawell, S.E., Mills, G.B., Chen, H., 2018. BRD4 facilitates replication stress-induced DNA damage response. Oncogene 37, 3763–3777. doi:10.1038/s41388-018-0194-3

100. Zhang, W., Prakash, C., Sum, C., Gong, Y., Li, Y., Kwok, J.J.T., Thiessen, N., Pettersson, S., Jones, S.J.M., Knapp, S., Yang, H., Chin, K.-C., 2012. Bromodomain-containing protein 4 (BRD4) regulates RNA polymerase II serine 2 phosphorylation in human CD4+ T cells. J Biol Chem 287, 43137–43155. doi:10.1074/jbc.M112.413047

101. Zhou, Y., Zhou, J., Lu, X., Tan, T.-Z., Chng, W.-J., 2018. BET Bromodomain inhibition promotes De-repression of TXNIP and activation of ASK1-MAPK pathway in acute myeloid leukemia. BMC Cancer 18, 731–11. doi:10.1186/s12885-018-4661-6

102. Zuber, J., Shi, J., Wang, E., Rappaport, A.R., Herrmann, H., Sison, E.A., Magoon, D., Qi, J., Blatt, K., Wunderlich, M., Taylor, M.J., Johns, C., Chicas, A., Mulloy, J.C., Kogan, S.C., Brown, P., Valent, P., Bradner, J.E., Lowe, S.W., Vakoc, C.R., 2011. RNAi screen identifies Brd4 as a therapeutic target in acute myeloid leukaemia. Nature 478, 524–528. doi:10.1038/nature10334

