## Supplementary material for "BRD4 Prevents R-Loop Formation and Transcription-Replication Conflicts by Ensuring Efficient Transcription Elongation": Source Data: 20191112_HeLa_dBET6.pptx

### Slide 1
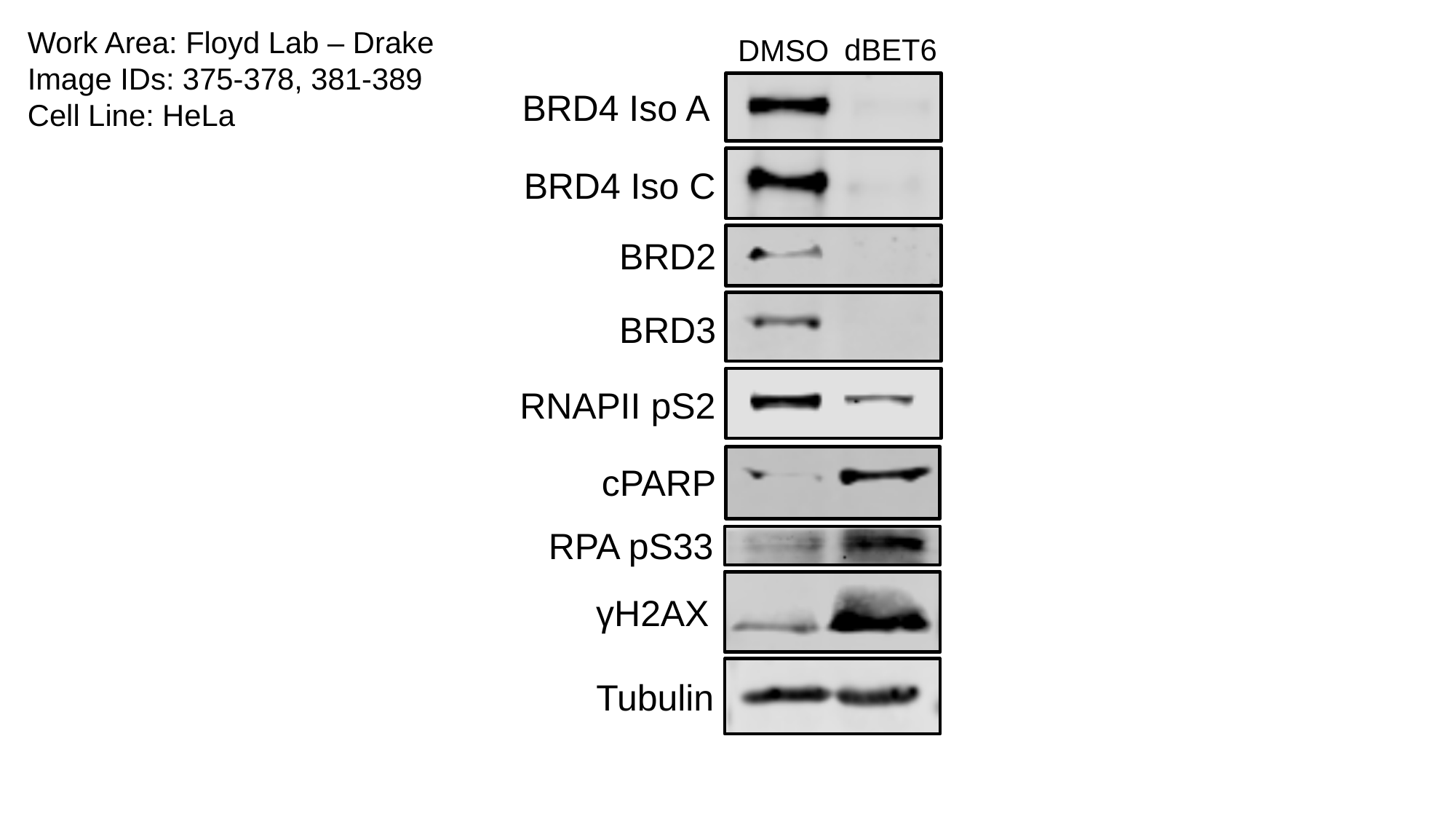

Work Area: Floyd Lab – Drake
Image IDs: 375-378, 381-389
Cell Line: HeLa
dBET6
DMSO
BRD4 Iso A
BRD4 Iso C
BRD2
BRD3
RNAPII pS2
cPARP
RPA pS33
γH2AX
Tubulin

### Slide 2
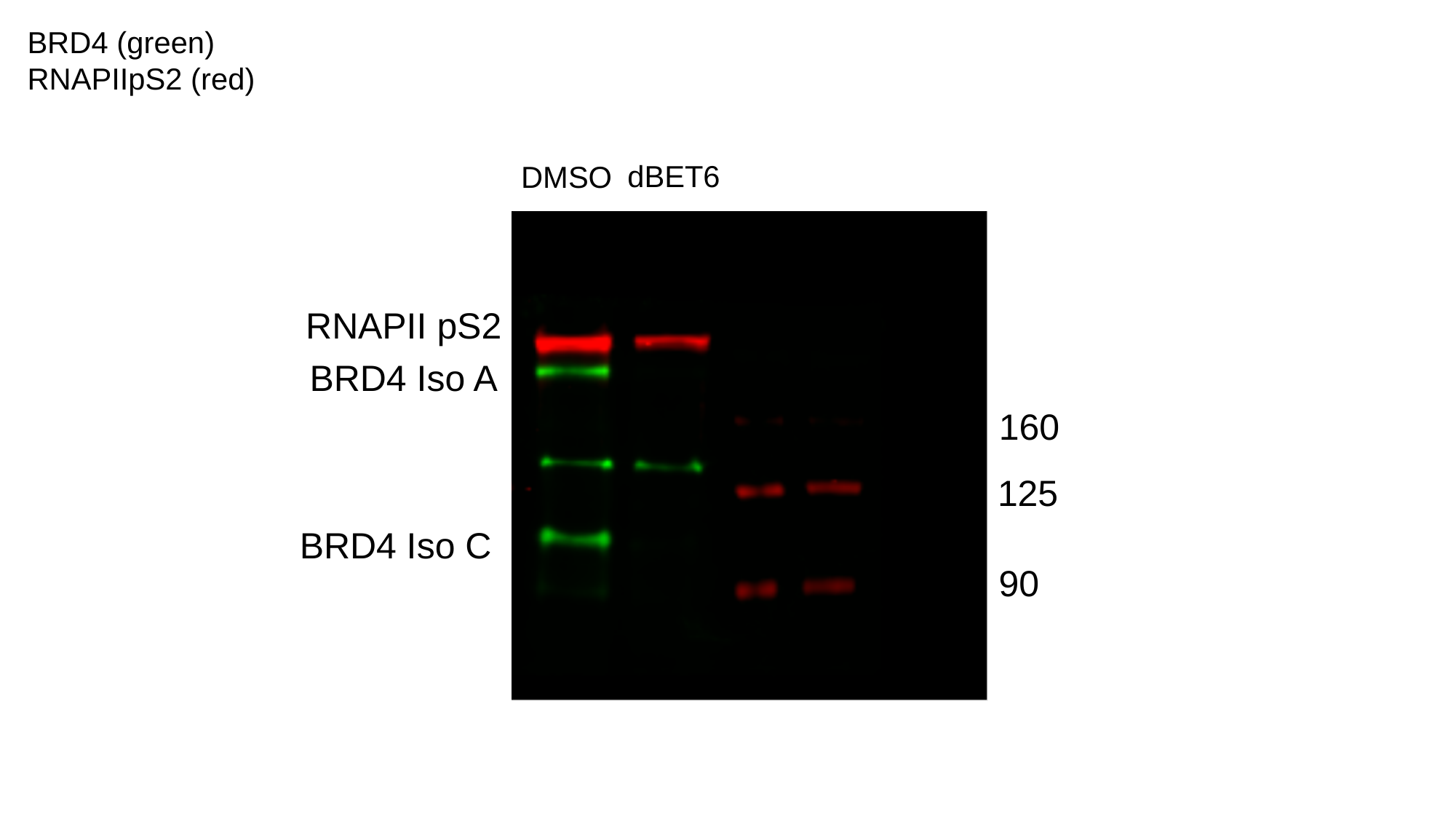

BRD4 (green)
RNAPIIpS2 (red)
dBET6
DMSO
RNAPII pS2
BRD4 Iso A
160
125
BRD4 Iso C
90

### Slide 3
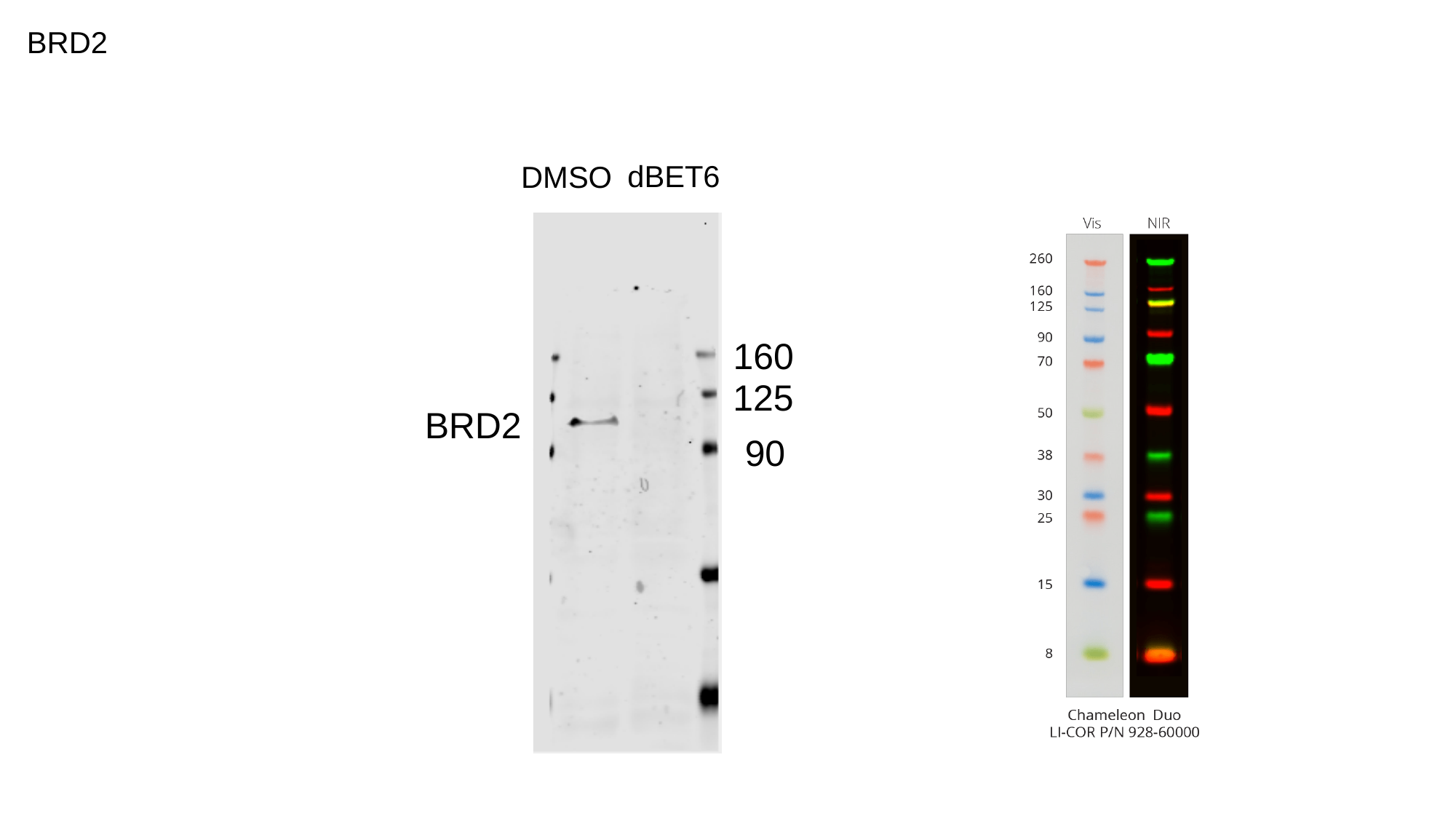

BRD2
dBET6
DMSO
160
125
BRD2
90

### Slide 4
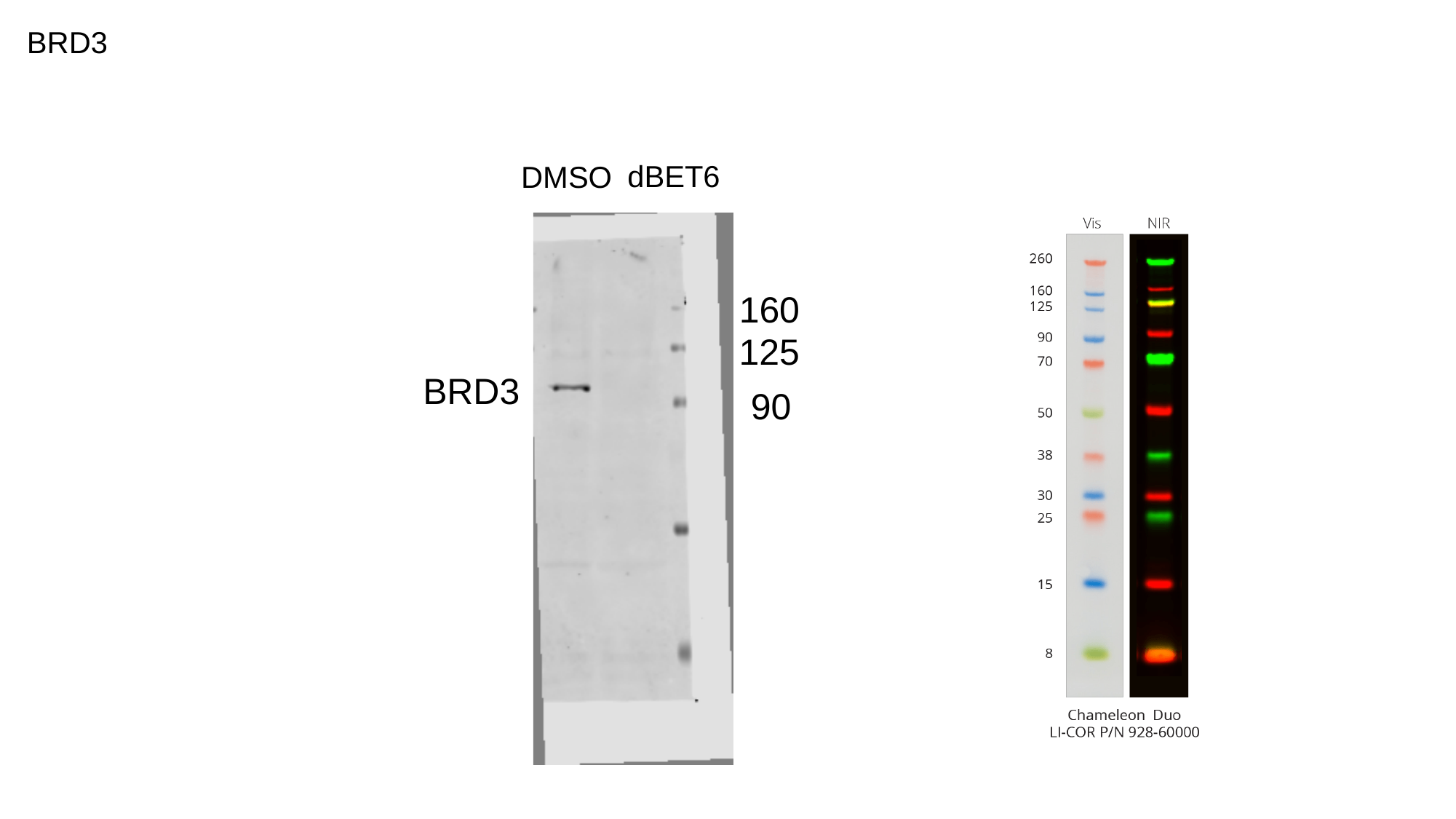

BRD3
dBET6
DMSO
160
125
BRD3
90

### Slide 5
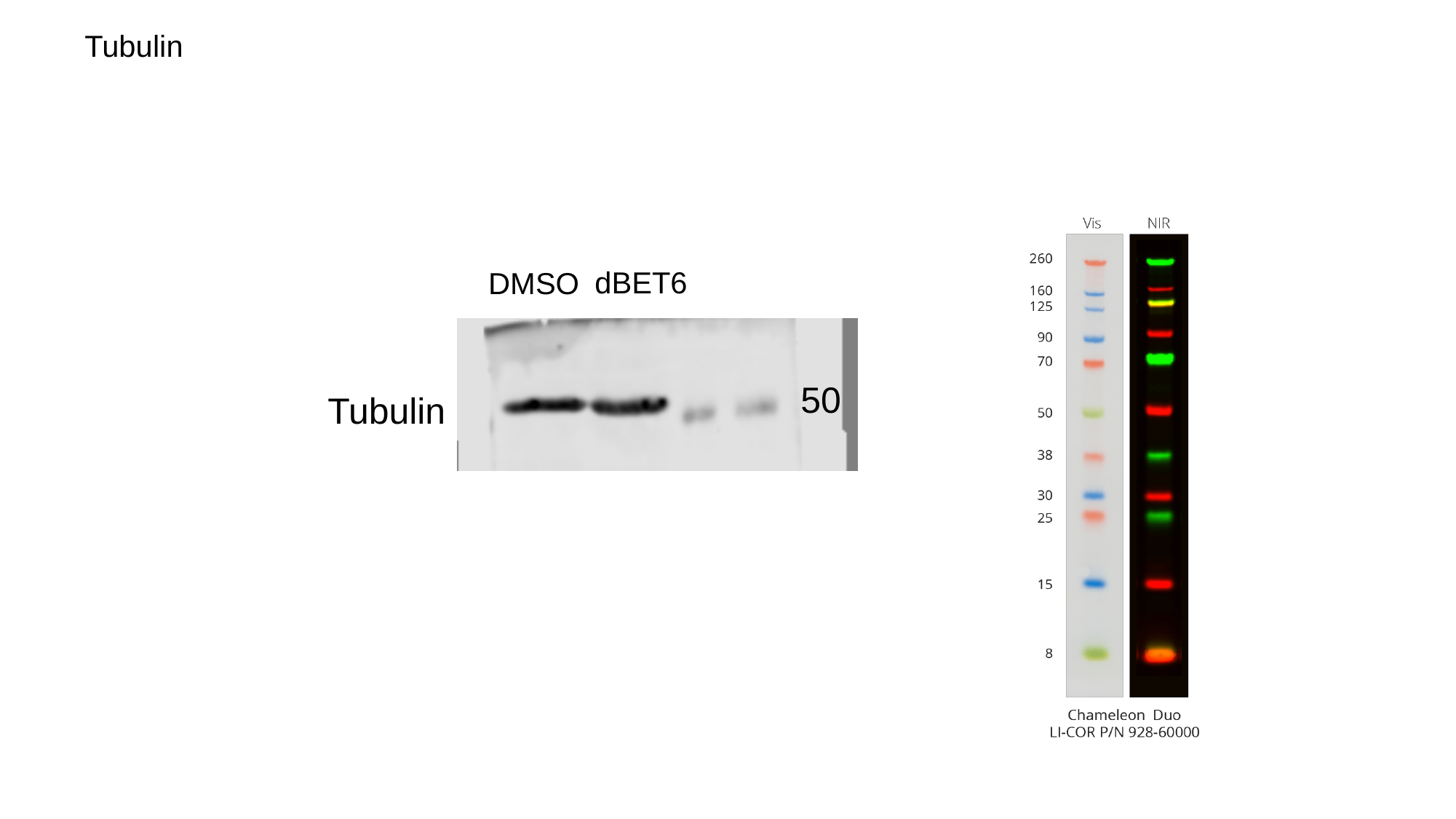

Tubulin
dBET6
DMSO
50
Tubulin

### Slide 6
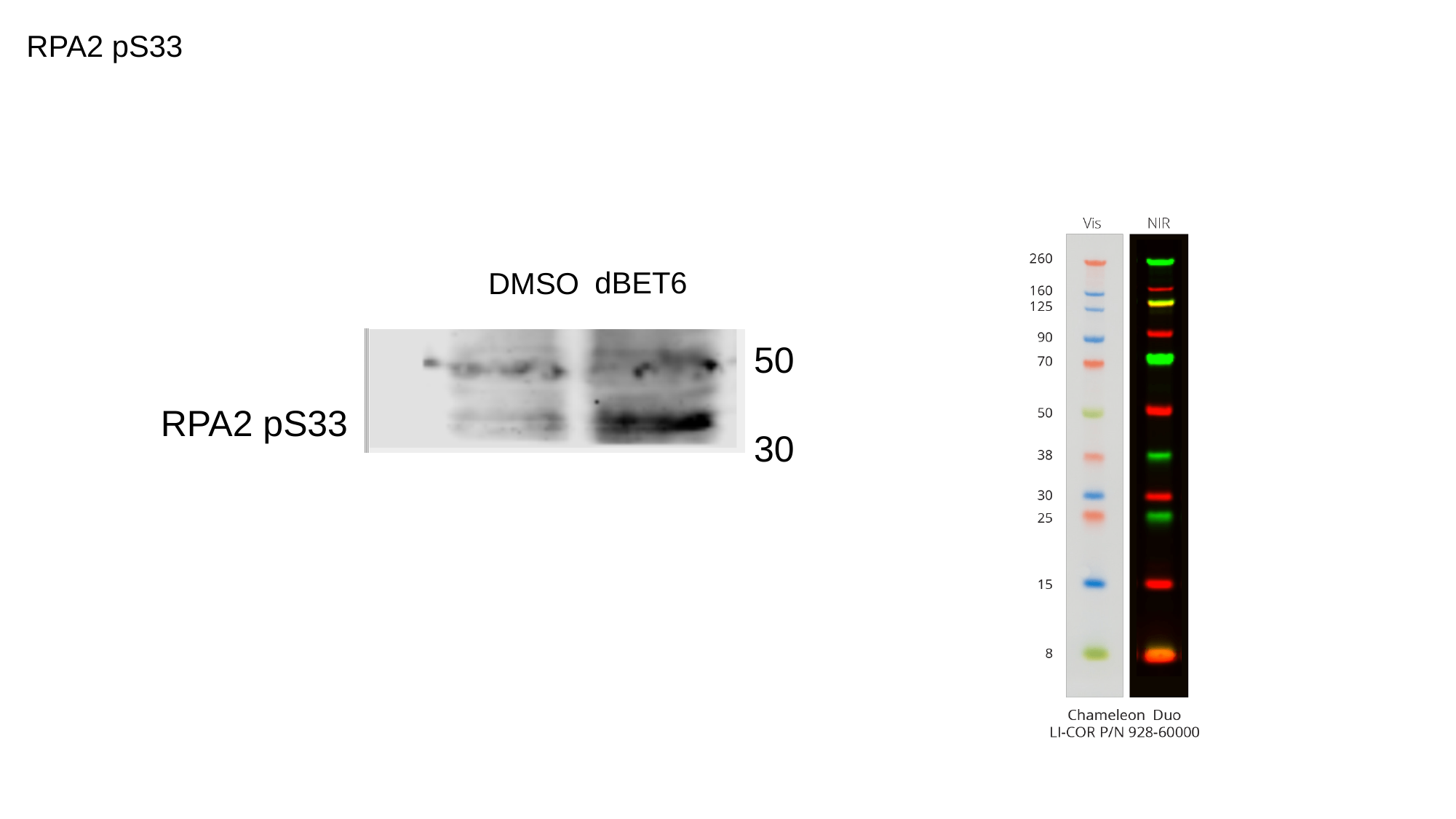

RPA2 pS33
dBET6
DMSO
50
RPA2 pS33
30

### Slide 7
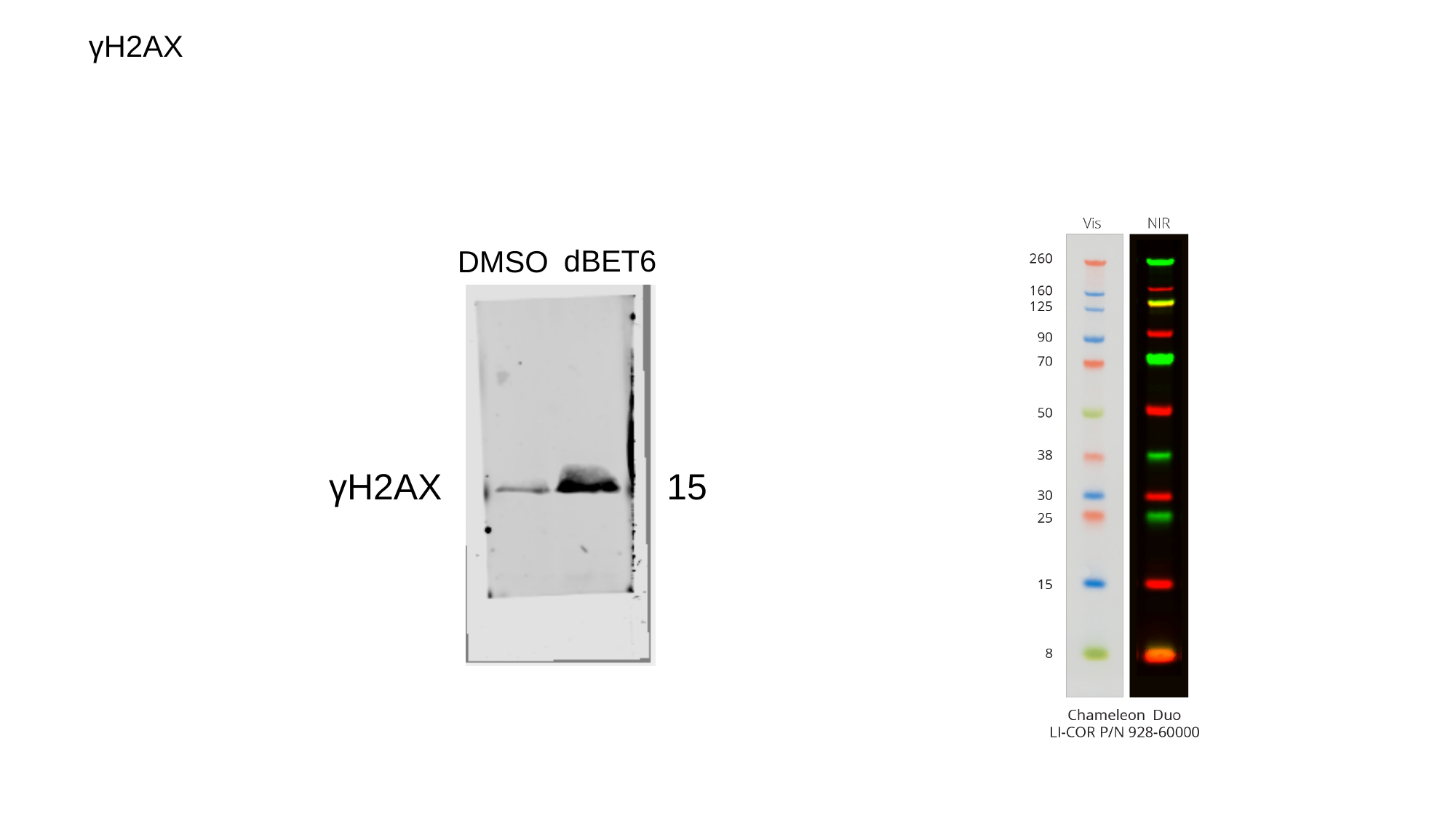

γH2AX
dBET6
DMSO
γH2AX
15

### Slide 8
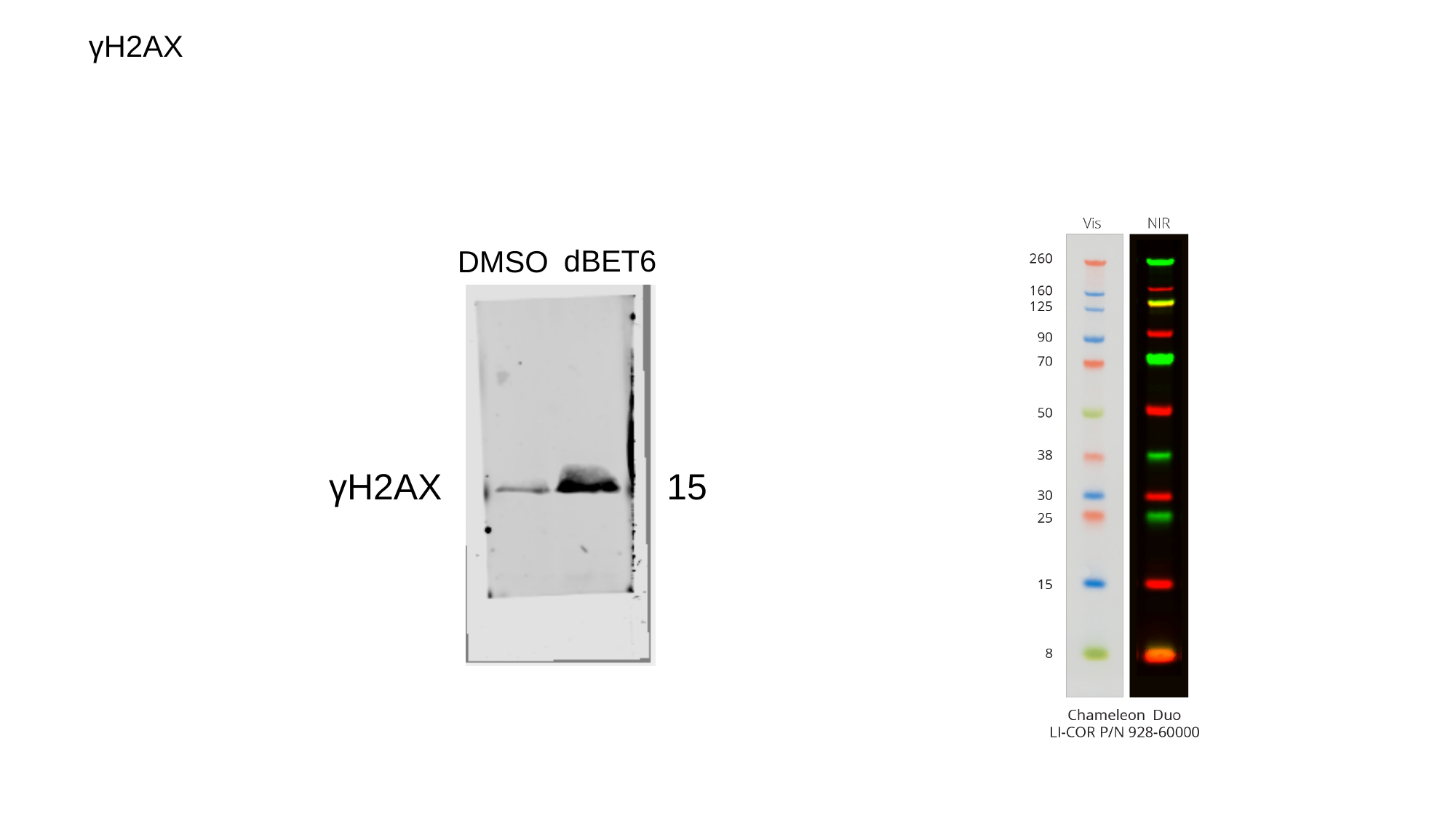

γH2AX
dBET6
DMSO
γH2AX
15

### Slide 9
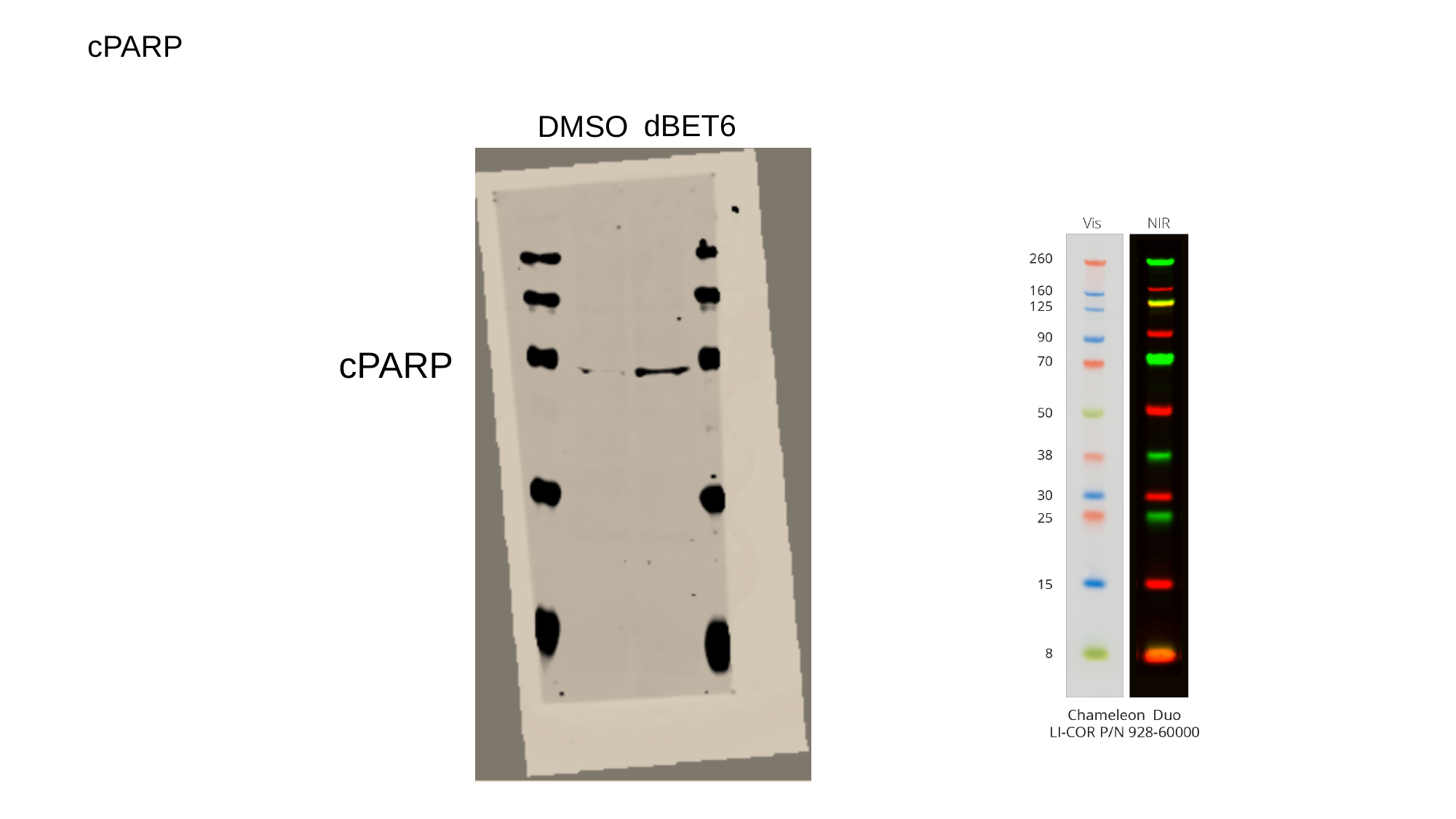

cPARP
dBET6
DMSO
cPARP
