## Supplementary material for "BRD4 Prevents R-Loop Formation and Transcription-Replication Conflicts by Ensuring Efficient Transcription Elongation": Source Data: 20171024.pptx

### Slide 1
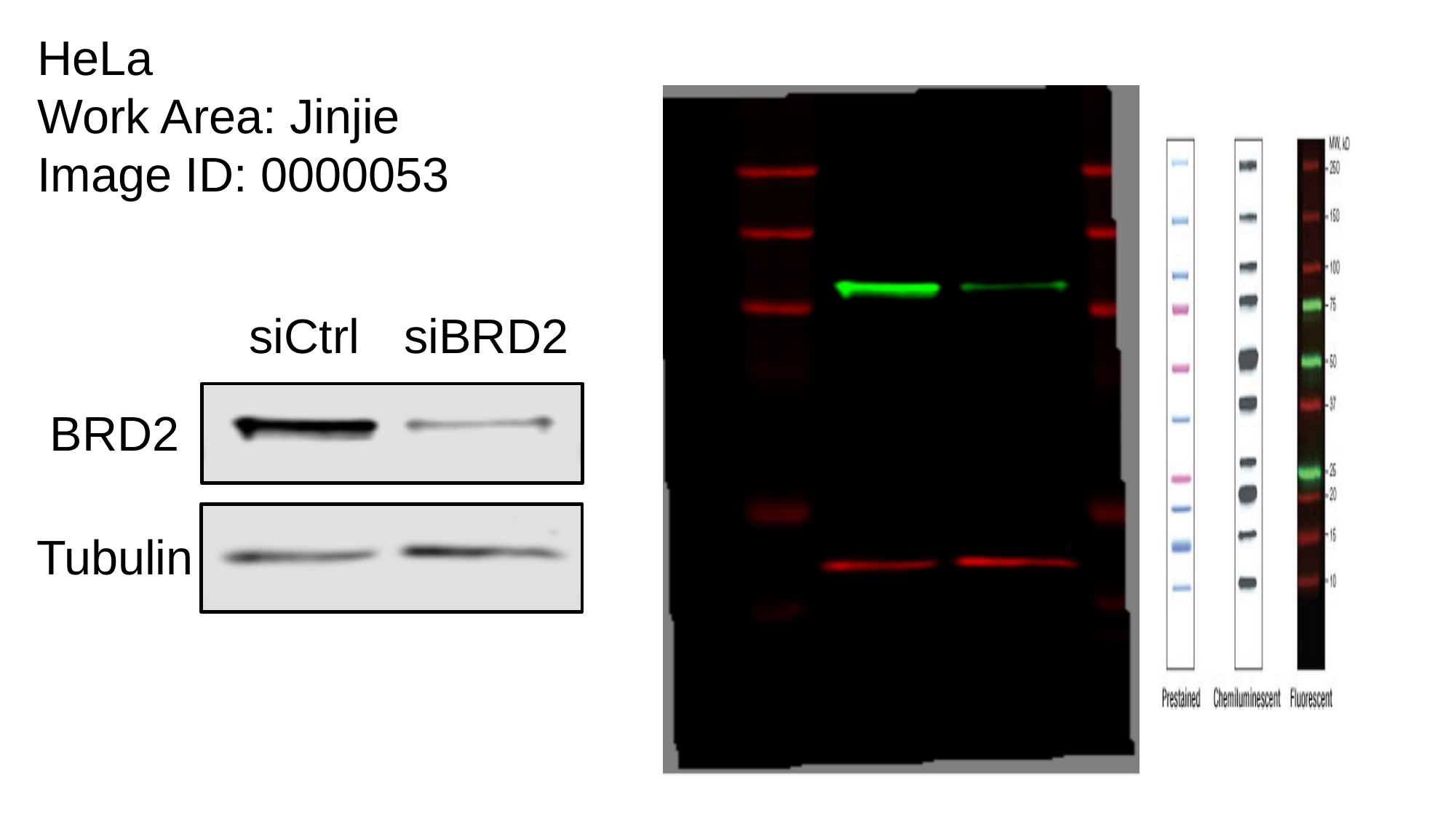

HeLa
Work Area: Jinjie
Image ID: 0000053
siBRD2
siCtrl
BRD2
Tubulin

### Slide 2
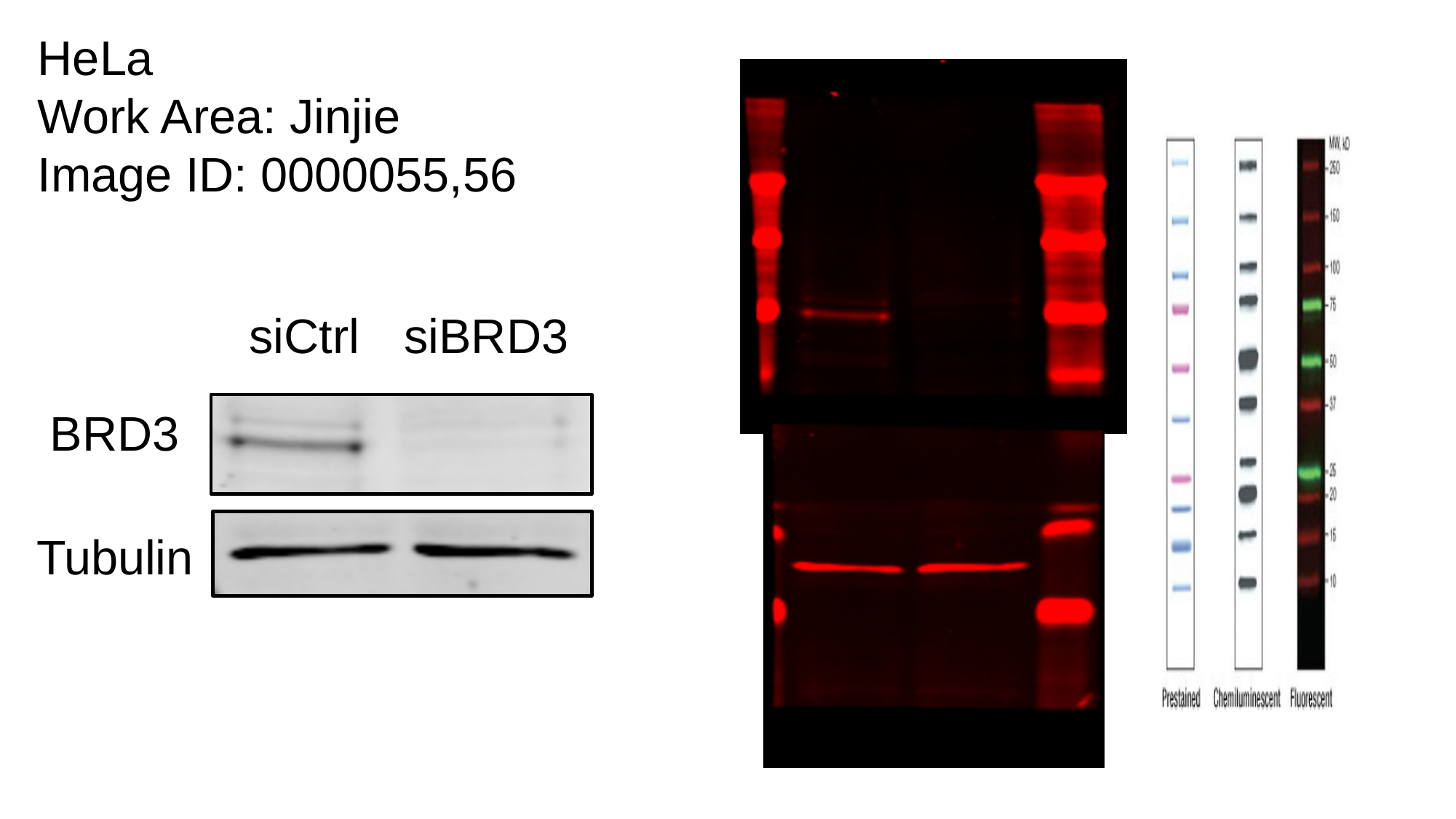

HeLa
Work Area: Jinjie
Image ID: 0000055,56
siBRD3
siCtrl
BRD3
Tubulin

### Slide 3
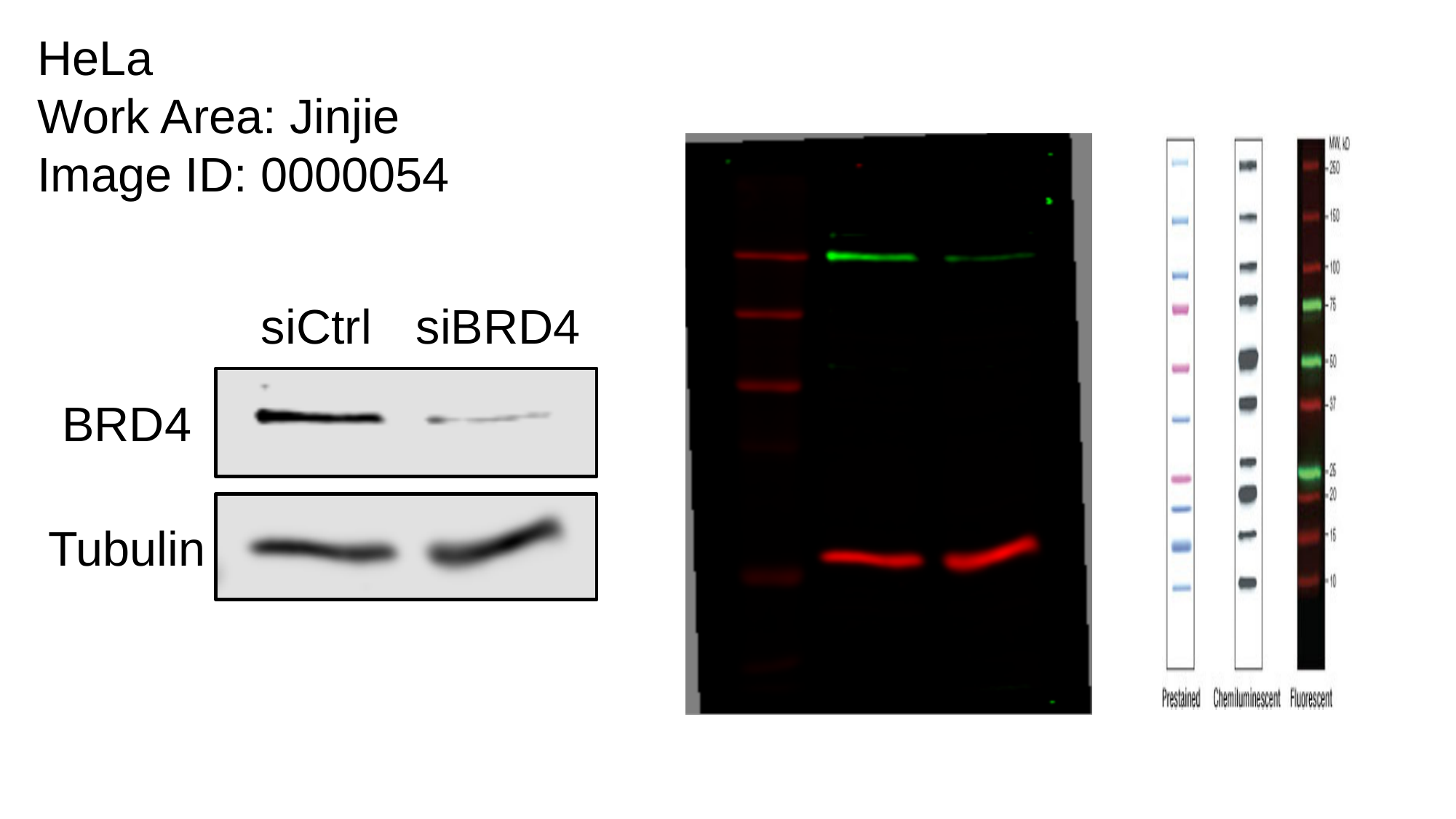

HeLa
Work Area: Jinjie
Image ID: 0000054
siBRD4
siCtrl
BRD4
Tubulin

### Slide 4
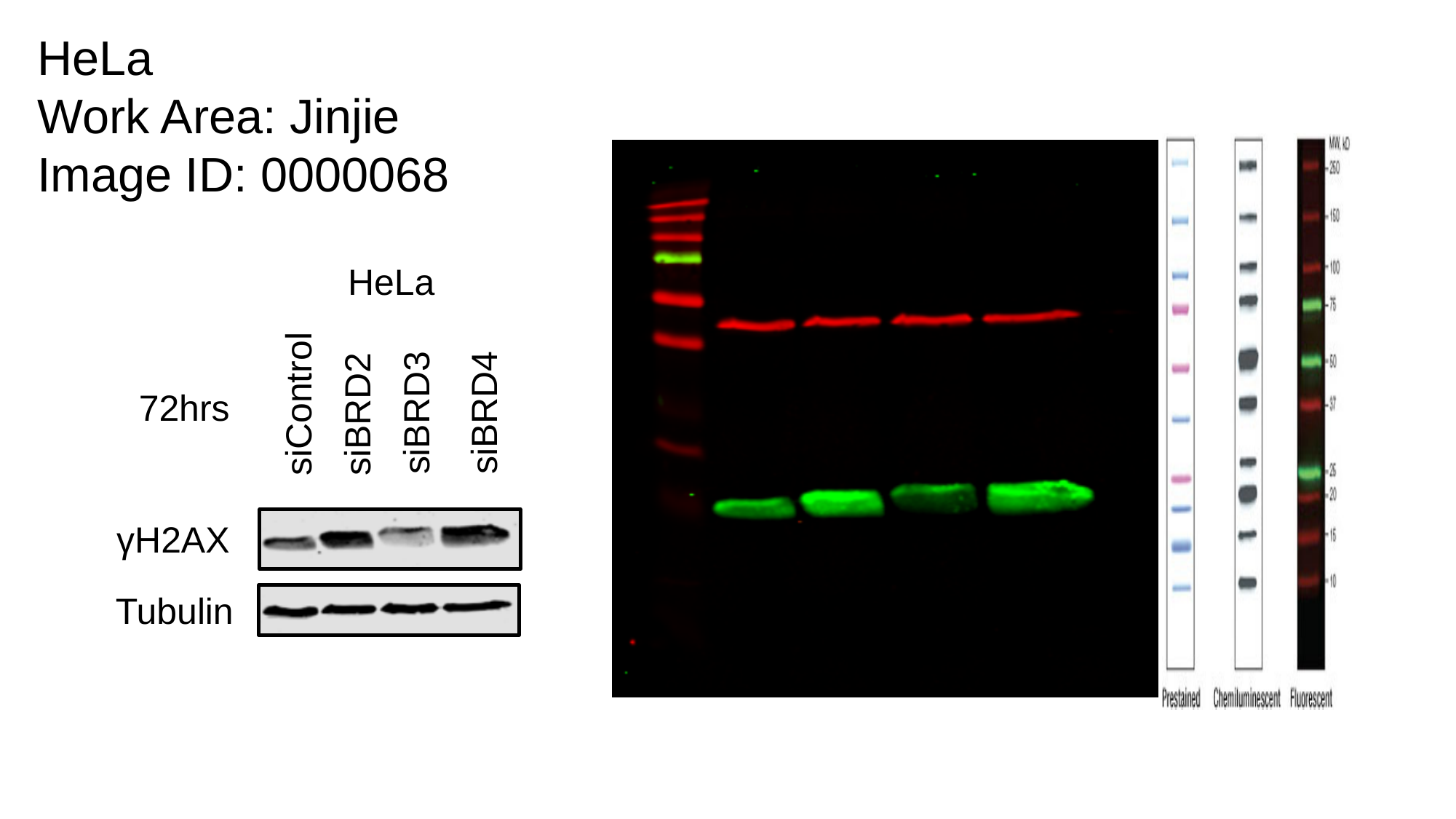

HeLa
Work Area: Jinjie
Image ID: 0000068
HeLa
siControl
72hrs
siBRD3
siBRD4
siBRD2
γH2AX
Tubulin
