## Supplementary material for "BRD4 Prevents R-Loop Formation and Transcription-Replication Conflicts by Ensuring Efficient Transcription Elongation": Source Data: 20191028.pptx

### Slide 1
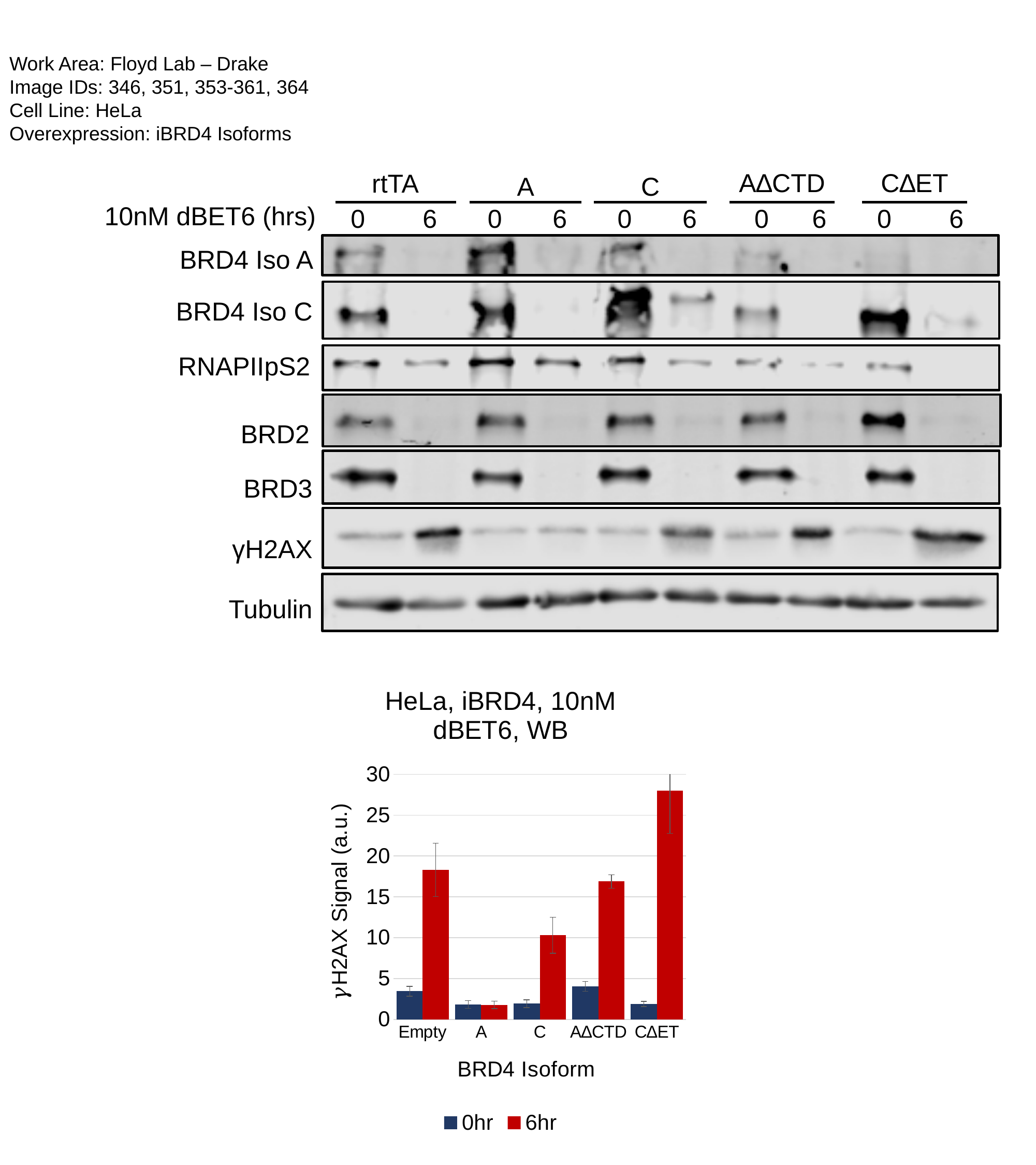

Work Area: Floyd Lab – Drake
Image IDs: 346, 351, 353-361, 364
Cell Line: HeLa
Overexpression: iBRD4 Isoforms
A∆CTD
C∆ET
rtTA
A
C
10nM dBET6 (hrs)
 0 6 0 6 0 6 0 6 0 6
BRD4 Iso A
BRD4 Iso C
RNAPIIpS2
BRD2
BRD3
γH2AX
Tubulin
#### Chart: HeLa, iBRD4, 10nM dBET6, WB
| Category | | |
|---|---|---|
| Empty | 3.441986806363989 | 18.30128205128205 |
| A | 1.816977225672878 | 1.7869955156950672 |
| C | 1.9205745669623997 | 10.298368298368297 |
| A∆CTD | 4.044890162368673 | 16.881542699724516 |
| C∆ET | 1.875 | 28.01474460242233 |

### Slide 2
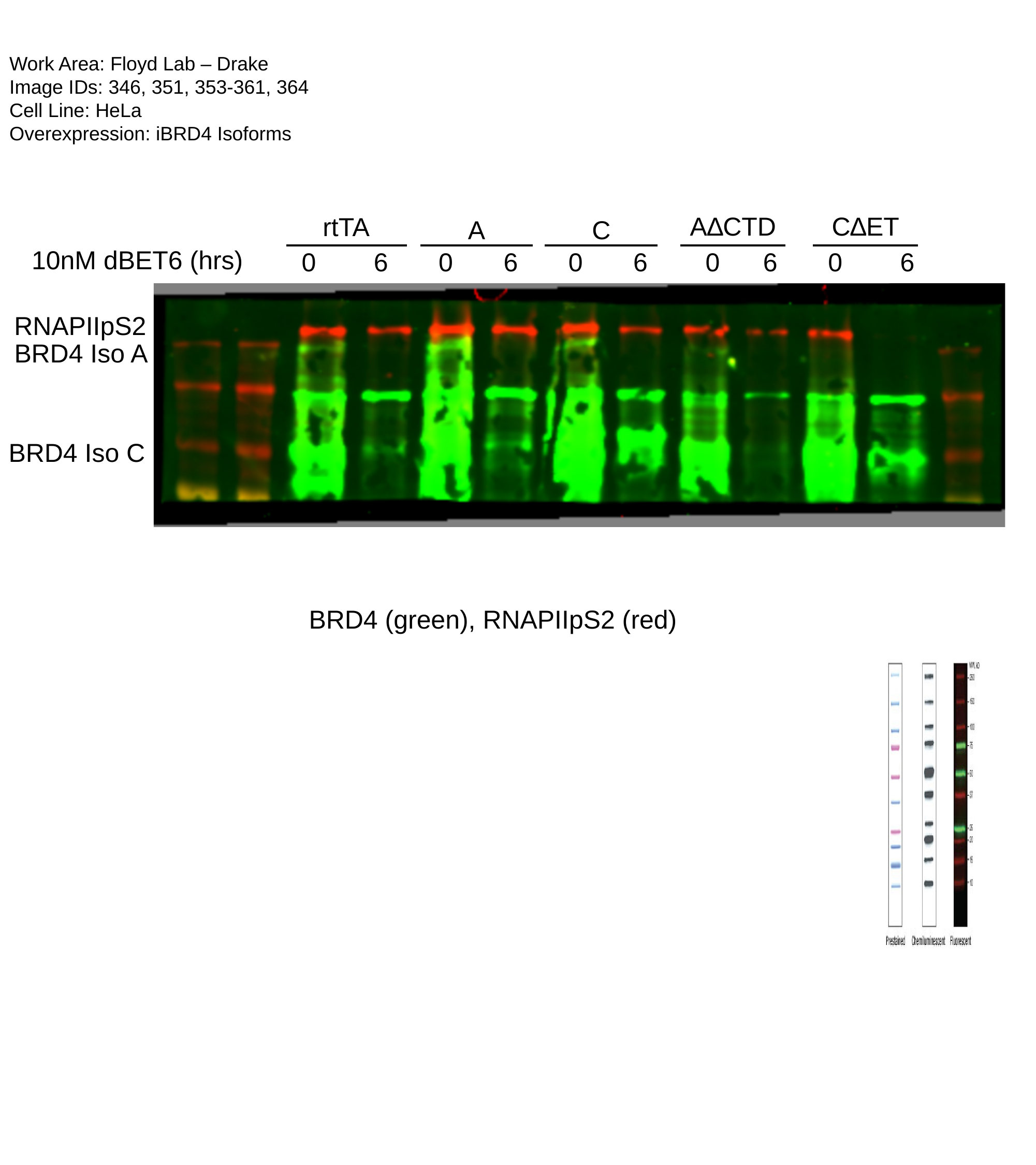

Work Area: Floyd Lab – Drake
Image IDs: 346, 351, 353-361, 364
Cell Line: HeLa
Overexpression: iBRD4 Isoforms
A∆CTD
C∆ET
rtTA
A
C
10nM dBET6 (hrs)
 0 6 0 6 0 6 0 6 0 6
RNAPIIpS2
BRD4 Iso A
BRD4 Iso C
BRD4 (green), RNAPIIpS2 (red)

### Slide 3
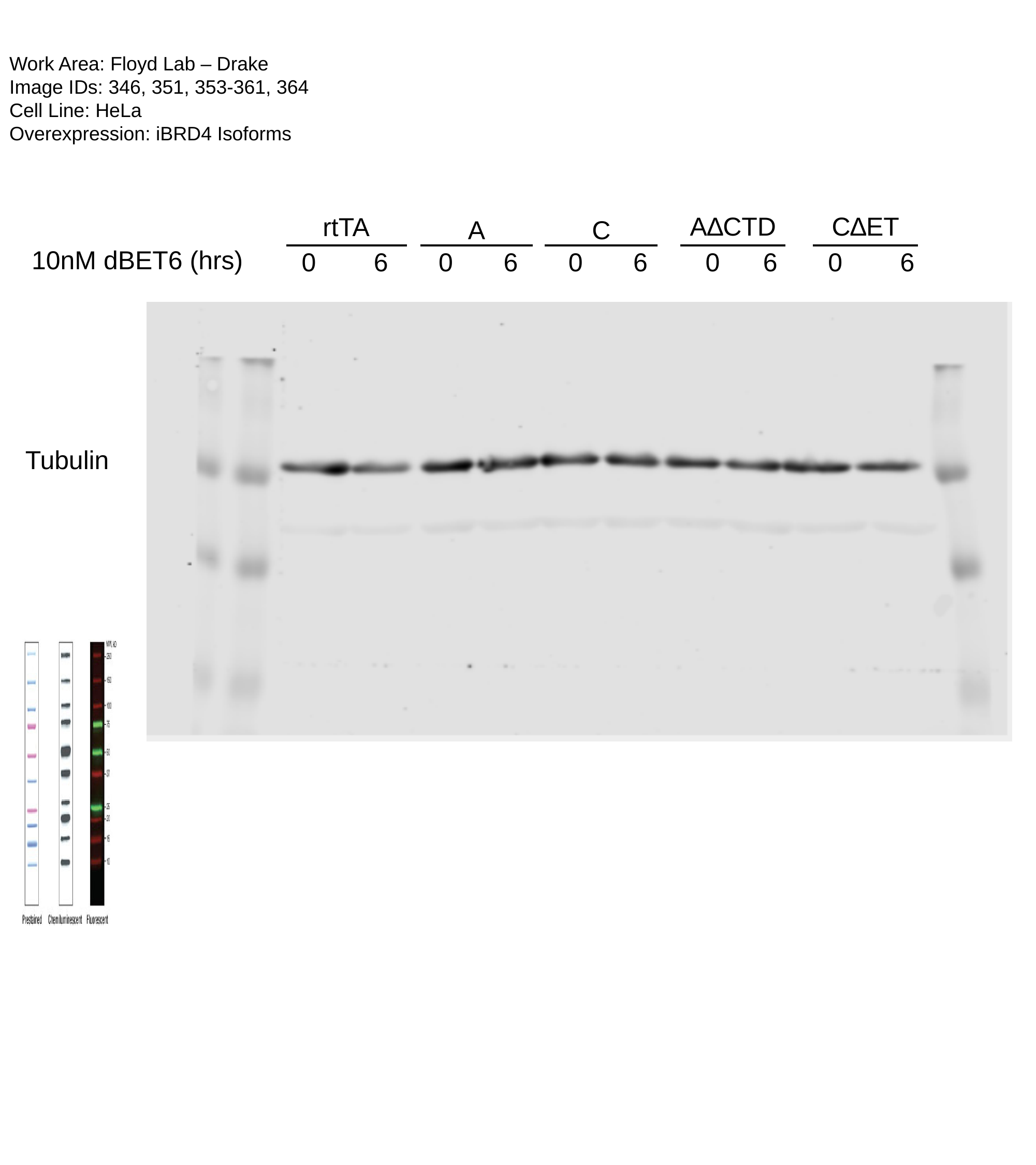

Work Area: Floyd Lab – Drake
Image IDs: 346, 351, 353-361, 364
Cell Line: HeLa
Overexpression: iBRD4 Isoforms
A∆CTD
C∆ET
rtTA
A
C
10nM dBET6 (hrs)
 0 6 0 6 0 6 0 6 0 6
Tubulin

### Slide 4
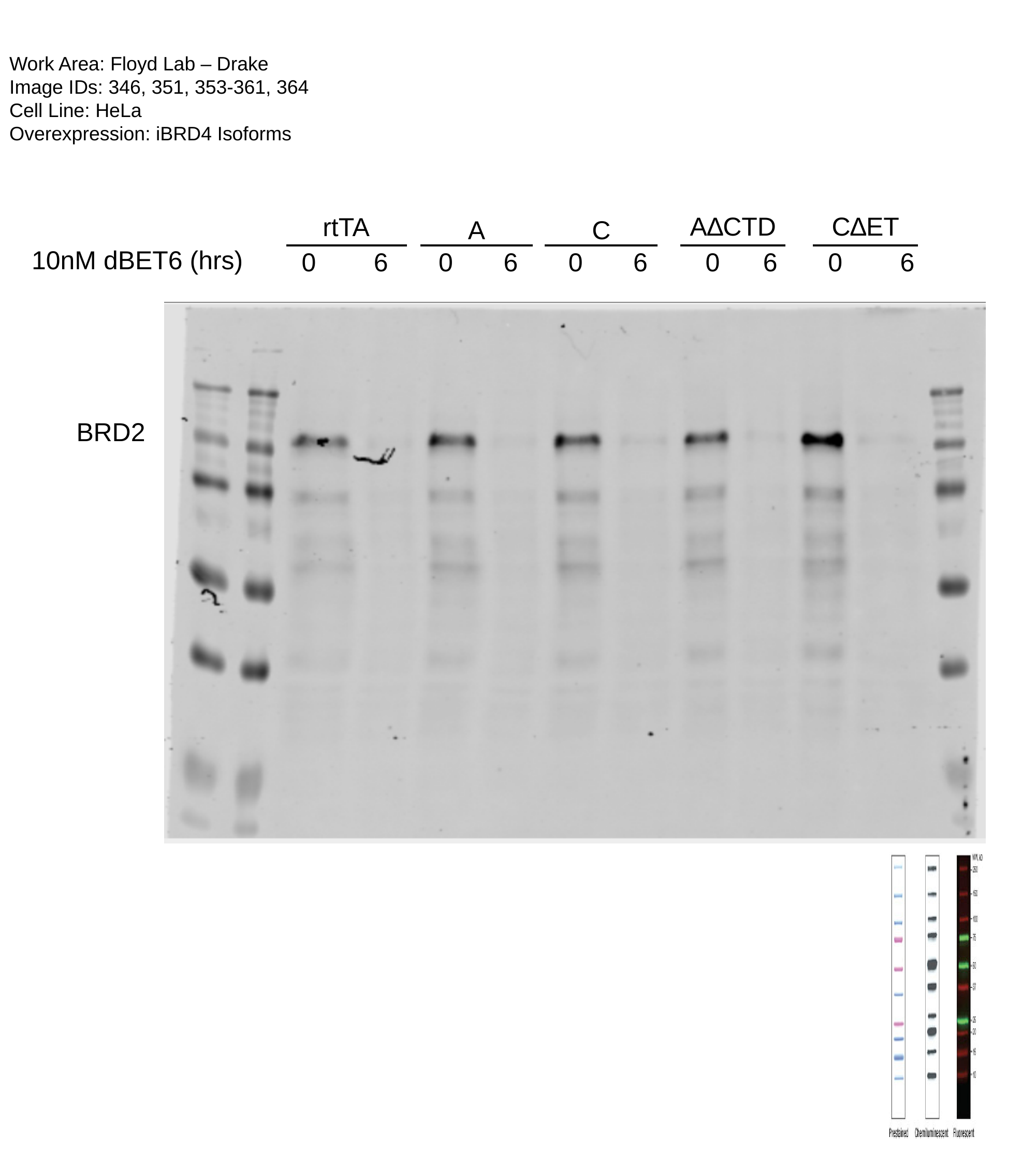

Work Area: Floyd Lab – Drake
Image IDs: 346, 351, 353-361, 364
Cell Line: HeLa
Overexpression: iBRD4 Isoforms
A∆CTD
C∆ET
rtTA
A
C
10nM dBET6 (hrs)
 0 6 0 6 0 6 0 6 0 6
BRD2

### Slide 5
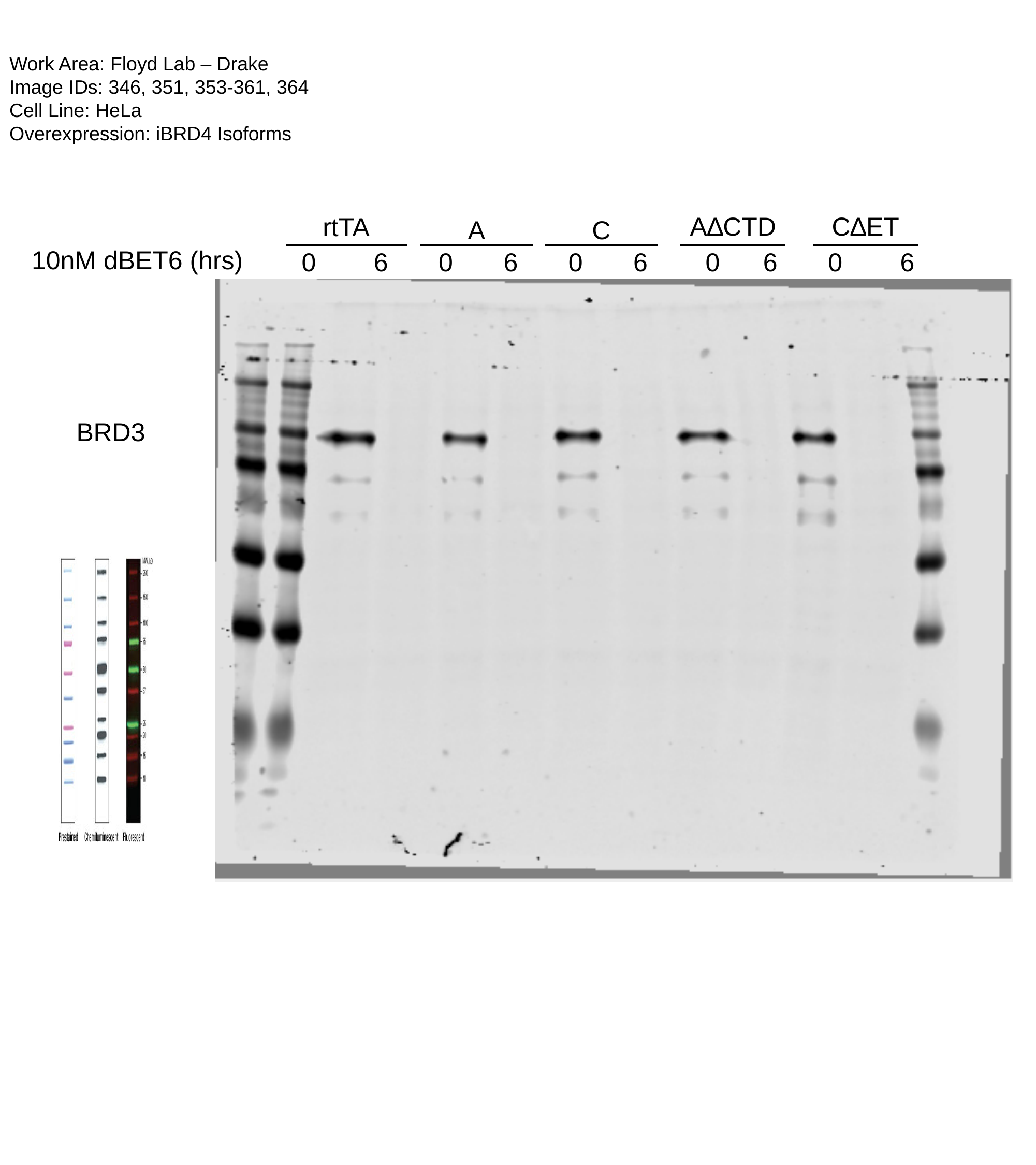

Work Area: Floyd Lab – Drake
Image IDs: 346, 351, 353-361, 364
Cell Line: HeLa
Overexpression: iBRD4 Isoforms
A∆CTD
C∆ET
rtTA
A
C
10nM dBET6 (hrs)
 0 6 0 6 0 6 0 6 0 6
BRD3

### Slide 6
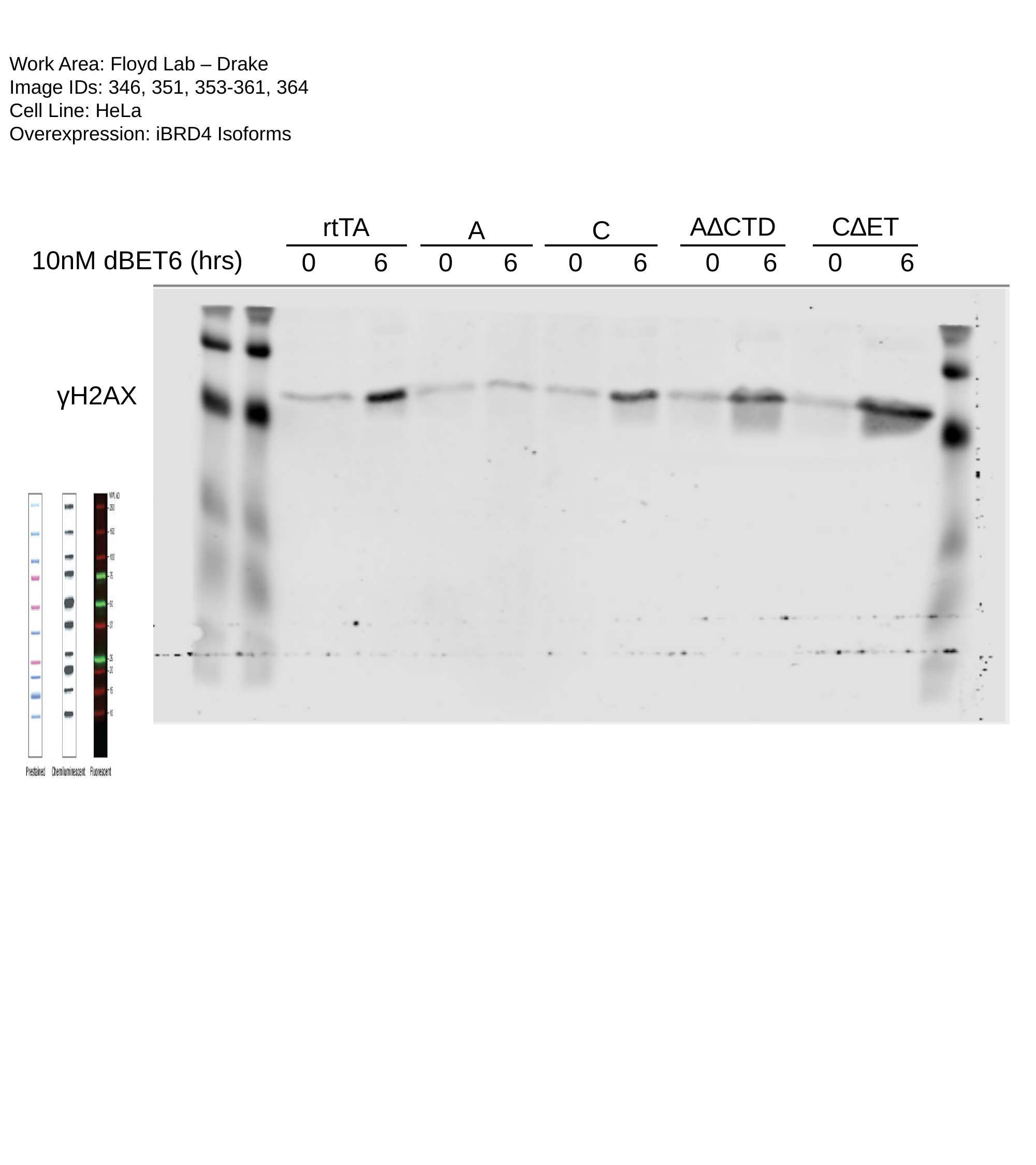

Work Area: Floyd Lab – Drake
Image IDs: 346, 351, 353-361, 364
Cell Line: HeLa
Overexpression: iBRD4 Isoforms
A∆CTD
C∆ET
rtTA
A
C
10nM dBET6 (hrs)
 0 6 0 6 0 6 0 6 0 6
γH2AX
