## Supplementary material for "BRD4 Prevents R-Loop Formation and Transcription-Replication Conflicts by Ensuring Efficient Transcription Elongation": Source Data: 20190604.pptx

### Slide 1
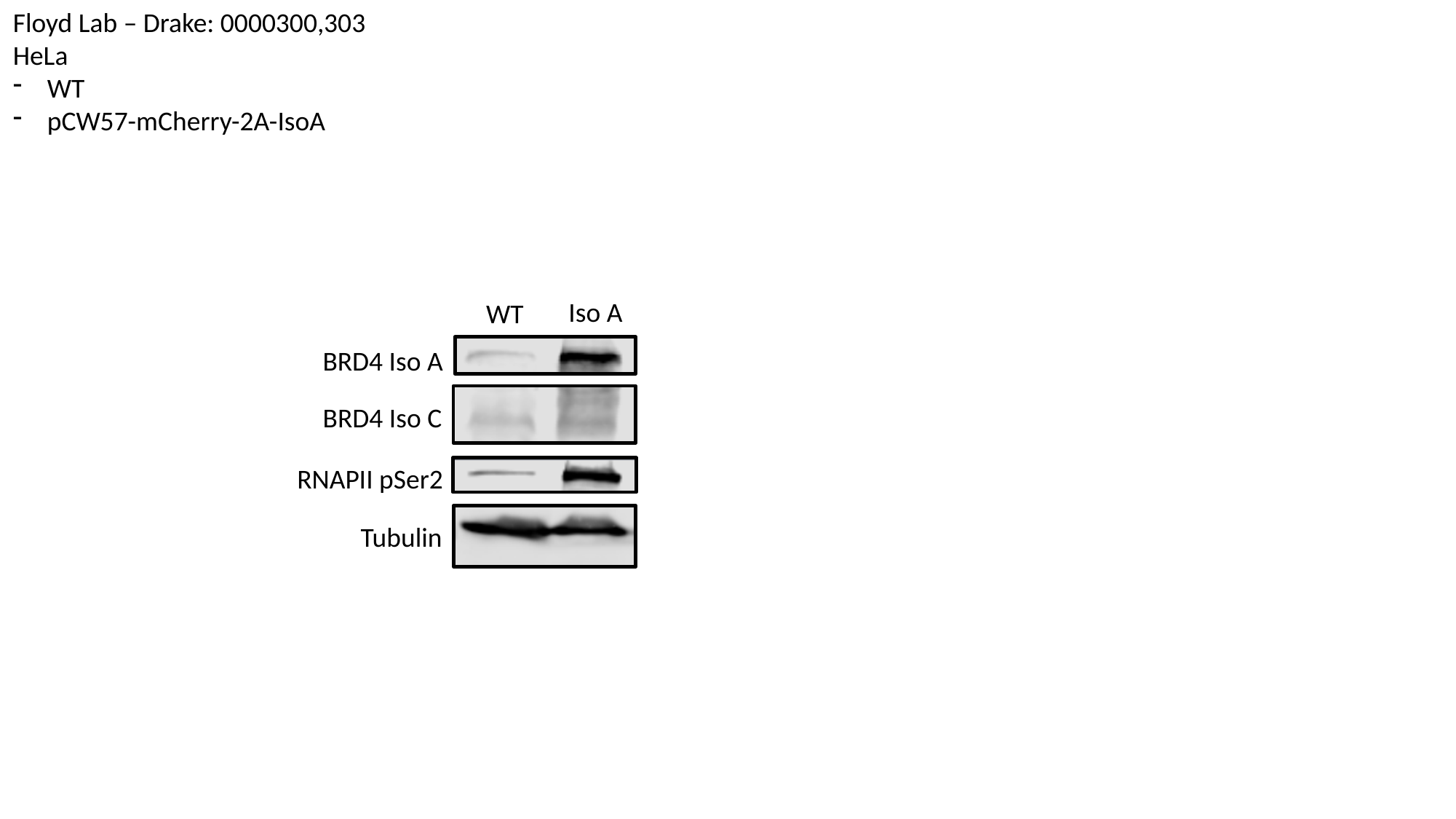

Floyd Lab – Drake: 0000300,303
HeLa
WT
pCW57-mCherry-2A-IsoA
Iso A
WT
BRD4 Iso A
BRD4 Iso C
RNAPII pSer2
Tubulin

### Slide 2
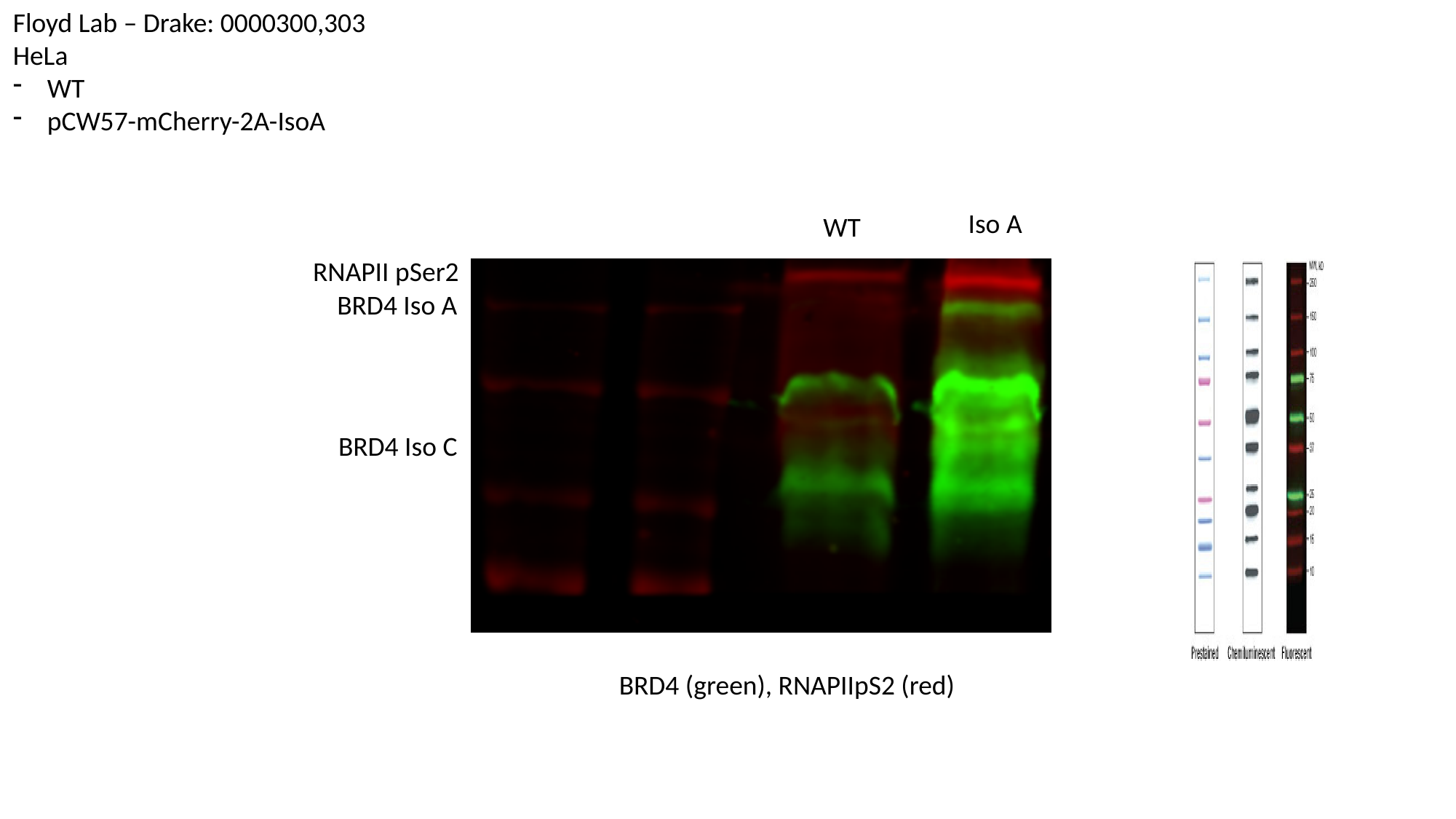

Floyd Lab – Drake: 0000300,303
HeLa
WT
pCW57-mCherry-2A-IsoA
Iso A
WT
RNAPII pSer2
BRD4 Iso A
BRD4 Iso C
BRD4 (green), RNAPIIpS2 (red)

### Slide 3
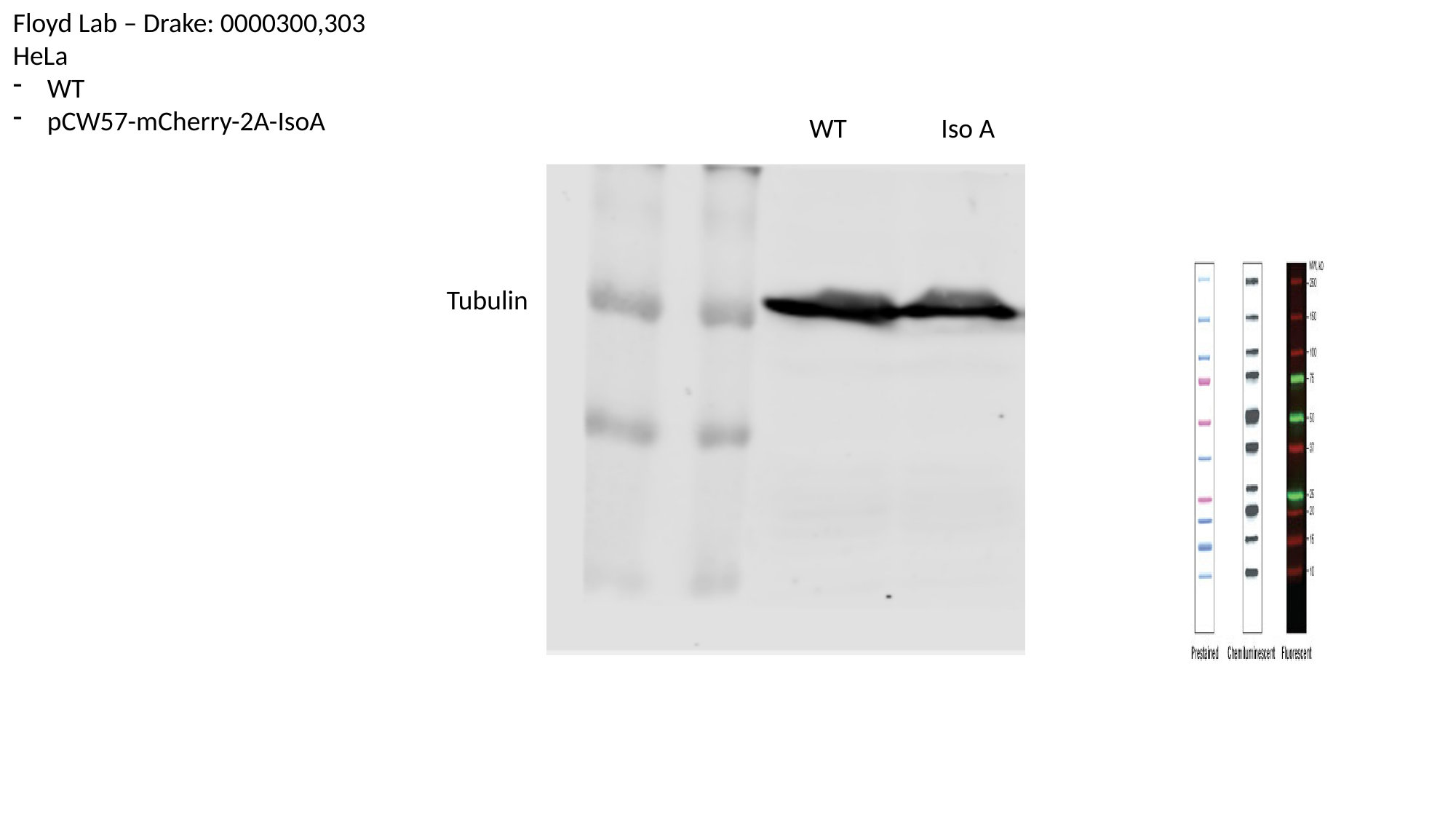

Floyd Lab – Drake: 0000300,303
HeLa
WT
pCW57-mCherry-2A-IsoA
WT
Iso A
Tubulin
