## Supplementary material for "BRD4 Prevents R-Loop Formation and Transcription-Replication Conflicts by Ensuring Efficient Transcription Elongation": Source Data: 293T_HeLa_dBET6.pptx

### Slide 1
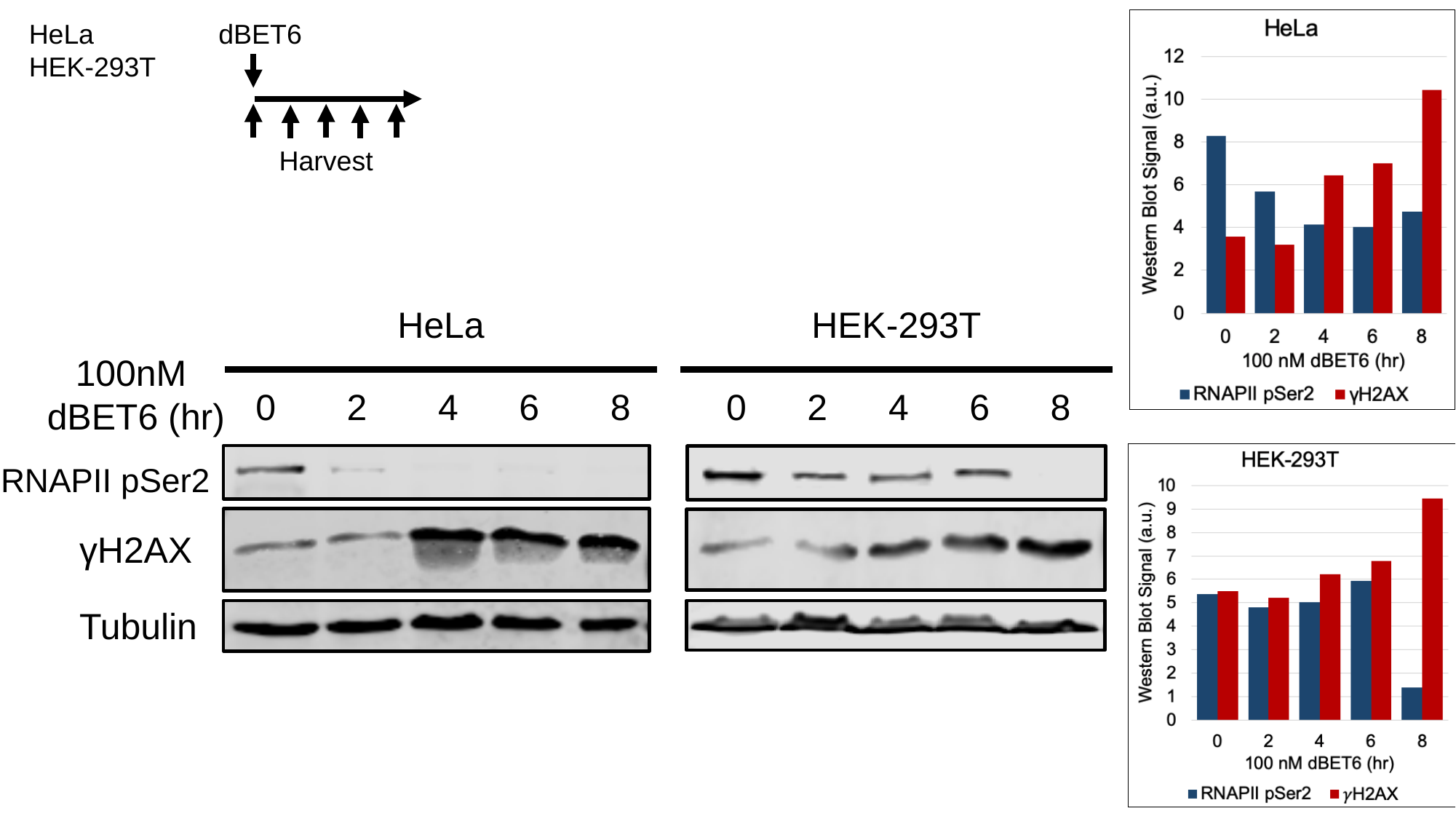

HeLa
HEK-293T
dBET6
Harvest
HeLa
HEK-293T
100nM
dBET6 (hr)
 0 2 4 6 8
 0 2 4 6 8
RNAPII pSer2
γH2AX
Tubulin

### Slide 2
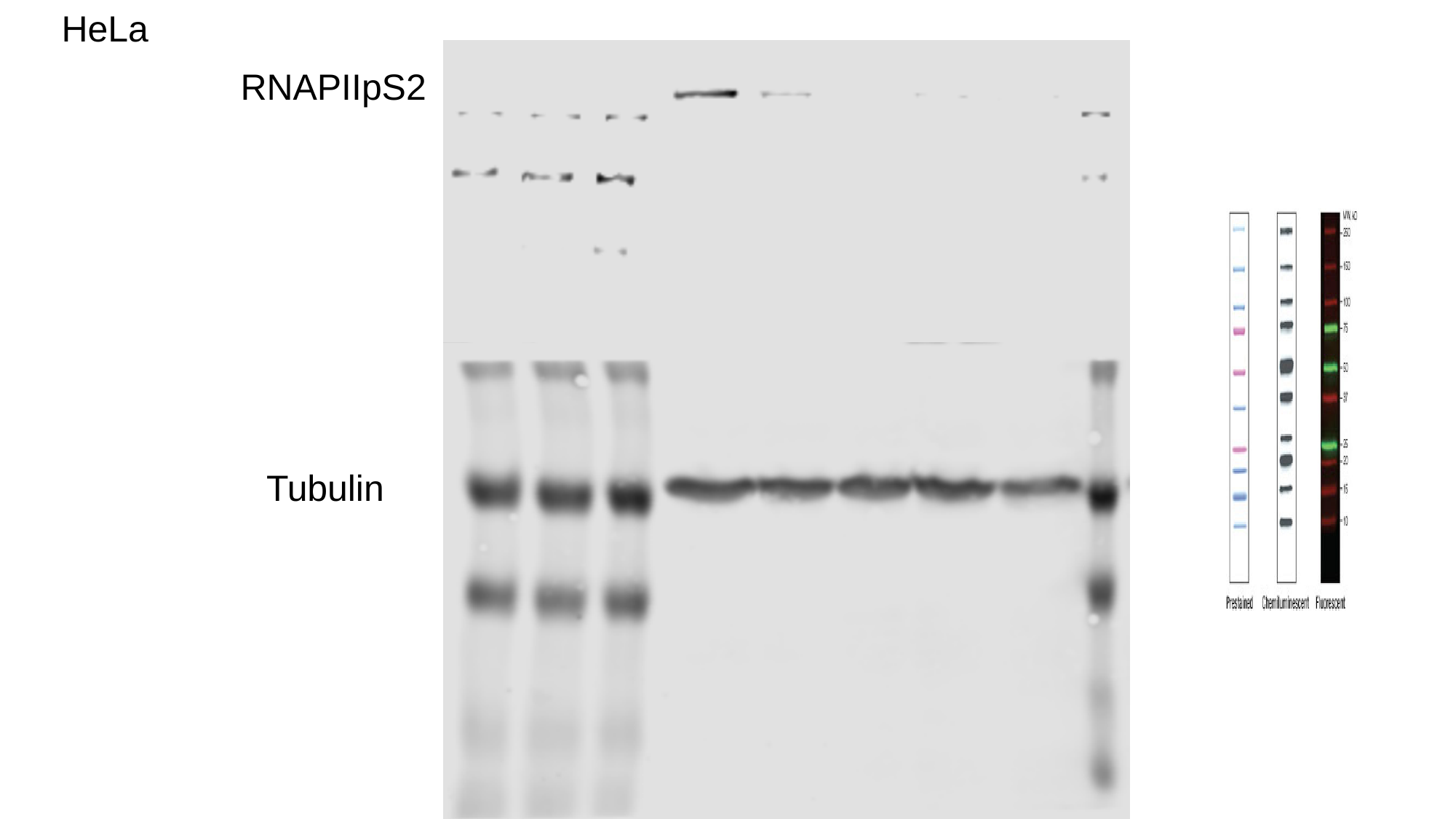

HeLa
RNAPIIpS2
Tubulin

### Slide 3
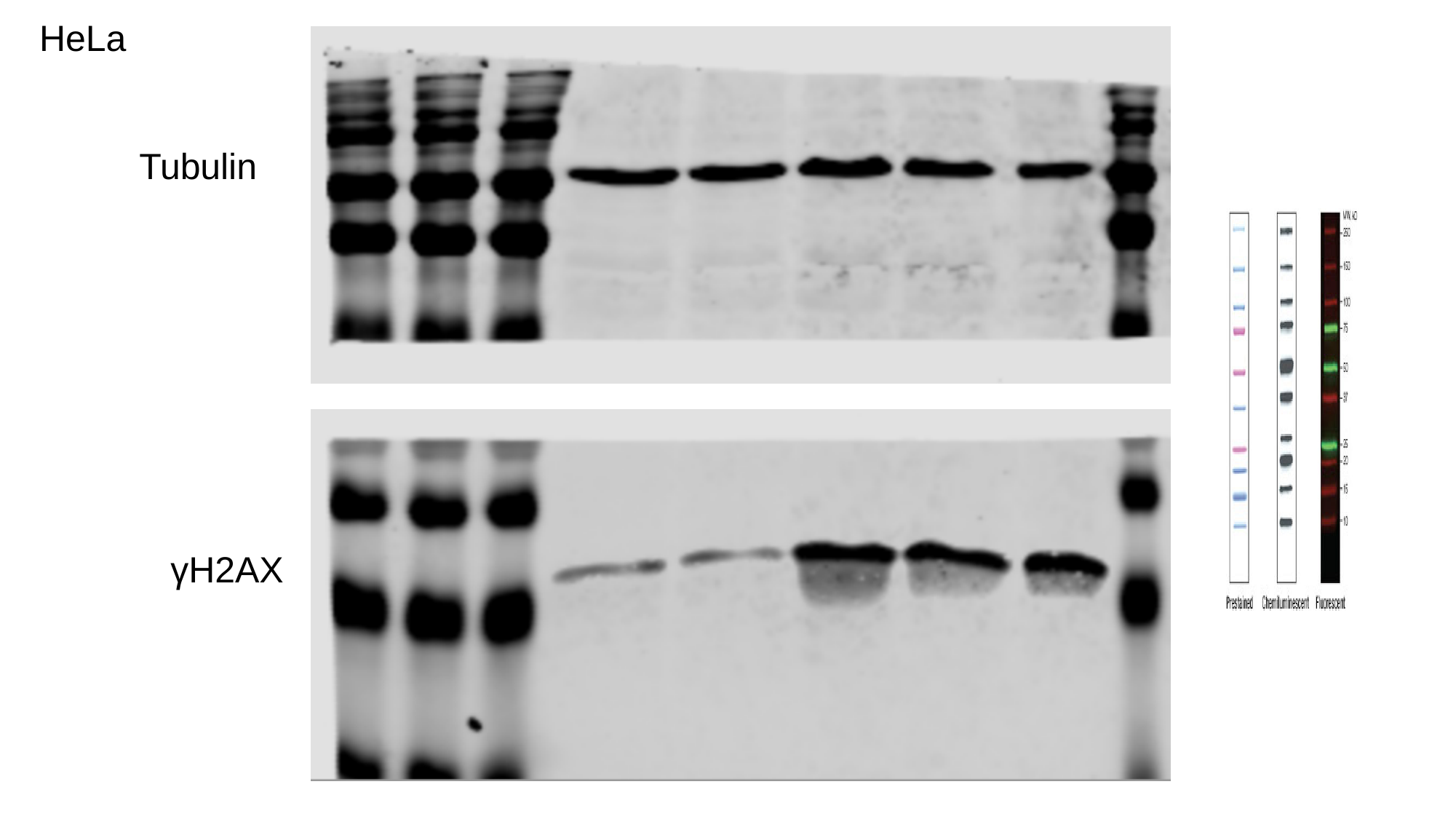

HeLa
Tubulin
γH2AX

### Slide 4
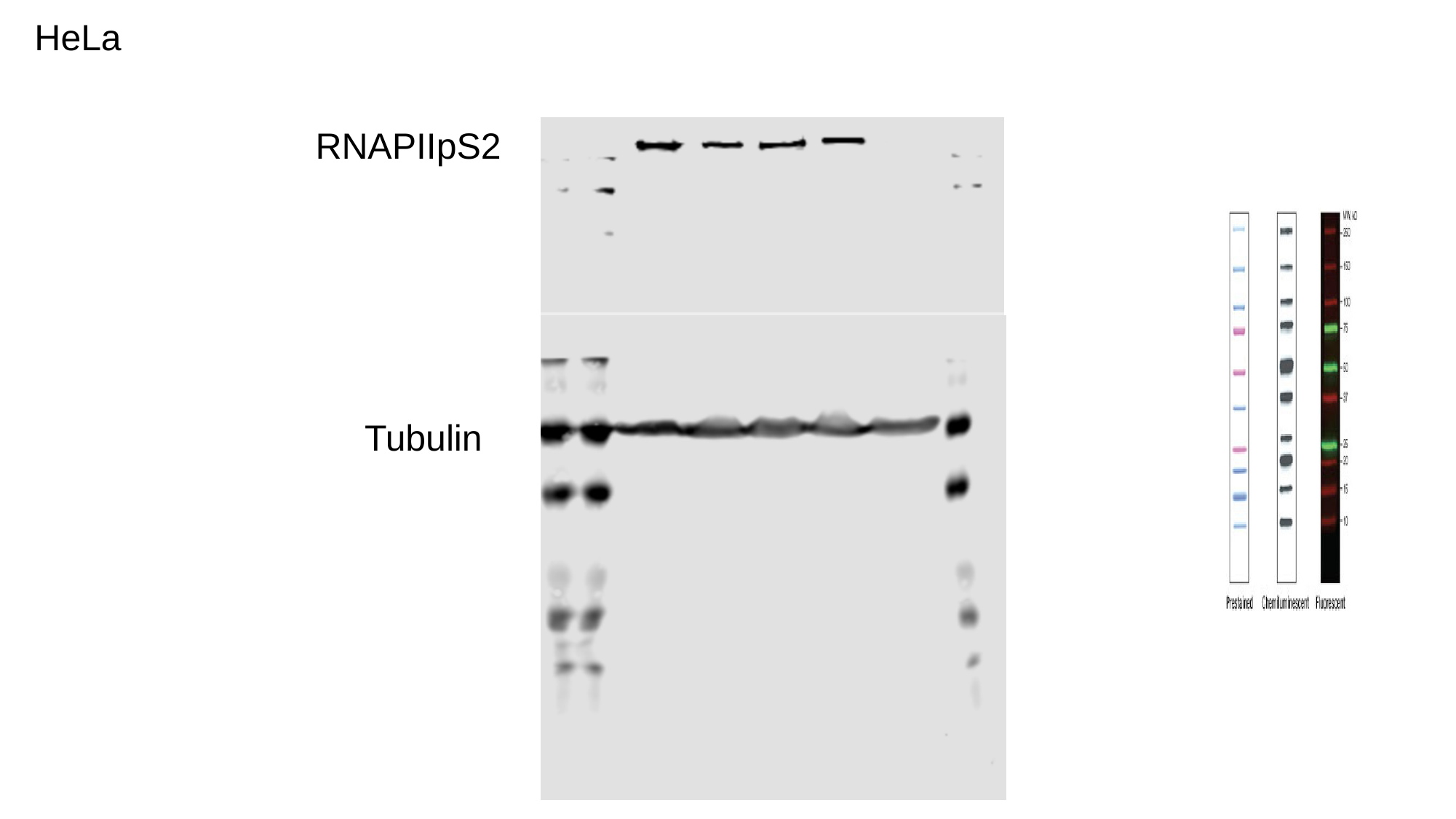

HeLa
RNAPIIpS2
Tubulin

### Slide 5
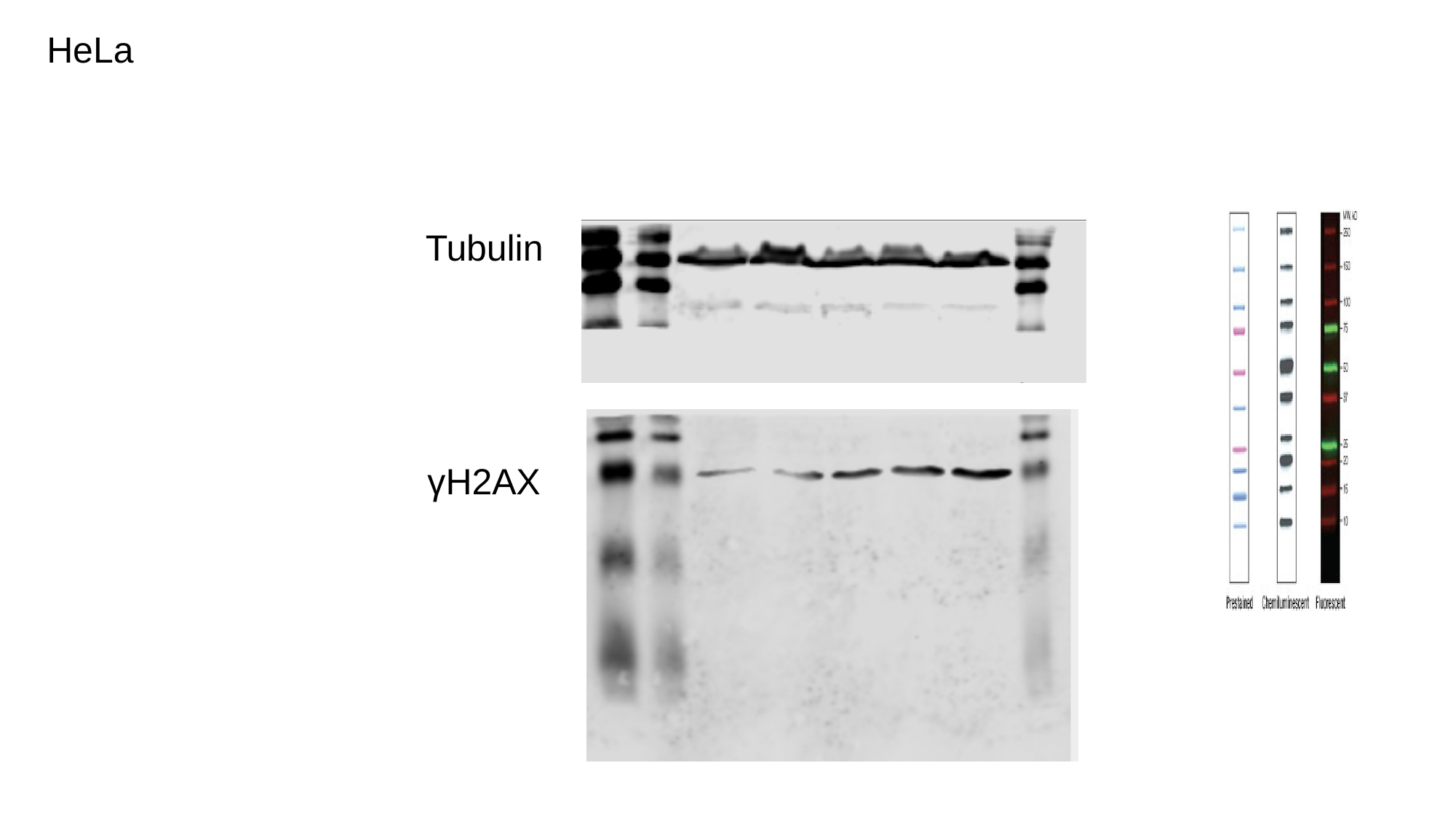

HeLa
Tubulin
γH2AX
