## Supplementary material for "BRD4 Prevents R-Loop Formation and Transcription-Replication Conflicts by Ensuring Efficient Transcription Elongation": Source Data: 20180512_MG132_dBET6.pptx

### Slide 1
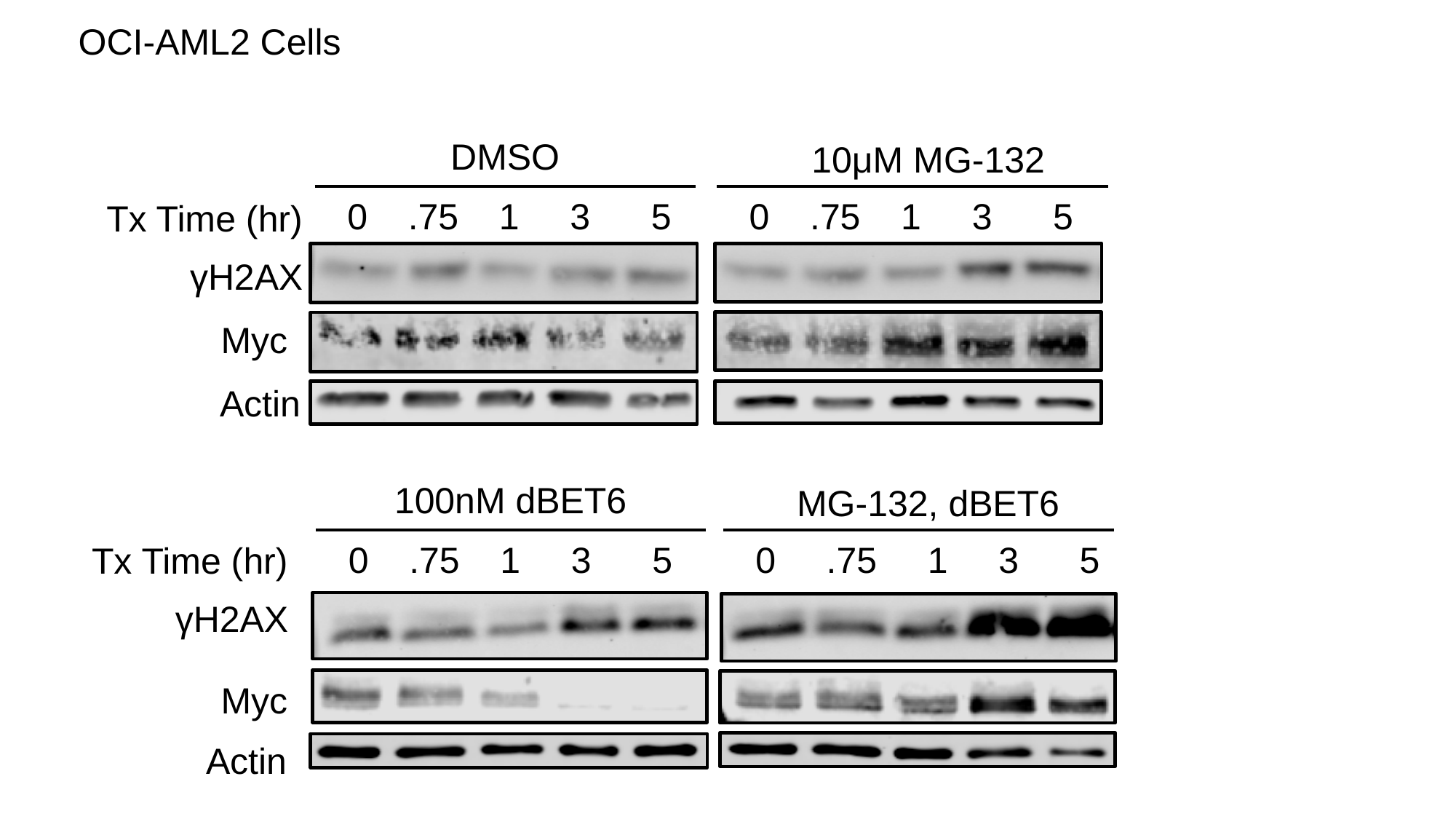

OCI-AML2 Cells
DMSO
10μM MG-132
0 .75 1 3 5
0 .75 1 3 5
Tx Time (hr)
γH2AX
Myc
Actin
100nM dBET6
MG-132, dBET6
0 .75 1 3 5
0 .75 1 3 5
Tx Time (hr)
γH2AX
Myc
Actin

### Slide 2
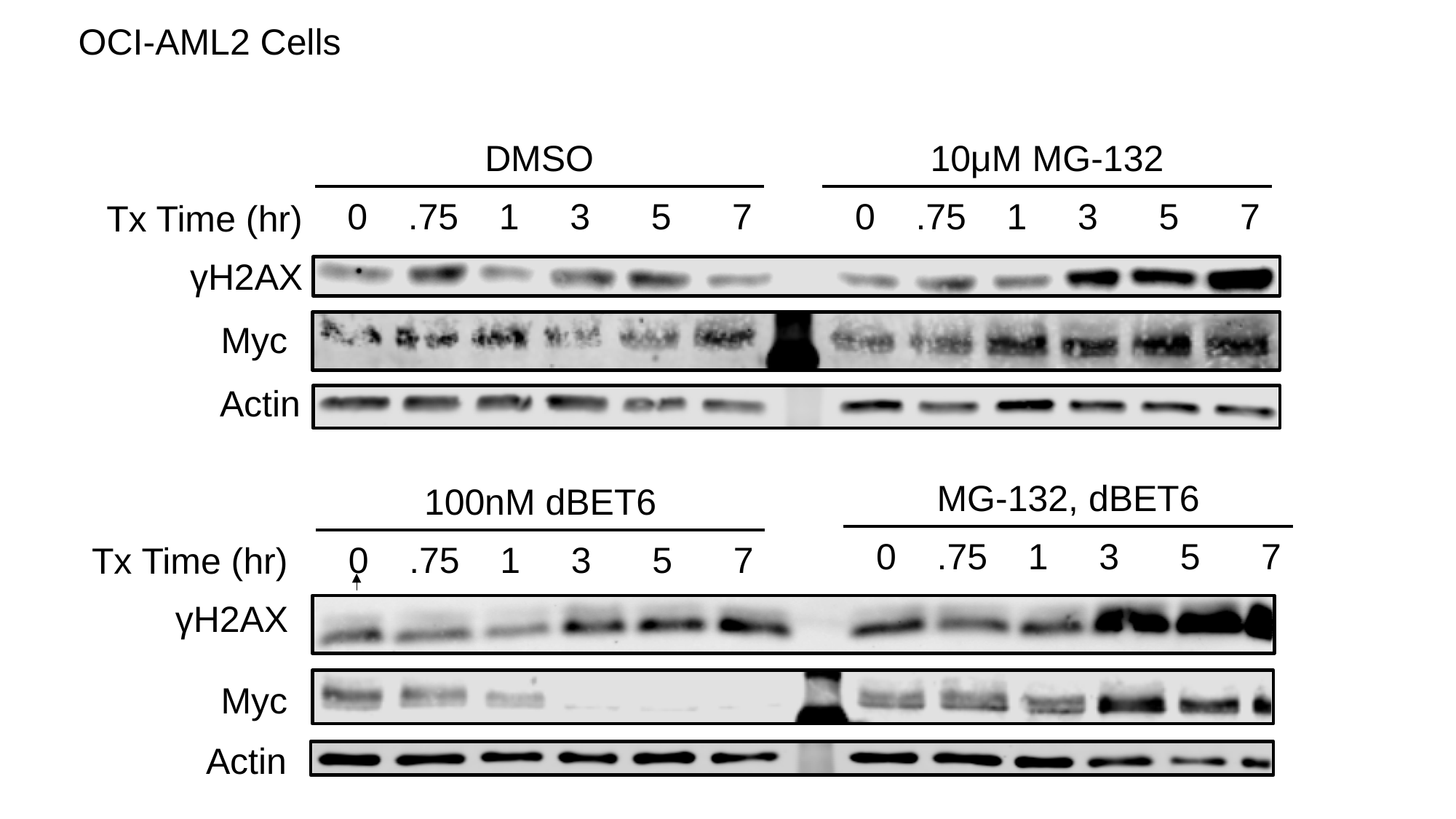

OCI-AML2 Cells
DMSO
10μM MG-132
0 .75 1 3 5 7
0 .75 1 3 5 7
Tx Time (hr)
γH2AX
Myc
Actin
MG-132, dBET6
100nM dBET6
0 .75 1 3 5 7
0 .75 1 3 5 7
Tx Time (hr)
γH2AX
Myc
Actin

### Slide 3
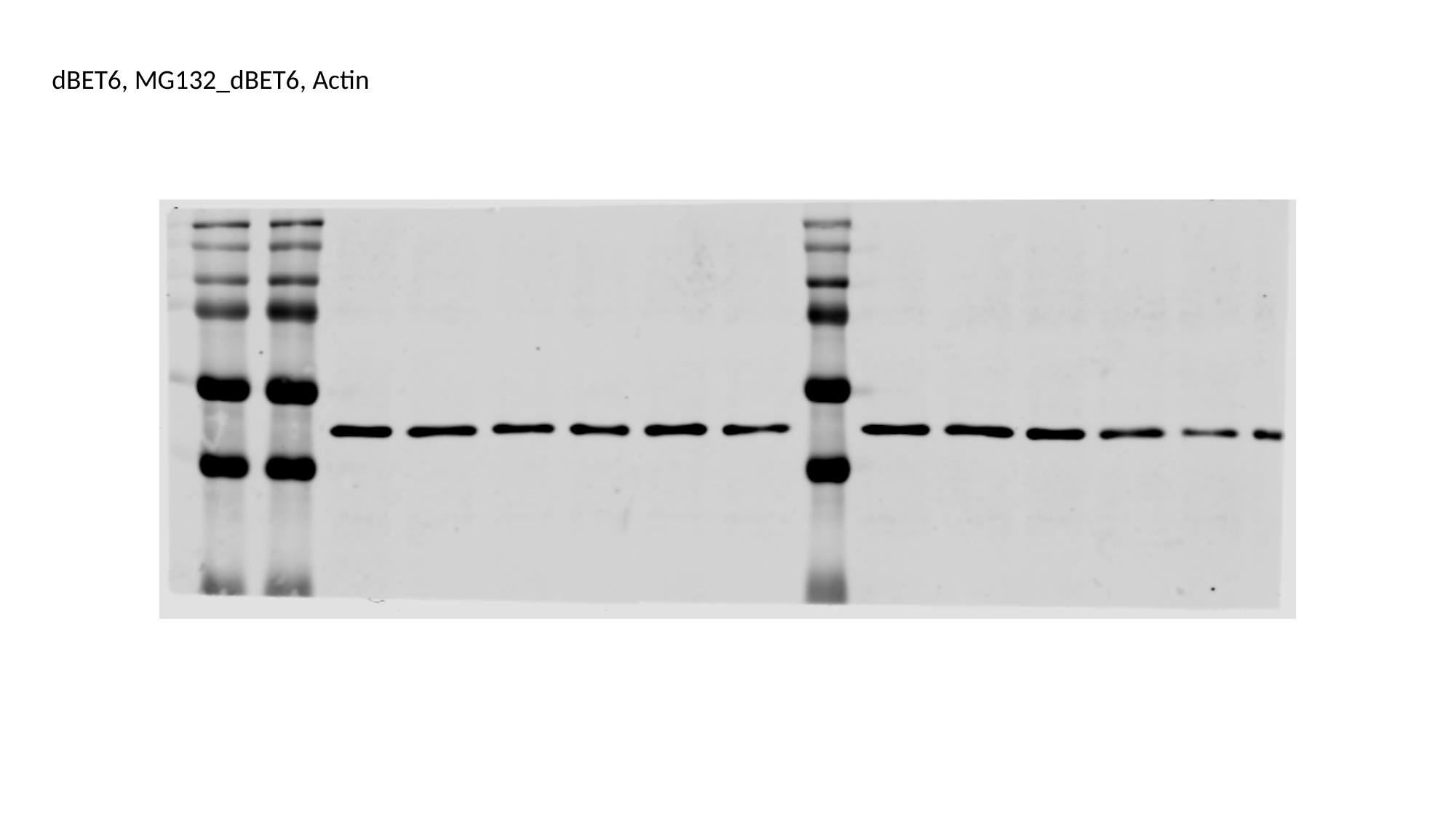

dBET6, MG132_dBET6, Actin

### Slide 4
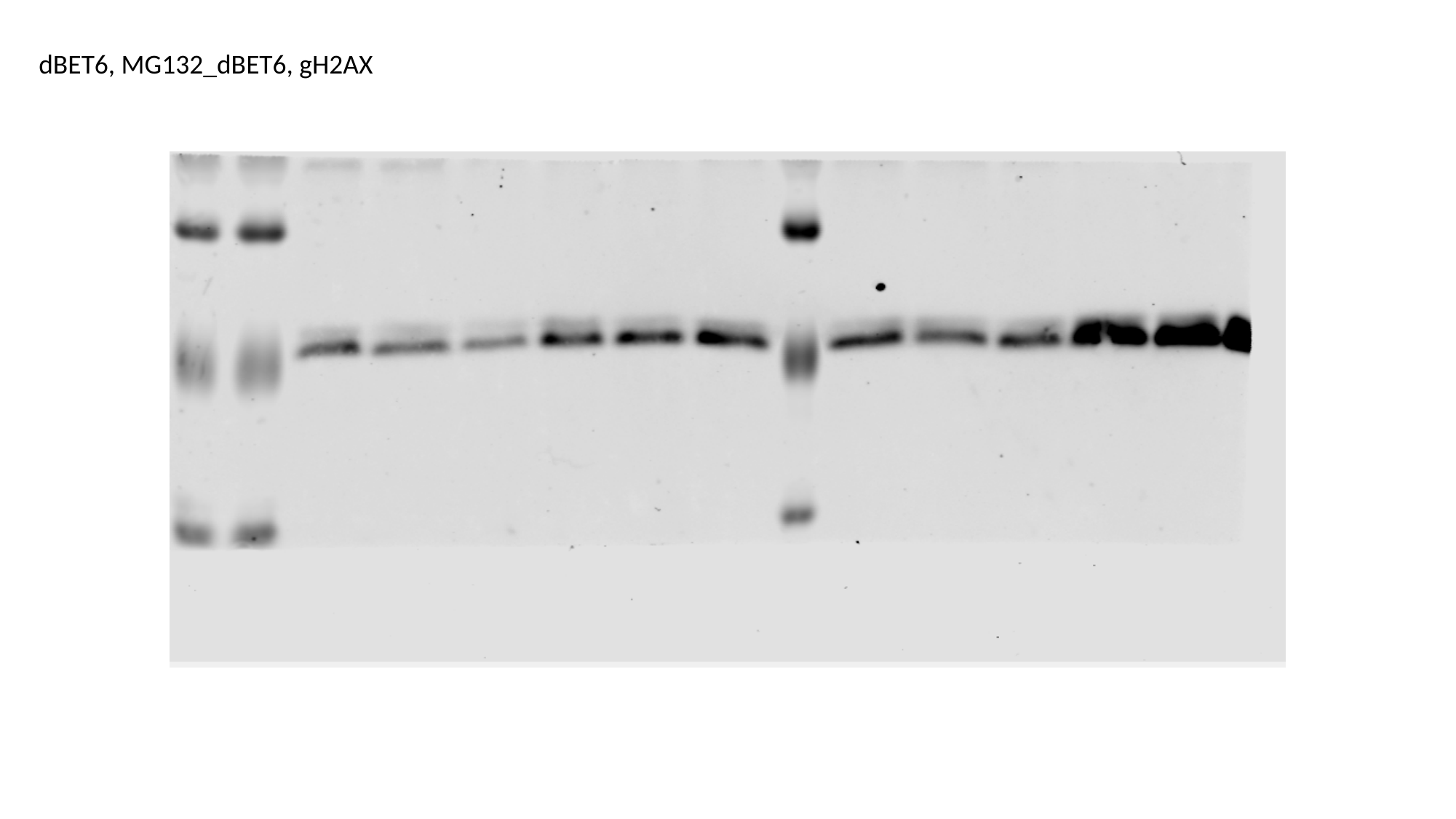

dBET6, MG132_dBET6, gH2AX

### Slide 5
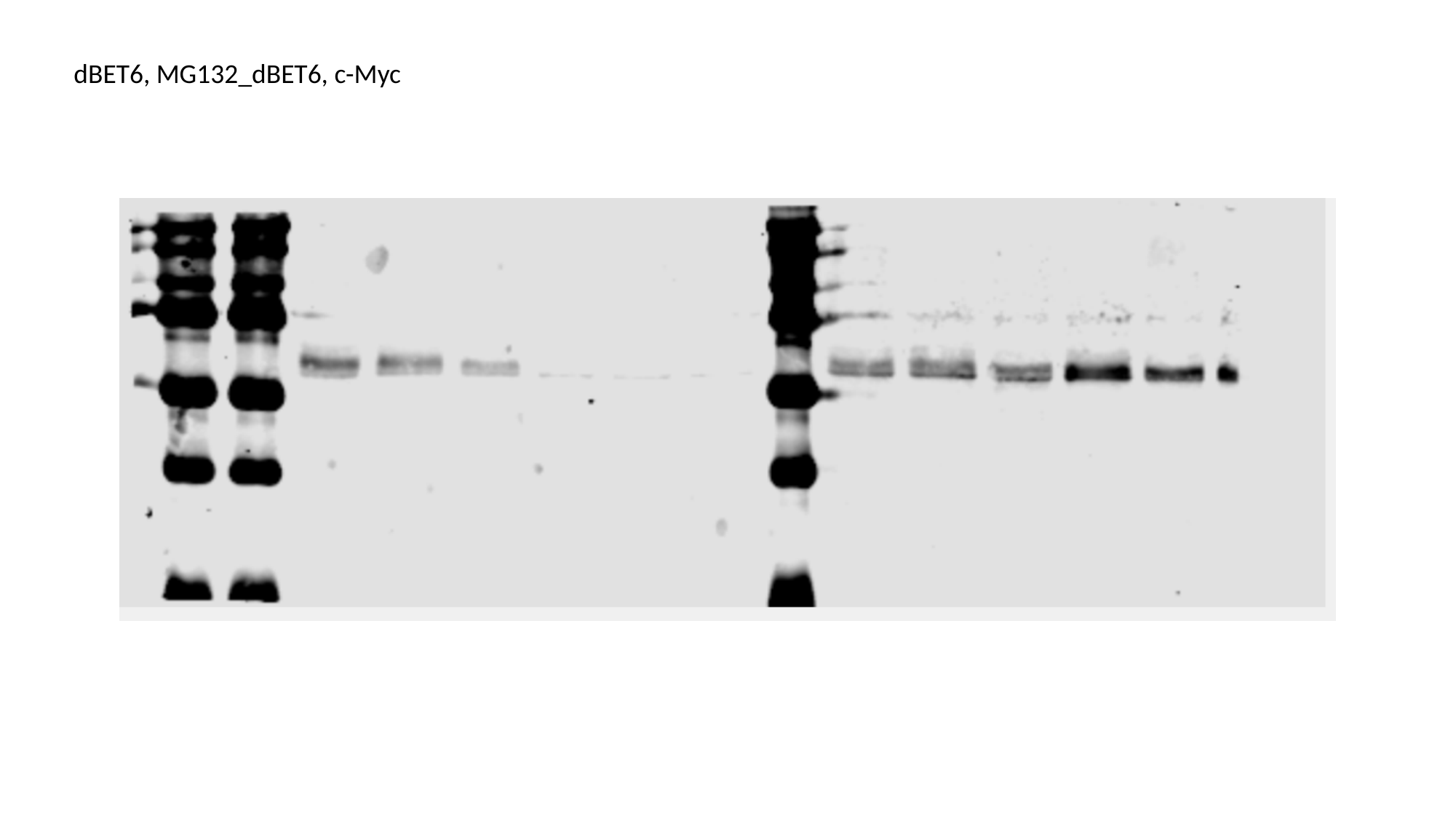

dBET6, MG132_dBET6, c-Myc

### Slide 6
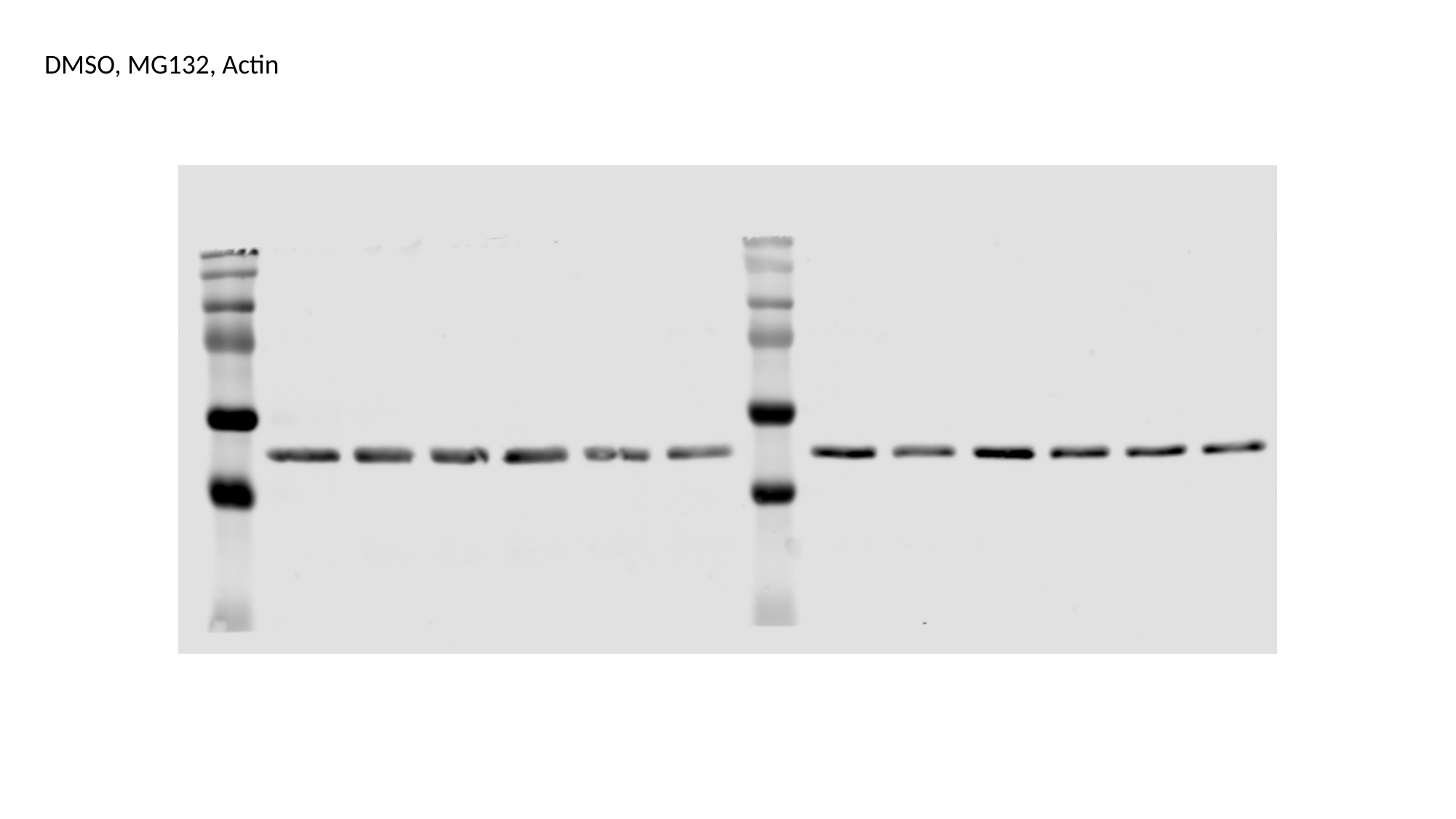

DMSO, MG132, Actin

### Slide 7
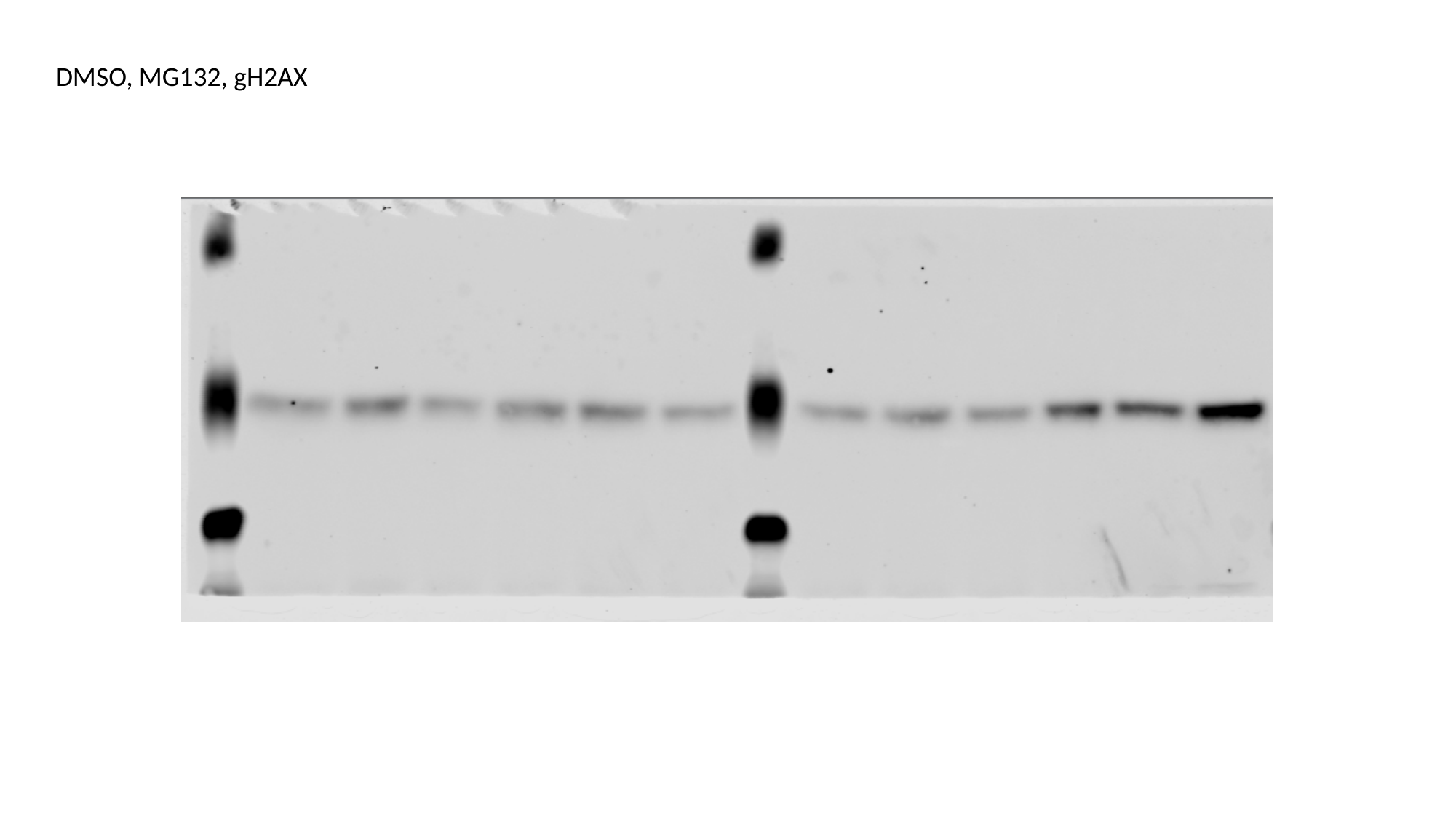

DMSO, MG132, gH2AX

### Slide 8
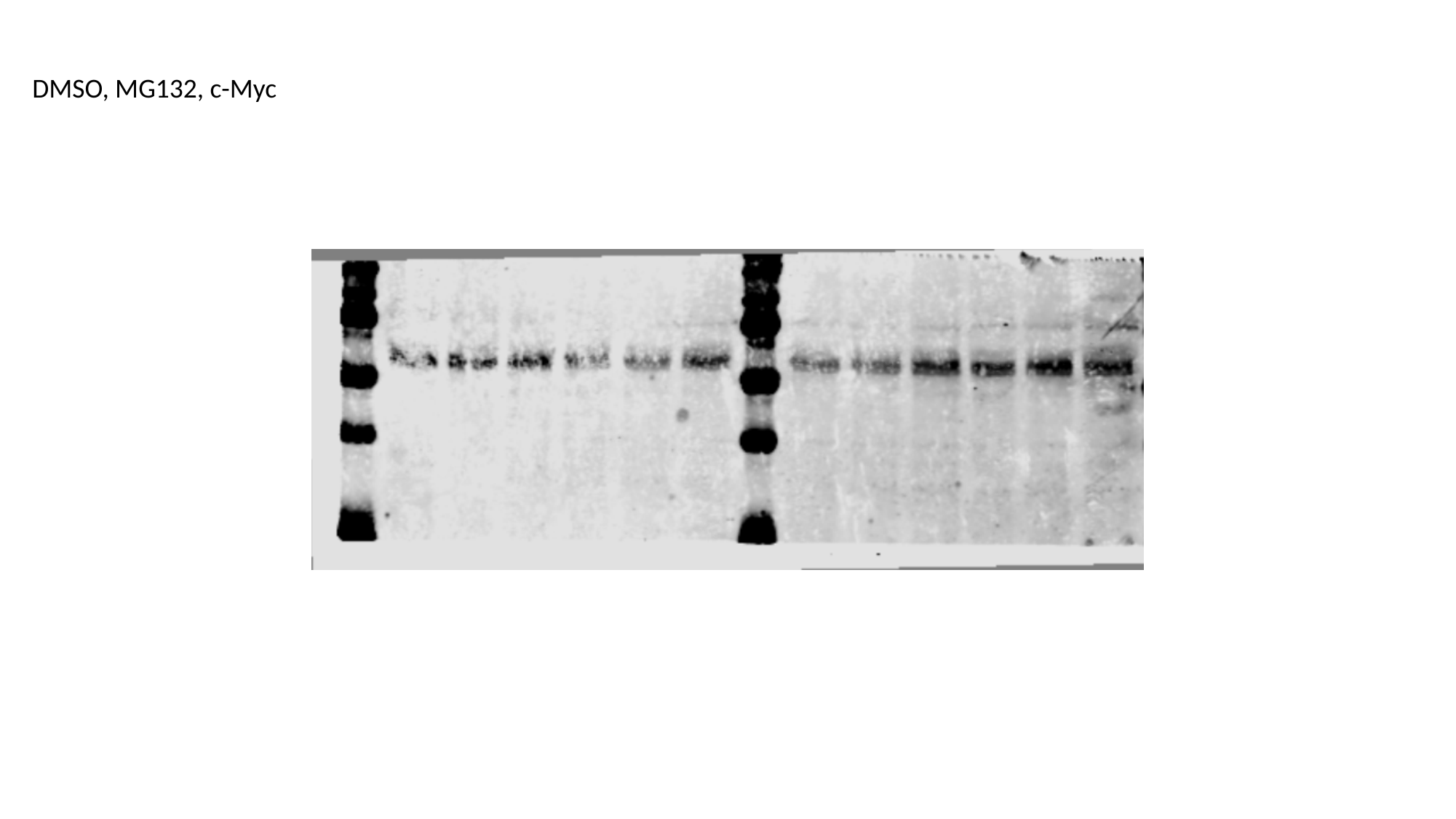

DMSO, MG132, c-Myc
