## Supplementary material for "BRD4 Prevents R-Loop Formation and Transcription-Replication Conflicts by Ensuring Efficient Transcription Elongation": Source Data: 20200305_BETsiRNA.pptx

### Slide 1
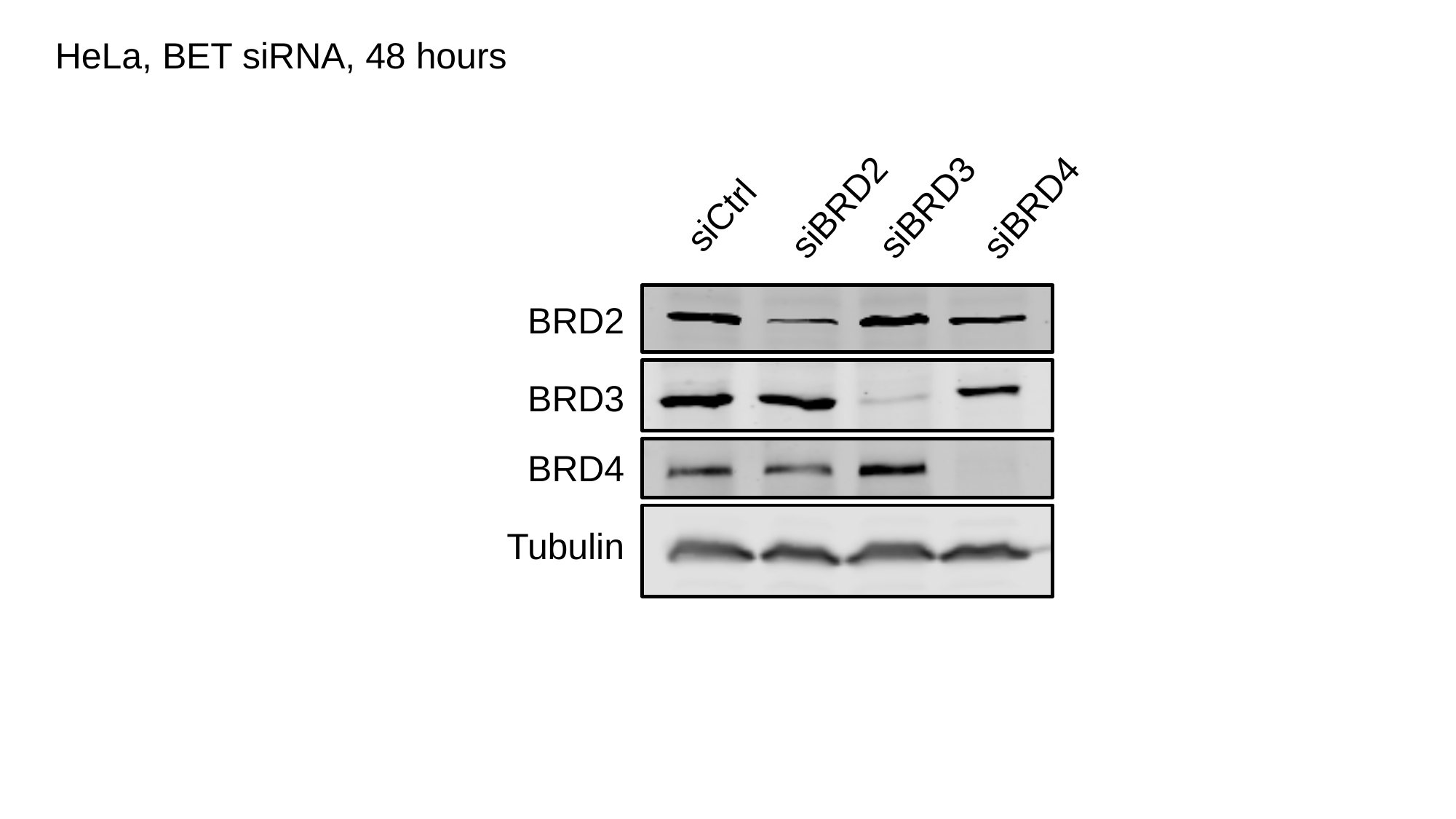

HeLa, BET siRNA, 48 hours
siBRD2
siBRD3
siBRD4
siCtrl
BRD2
BRD3
BRD4
Tubulin

### Slide 2
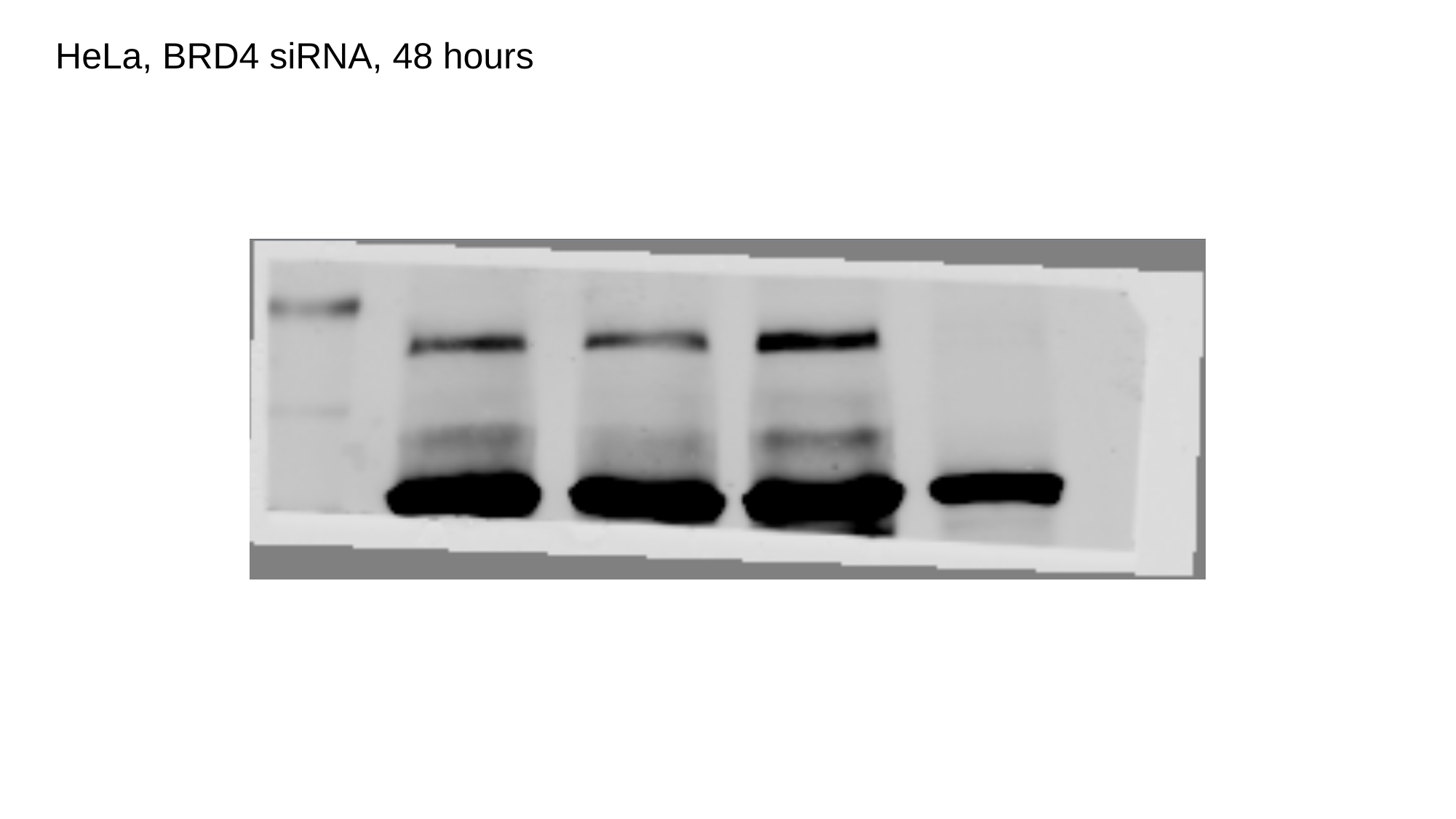

HeLa, BRD4 siRNA, 48 hours

### Slide 3
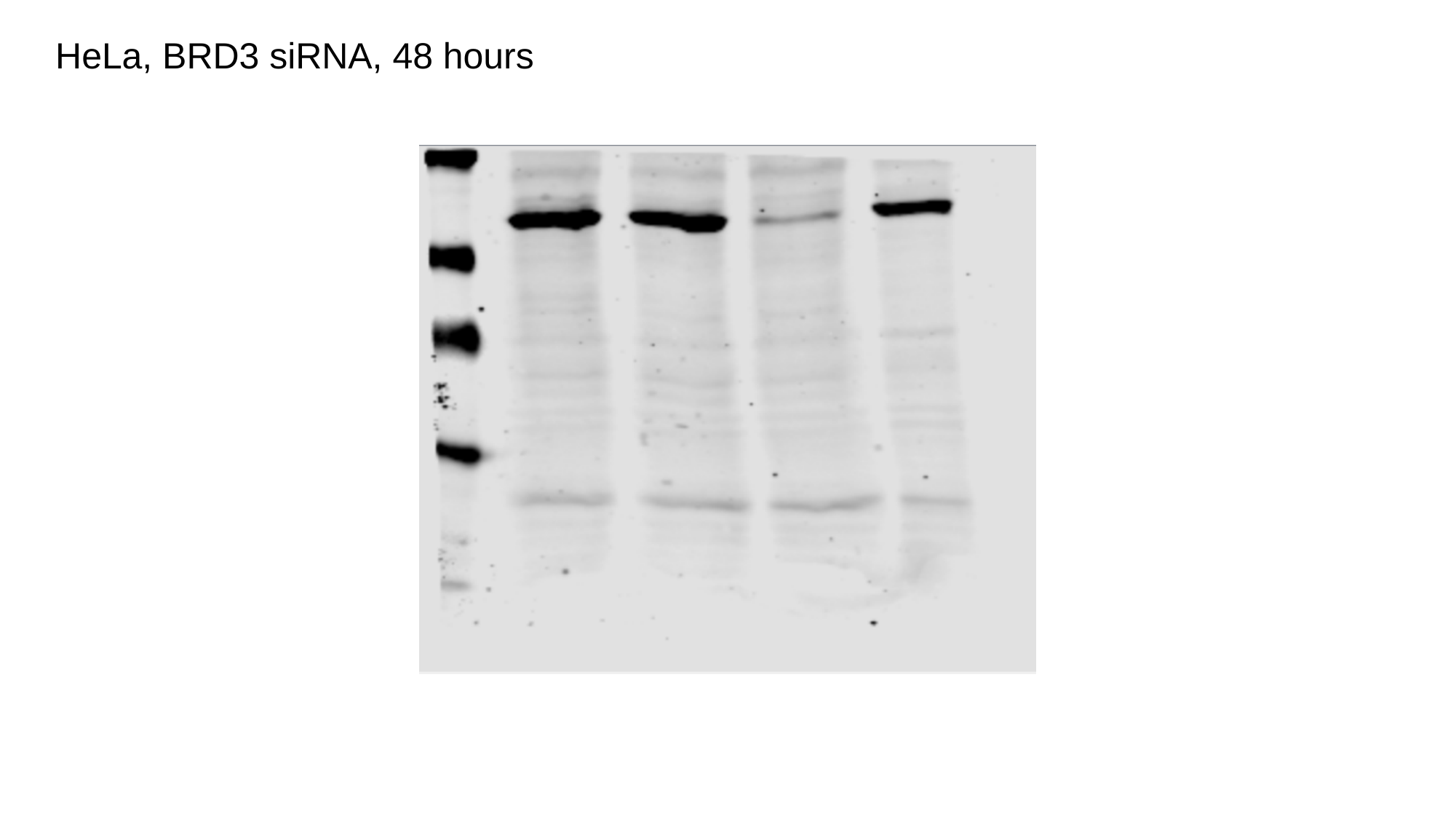

HeLa, BRD3 siRNA, 48 hours

### Slide 4
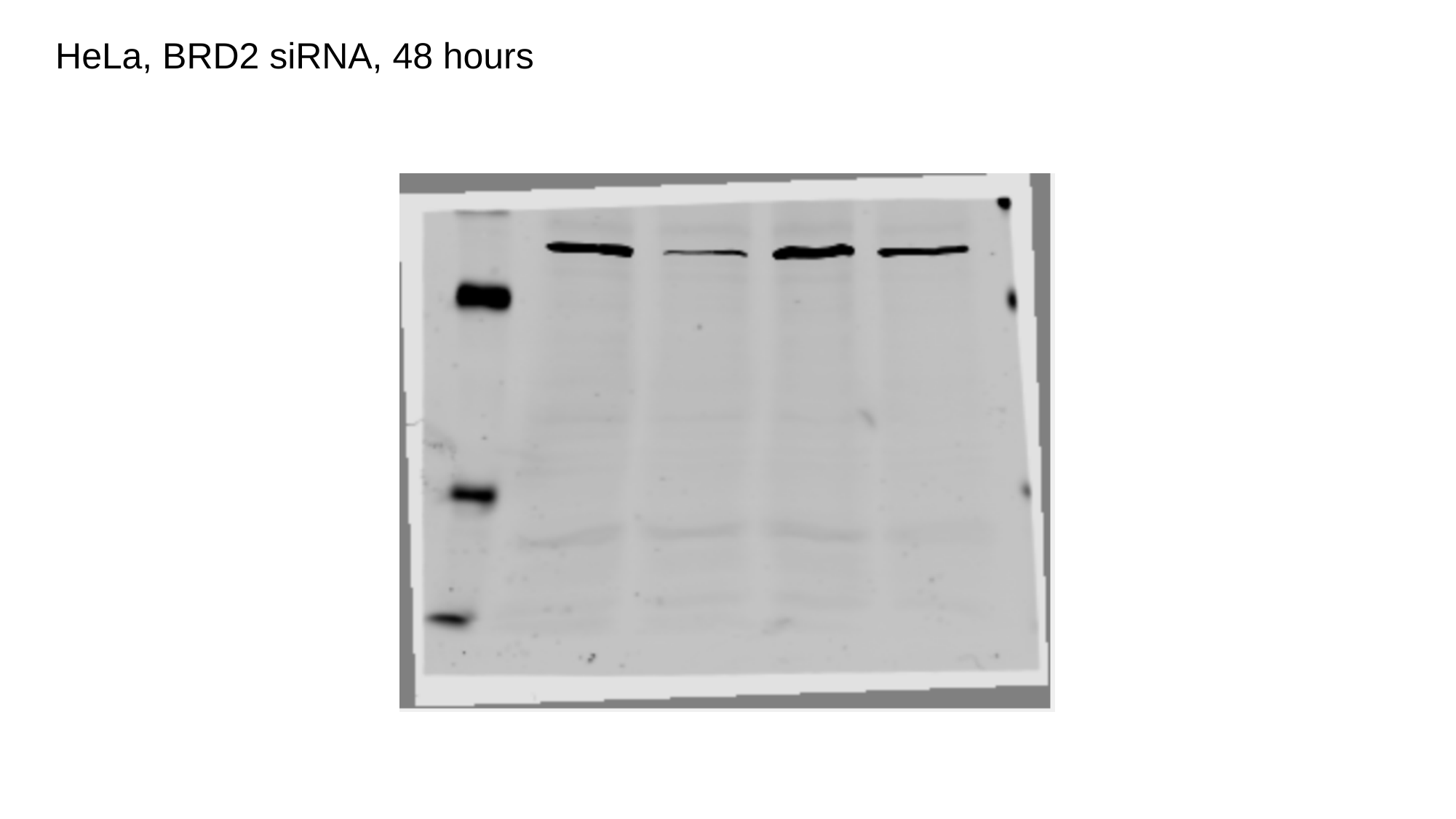

HeLa, BRD2 siRNA, 48 hours

### Slide 5
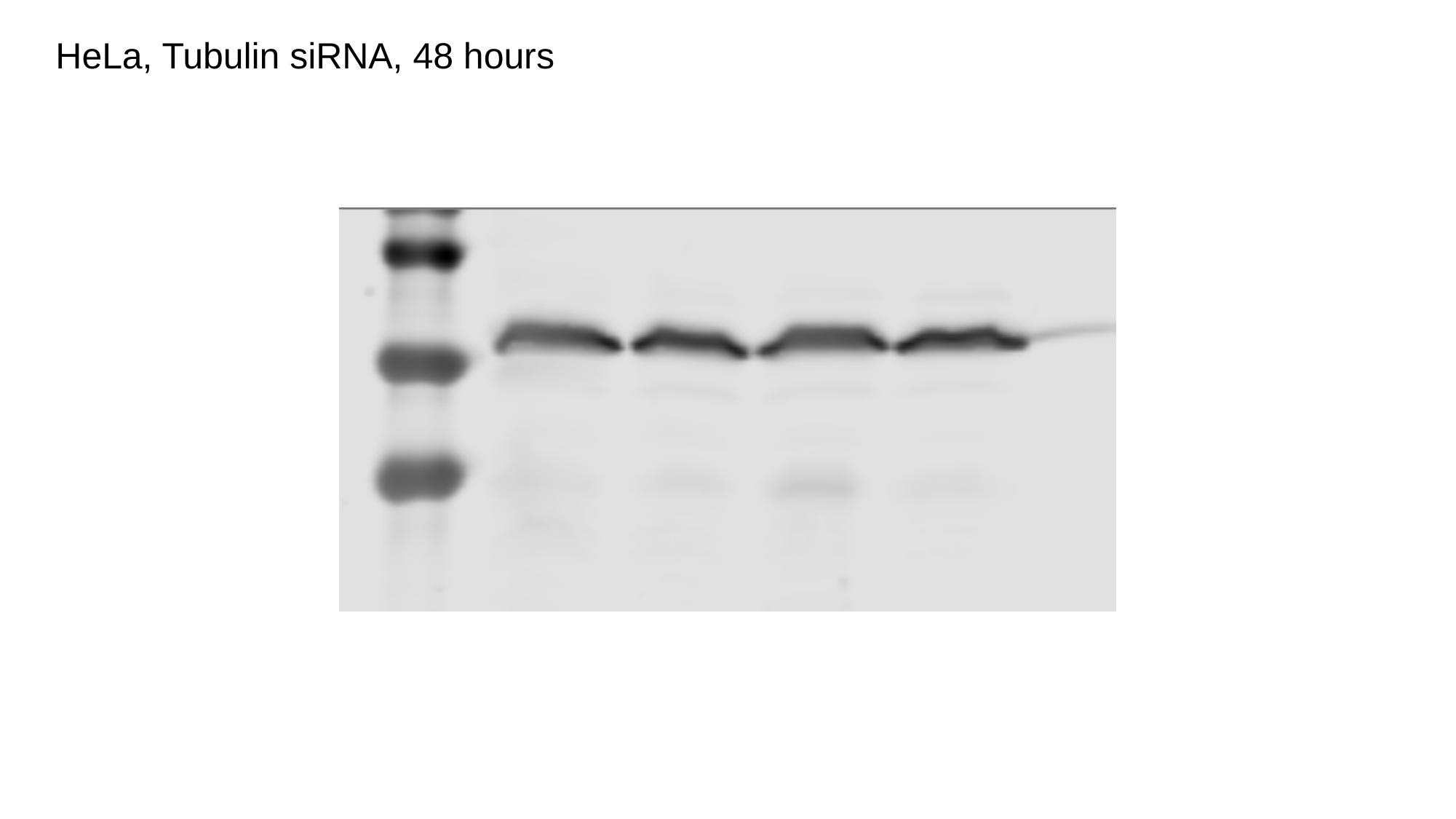

HeLa, Tubulin siRNA, 48 hours
