## Supplementary material for "BRD4 Prevents R-Loop Formation and Transcription-Replication Conflicts by Ensuring Efficient Transcription Elongation": Source Data: 20190604_A.pptx

### Slide 1
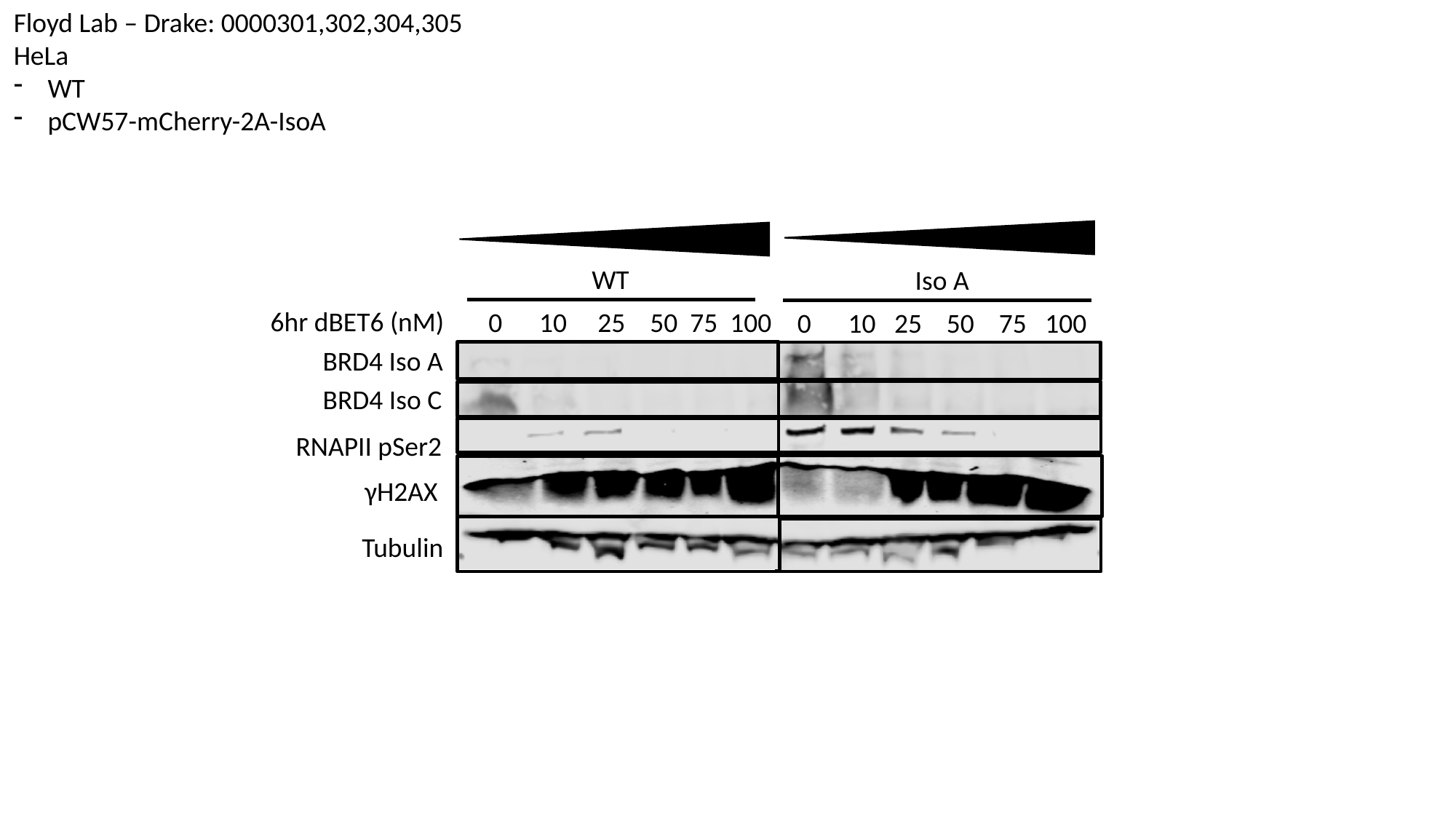

Floyd Lab – Drake: 0000301,302,304,305
HeLa
WT
pCW57-mCherry-2A-IsoA
WT
Iso A
6hr dBET6 (nM)
 0 10 25 50 75 100
0 10 25 50 75 100
BRD4 Iso A
BRD4 Iso C
RNAPII pSer2
γH2AX
Tubulin

### Slide 2
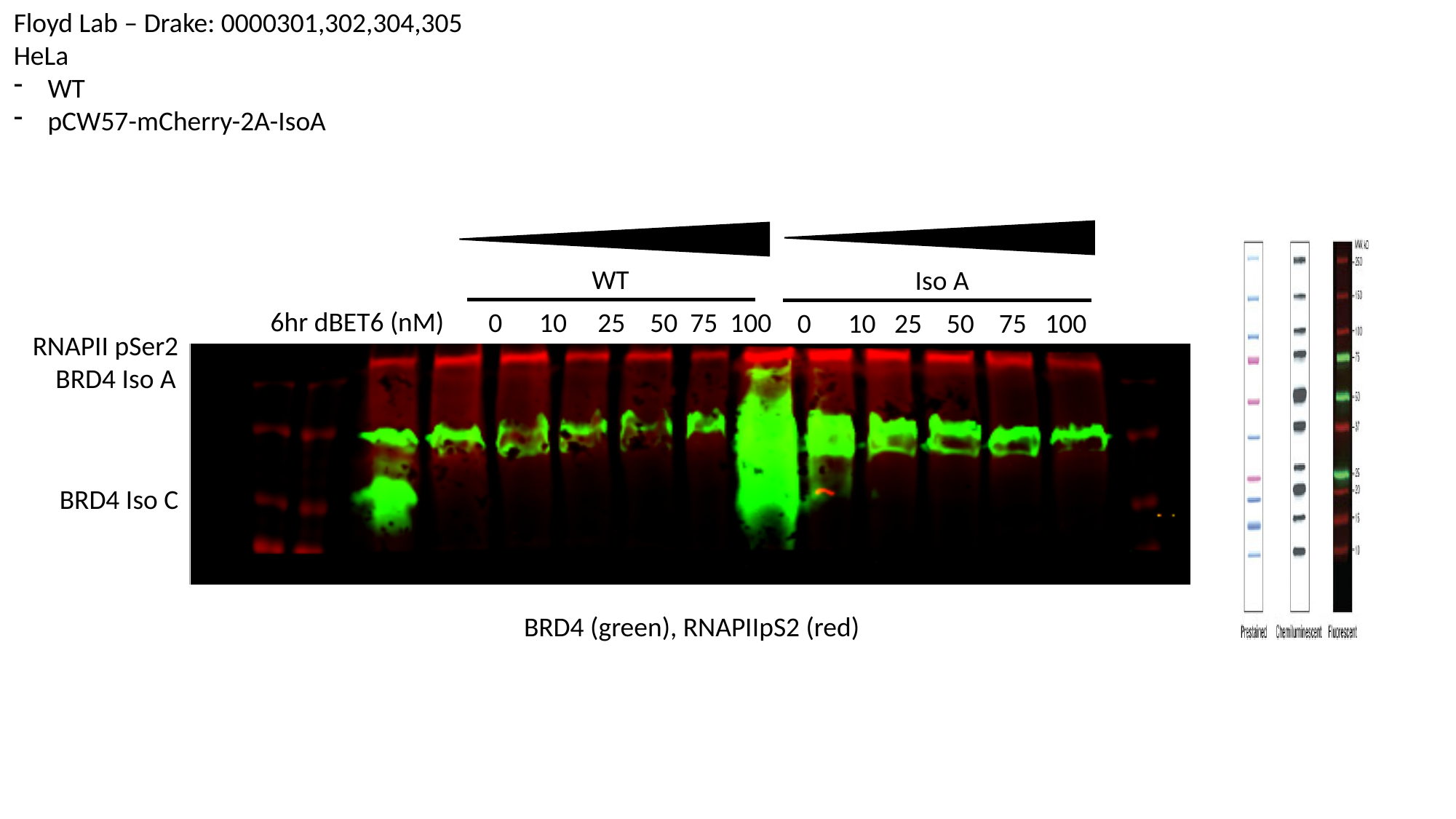

Floyd Lab – Drake: 0000301,302,304,305
HeLa
WT
pCW57-mCherry-2A-IsoA
WT
Iso A
6hr dBET6 (nM)
 0 10 25 50 75 100
0 10 25 50 75 100
RNAPII pSer2
BRD4 Iso A
BRD4 Iso C
BRD4 (green), RNAPIIpS2 (red)

### Slide 3
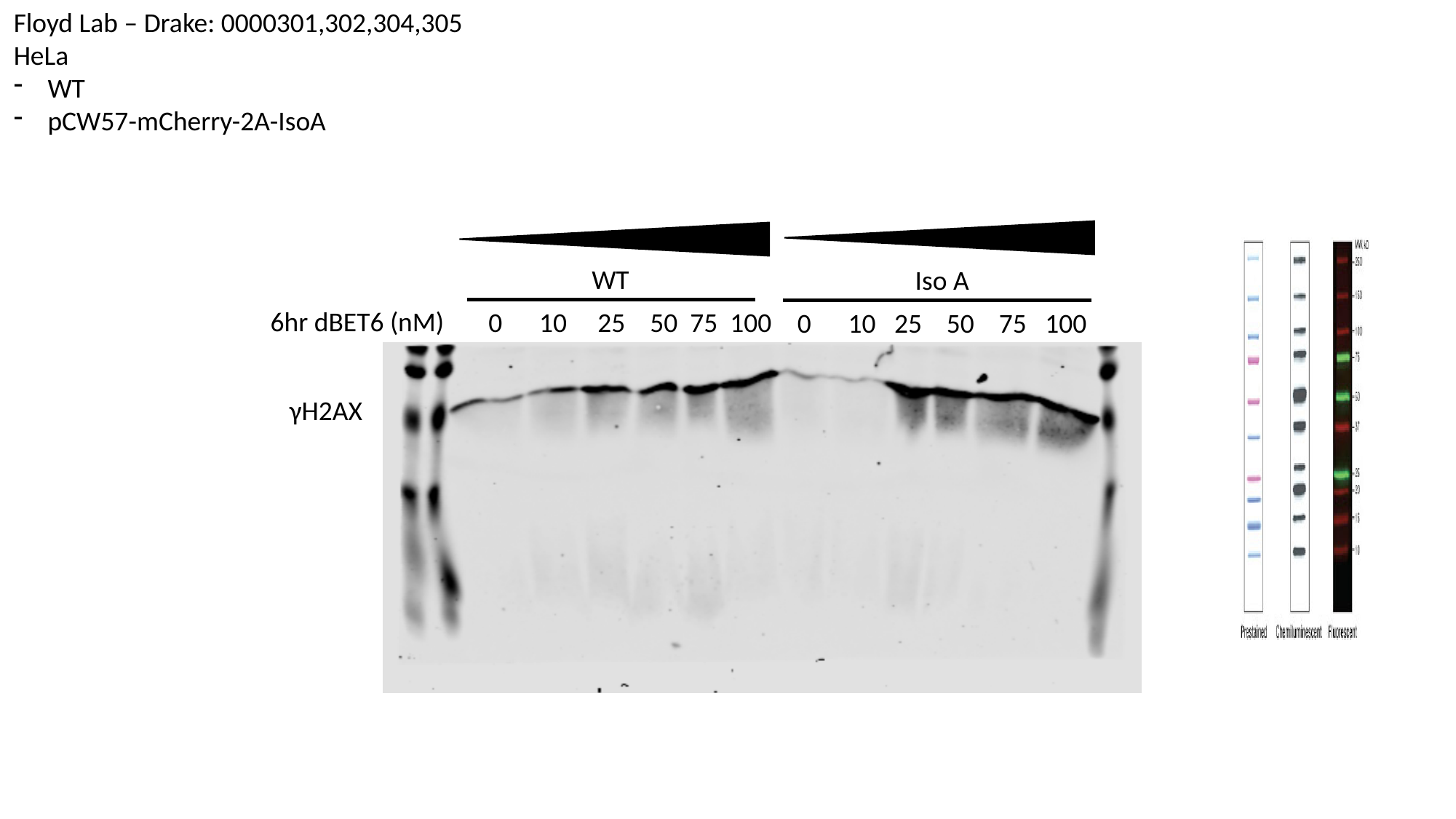

Floyd Lab – Drake: 0000301,302,304,305
HeLa
WT
pCW57-mCherry-2A-IsoA
WT
Iso A
6hr dBET6 (nM)
 0 10 25 50 75 100
0 10 25 50 75 100
γH2AX

### Slide 4
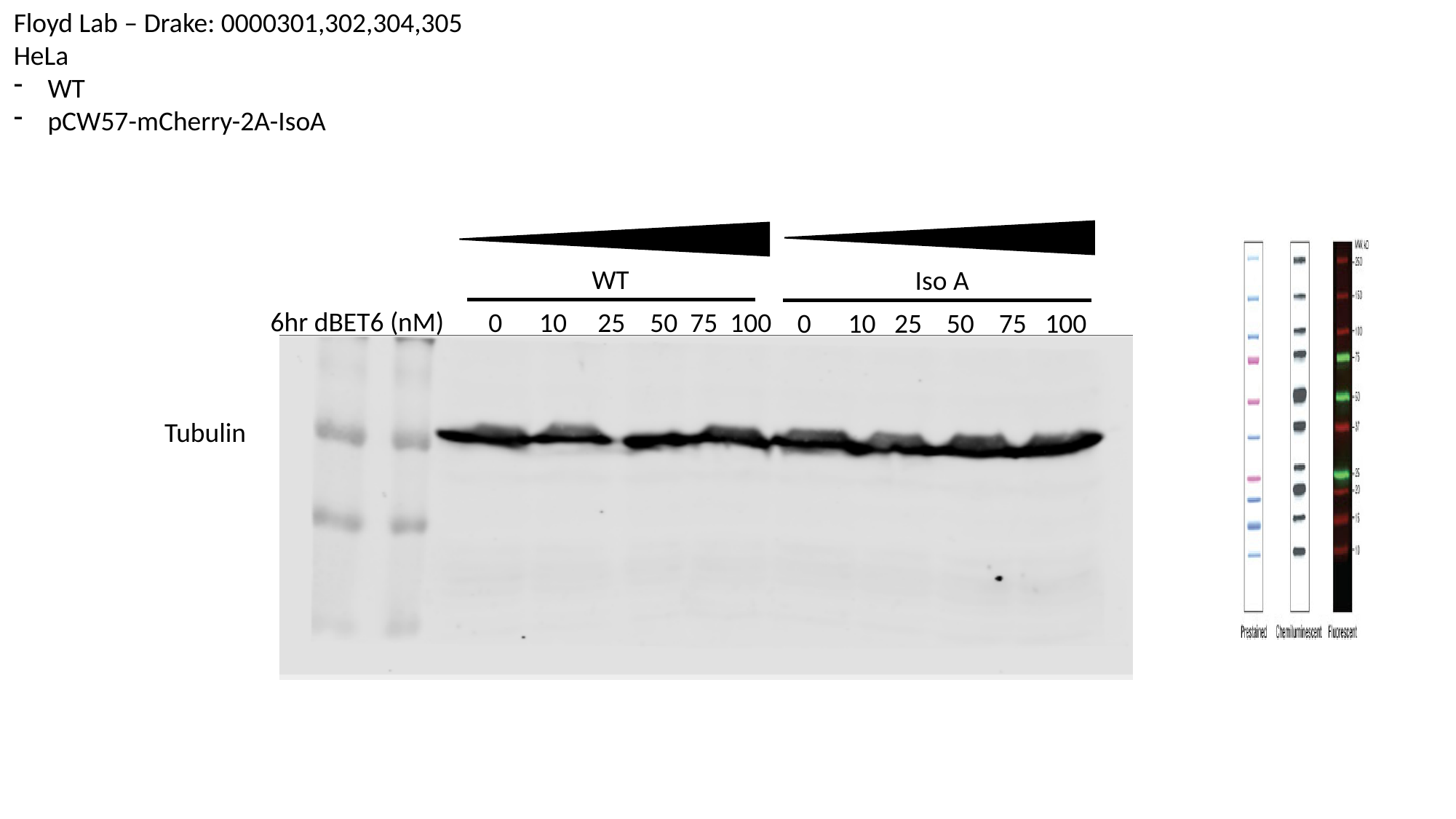

Floyd Lab – Drake: 0000301,302,304,305
HeLa
WT
pCW57-mCherry-2A-IsoA
WT
Iso A
6hr dBET6 (nM)
 0 10 25 50 75 100
0 10 25 50 75 100
Tubulin

### Slide 5
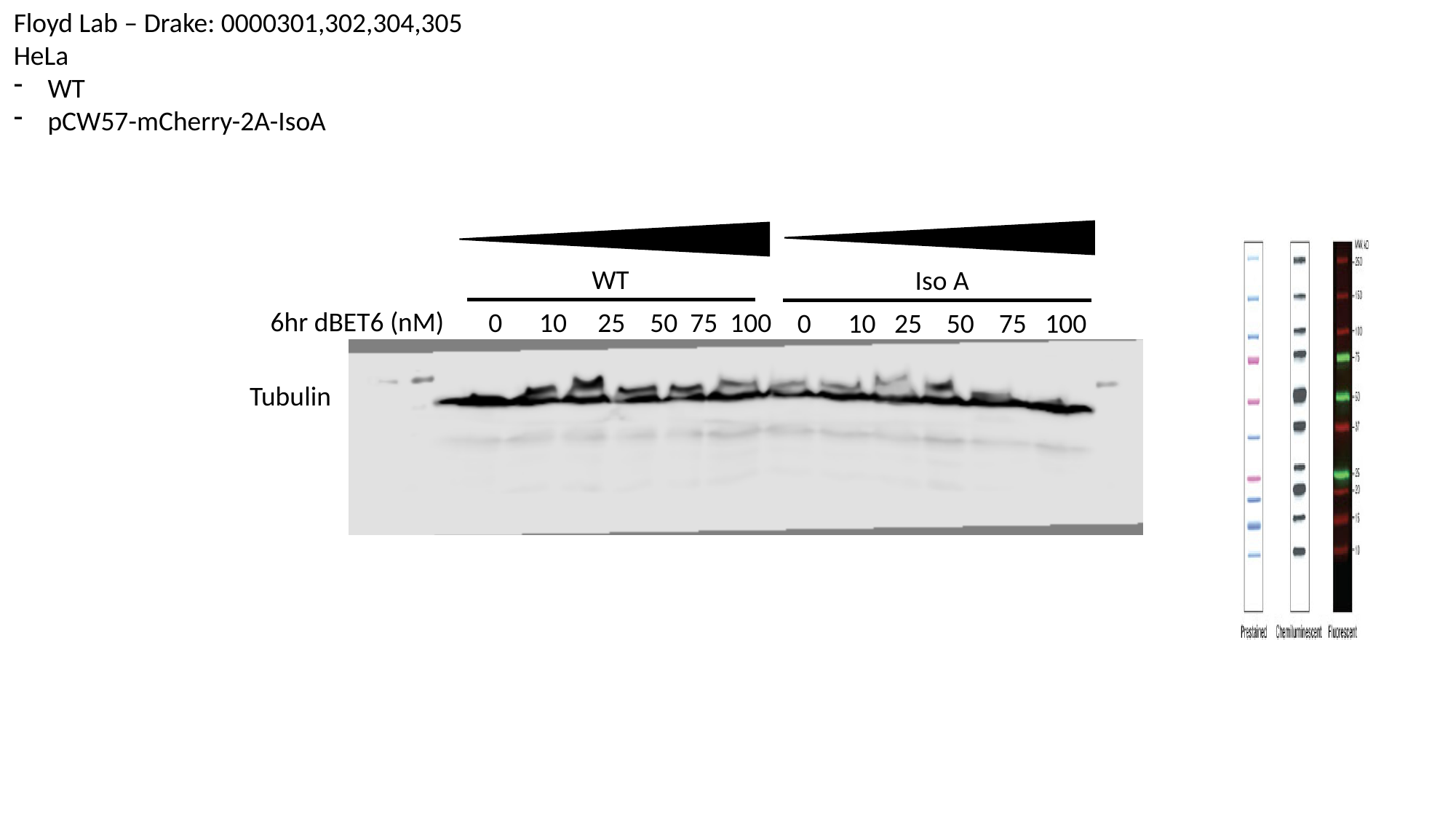

Floyd Lab – Drake: 0000301,302,304,305
HeLa
WT
pCW57-mCherry-2A-IsoA
WT
Iso A
6hr dBET6 (nM)
 0 10 25 50 75 100
0 10 25 50 75 100
Tubulin
