## Supplementary material for "BRD4 Prevents R-Loop Formation and Transcription-Replication Conflicts by Ensuring Efficient Transcription Elongation": Source Data: 20180626 - RNaseH1 D210N, WKKD V5 IP.pptx

### Slide 1
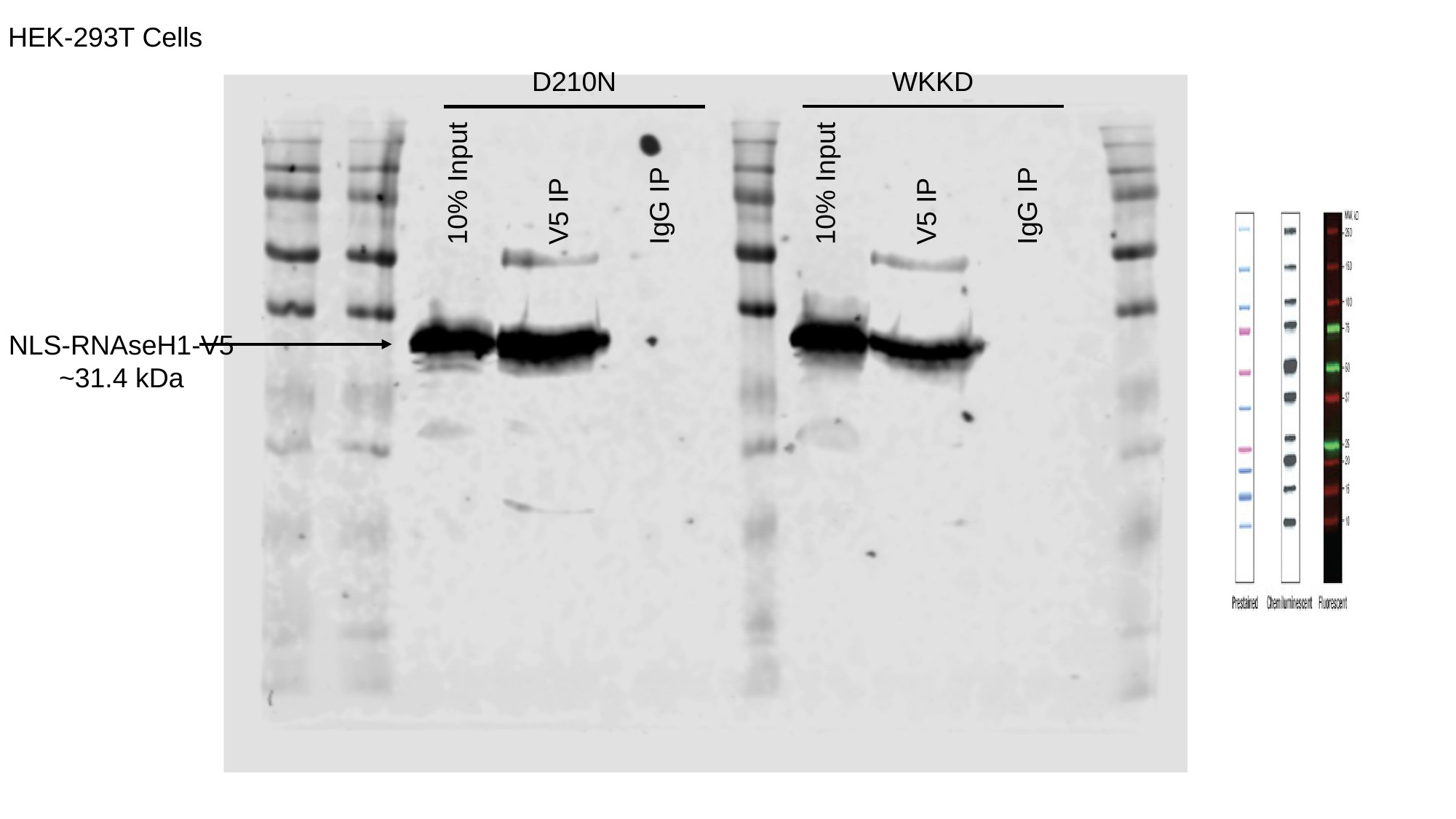

HEK-293T Cells
D210N
WKKD
10% Input
V5 IP
IgG IP
10% Input
V5 IP
IgG IP
NLS-RNAseH1-V5
~31.4 kDa

### Slide 2
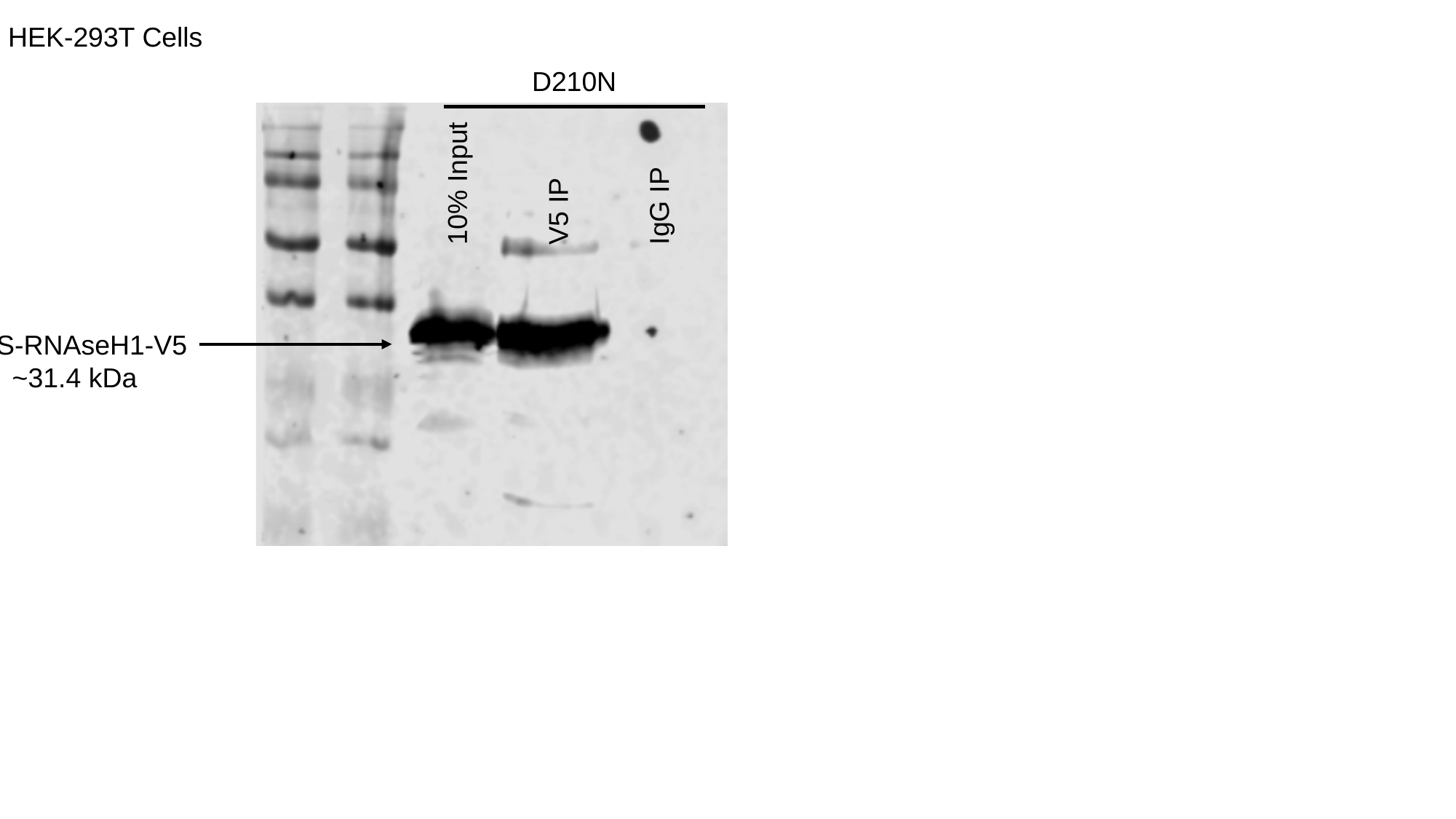

HEK-293T Cells
D210N
10% Input
V5 IP
IgG IP
NLS-RNAseH1-V5
~31.4 kDa
