## Supplementary material for "BRD4 Prevents R-Loop Formation and Transcription-Replication Conflicts by Ensuring Efficient Transcription Elongation": Source Data: 20190206 - V5-RH1 WT and WKKD WB.pptx

### Slide 1
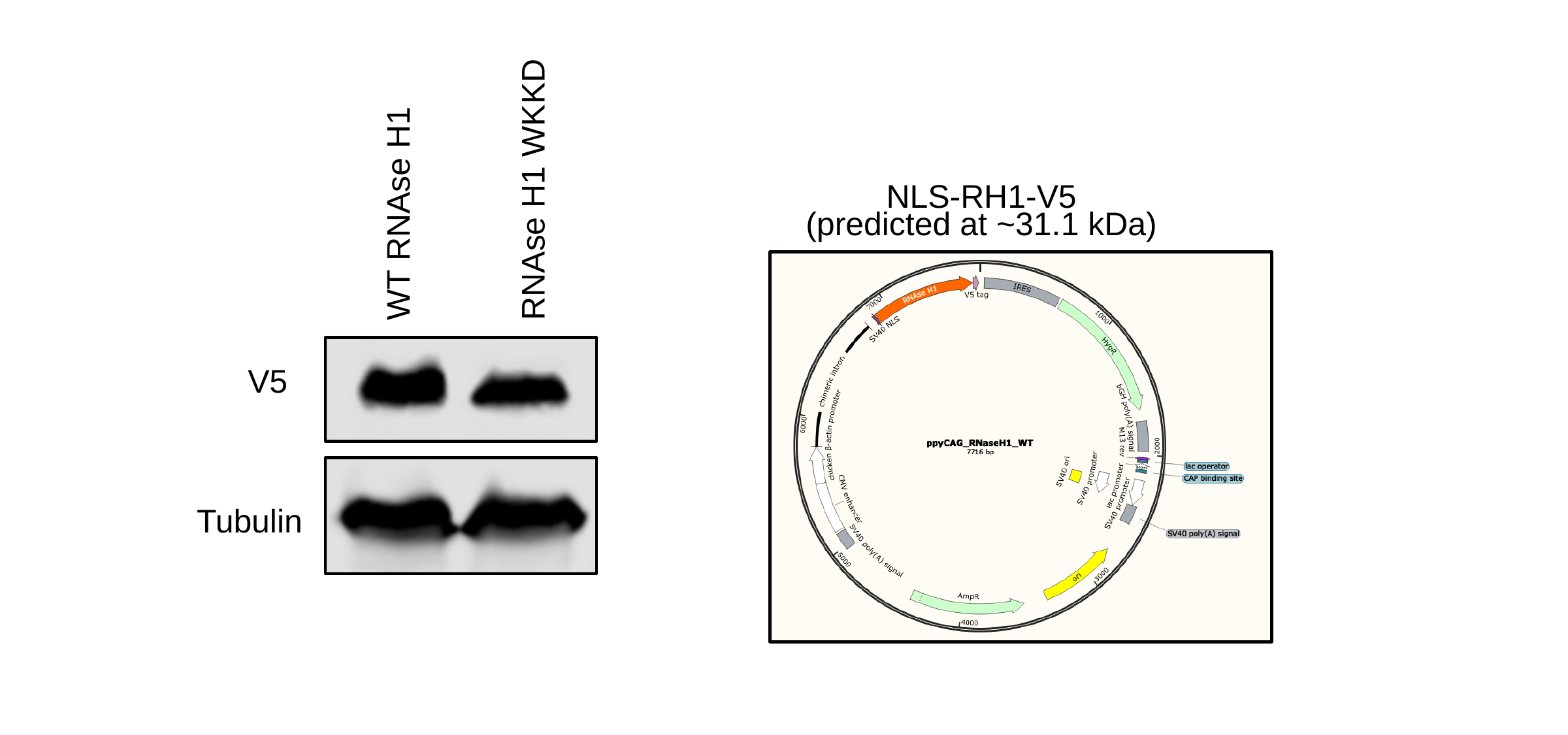

RNAse H1 WKKD
NLS-RH1-V5
WT RNAse H1
(predicted at ~31.1 kDa)
V5
Tubulin

### Slide 2
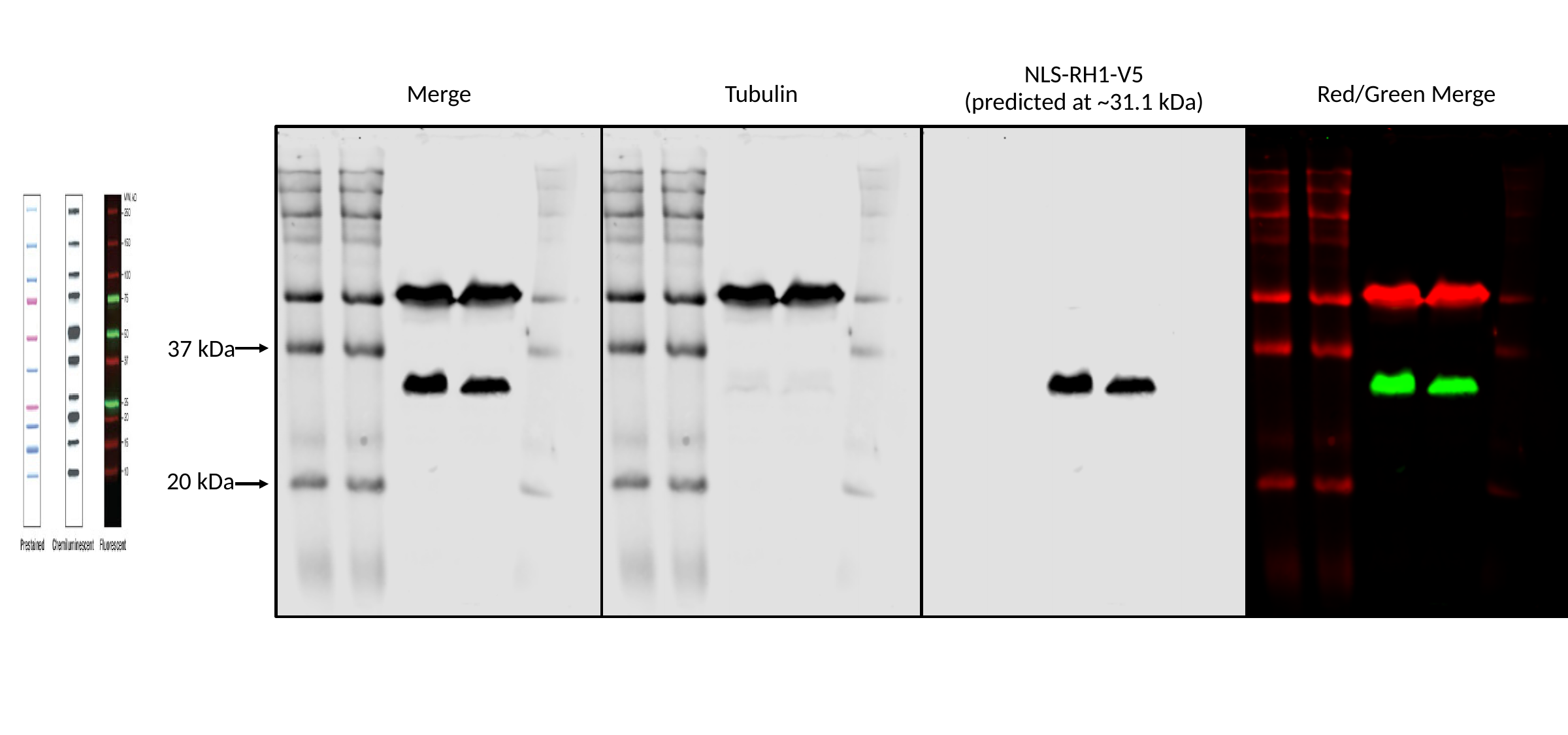

NLS-RH1-V5
Merge
Tubulin
Red/Green Merge
(predicted at ~31.1 kDa)
37 kDa
20 kDa
