## Supplementary material for "BRD4 Prevents R-Loop Formation and Transcription-Replication Conflicts by Ensuring Efficient Transcription Elongation": Source Data: 20191003_SRSF1_SETX_DHX9.pptx

### Slide 1
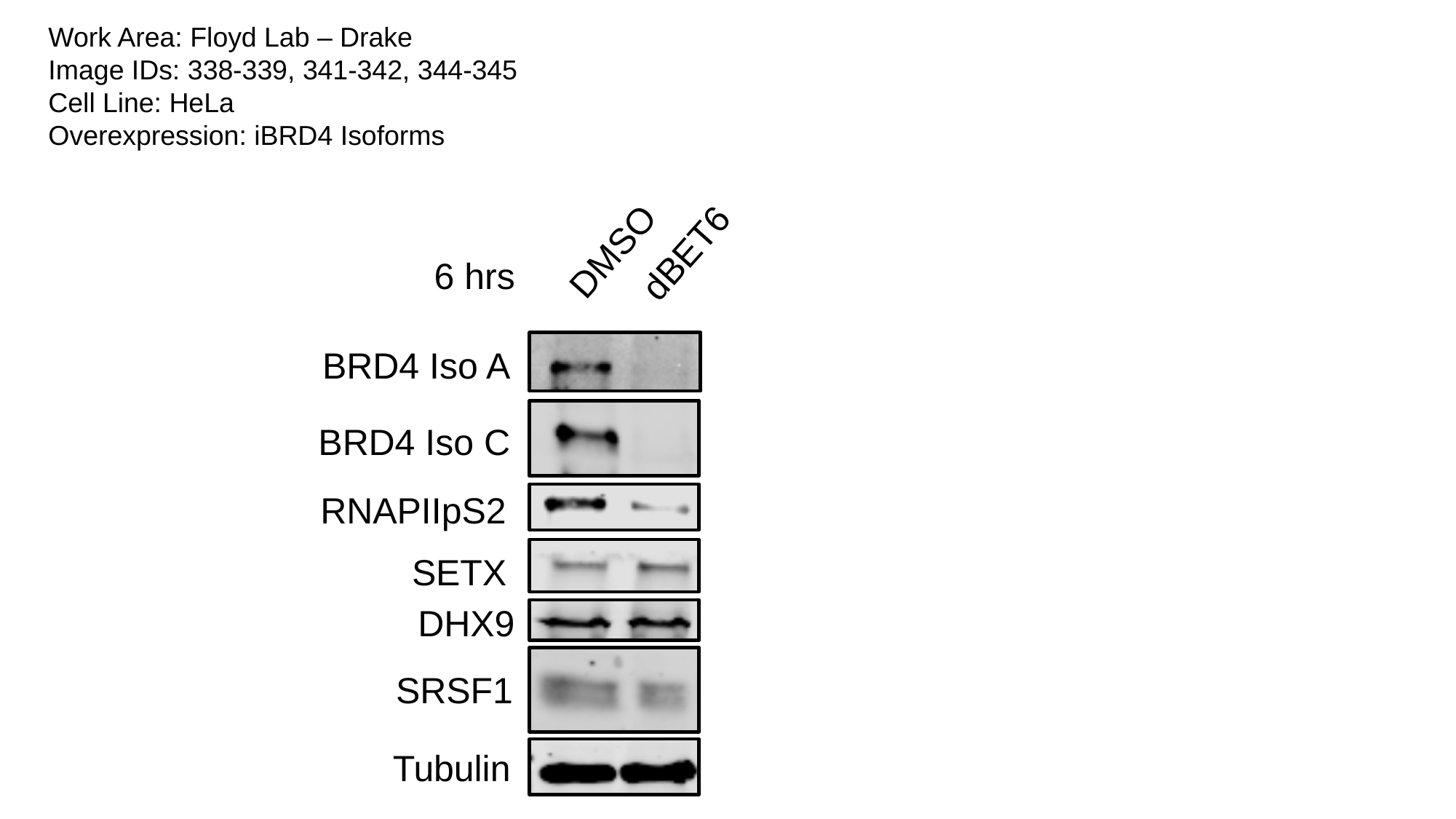

Work Area: Floyd Lab – Drake
Image IDs: 338-339, 341-342, 344-345
Cell Line: HeLa
Overexpression: iBRD4 Isoforms
DMSO
dBET6
6 hrs
BRD4 Iso A
BRD4 Iso C
RNAPIIpS2
SETX
DHX9
SRSF1
Tubulin

### Slide 2
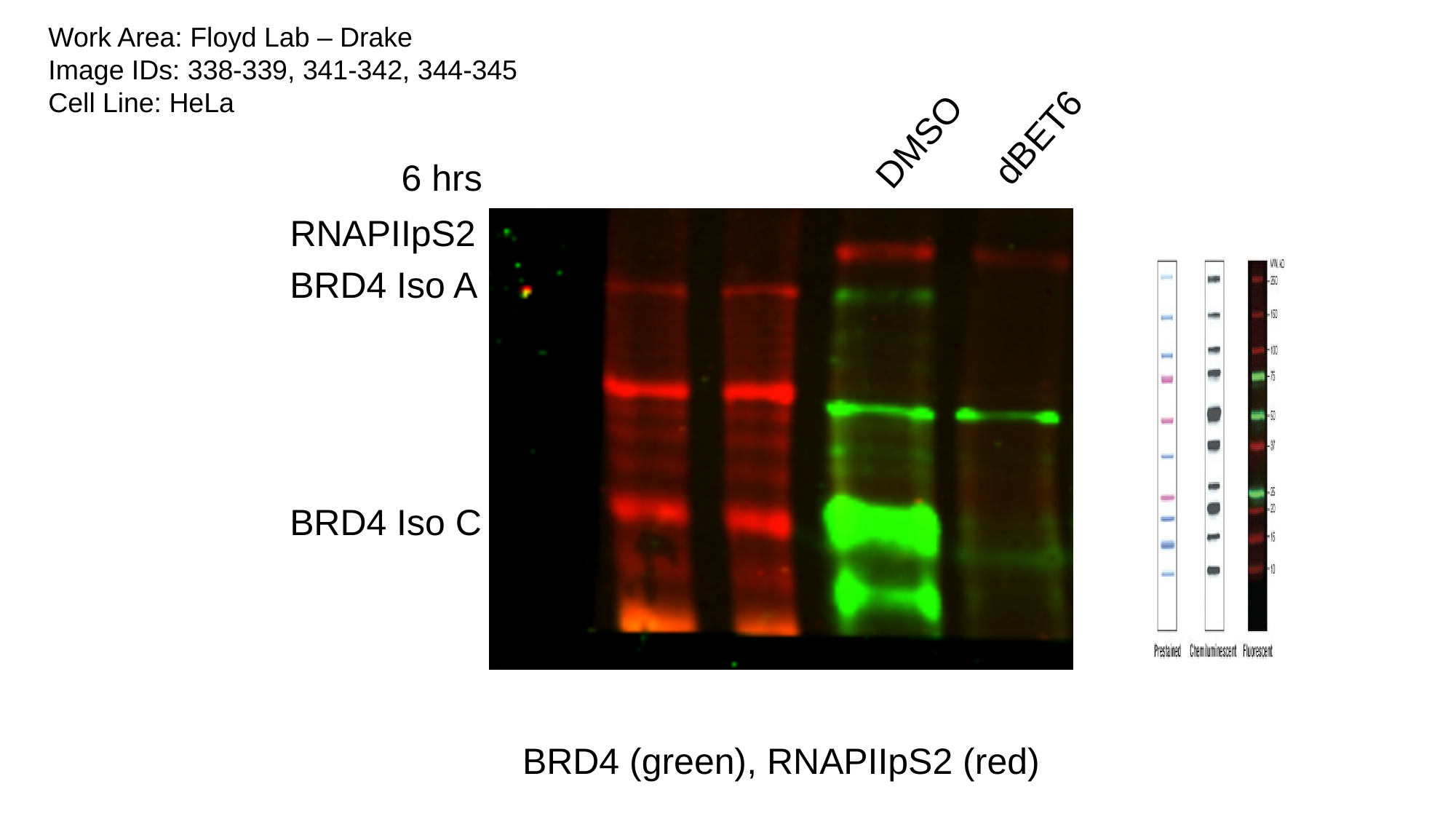

Work Area: Floyd Lab – Drake
Image IDs: 338-339, 341-342, 344-345
Cell Line: HeLa
dBET6
DMSO
6 hrs
RNAPIIpS2
BRD4 Iso A
BRD4 Iso C
BRD4 (green), RNAPIIpS2 (red)

### Slide 3
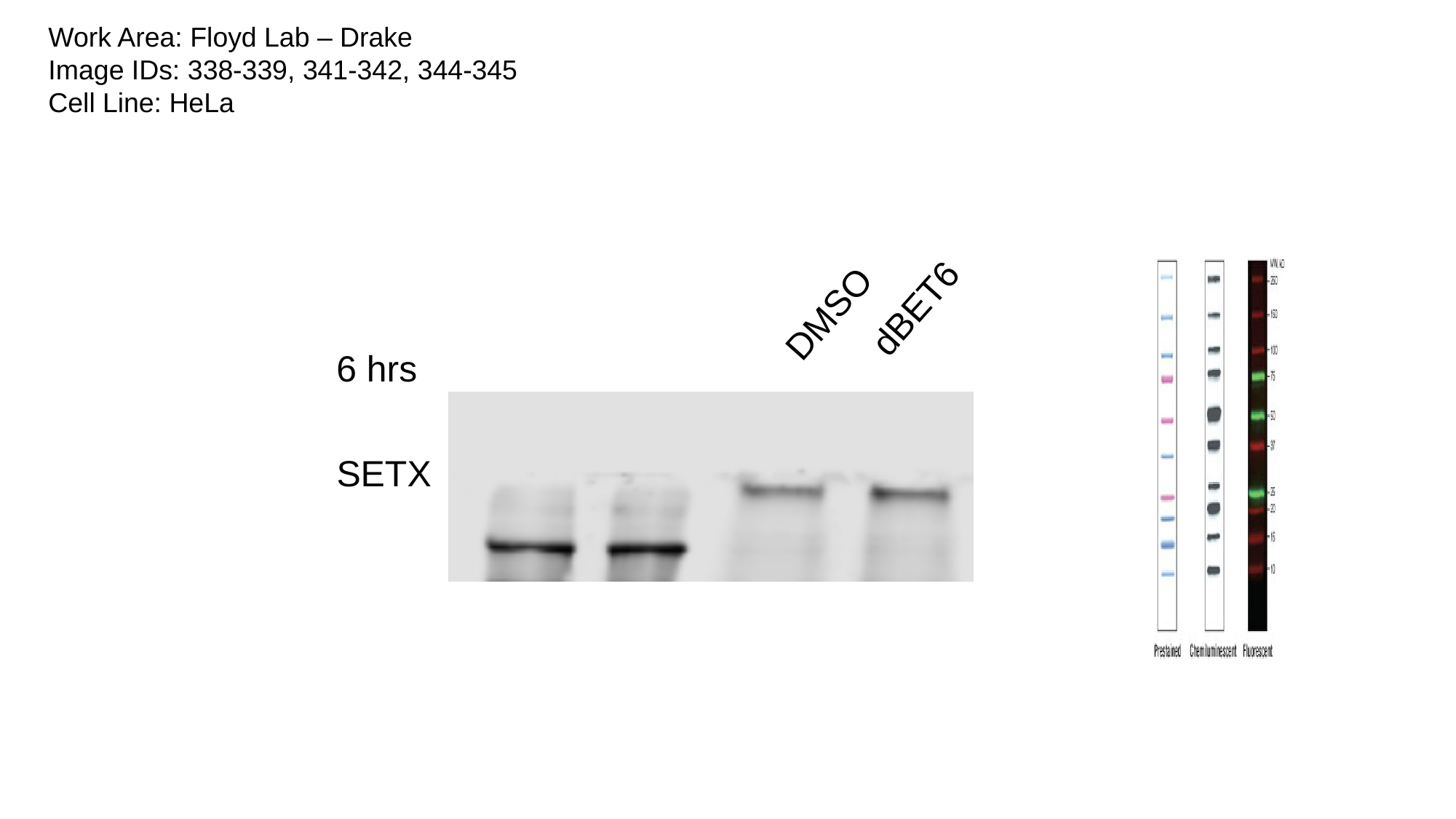

Work Area: Floyd Lab – Drake
Image IDs: 338-339, 341-342, 344-345
Cell Line: HeLa
dBET6
DMSO
6 hrs
SETX

### Slide 4
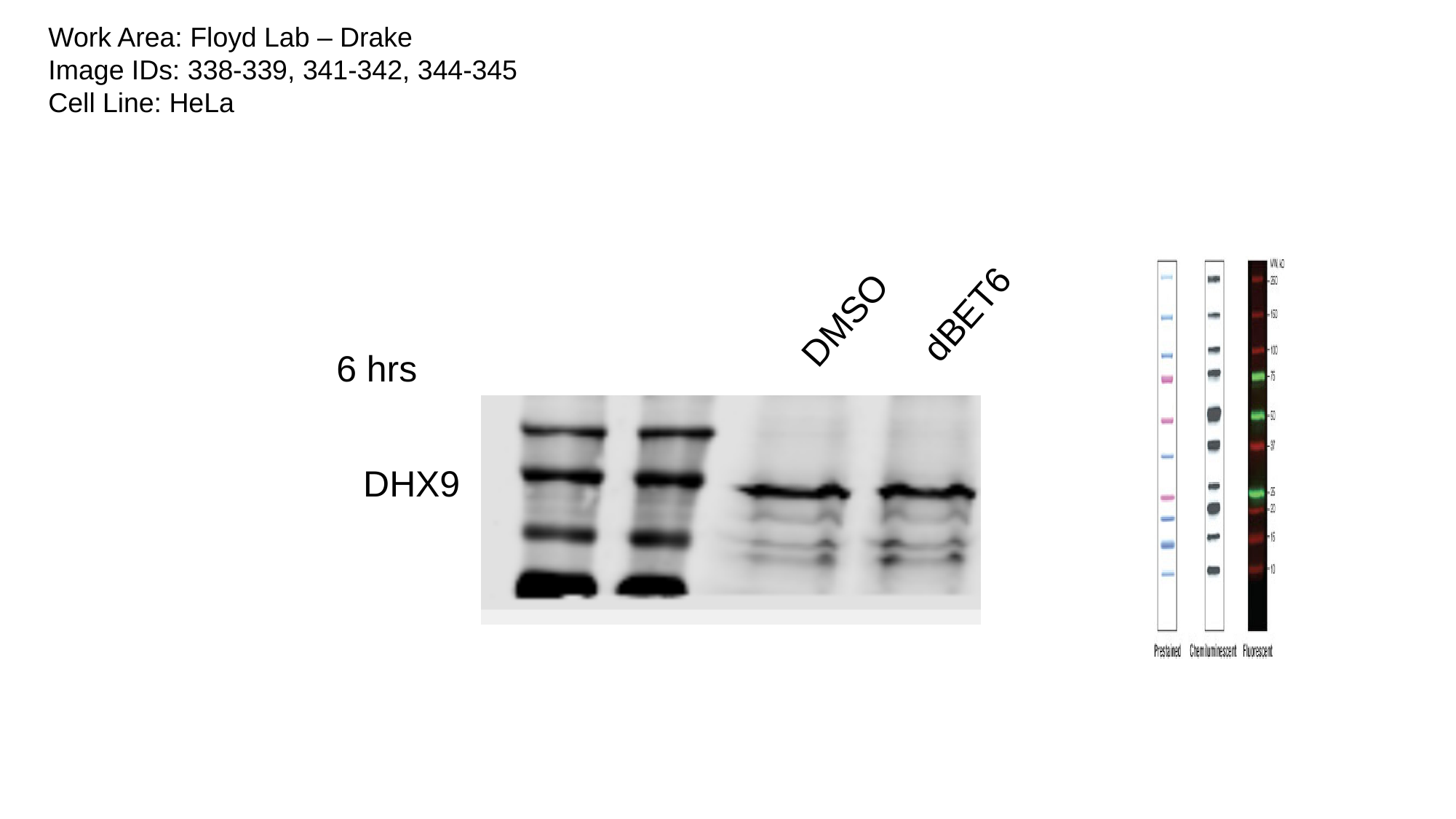

Work Area: Floyd Lab – Drake
Image IDs: 338-339, 341-342, 344-345
Cell Line: HeLa
dBET6
DMSO
6 hrs
DHX9

### Slide 5
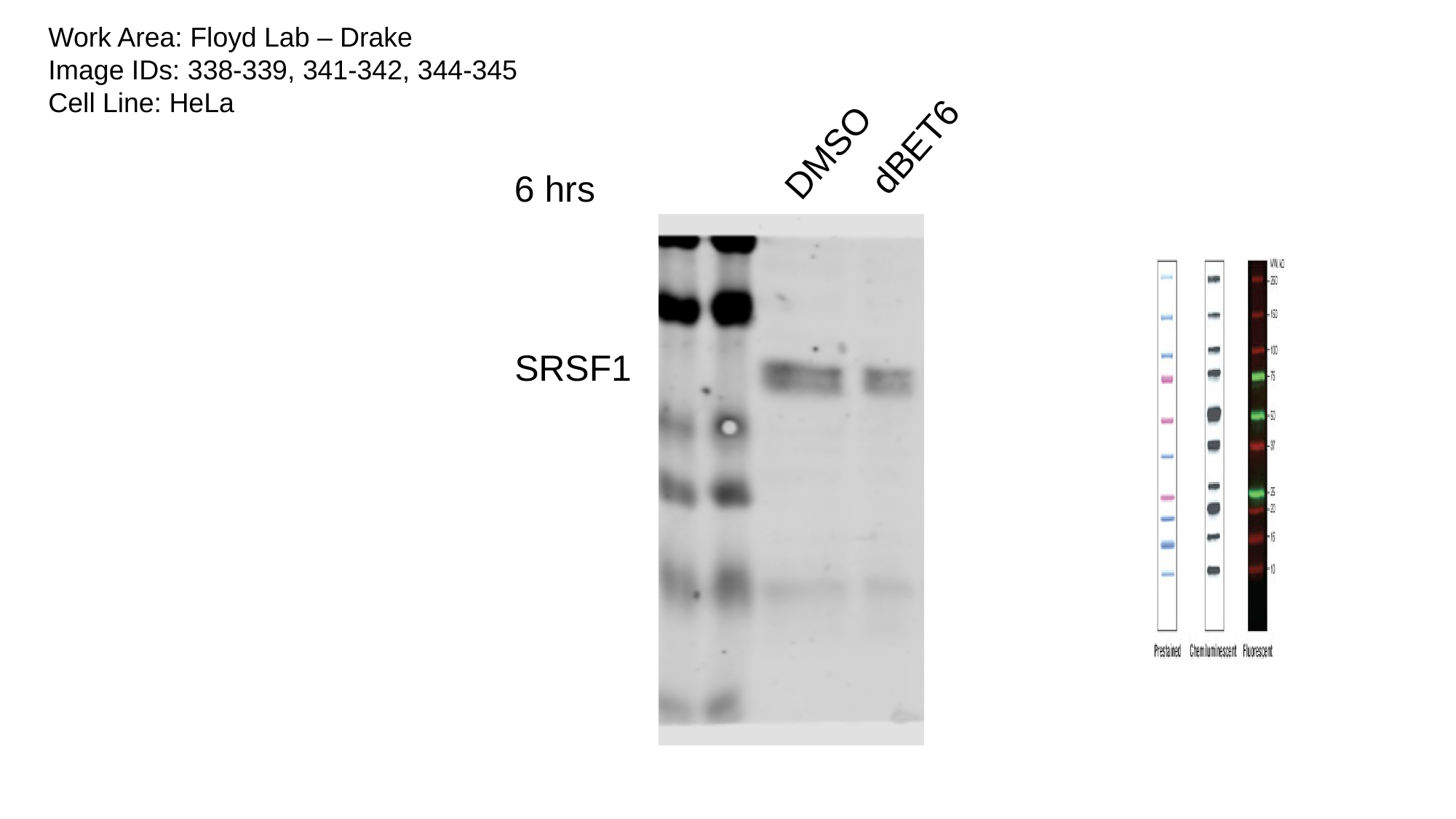

Work Area: Floyd Lab – Drake
Image IDs: 338-339, 341-342, 344-345
Cell Line: HeLa
dBET6
DMSO
6 hrs
SRSF1

### Slide 6
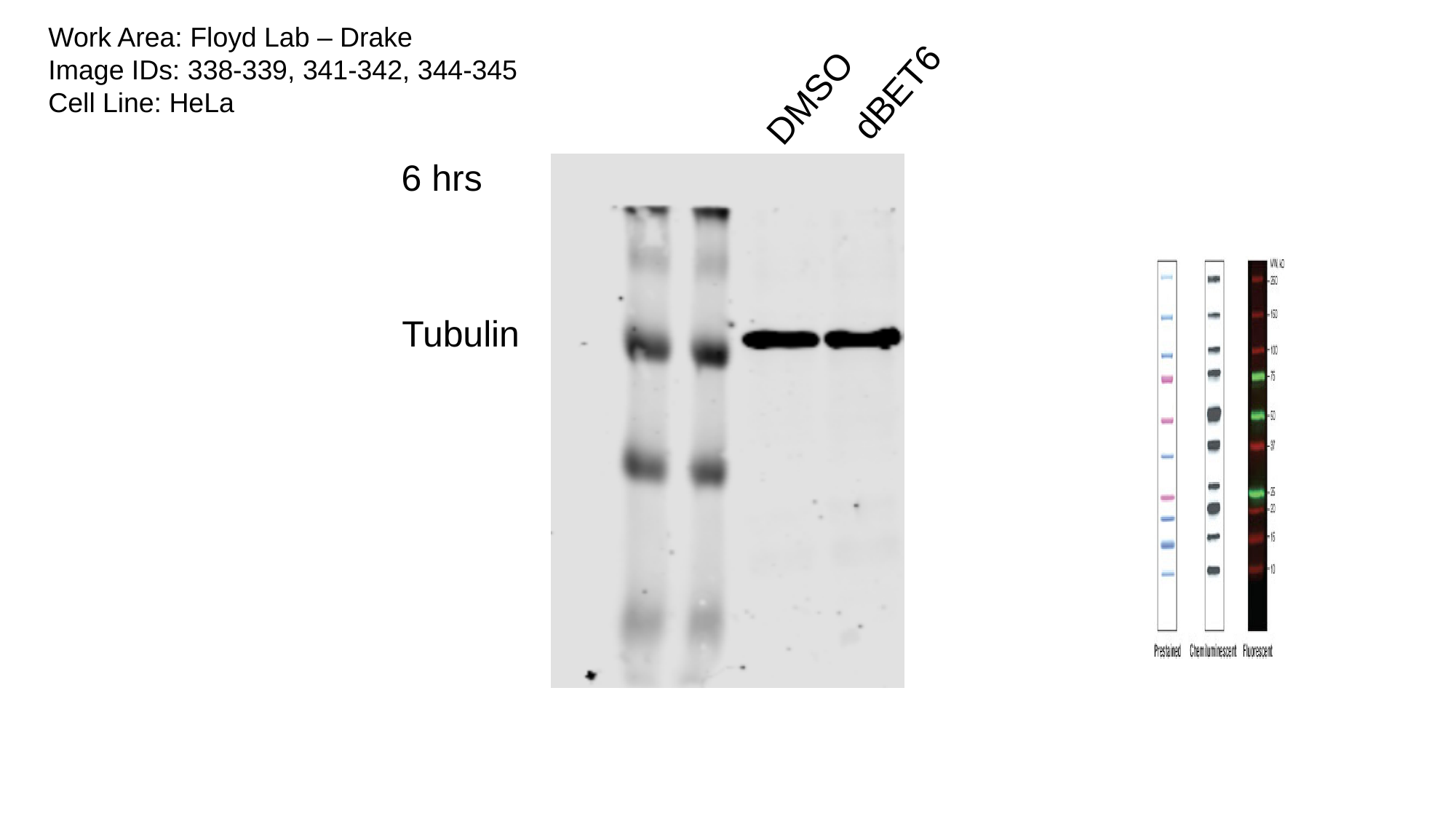

Work Area: Floyd Lab – Drake
Image IDs: 338-339, 341-342, 344-345
Cell Line: HeLa
dBET6
DMSO
6 hrs
Tubulin
