## Supplementary material for "BRD4 Prevents R-Loop Formation and Transcription-Replication Conflicts by Ensuring Efficient Transcription Elongation": Source Data: 20200305_sivarious_dbet6.pptx

### Slide 1
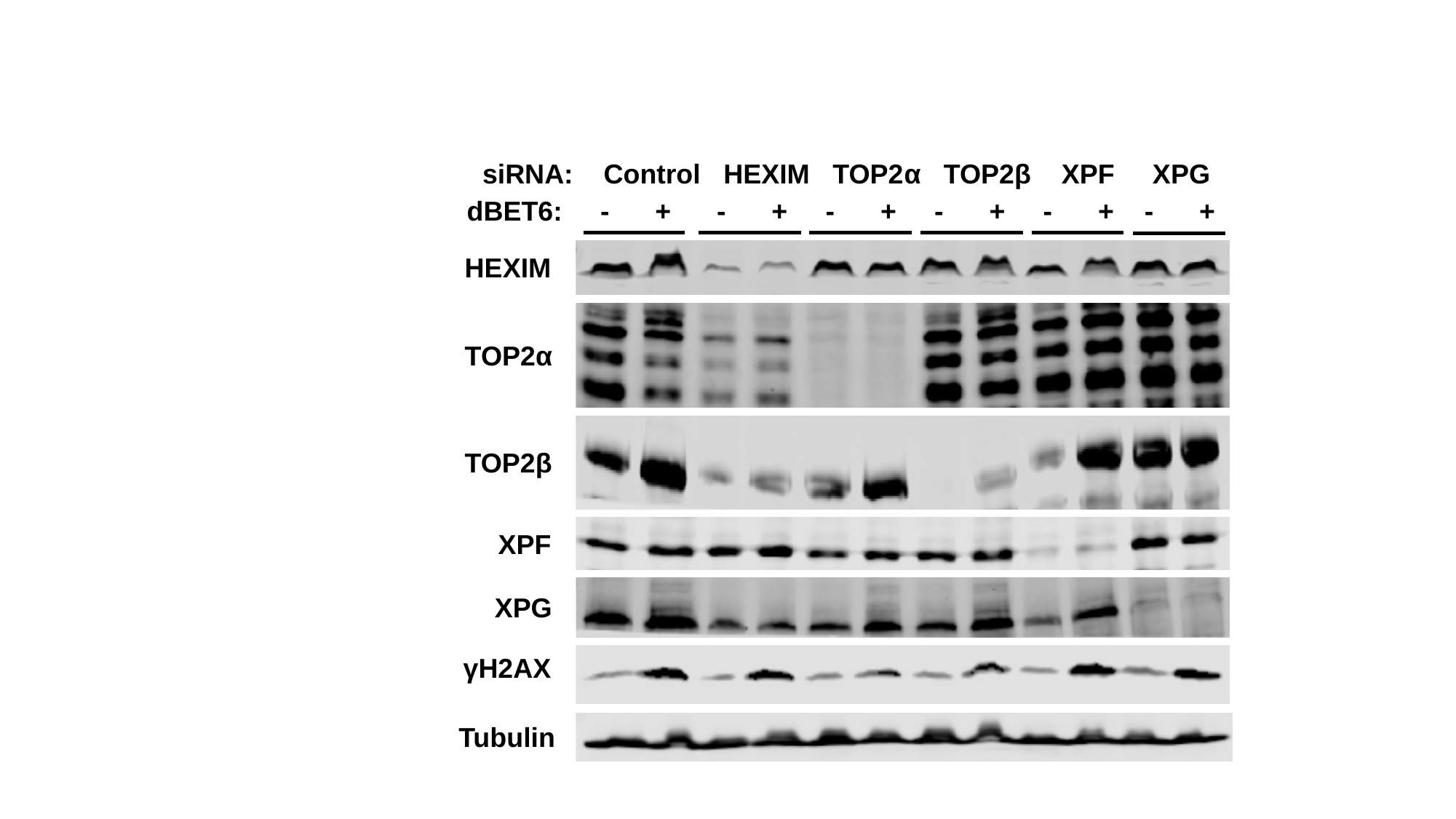

siRNA: Control HEXIM TOP2α TOP2β XPF XPG
dBET6: - + - + - + - + - + - +
HEXIM
TOP2α
TOP2β
XPF
XPG
γH2AX
Tubulin

### Slide 2
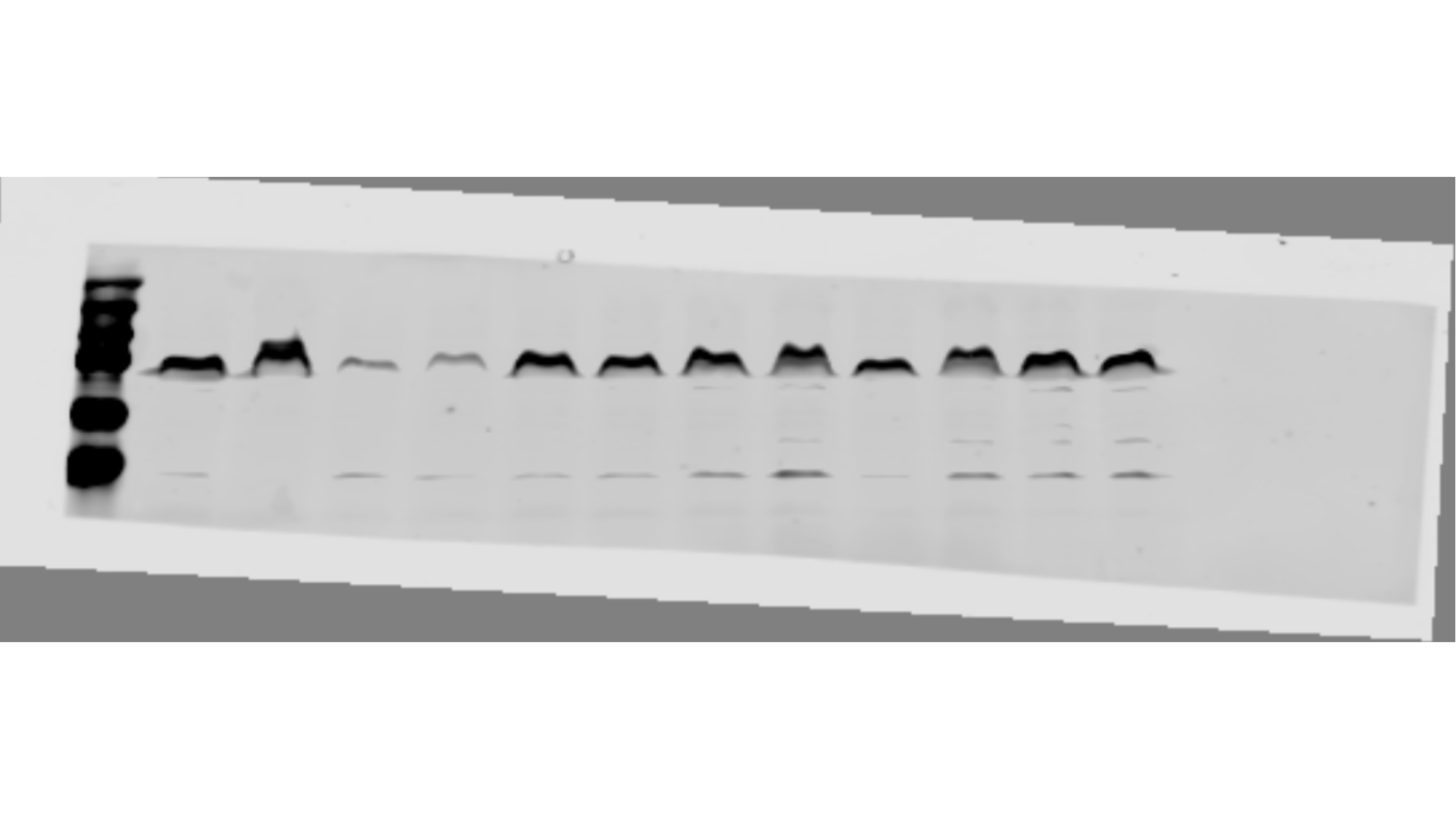

### Slide 3
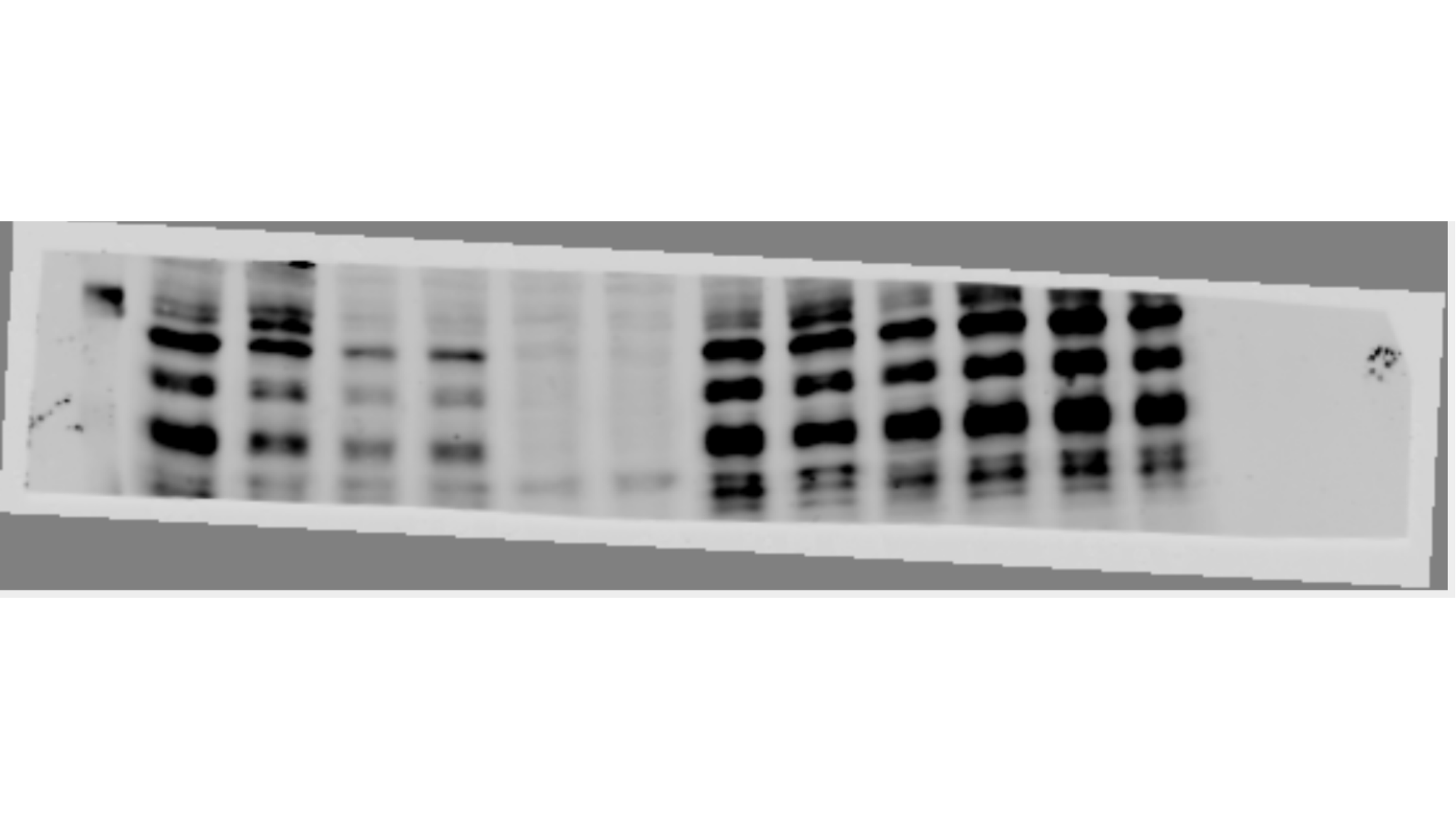

### Slide 4
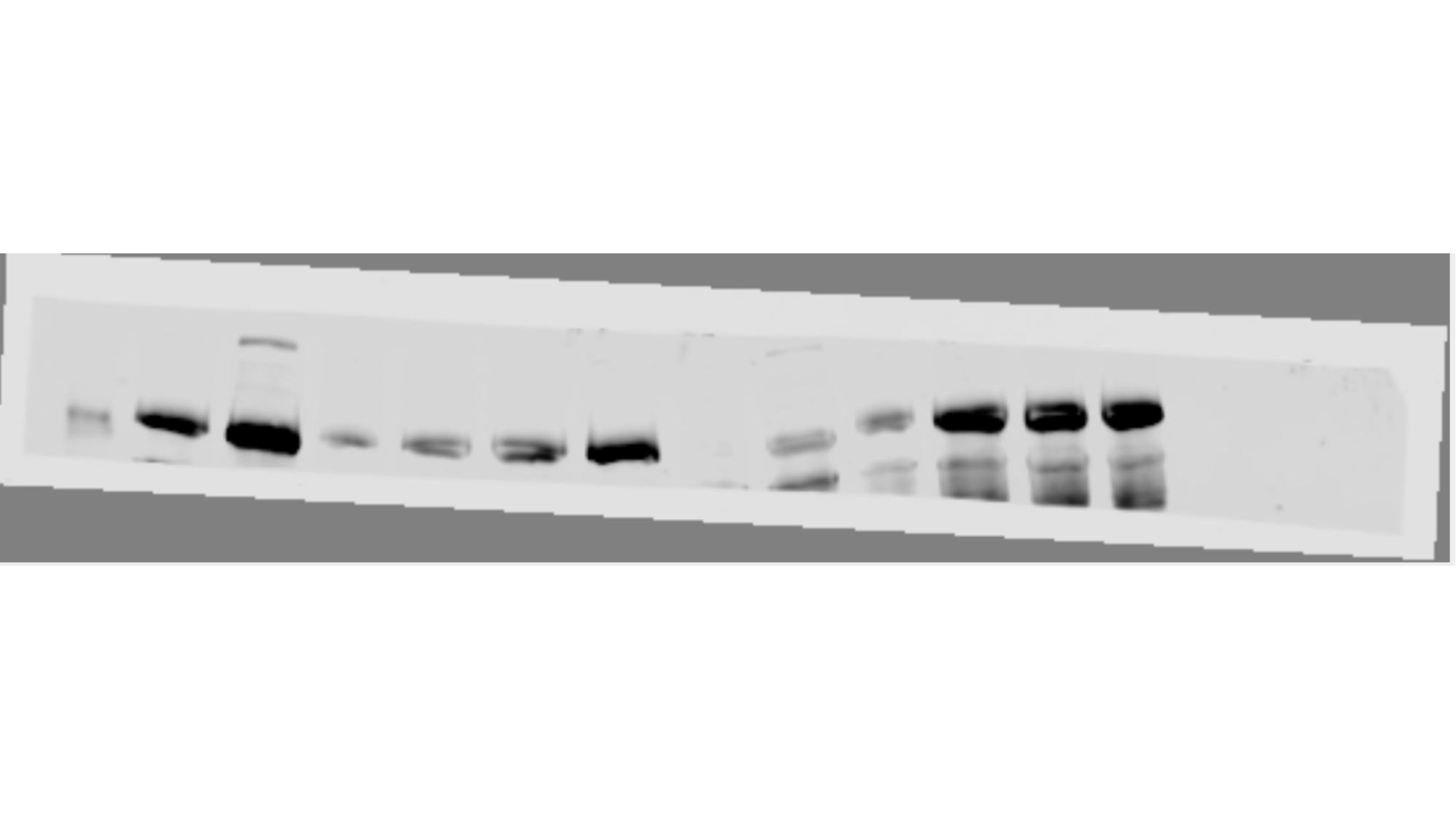

### Slide 5
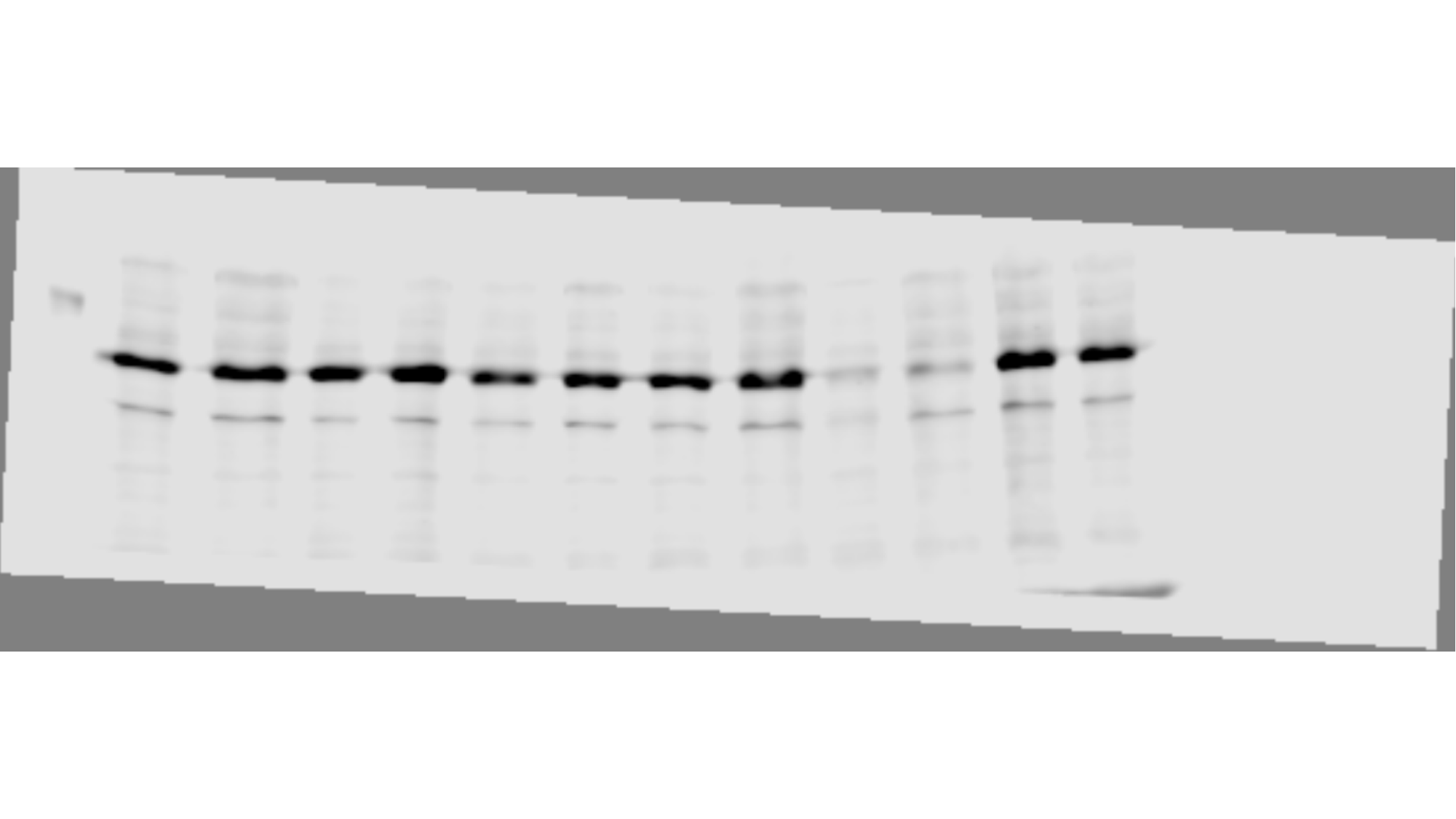

### Slide 6
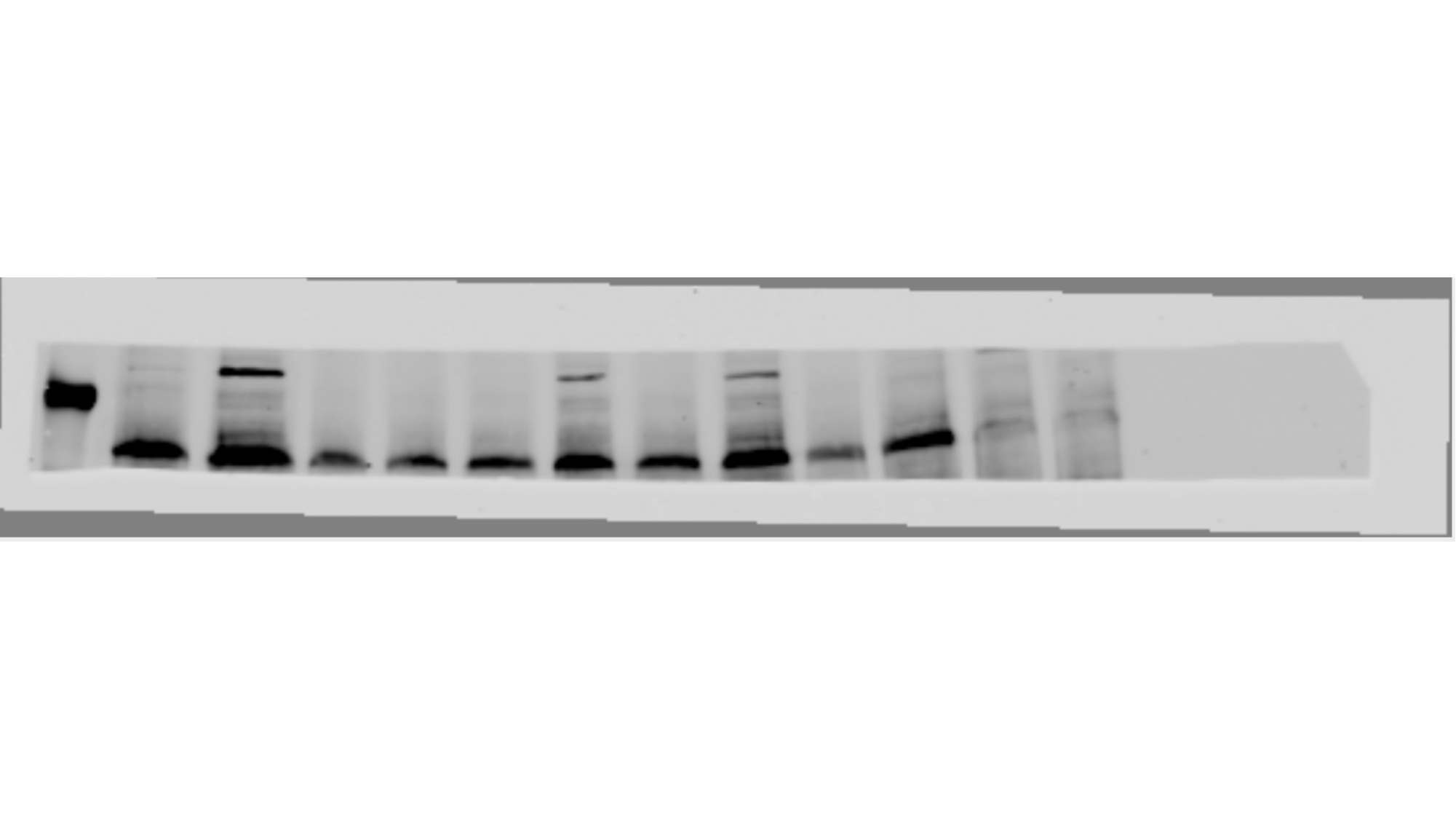

### Slide 7
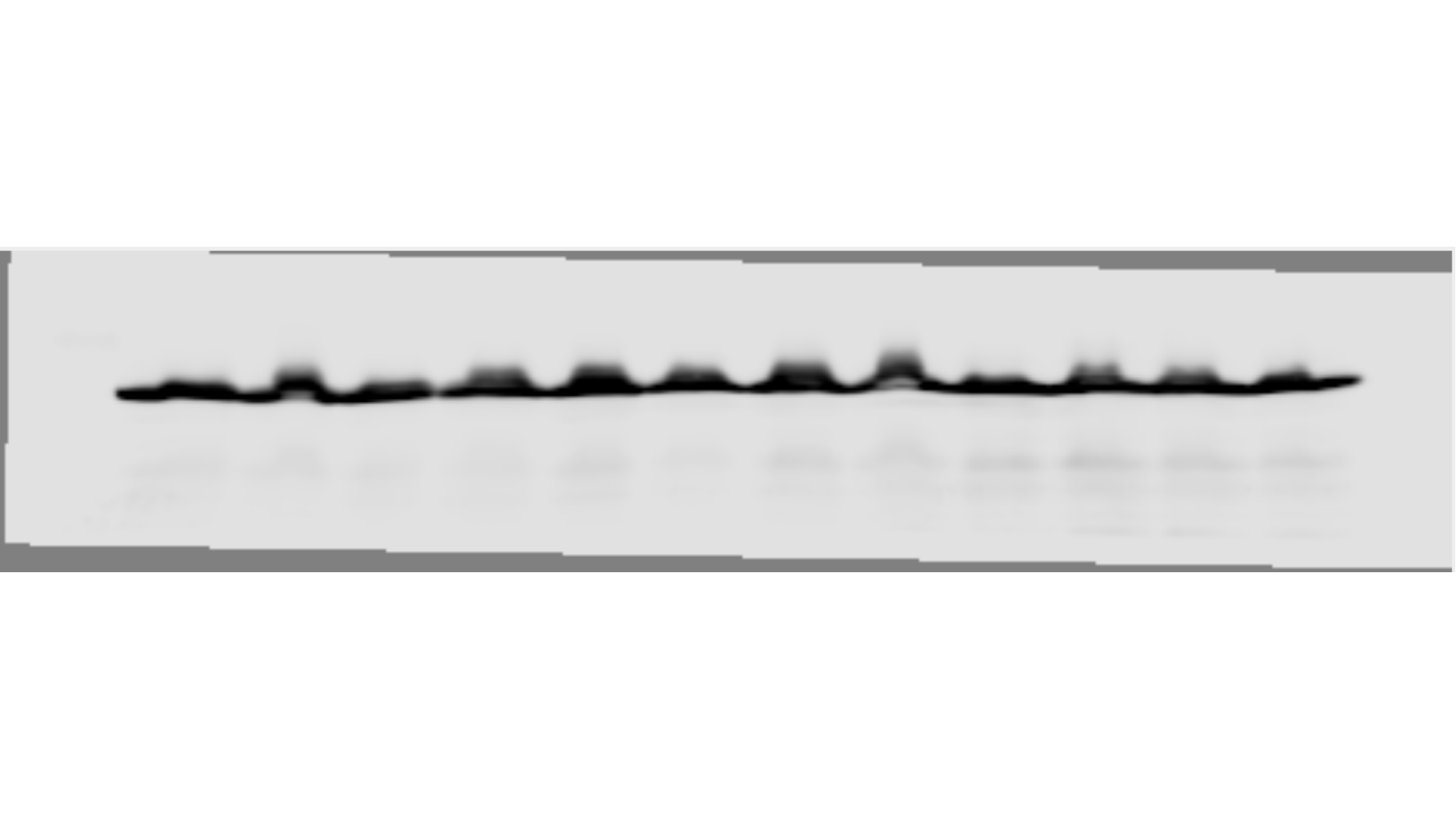

### Slide 8
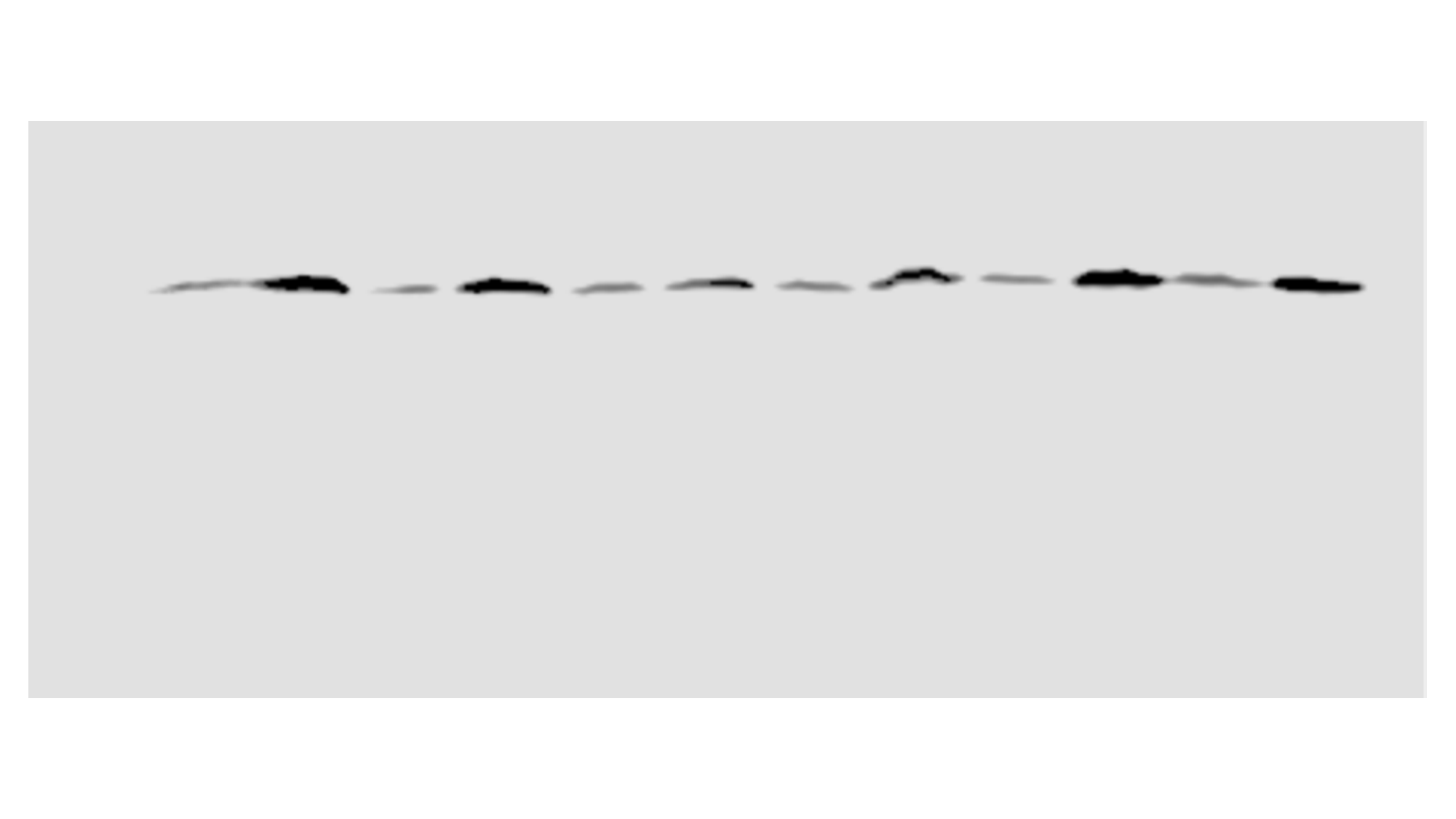

### Slide 9
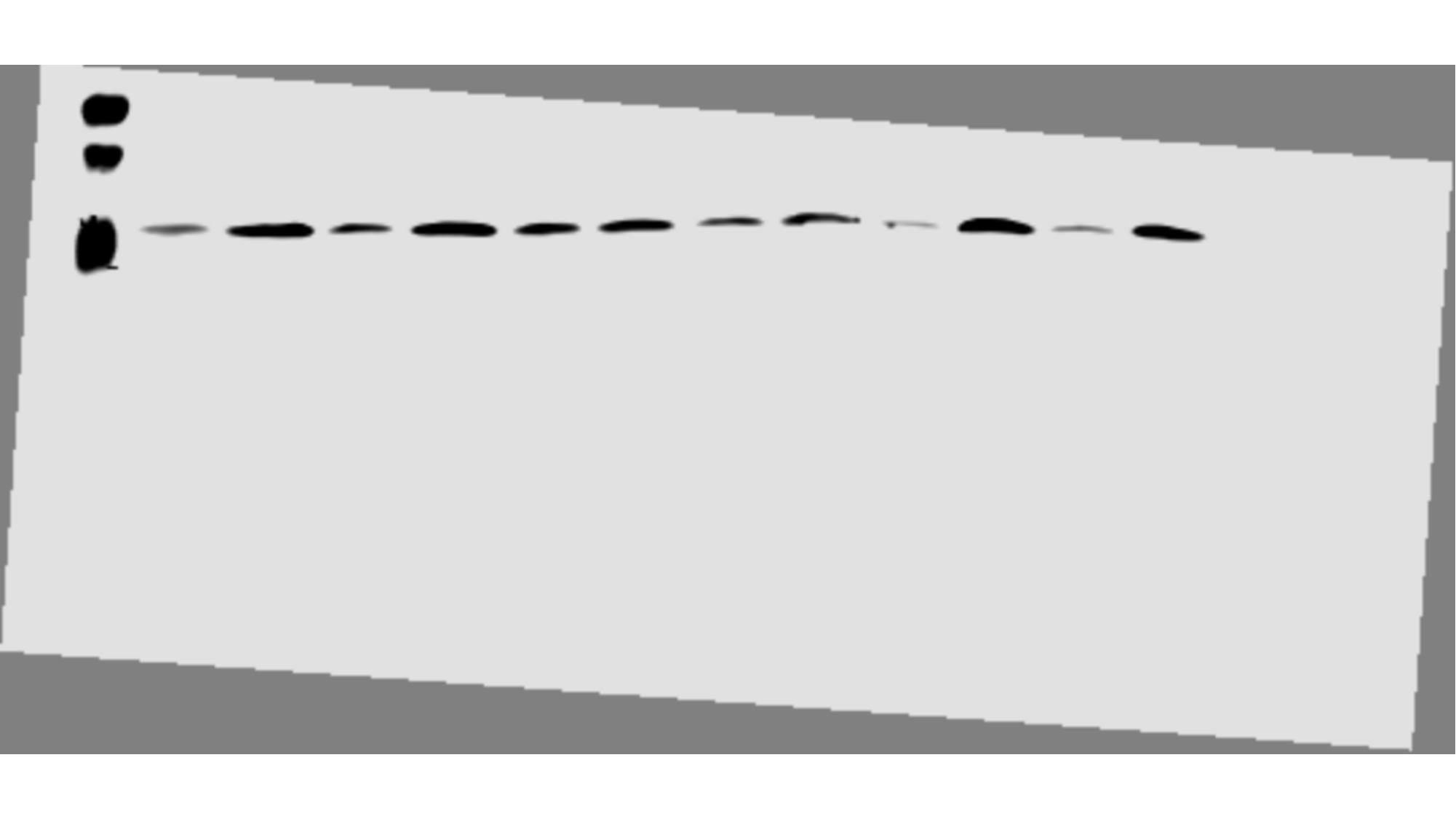

### Slide 10
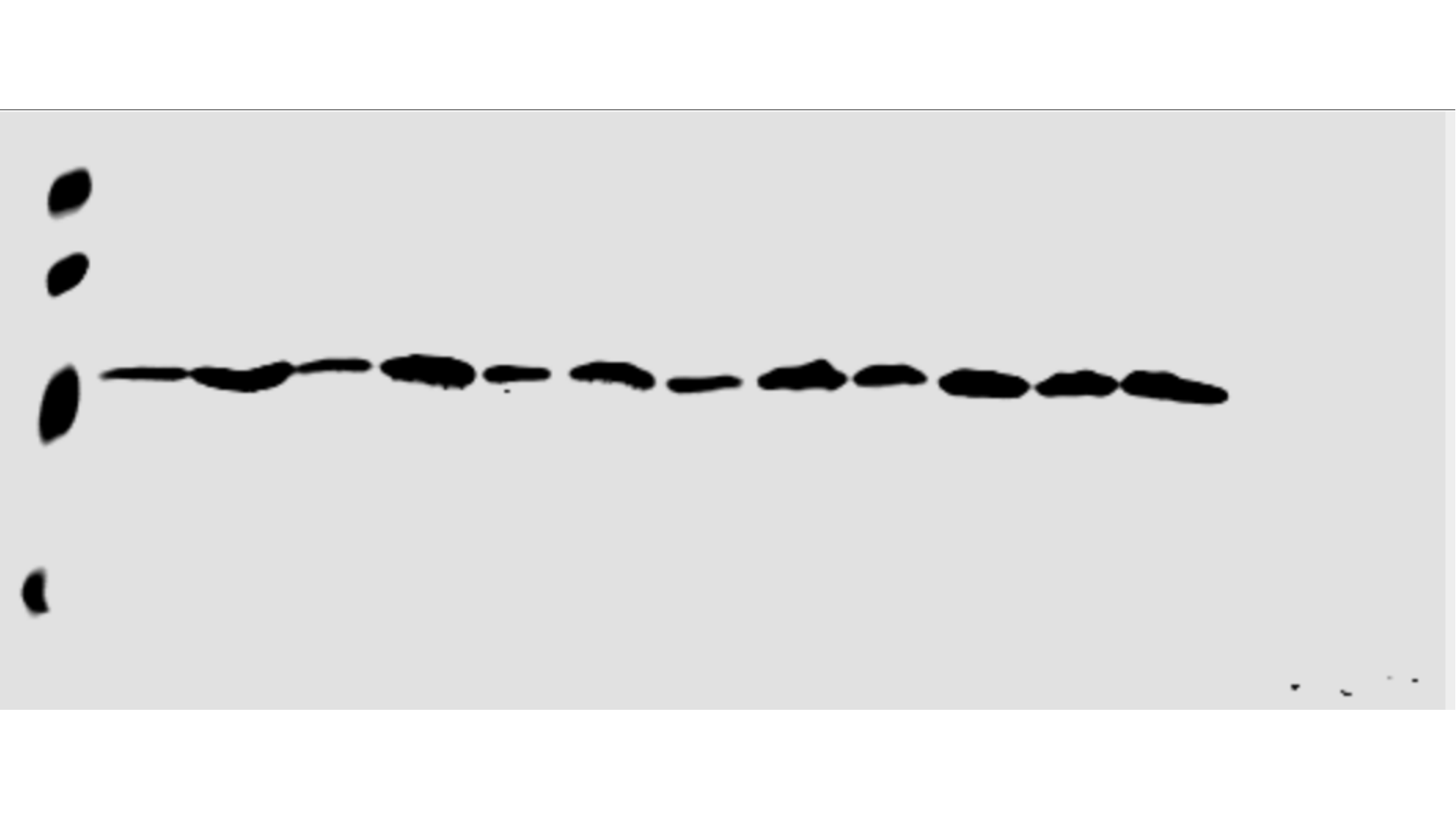
