## Supplementary material for "BRD4 Prevents R-Loop Formation and Transcription-Replication Conflicts by Ensuring Efficient Transcription Elongation": Source Data: 20200330_dex_camp.pptx

### Slide 1
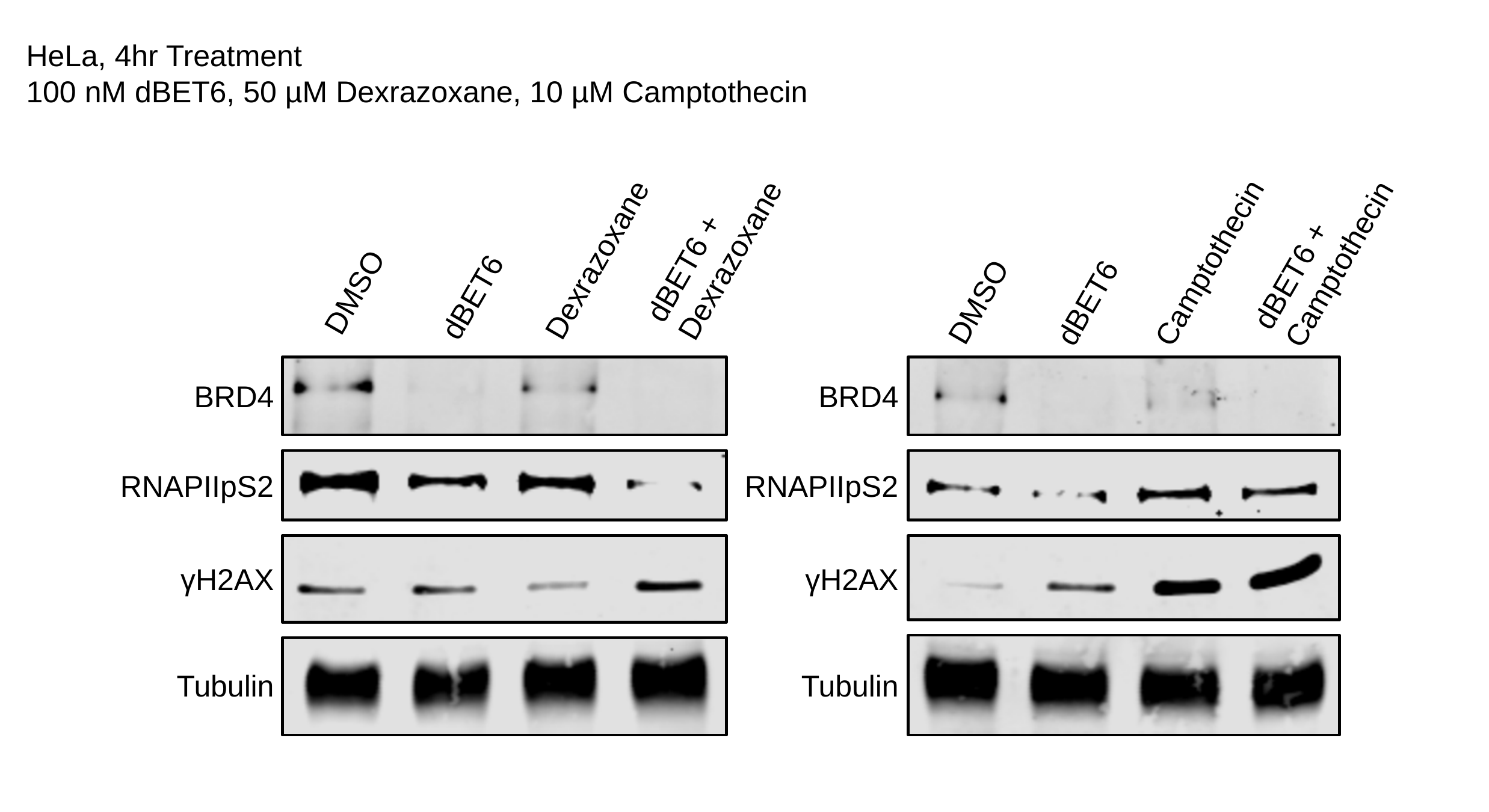

HeLa, 4hr Treatment
100 nM dBET6, 50 µM Dexrazoxane, 10 µM Camptothecin
dBET6 +
Dexrazoxane
dBET6 +
Camptothecin
Dexrazoxane
Camptothecin
DMSO
dBET6
DMSO
dBET6
BRD4
BRD4
RNAPIIpS2
RNAPIIpS2
γH2AX
γH2AX
Tubulin
Tubulin

### Slide 2
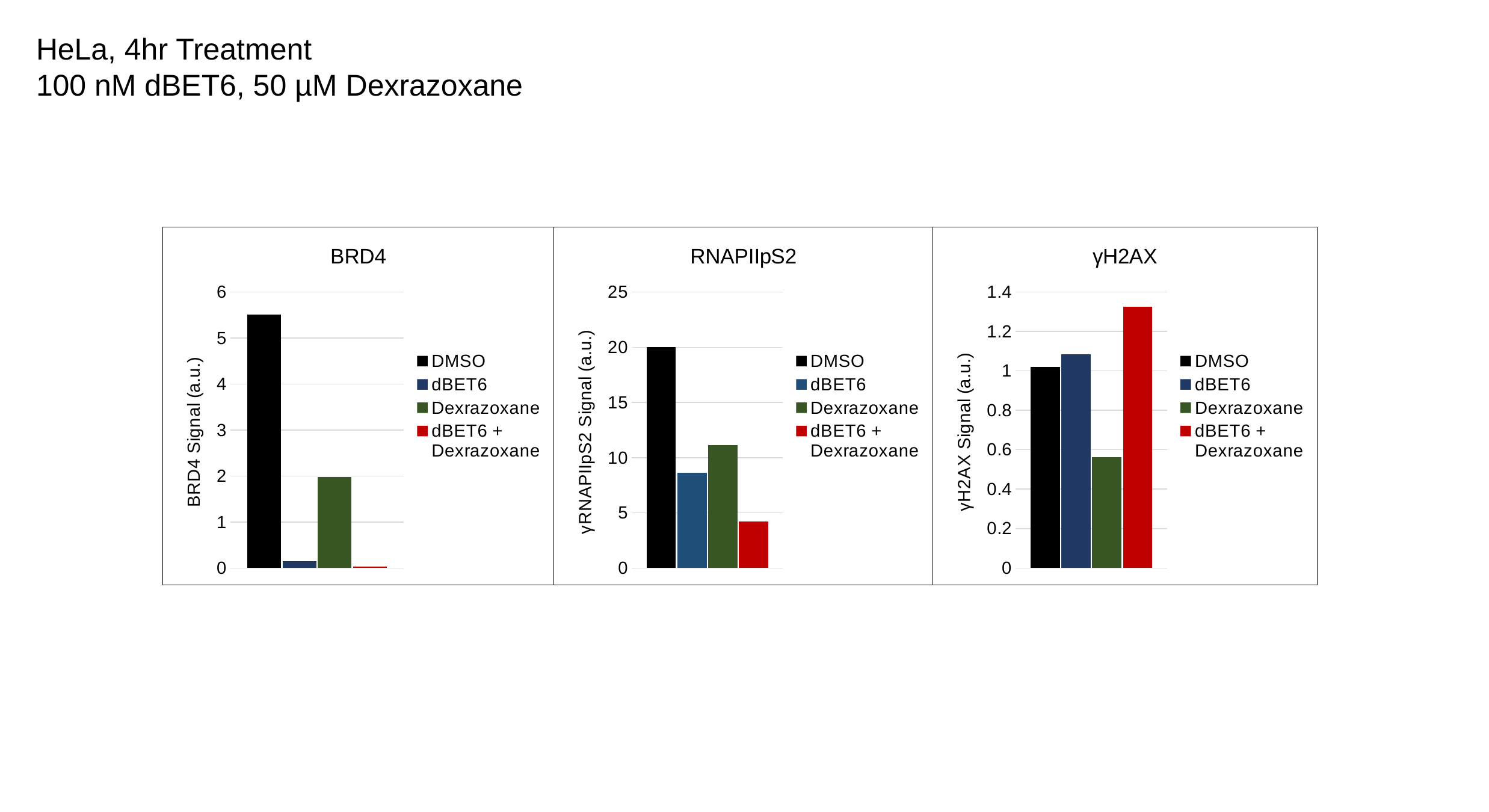

HeLa, 4hr Treatment
100 nM dBET6, 50 µM Dexrazoxane
#### Chart: RNAPIIpS2
| Category | | | | |
|---|---|---|---|---|
| RNAPIIpS2 | 20.03454231433506 | 8.62669245647969 | 11.15 | 4.217252396166135 |
#### Chart: γH2AX
| Category | | | | |
|---|---|---|---|---|
| γH2AX | 1.0210280373831777 | 1.0829145728643217 | 0.5622641509433962 | 1.3249475890985325 |
#### Chart: BRD4
| Category | | | | |
|---|---|---|---|---|
| BRD4 | 5.509499136442142 | 0.14777562862669244 | 1.9833333333333334 | 0.03578274760383387 |

### Slide 3
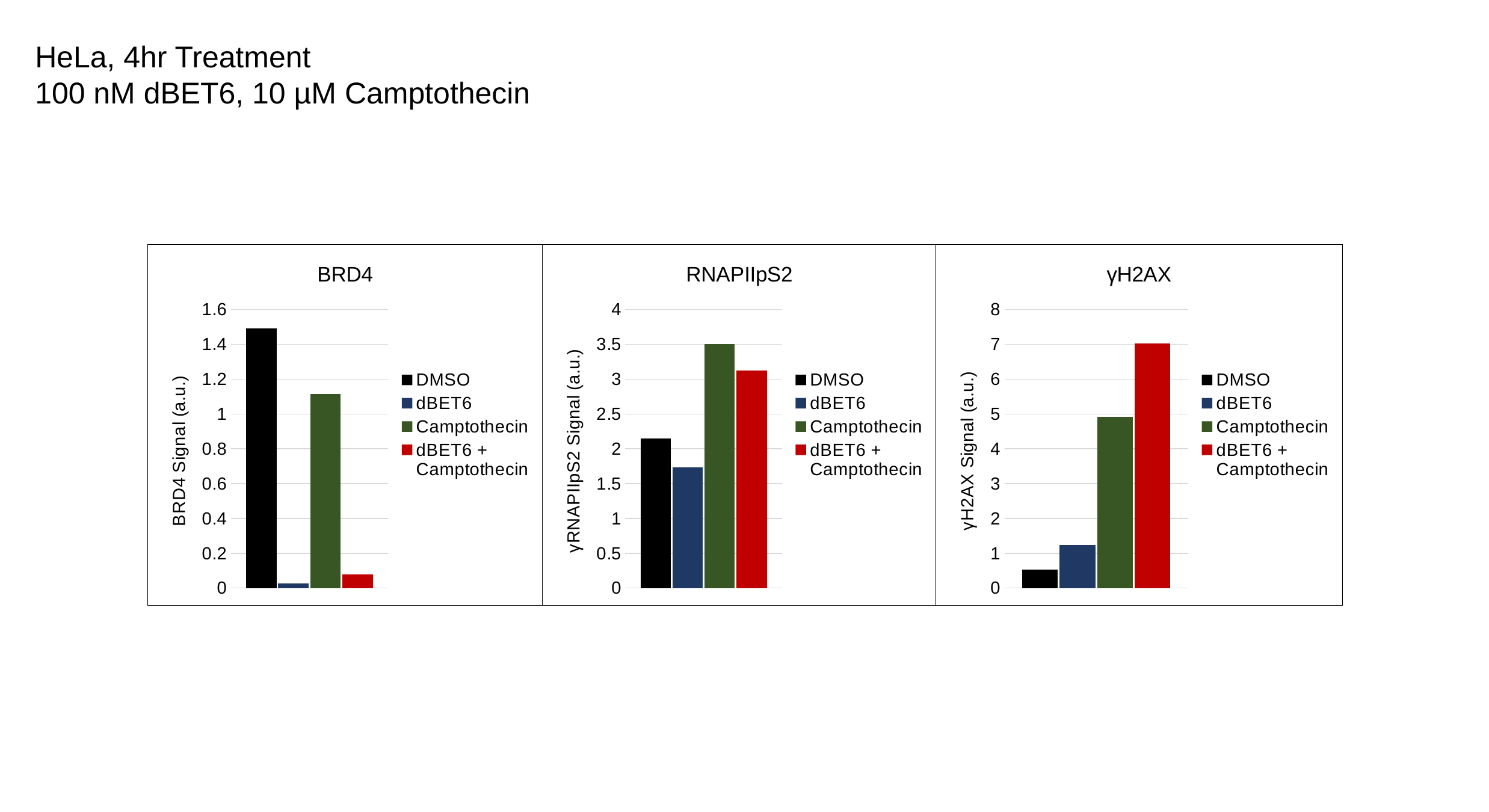

HeLa, 4hr Treatment
100 nM dBET6, 10 µM Camptothecin
#### Chart: BRD4
| Category | | | | |
|---|---|---|---|---|
| BRD4 | 1.4907651715039578 | 0.027044025157232705 | 1.1168384879725086 | 0.07678300455235204 |
#### Chart: RNAPIIpS2
| Category | | | | |
|---|---|---|---|---|
| RNAPIIpS2 | 2.150395778364116 | 1.729559748427673 | 3.5051546391752577 | 3.125948406676783 |
#### Chart: γH2AX
| Category | | | | |
|---|---|---|---|---|
| γH2AX | 0.528604118993135 | 1.2402088772845952 | 4.920212765957447 | 7.0212765957446805 |

### Slide 4
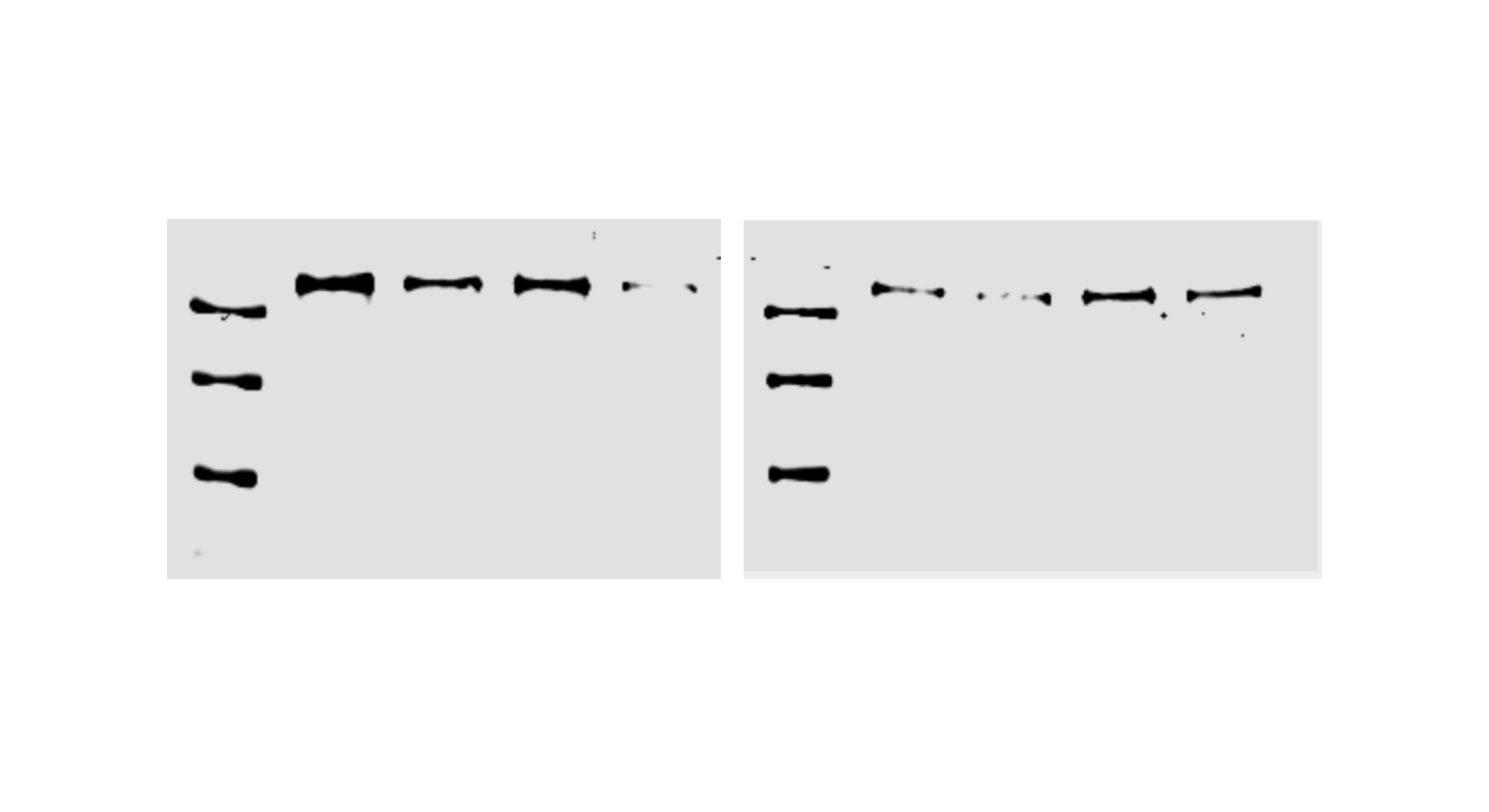

### Slide 5
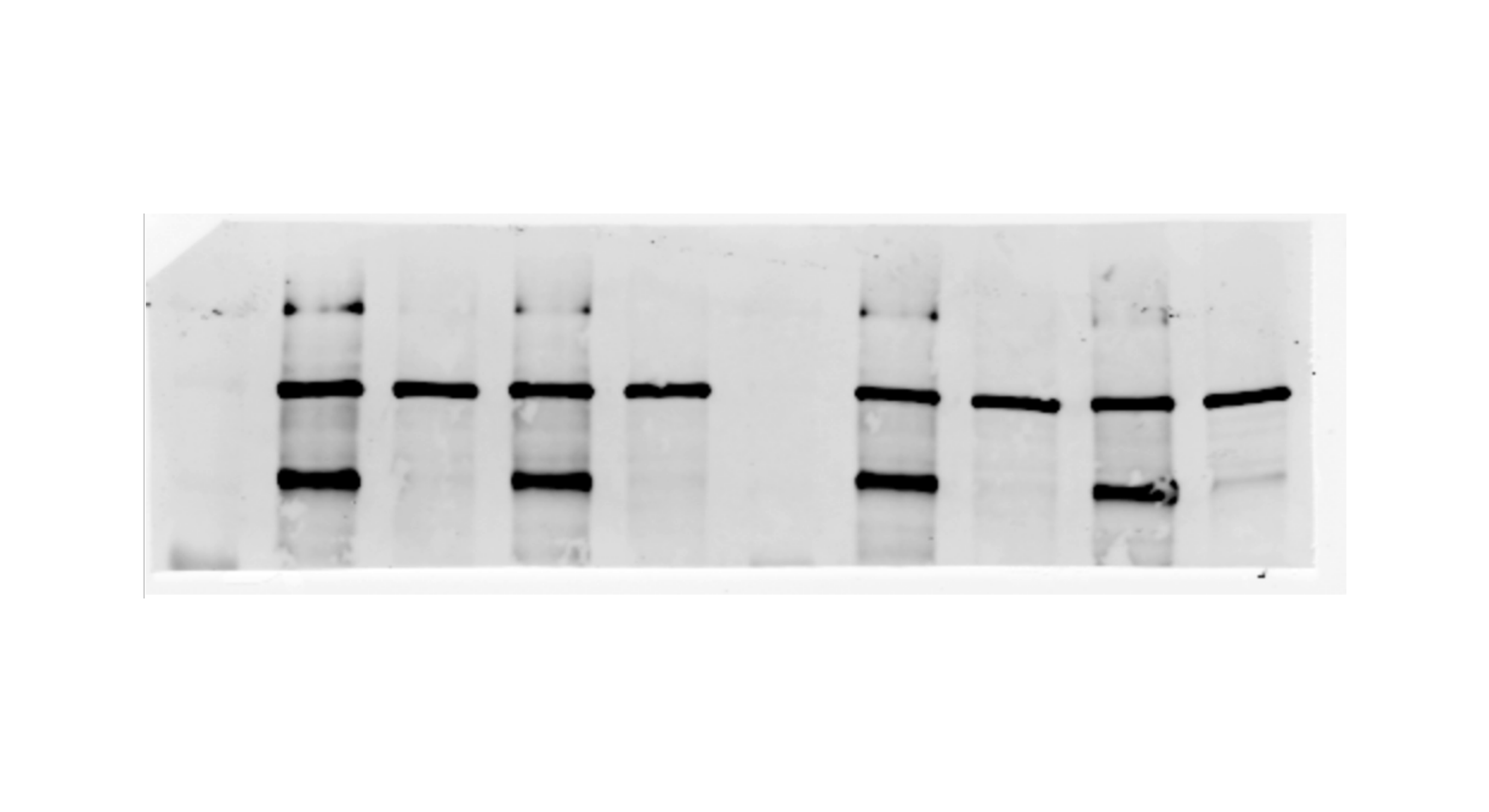

### Slide 6
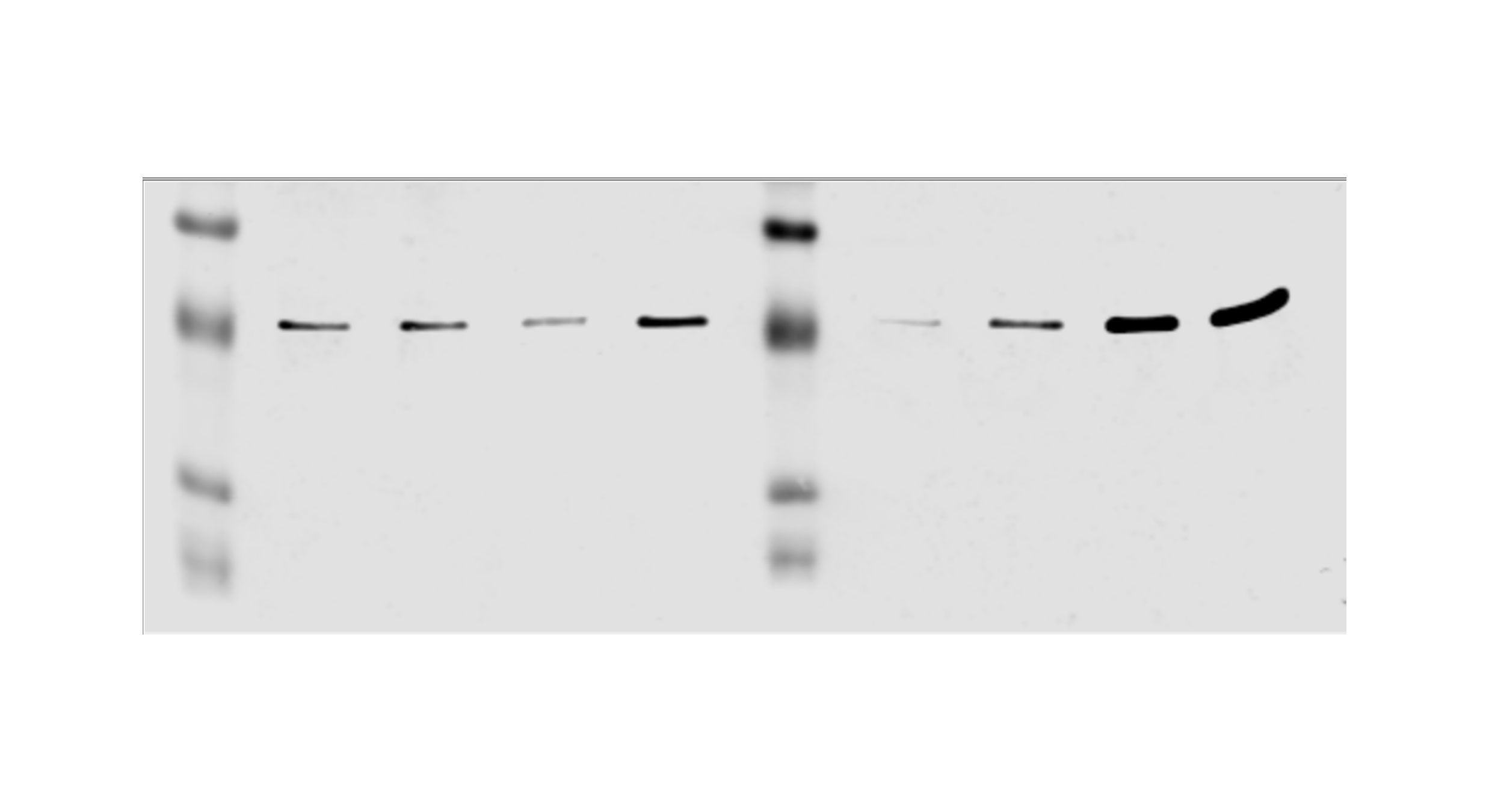

### Slide 7
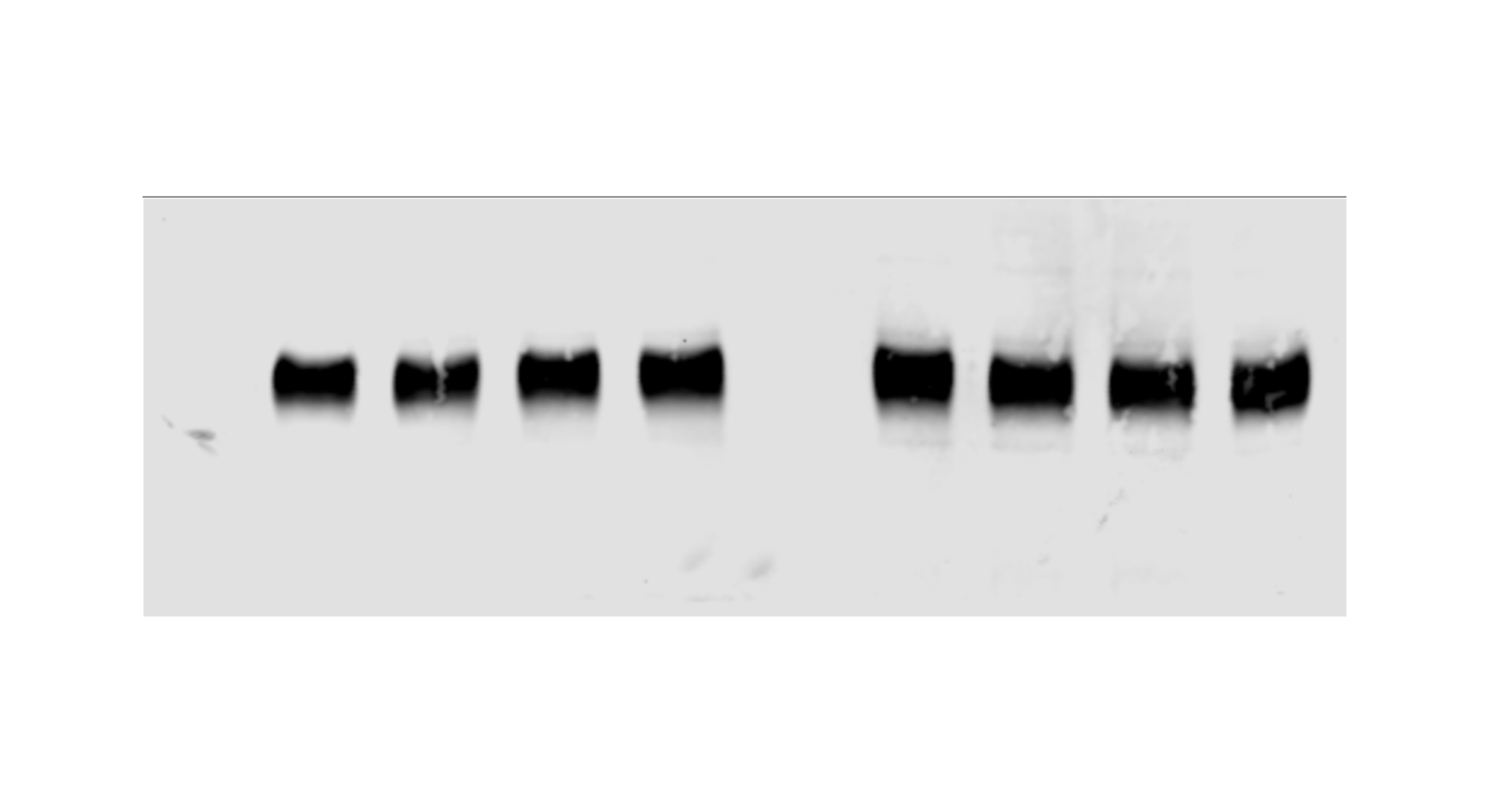

### Slide 8
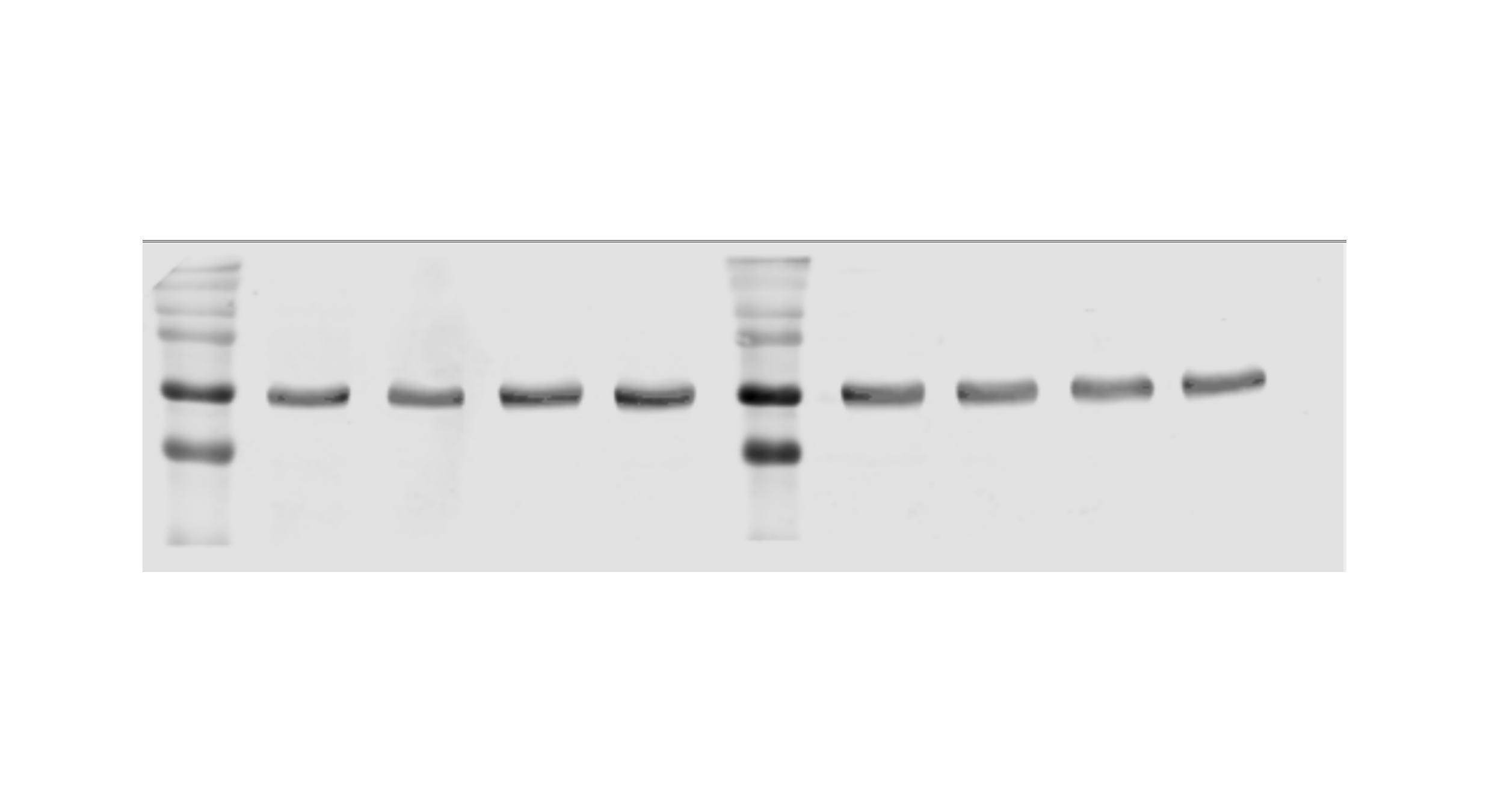
