## Supplementary material for "BRD4 Prevents R-Loop Formation and Transcription-Replication Conflicts by Ensuring Efficient Transcription Elongation": Source Data: DRB- TRP Dosage Titration.pptx

### Slide 1
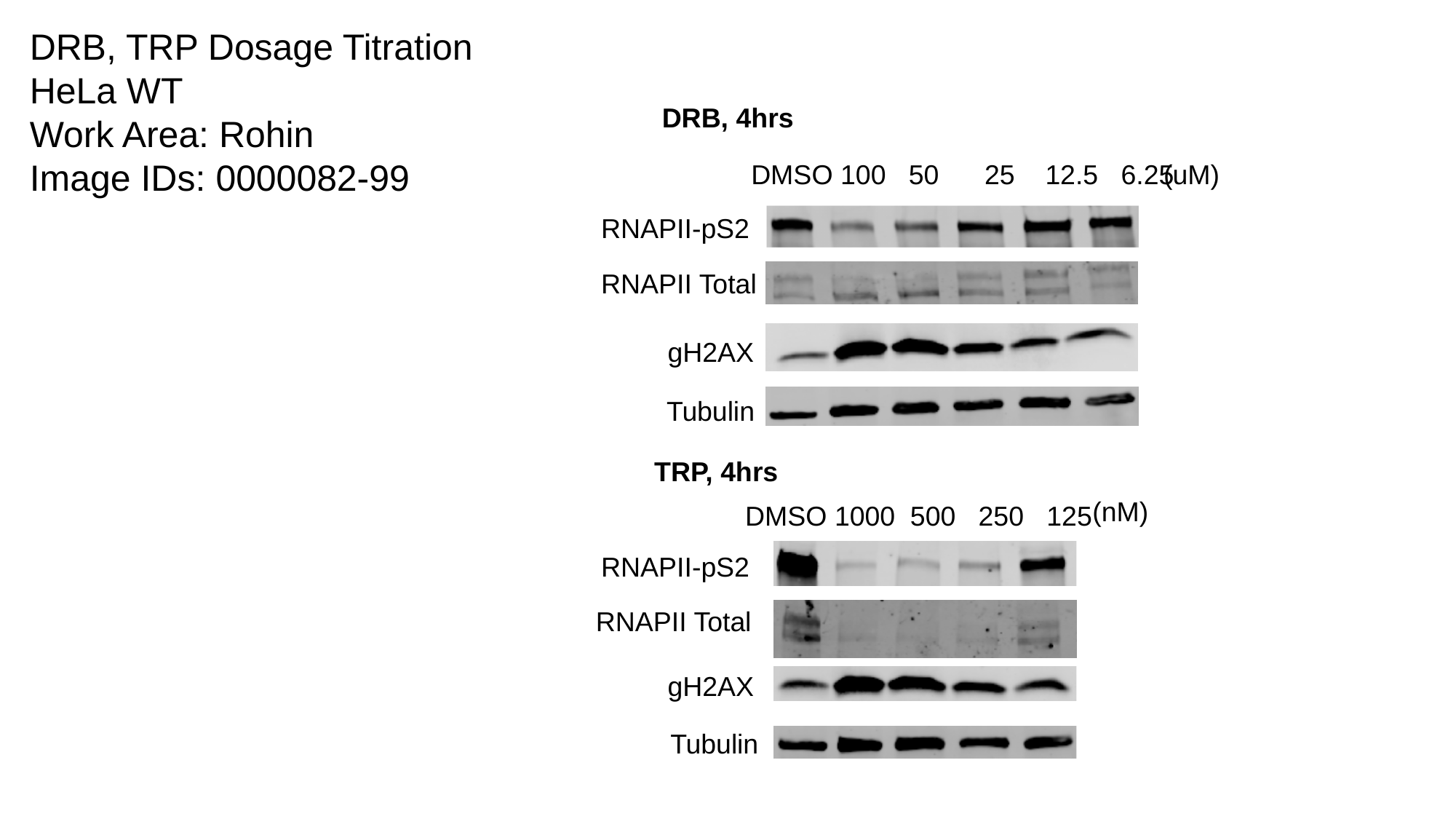

DRB, TRP Dosage Titration
HeLa WT
Work Area: Rohin
Image IDs: 0000082-99
DRB, 4hrs
(uM)
DMSO 100 50 25 12.5 6.25
RNAPII-pS2
RNAPII Total
gH2AX
Tubulin
TRP, 4hrs
(nM)
DMSO 1000 500 250 125
RNAPII-pS2
RNAPII Total
gH2AX
Tubulin

### Slide 2
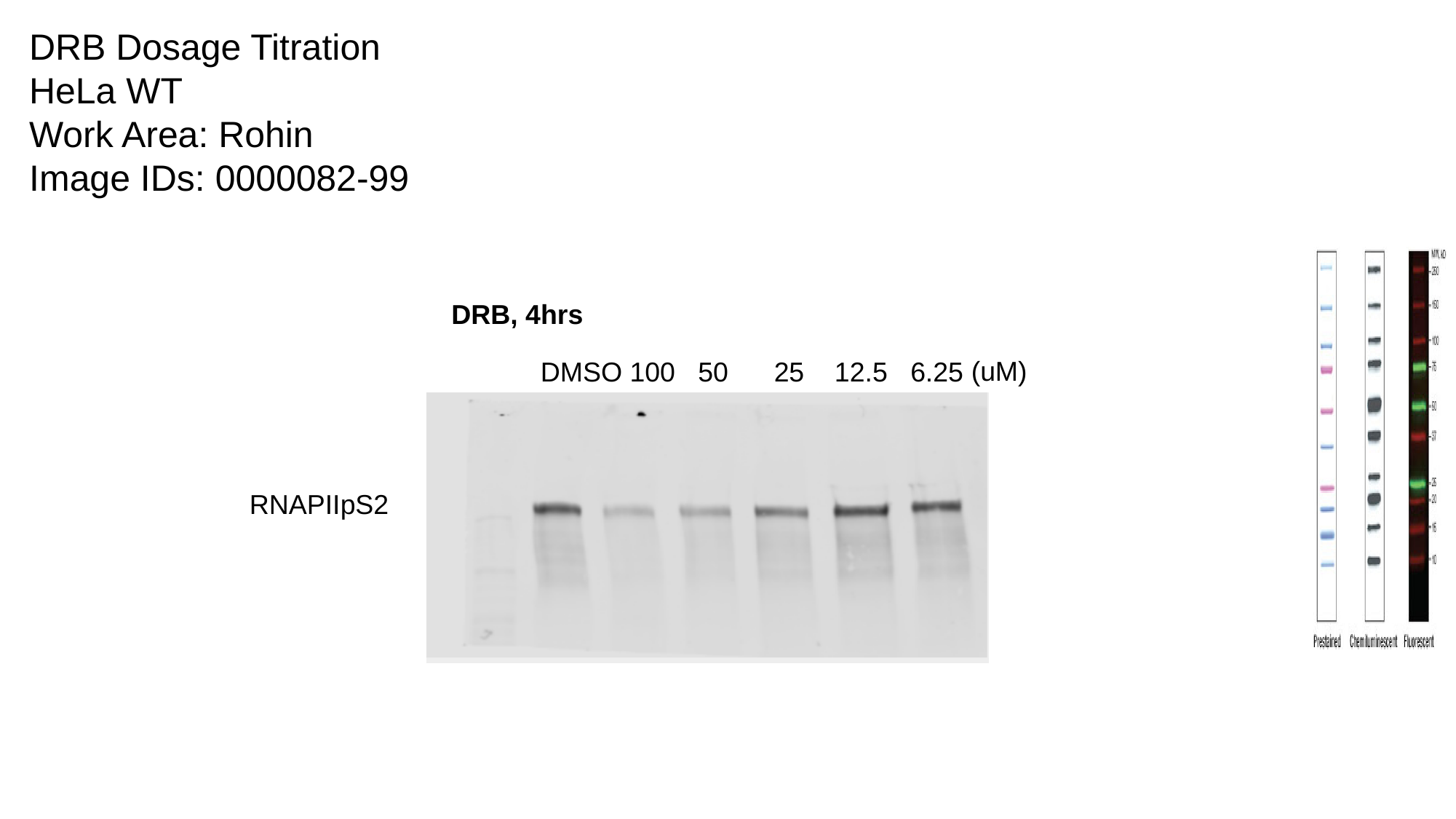

DRB Dosage Titration
HeLa WT
Work Area: Rohin
Image IDs: 0000082-99
DRB, 4hrs
(uM)
DMSO 100 50 25 12.5 6.25
RNAPIIpS2

### Slide 3
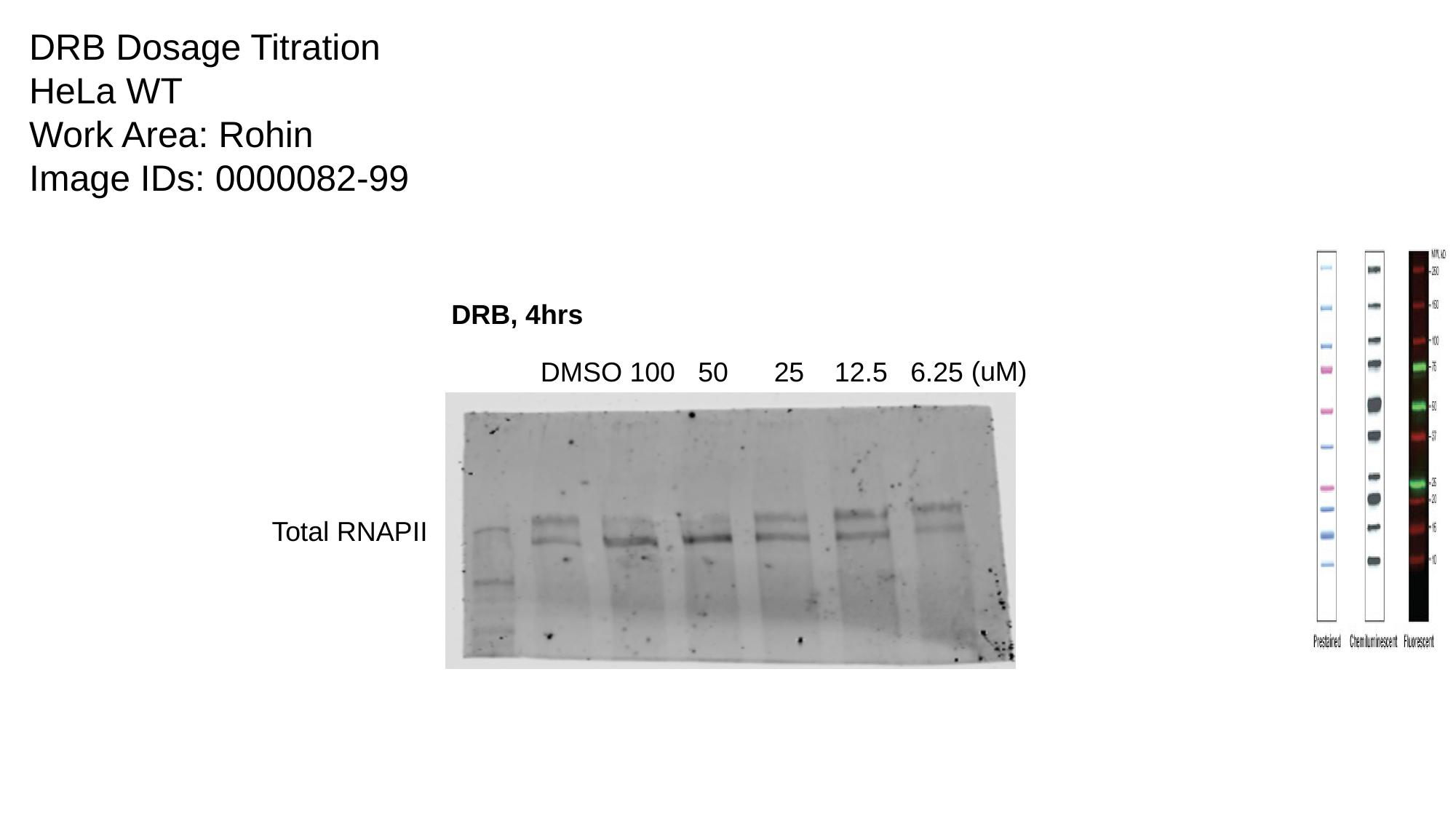

DRB Dosage Titration
HeLa WT
Work Area: Rohin
Image IDs: 0000082-99
DRB, 4hrs
(uM)
DMSO 100 50 25 12.5 6.25
Total RNAPII

### Slide 4
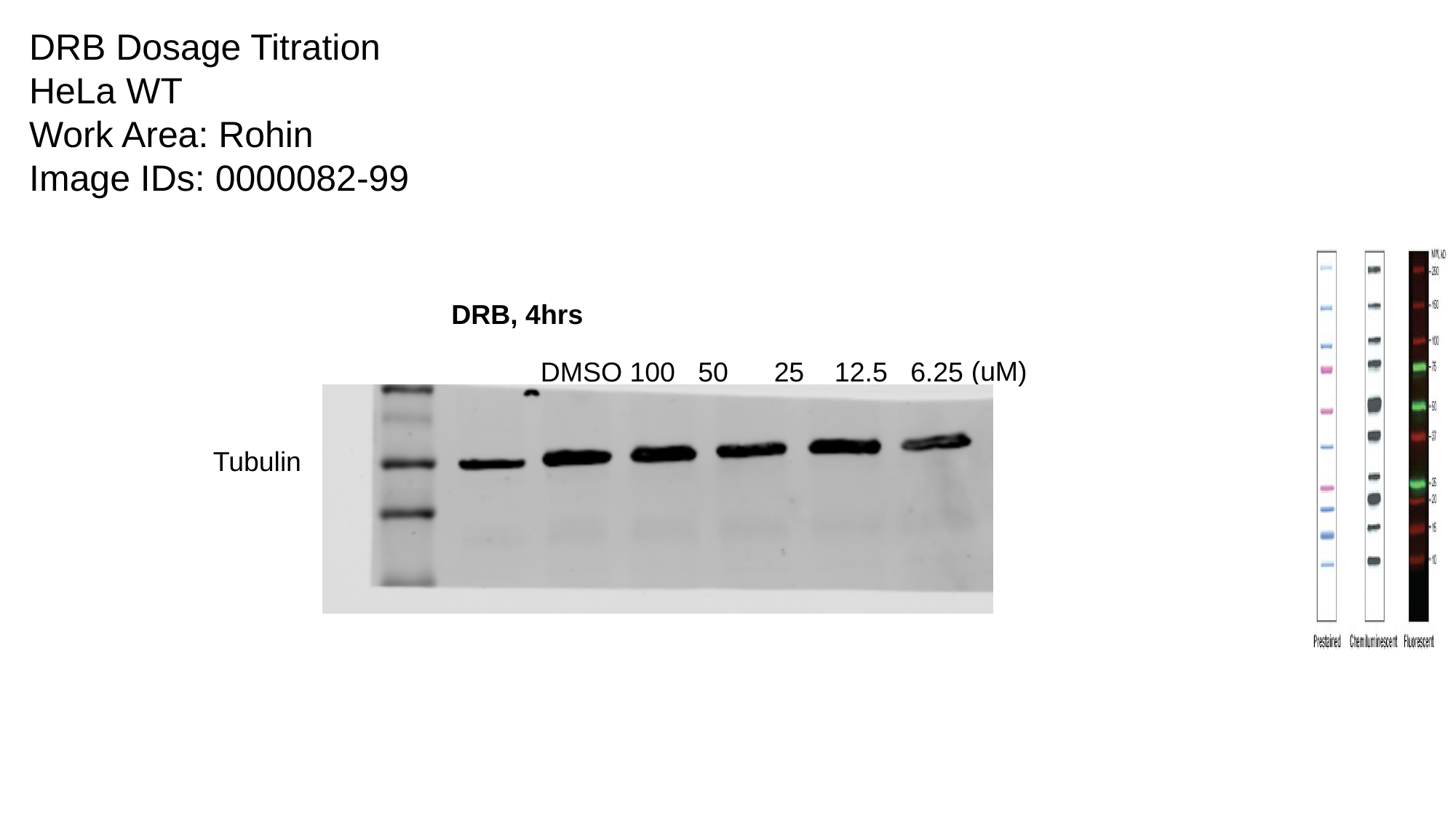

DRB Dosage Titration
HeLa WT
Work Area: Rohin
Image IDs: 0000082-99
DRB, 4hrs
(uM)
DMSO 100 50 25 12.5 6.25
Tubulin

### Slide 5
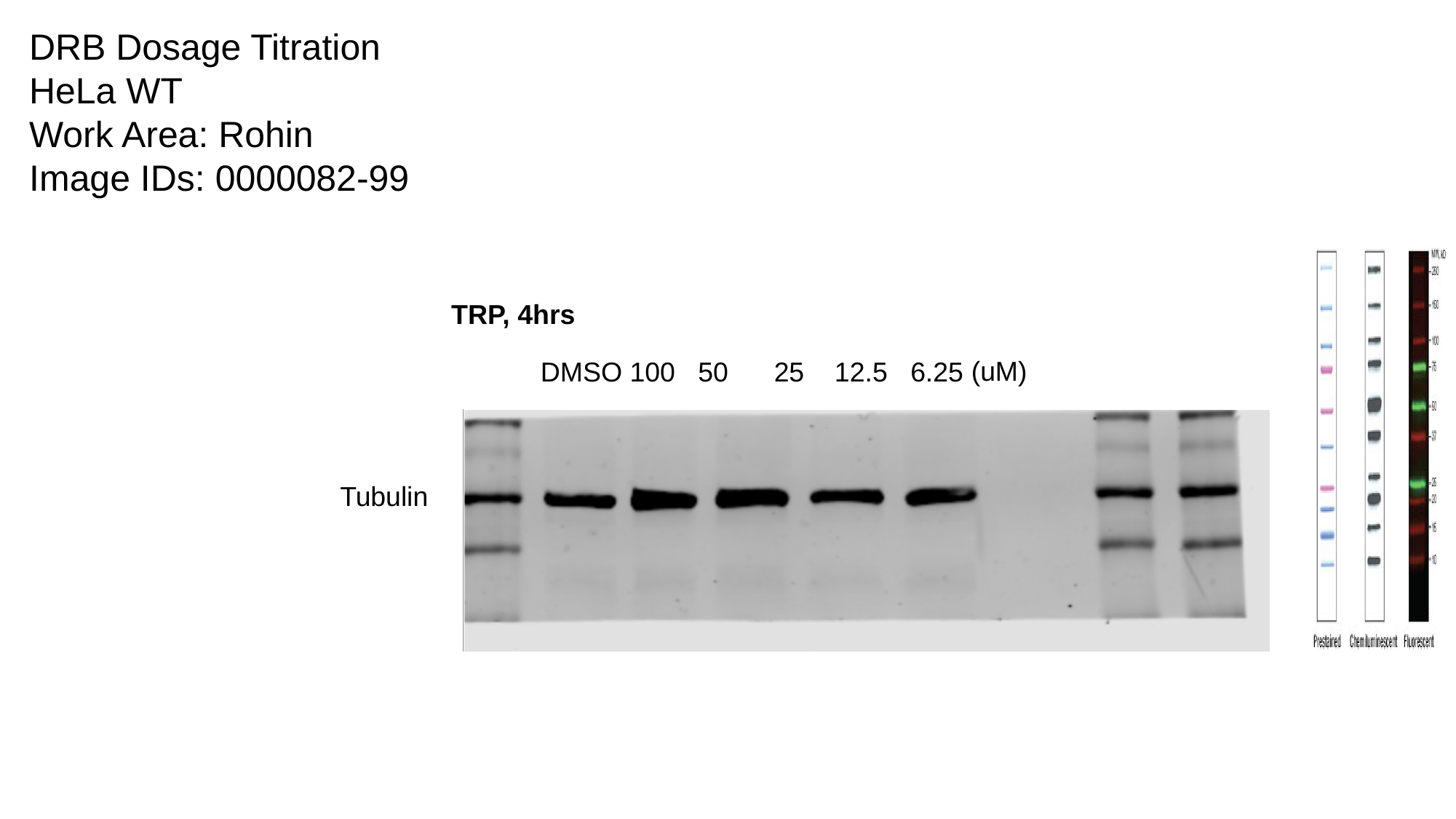

DRB Dosage Titration
HeLa WT
Work Area: Rohin
Image IDs: 0000082-99
TRP, 4hrs
(uM)
DMSO 100 50 25 12.5 6.25
Tubulin

### Slide 6
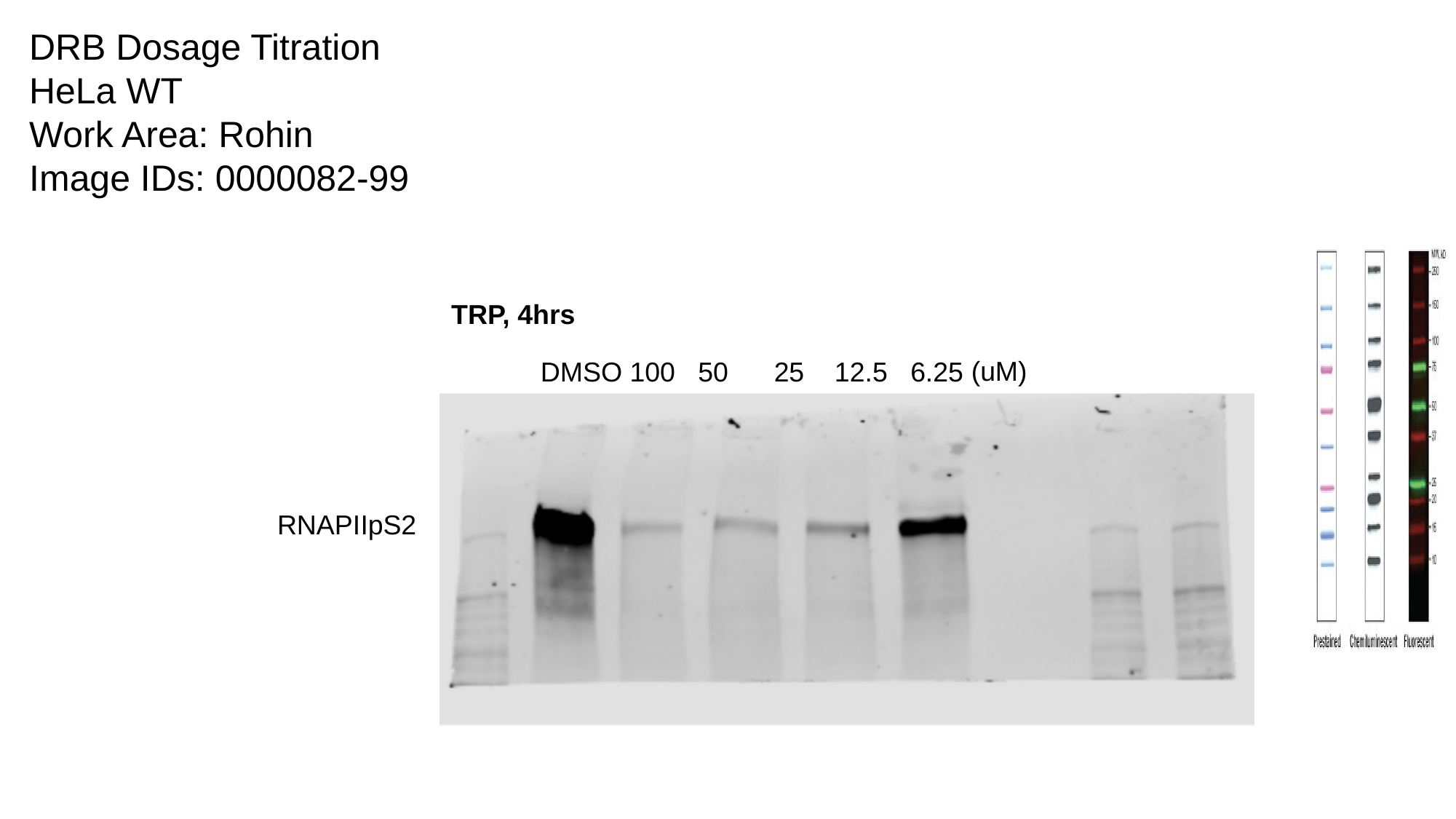

DRB Dosage Titration
HeLa WT
Work Area: Rohin
Image IDs: 0000082-99
TRP, 4hrs
(uM)
DMSO 100 50 25 12.5 6.25
RNAPIIpS2

### Slide 7
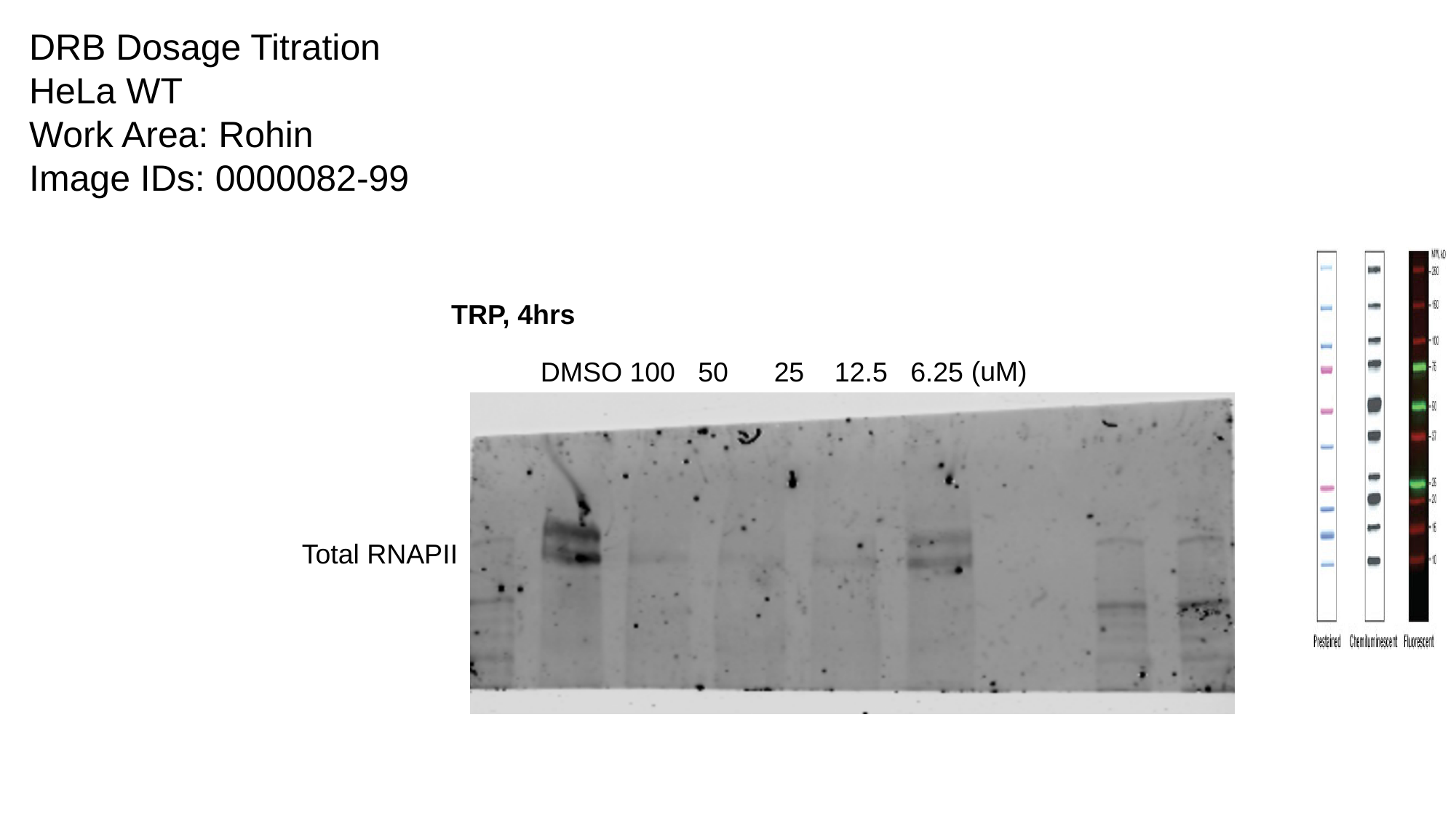

DRB Dosage Titration
HeLa WT
Work Area: Rohin
Image IDs: 0000082-99
TRP, 4hrs
(uM)
DMSO 100 50 25 12.5 6.25
Total RNAPII
