## Supplementary material for "BRD4 Prevents R-Loop Formation and Transcription-Replication Conflicts by Ensuring Efficient Transcription Elongation": Source Data: Trp-dBET6 Rescue Triplicate.pptx

### Slide 1
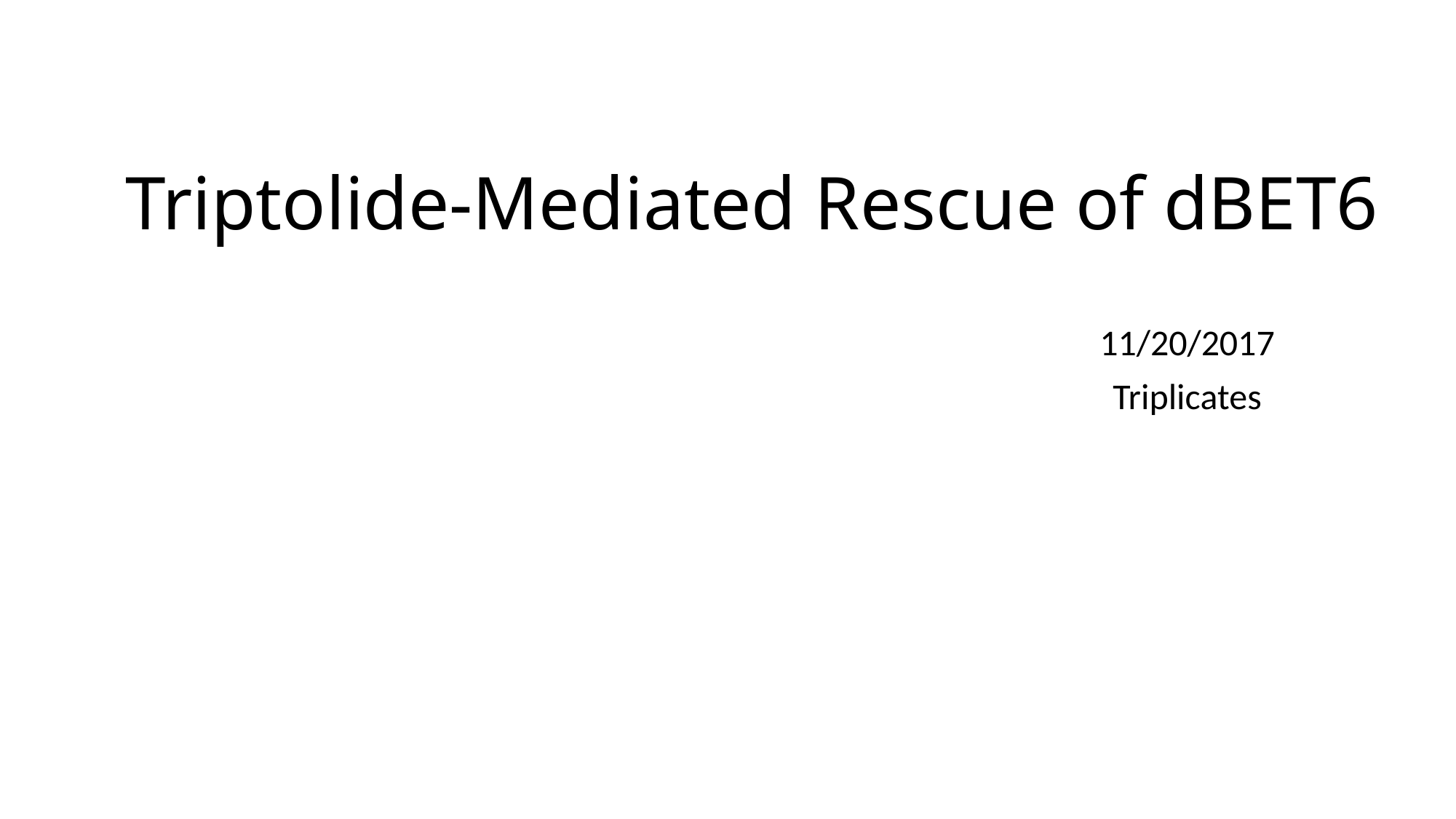

#### Triptolide-Mediated Rescue of dBET6
11/20/2017
Triplicates

### Slide 2
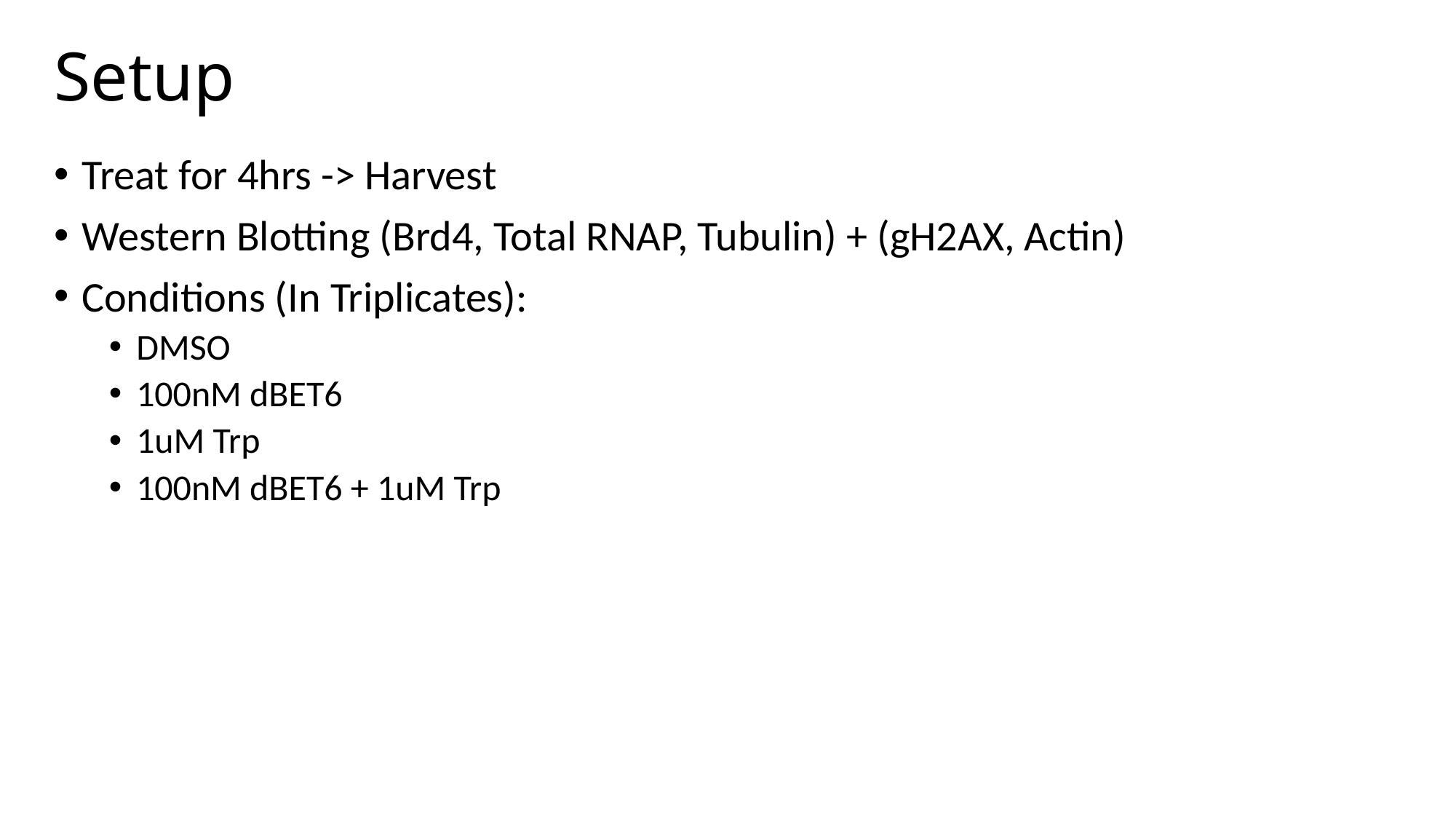

#### Setup
Treat for 4hrs -> Harvest
Western Blotting (Brd4, Total RNAP, Tubulin) + (gH2AX, Actin)
Conditions (In Triplicates):
DMSO
100nM dBET6
1uM Trp
100nM dBET6 + 1uM Trp

### Slide 3
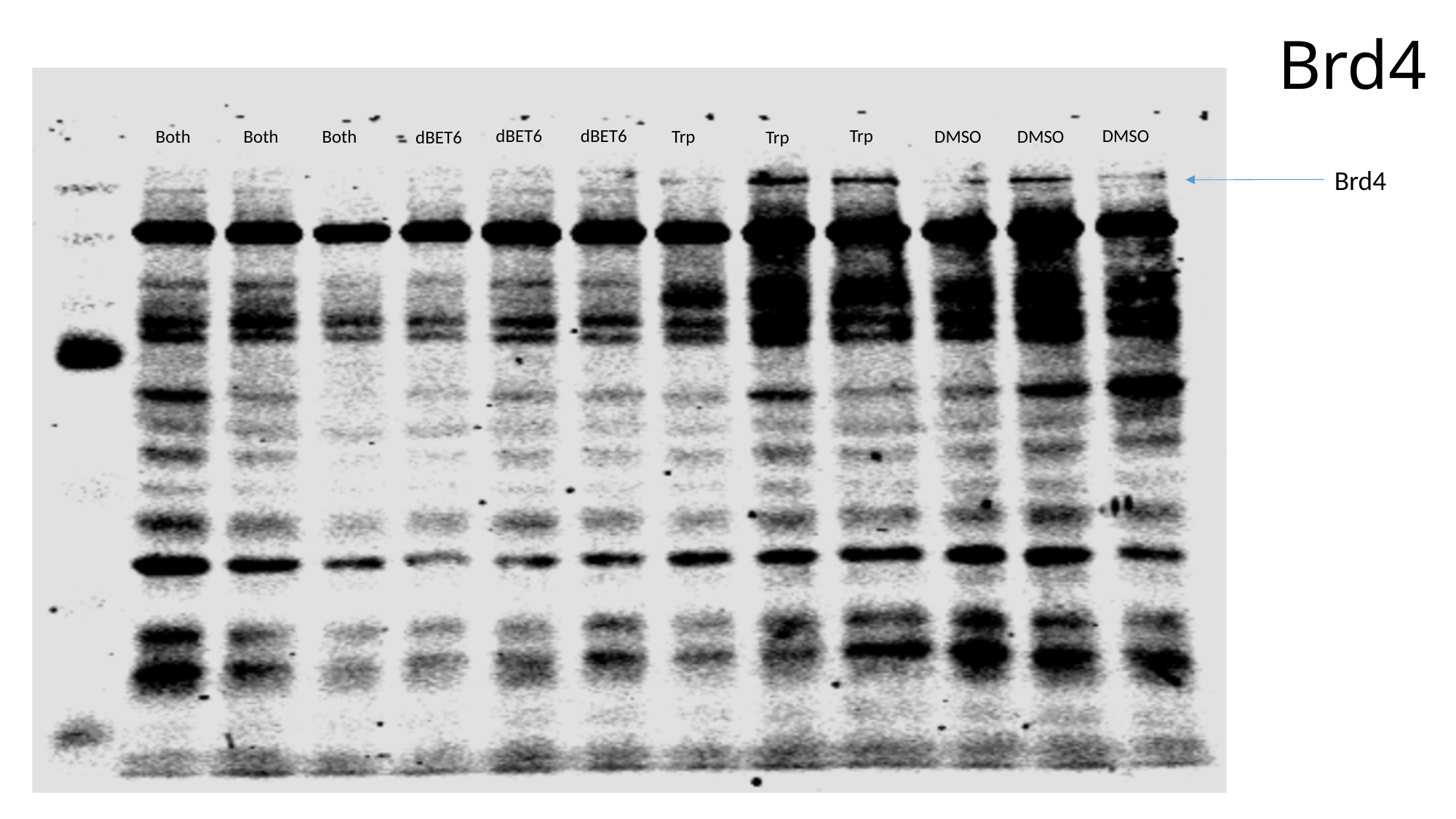

#### Brd4
dBET6
dBET6
Trp
DMSO
Both
DMSO
Both
Both
Trp
DMSO
dBET6
Trp
Brd4

### Slide 4
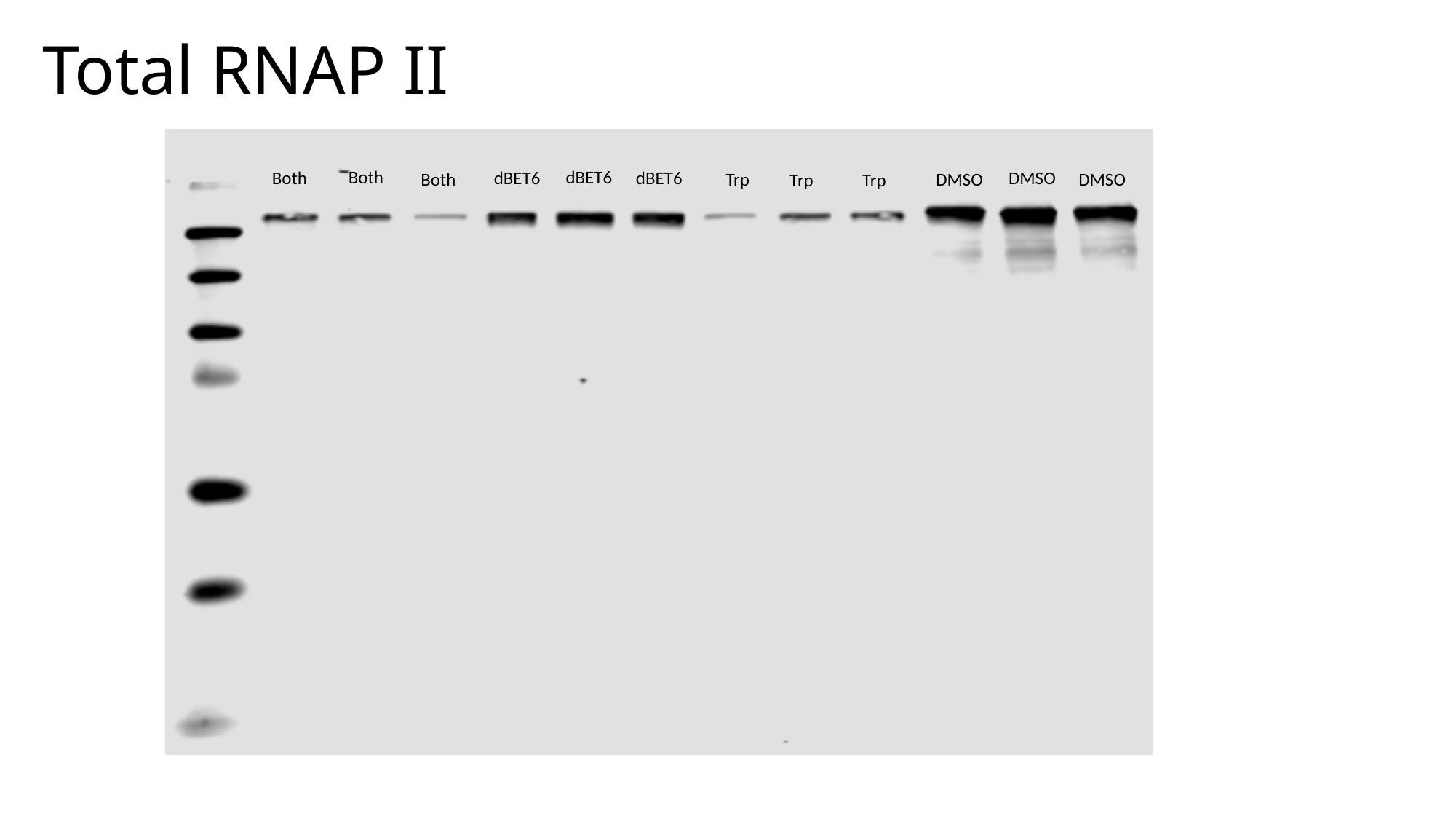

#### Total RNAP II
Both
dBET6
Both
dBET6
dBET6
DMSO
Both
Trp
DMSO
DMSO
Trp
Trp

### Slide 5
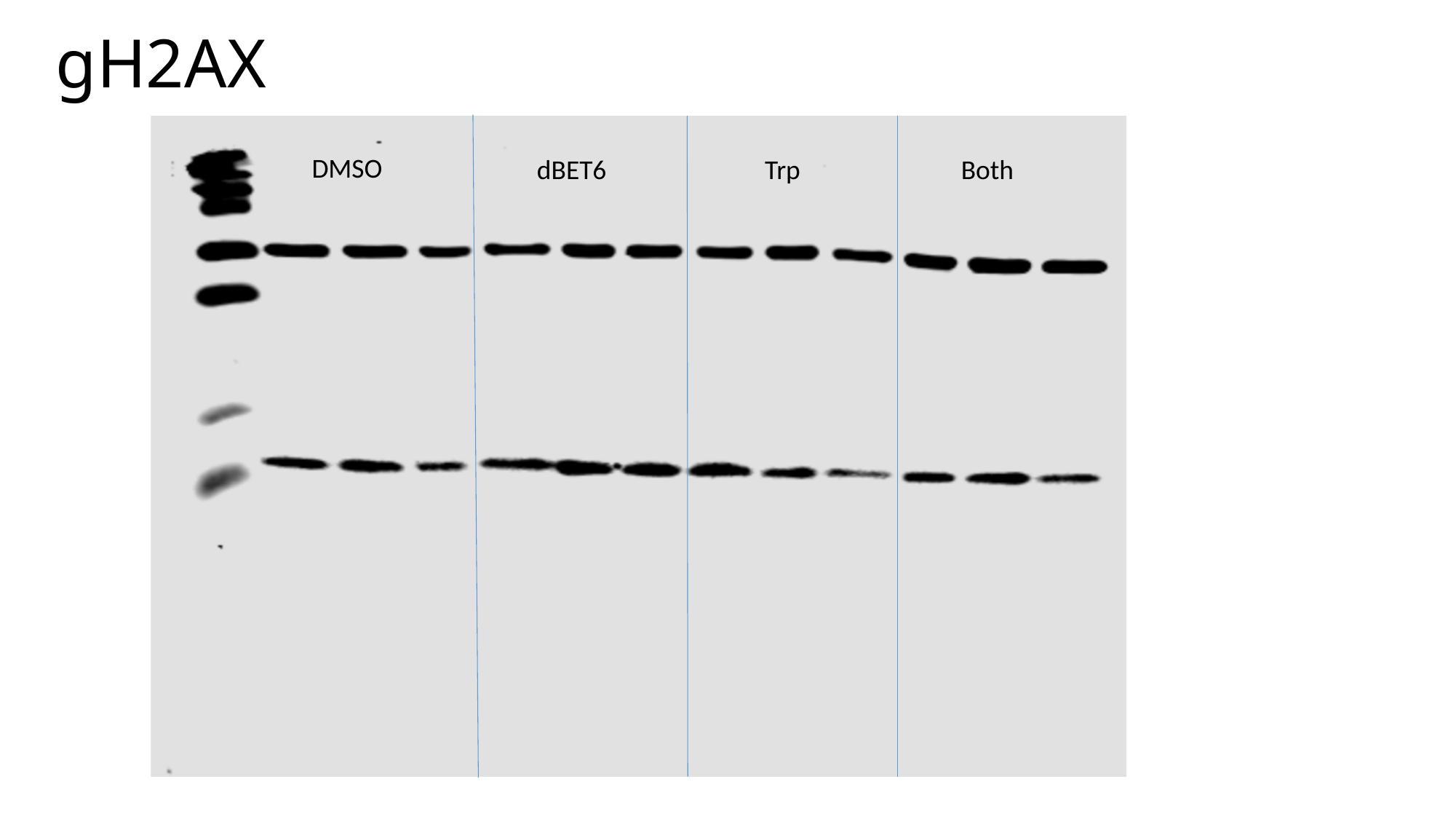

# gH2AX
DMSO
dBET6
Trp
Both

### Slide 6
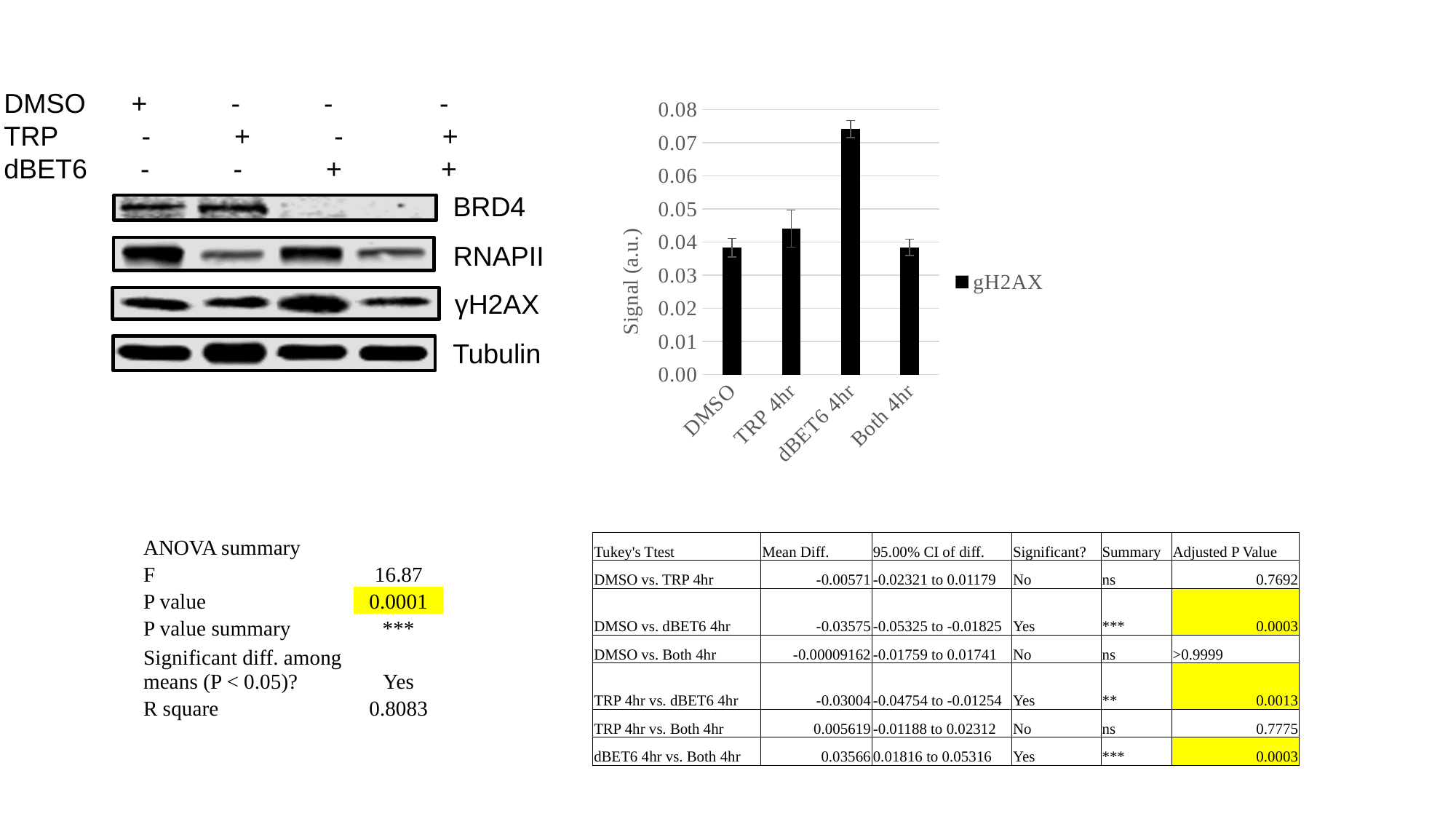

DMSO + - - -
TRP - + - +
dBET6 - - + +
###### Chart
| Category | |
|---|---|
| DMSO | 0.038310128344974305 |
| TRP 4hr | 0.044020342848660715 |
| dBET6 4hr | 0.07406086069457445 |
| Both 4hr | 0.038401992366060456 |BRD4
RNAPII
γH2AX
Tubulin
| ANOVA summary | |
| --- | --- |
| F | 16.87 |
| P value | 0.0001 |
| P value summary | \*\*\* |
| Significant diff. among means (P < 0.05)? | Yes |
| R square | 0.8083 |
| Tukey's Ttest | Mean Diff. | 95.00% CI of diff. | Significant? | Summary | Adjusted P Value |
| --- | --- | --- | --- | --- | --- |
| DMSO vs. TRP 4hr | -0.00571 | -0.02321 to 0.01179 | No | ns | 0.7692 |
| DMSO vs. dBET6 4hr | -0.03575 | -0.05325 to -0.01825 | Yes | \*\*\* | 0.0003 |
| DMSO vs. Both 4hr | -0.00009162 | -0.01759 to 0.01741 | No | ns | >0.9999 |
| TRP 4hr vs. dBET6 4hr | -0.03004 | -0.04754 to -0.01254 | Yes | \*\* | 0.0013 |
| TRP 4hr vs. Both 4hr | 0.005619 | -0.01188 to 0.02312 | No | ns | 0.7775 |
| dBET6 4hr vs. Both 4hr | 0.03566 | 0.01816 to 0.05316 | Yes | \*\*\* | 0.0003 |
