## Supplementary figures and images for "BRD4 Prevents R-Loop Formation and Transcription-Replication Conflicts by Ensuring Efficient Transcription Elongation"

### 20160525_MEF_KPR8.pptx

#### Slide 1
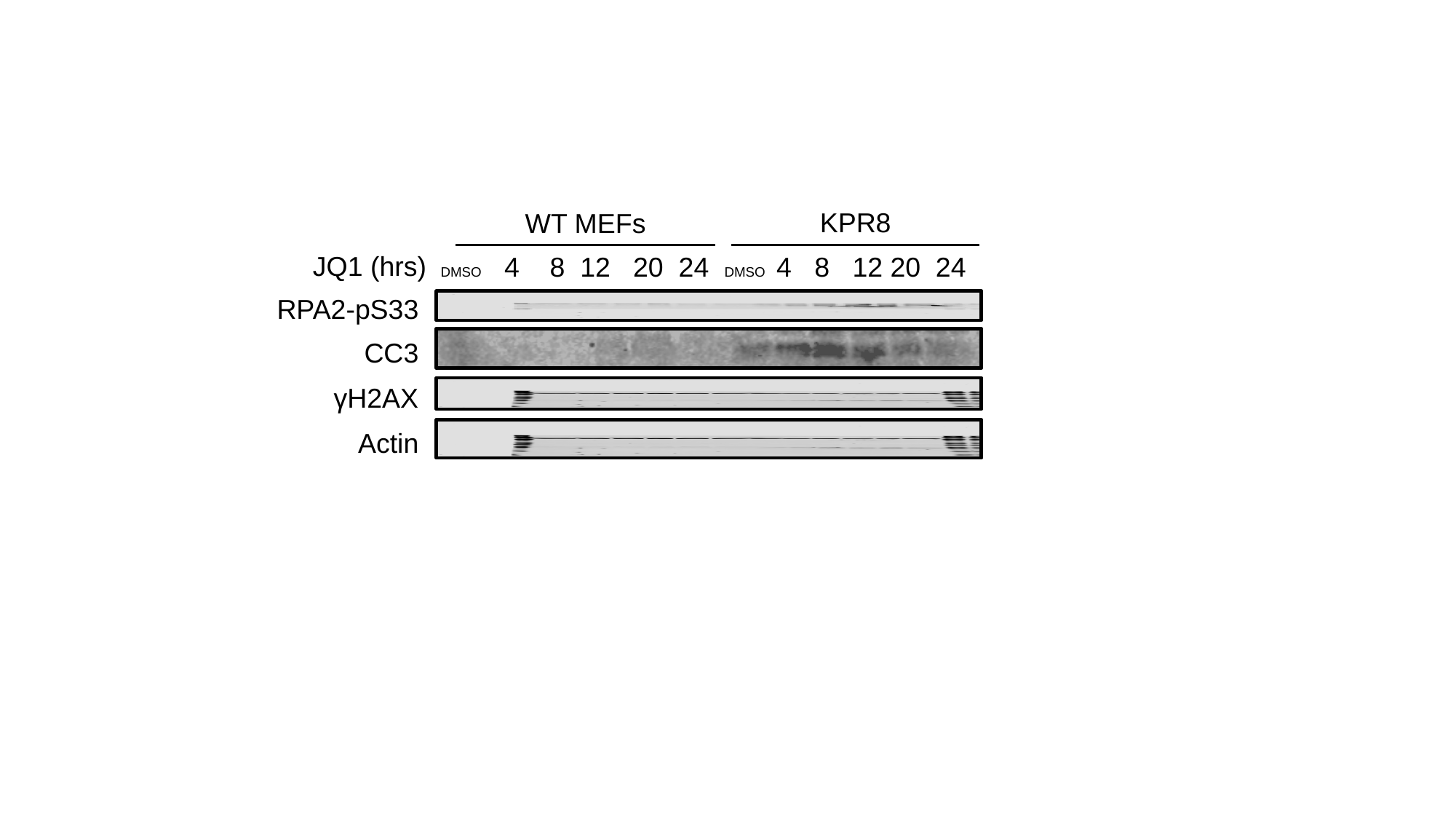

KPR8
WT MEFs
JQ1 (hrs)
DMSO 4 8 12 20 24 DMSO 4 8 12 20 24
RPA2-pS33
CC3
γH2AX
Actin

#### Slide 2
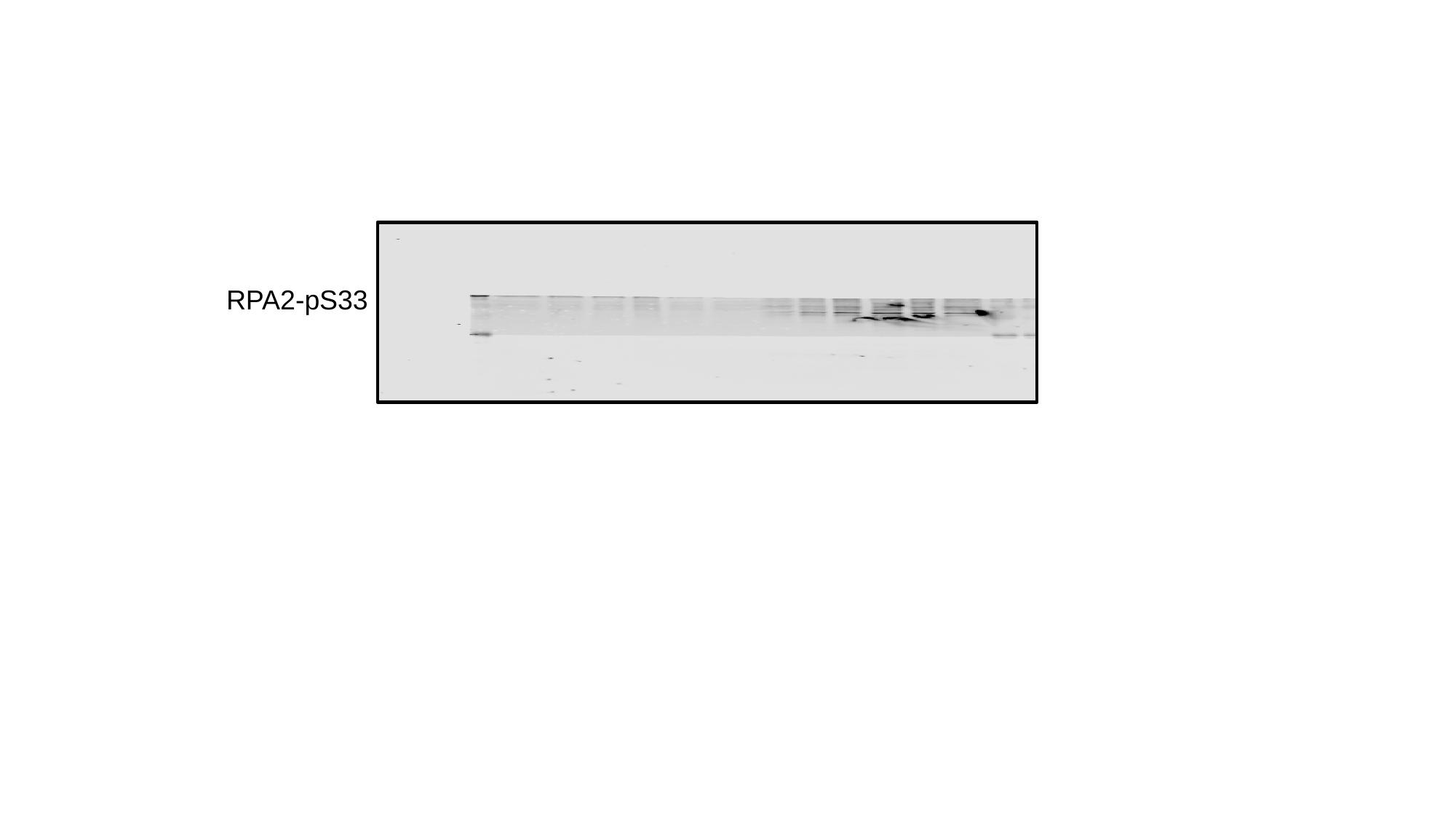

RPA2-pS33

#### Slide 3
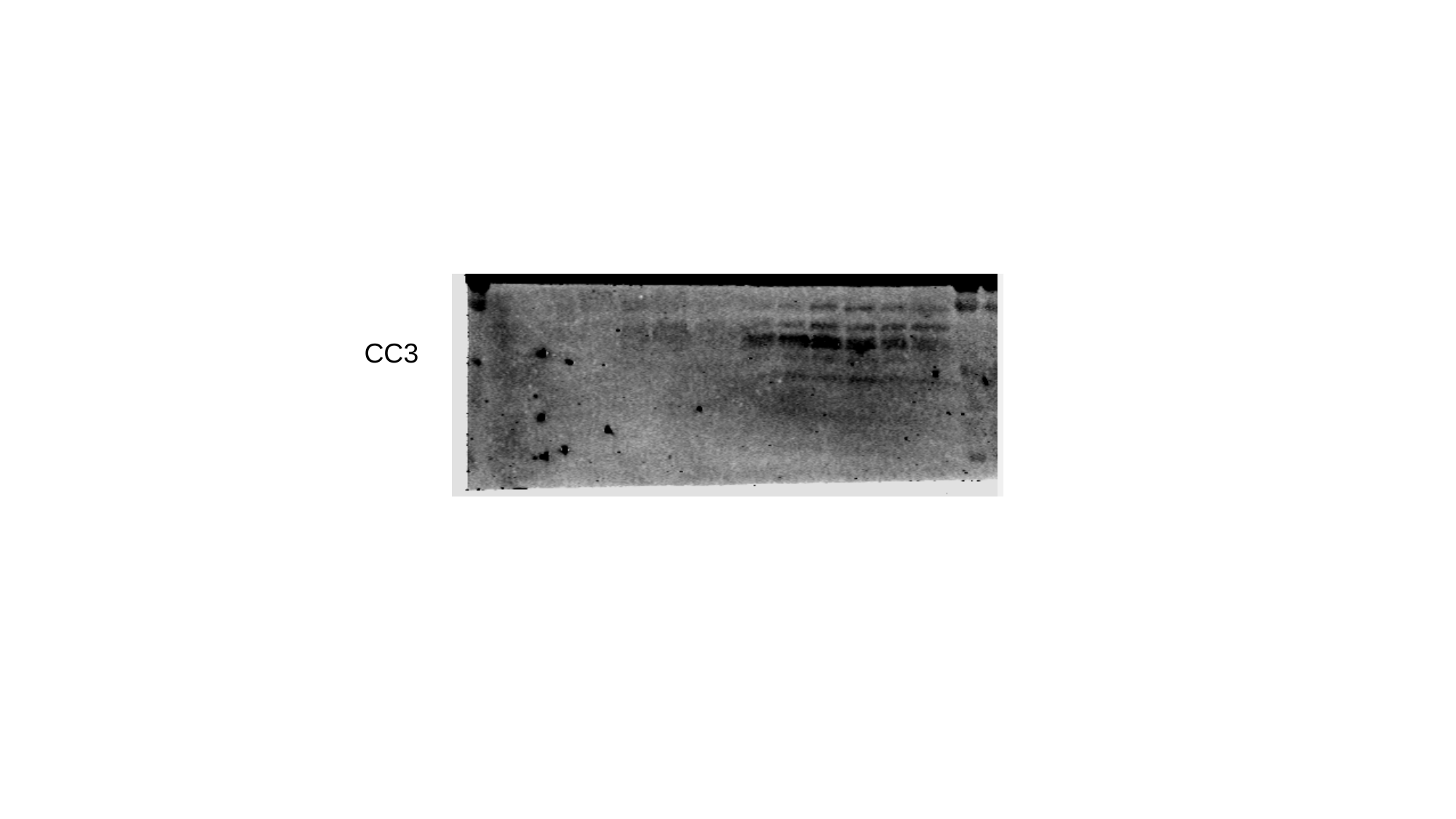

CC3

#### Slide 4
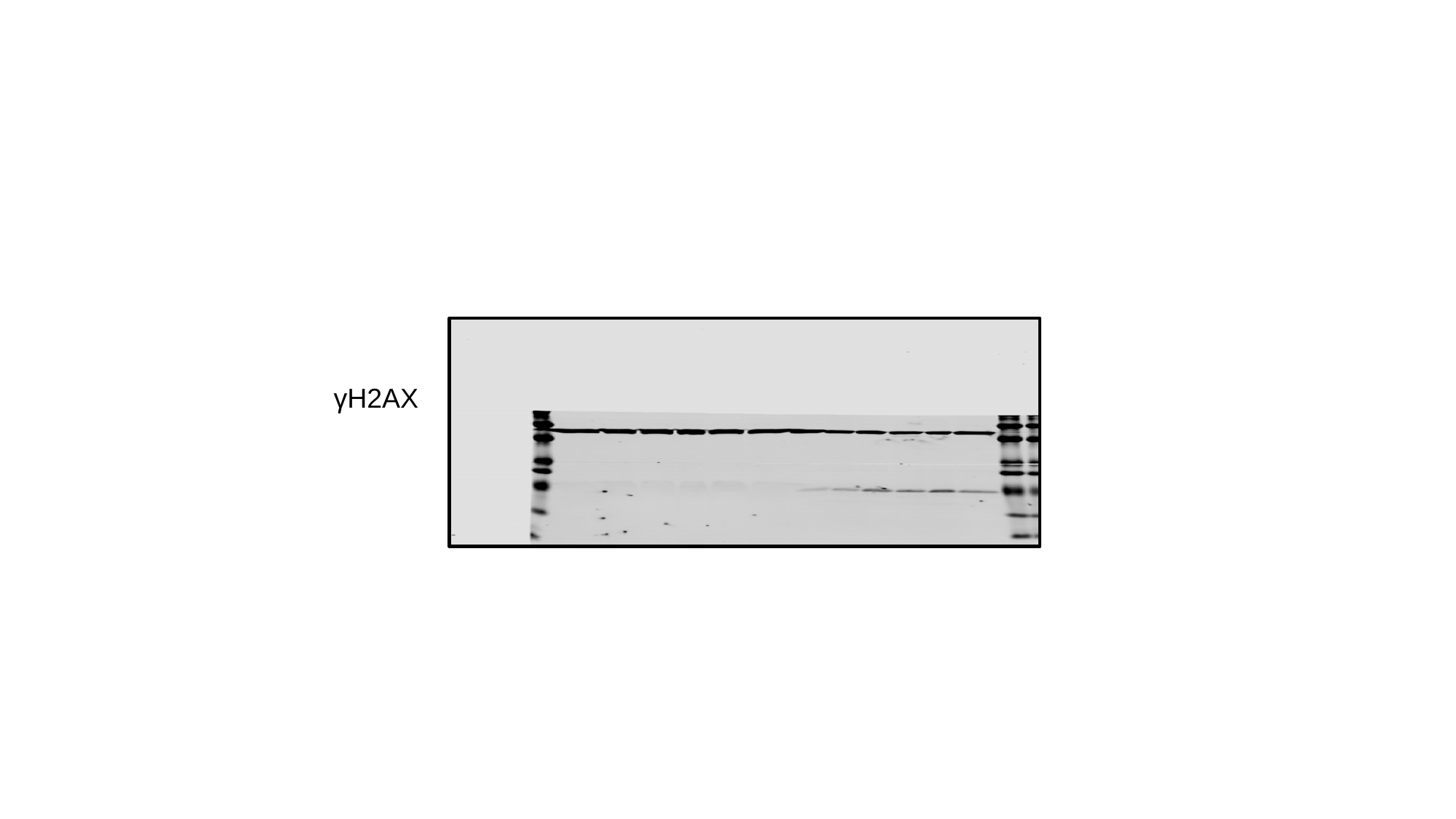

γH2AX

#### Slide 5
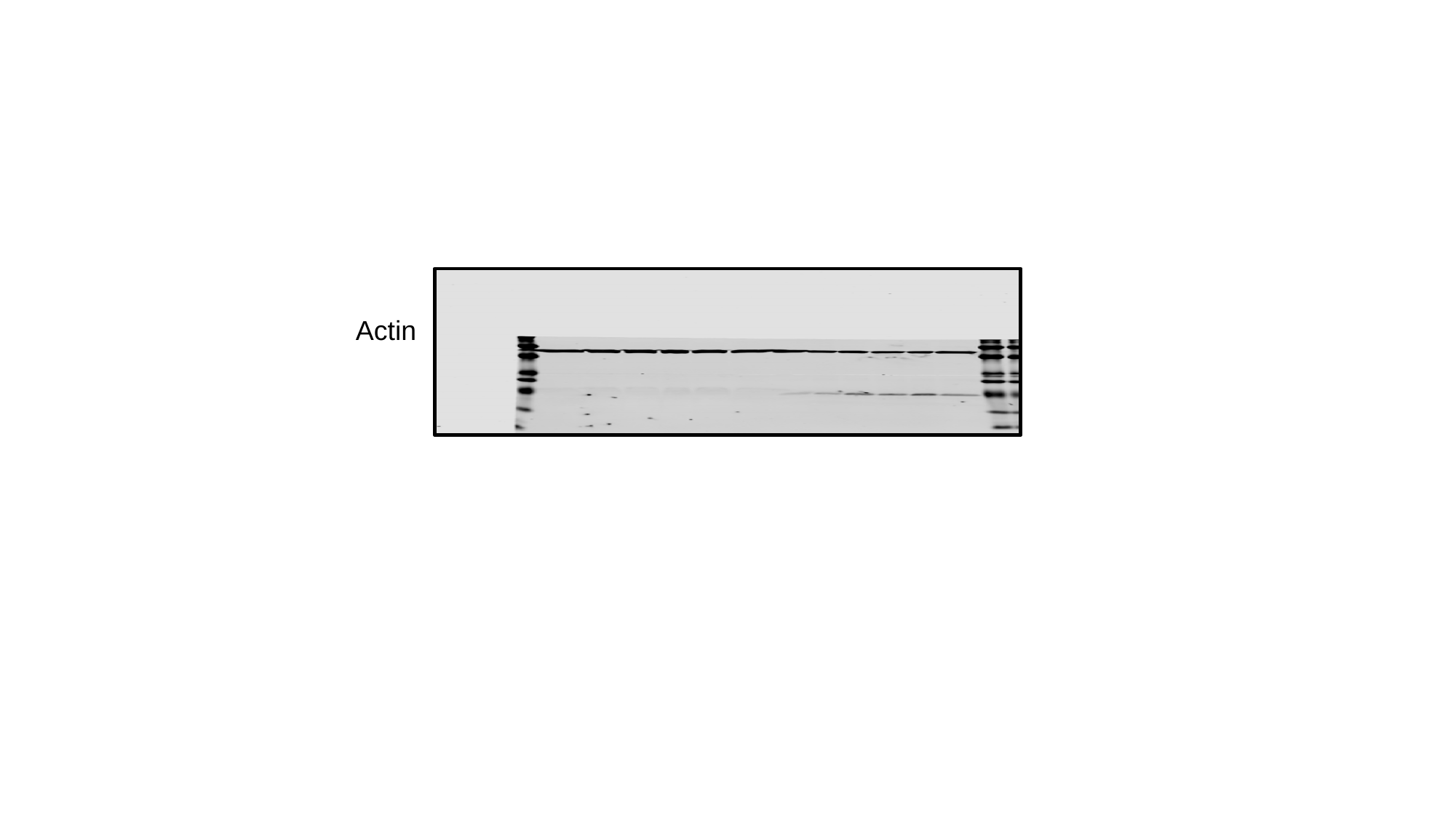

Actin

### 20190516_CellCycleSubG1.pptx

#### Slide 1
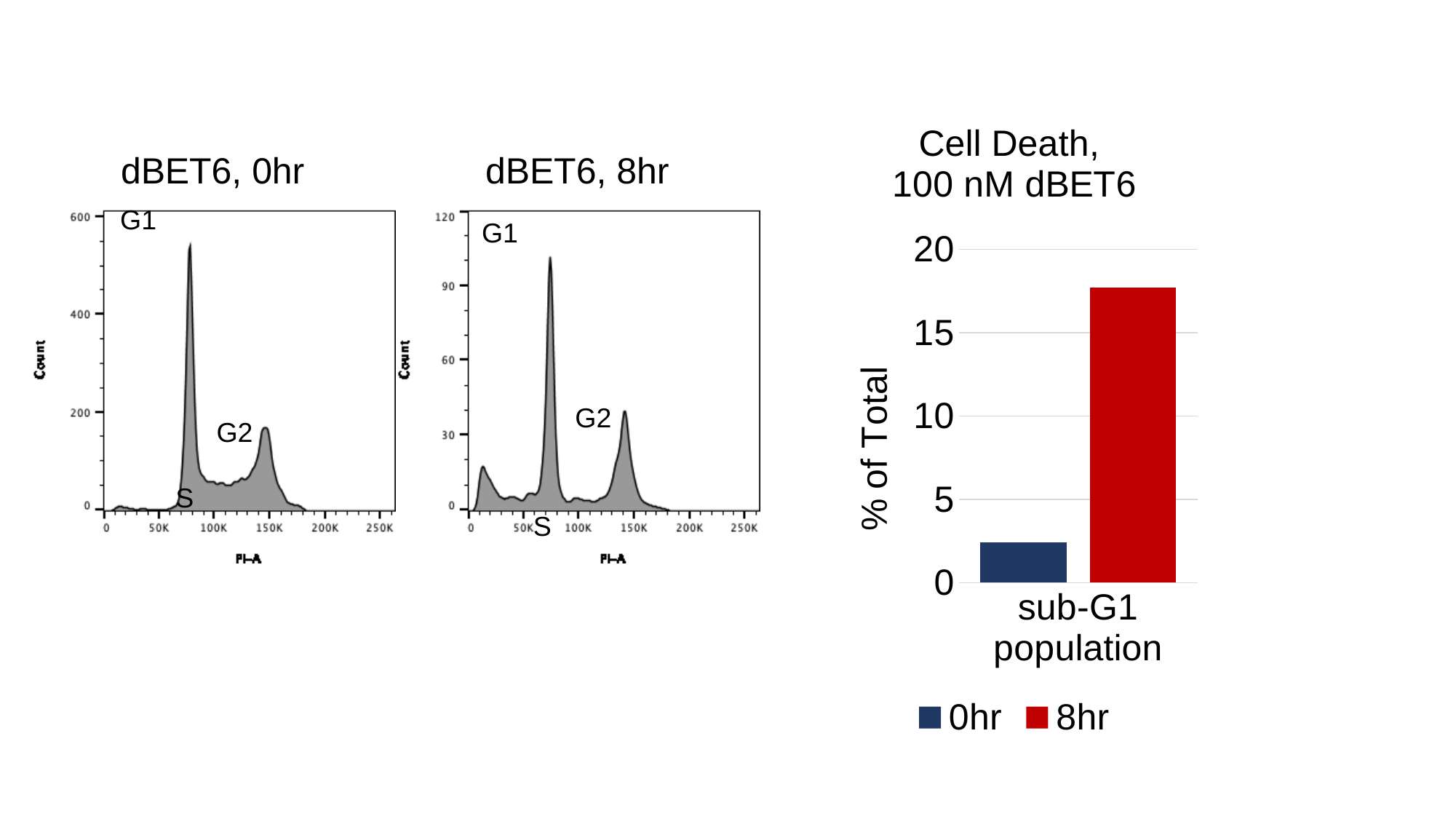

##### Chart: Cell Death,
100 nM dBET6
| Category |
|---|
| sub-G1 population | 2.4 | 17.7 |dBET6, 0hr
dBET6, 8hr
G1
G1
G2
G2
S
S

### 20200204_BETihibitors.pptx

#### Slide 1
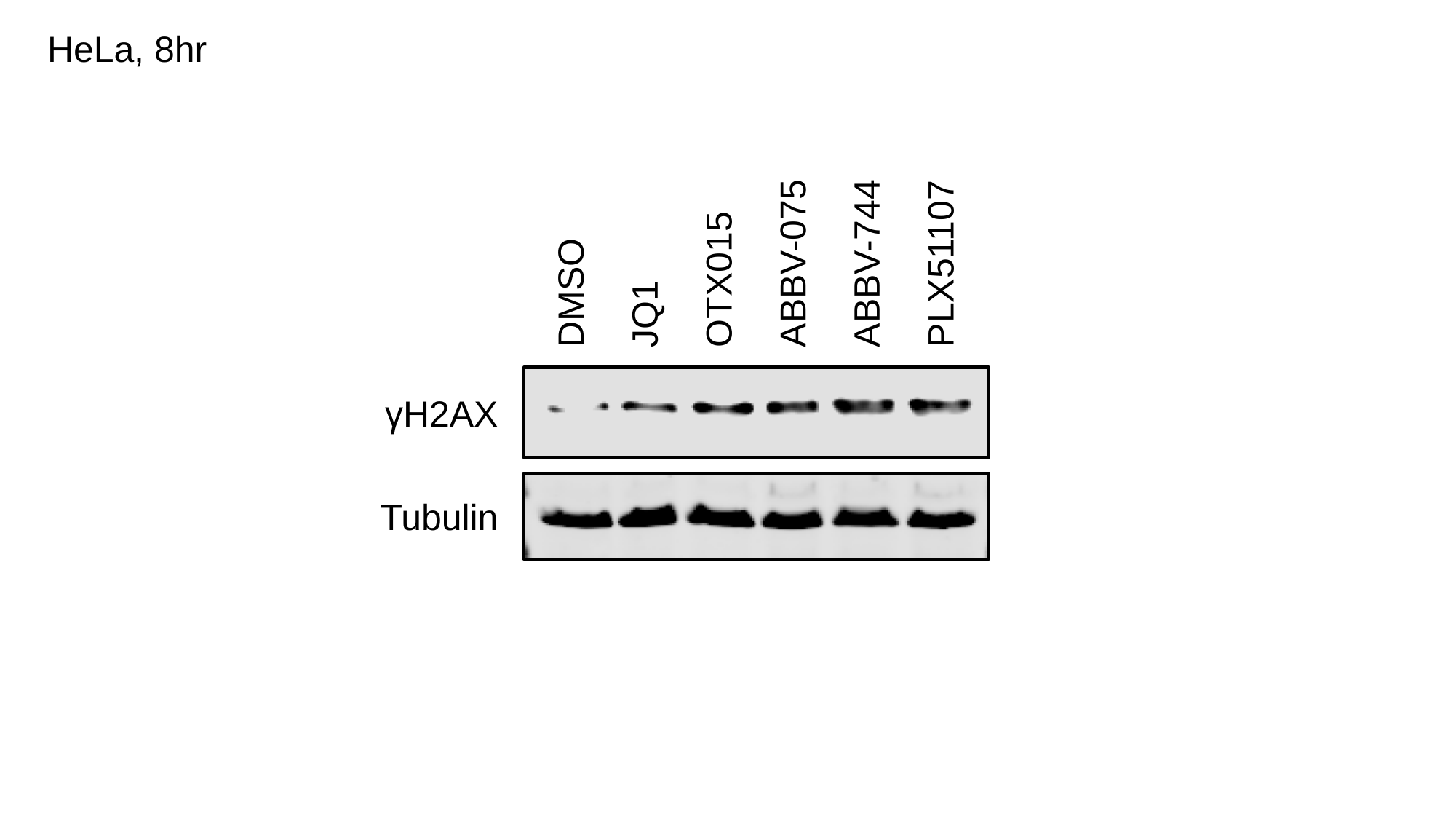

HeLa, 8hr
DMSO
JQ1
OTX015
ABBV-075
ABBV-744
PLX51107
γH2AX
Tubulin

#### Slide 2
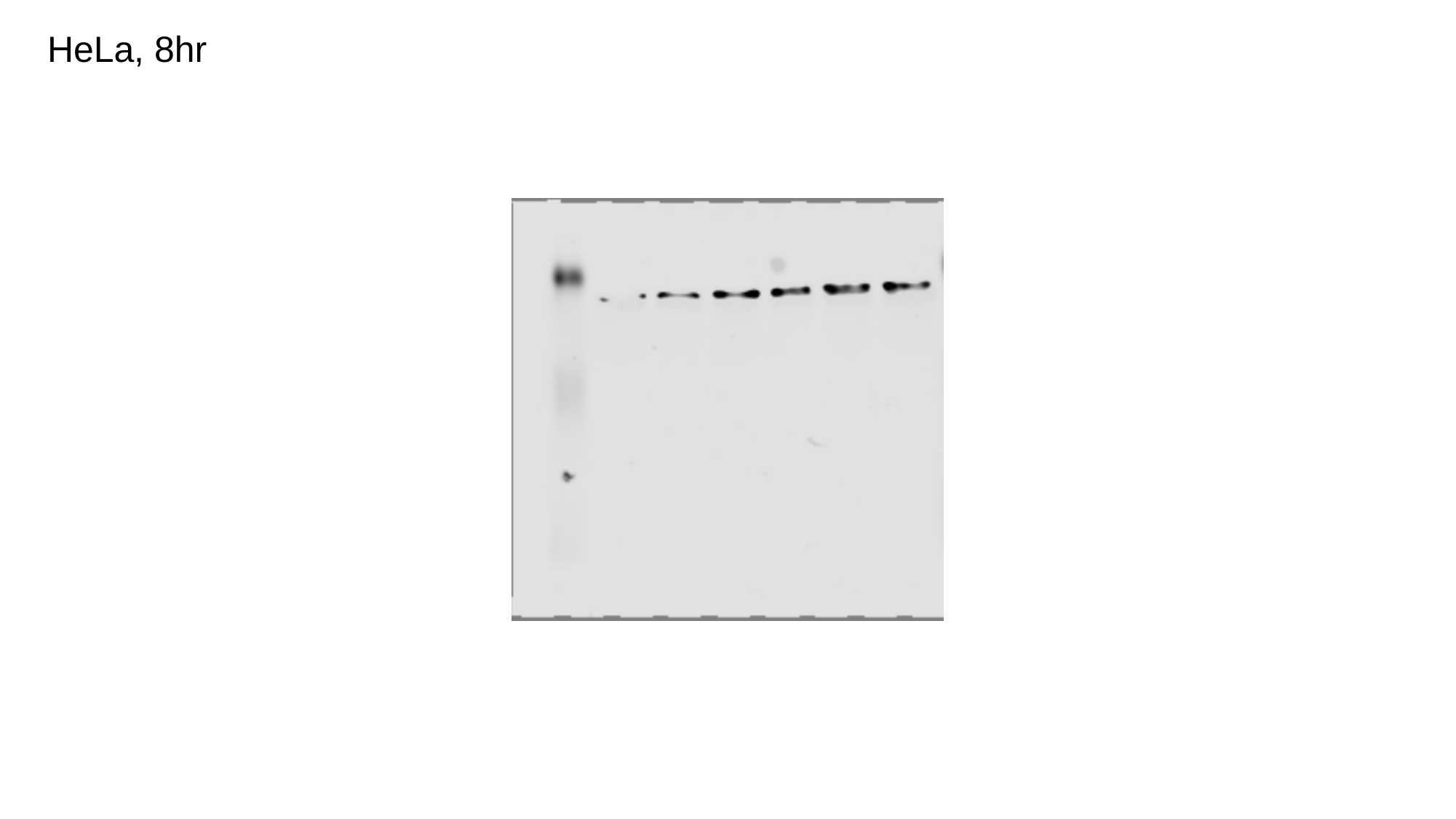

HeLa, 8hr

#### Slide 3
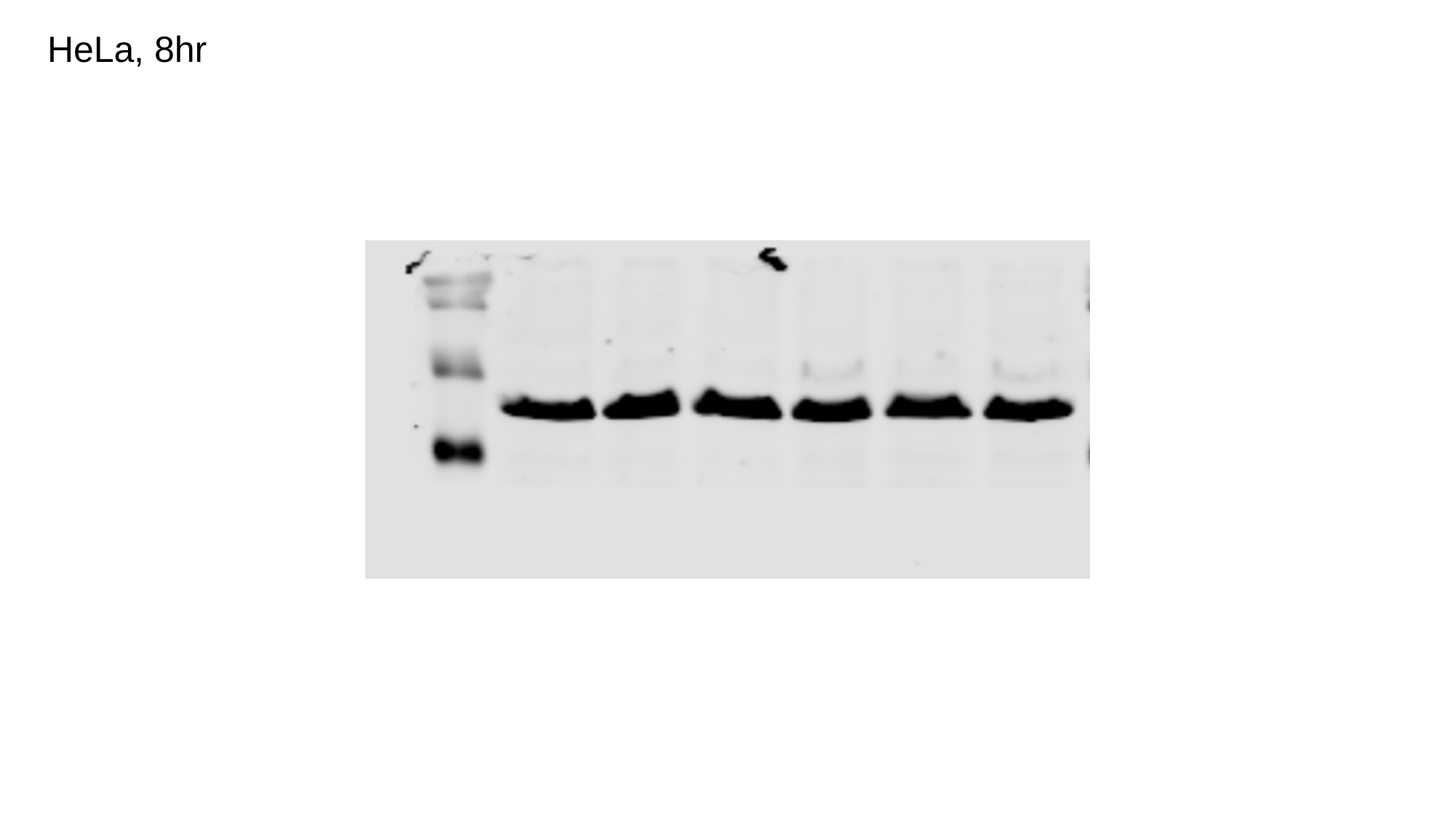

HeLa, 8hr
